# Drift, selection and convergence in the evolution of a nonribosomal peptide

**DOI:** 10.64898/2025.12.15.694437

**Authors:** Sebastian Kaiser, Niayesh Shahmohammadi, Trinetri S. M. Goel, Charles J. Buchanan, Sriram G. Garg, Daniela Vidaurre Barahona, Zhengyi Qian, Claire Lefèvre, Daniel Schindler, Yonggyun Kim, Helge B. Bode, Georg K. A. Hochberg

**Affiliations:** Department of Natural Products in Organismic Interactions, Max-Planck-Institute for Terrestrial Microbiology, 35043 Marburg, Germany; Evolutionary Biochemistry Group, Max-Planck-Institute for Terrestrial Microbiology, 35043 Marburg, Germany; School of Life Sciences and Engineering, Gyeongkuk National University, Andong 36729, Korea; Department of Structural and Molecular Biology, Division of Biosciences, University College London; London, UK; The Francis Crick Institute; London NW1 1AT, UK; Department of Biology, Phillips University Marburg, 35043 Marburg, Germany; Max Planck Institute for Terrestrial Microbiology, 35043 Marburg, Germany; Center for Molecular Biology of Heidelberg University (ZMBH), 69120 Heidelberg, Germany; Center for Synthetic Microbiology (SYNMIKRO), Phillips University Marburg, 35043 Marburg, Germany; Department of Chemistry, Phillips University Marburg, 35043 Marburg, Germany

## Abstract

Nonribosomal peptides play important roles in microbial ecology and medicine, for example as toxins and antibiotics such as penicillin^1^. Like ordinary proteins, nonribosomal peptides are subject to evolutionary change, which is driven by rampant recombinations of their synthetases^2^. This makes it difficult to study the evolutionary histories of nonribosomal peptides, because recombination is a serious challenge for phylogenetic inference^3^. Here we use recombination-aware phylogenetics and ancestral sequence reconstruction to retrace the evolution of GameXPeptides, nonribosomal peptides entomopathogenic bacteria use to kill their insect prey^4^. By untangling the complex histories of their synthetases, we discover dozens of new GameXPeptides and show striking patterns of structural convergence. In analogy to classic statistical test for natural selection acting on nucleotide mutations, we develop a similar test for whether selection acts to fix recombinations. We provide statistical and experimental evidence of both neutral genetic drift and natural selection in the diversification of GameXPeptides. GameXPeptides variants rapidly evolved highly insect specific toxicities, suggesting that their diversification may be driven by an evolutionary arms race. Our work suggests that small, fast evolving alteration to nonribosomal peptides can overcome host resistance on short timescales, with important implications for the evolution of resistance evading antibiotics.

---

Nonribosomal peptide synthetases (NRPS) are large, multi-domain assembly line enzymes that synthesize peptides which, in addition to the standard set of proteinogenic amino acids, can incorporate a variety of non-proteinogenic amino-acids and other moieties with accessory domains or separate enzymes^5^. NRPS consist of repeating <u>CAT</u>-tridomains, which usually incorporate one amino acid each. Within each tridomain, the <u>A</u>denylation (A)-domain selects the amino acid and then transfers it to the <u>T</u>hiolation (T)-domain, which covalently binds it to the synthetase. The <u>C</u>ondensation (C)-domain catalyzes peptide bond formation and, in the case of dual condensation/epimerization (C/E)-domains, includes switching the stereochemistry of the α-carbon of the amino acid from L to D configuration. The peptide is released by the <u>T</u>hio<u>e</u>sterase (TE)-domain, which comes last in the linear sequence of an NRPS. Some TE-domains liberate linear peptides, whereas others cyclize the peptide during release^5^ (Fig. 1a).

**Fig. 1.**
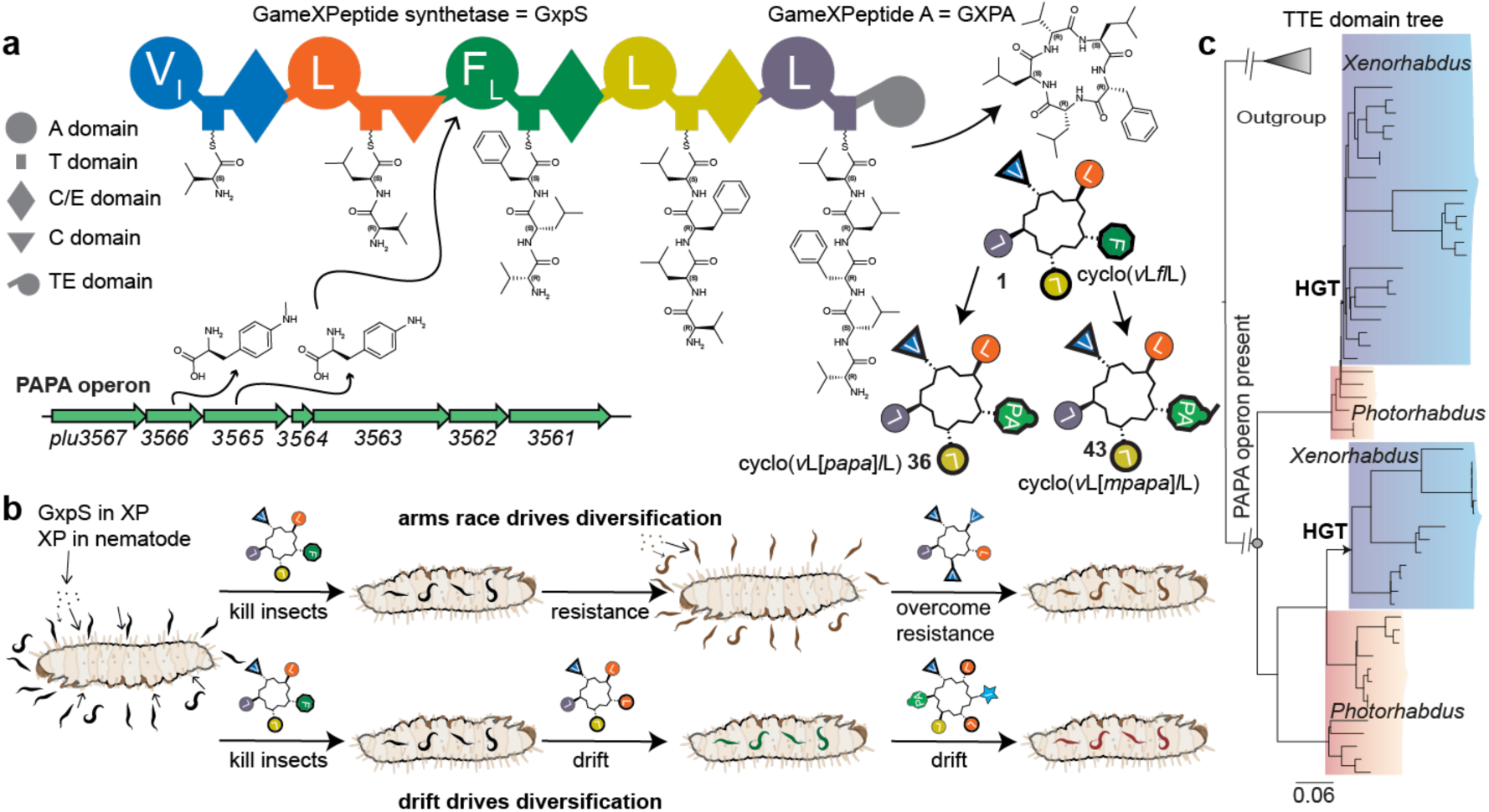
Evolutionary analysis of the NRPS GxpS. (**a**) The biosynthesis of **1** by GxpS. Four ATC-tridomains and one ATTE domain assemble different amino acids into a cyclic peptide. A-domains are labelled with the amino acid they select using single-letter amino acid code. Subscripts denote a side-specificity. The TE-domain releases the peptide and enables cyclization of the peptide. The PAPA operon is shown below. It produces PAPA and mPAPA, which can be accepted by GxpS’ A3-domain, shifting its major products to **36** and **42**. Amino acids in NRP are shown in one-letter code as well as specific symbol and color code according to the domains they originate from. D-amino acids in peptide sequences are shown in italics and in the cartoon peptide structure with bold black borders. *p*-aminophenylalanine and its methylated derivative is written as PAPA and mPAPA in brackets. (**b**) Hypothetical causes of GXP diversification. *Xenorhabdus* and *Photorhabdus* bacteria live symbiotically in nematodes and infest insects, using GXP to kill their insect prey. NRP diversification could be driven by an arms race, or neutral drift through a space of structurally different, but equally active GXP variants. (**c**) Phylogenetic tree of the TTE-domain of GxpS from *Xenorhabdus* (blue) and *Photorhabdus* (red). HGT: Horizontal gene transfer of GxpS from *Photorhabdus* to *Xenorhabdus*. Presence of the PAPA operon in the genome that contained the last common ancestor of GxpS is indicated.

The structure of NRPS-encoded natural products (NRP) evolves. This usually does not happen through point mutations that alter the active site geometries of the relevant domains. Instead, NRPS with altered specificities evolve mostly through recombination: e.g. one A-domain overwrites another A-domain, resulting in an altered NRP^2,6^. NRPS synthesize many compounds that act as antibiotics or as instruments of biological warfare, so one probable driver of NRP diversification is a molecular arms race between an antibiotic or toxin and its biological target^7–9^. As the organismal targets of NRP acquire resistance, NRP probably evolve to overcome that resistance. There is evidence for this in the evolution of glycopeptide^10^ and lipopeptide antibiotics, and this principle has been exploited for the discovery of novel classes of antibiotics^7,11–13^. In these cases, large structural changes result in new compound classes that confer escape from specific resistance mechanisms. It is unclear, however, if smaller, potentially more rapidly evolving changes within one compound class can similarly overcome resistance mechanisms. If they could, they would represent a promising reservoir for battling anti-microbial resistance.

Understanding the extent and evolutionary causes of NRP diversification is hampered by the fact that their structural diversity is mostly generated through recombinations, and only rarely through mutations, of NRPS genes^2^. Rampant recombinations make it difficult to infer phylogenetic histories of NRPS: each recombination event introduces a unique history, specific only to the stretch that underwent recombination. Conflicting histories within one alignment violate the assumptions of standard phylogenetics and can lead to phylogenies that do not accurately reflect the history of any part of the gene^3^. This makes it difficult to establish accurate ancestor-descendant relationships among NRPS and their products. For this reason, families of NRP are historically defined by structural similarity, not common descent. How often, in what order, and why particular NRP variants evolved is therefore largely unknown.

Here we tackle these challenges by studying the evolution of an NRP from the entomopathogenic bacteria *Xenorhabdus* and *Photorhabdus* (XP) which uses natural products to kill its insect prey^14,15^. The most conserved NRP among XP is the GameXPeptide (GXP) and its synthetase GxpS, which we study here. The canonical GxpS consists of five modules (specific for D-Val, L-Leu, D-Phe, D-Leu, L-Leu)^16–18^ (Fig. 1a). Due to the promiscuity of modules 1 and 3, the enzyme also produces more than one peptide. This family of peptides plausibly experiences both strong selection and neutral genetic drift: its function as an insecticide^4^ in a predator-prey system may lead to cycles of resistance and escape from resistance that diversify the NRP (Fig. 1b). Similarly, variants of GXP could evolve to expand the range of susceptible prey. At the same time, this system probably experiences significant drift: each nematode of the genera *Steinernema* and *Heterorhabditis* is initially colonized by a small number of XP bacteria, which then grow up to around 100 cells within each nematode. This bottlenecking in the XP life-cycle could lead to high rates of drift in the exact structure of GXP^19,20^, potentially producing many structurally different but equally active GXP variants (Fig. 1b).

Here we develop an explicit phylogenetic framework for understanding the evolution of NRPS and use it to discover a vast diversity of novel GXP variants. We retrace how these products diversified in history and provide evidence that both natural selection and neutral drift are important forces in the diversification of NRP families. Finally, we show that GXP variants show strong differences in their effectiveness as toxins against specific insects.

## Results

### Gene-level history of GxpS

Our first goal was to discover if there is structural variation among GXP. To do this, we first needed a phylogeny of GxpS that defines what counts as a GXP based on common descent and orthology of their synthetases, instead of structural similarity of the NRP. This is made difficult by the complex and hierarchical histories of NRPS: whole NRPS genes are subject to duplication events, gene fusions and/or horizontal transfer between bacteria^21^. In addition, individual domains of NRPS also duplicate to give rise to different kinds of A– and C-domains. Recombination between different modules adds a final layer of complexity and can also occur between domains from different NRPS. Each of these processes introduces different phylogenetic incongruences between the species these genes evolve in and the genes themselves, which complicates orthology assignments.

In order to untangle this complex hierarchy, we used the TE-domain as our phylogenetic yardstick (Fig. 1c and Extended Data Fig. 1a). TE-domains occur usually just once in NRPS, as is the case in GxpS. Further, recombination aware methods could not identify any recombinations in TE-domains from GxpS (Supp. Data S1). This means we can use the TE-domain in analogy to a conventional gene tree, which reports on whole-NRPS duplications and transfers, but is unaffected by individual domain duplications and recombinations. We inferred a tree of TTE-domains (T5 and TE domain) of 54 putative GxpS orthologs. We added the T-domain to the TTE-domain because it added more sequence information and lead to a better congruence to the species and individual domain trees overall. We rooted this tree with TTE-domains of a related NRPS called KolS^22^. All putative GxpS sequences formed a monophyletic group. In addition, a four modular NRPS called Xenotetrapeptide synthetase (XtpS)^23^, which was previously suspected to be a close relative of GxpS^14^, nests clearly inside the GxpS clade. So did a previously unknown six-modular NRPS. We then inferred a species tree of XP (Extended Data Fig. 1b) and compared it to our TTE-domain gene tree. This tree was incongruent with the TTE-domain tree, implying that GxpS has experienced whole NRPS horizontal gene transfer (HGT). Gene-tree-species tree reconciliation^24^ of the TTE and species tree indicated that GxpS first arose in *Photorhabdus* and was subsequently transferred to *Xenorhabdus* twice (Fig. 1c; Extended Data Fig. 1, Supplementary Fig. 1).

Like other NRPS, GxpS can incorporate non-proteinogenic amino acids. Specifically, *P. laumondii* TT01 harbors an operon that synthesizes the non-proteinogenic phenylalanine analog *p*-aminophenylalanine (PAPA) and its methylated derivative (mPAPA)^16^. These amino acids can be also incorporated by GxpS_TT01_’s A3-domain (Fig. 1A). When available, PAPA and mPAPA-incorporating derivatives become GxpS’ main natural products. By analyzing the phylogenies of genes in the PAPA operon, we deduced that it was present in the last common ancestor (LCA) of XP and therefore present by the time GxpS evolved in an ancestral *Photorhabdus* (Fig. 1c; Extended Data Figs. 2 and 3; Supplementary Fig. 2). The operon was lost twice in different groups of *Xenorhabdus*, and then reacquired in *Xenorhabdus doucetiae* via HGT. In this specific organism, however, the operon is littered with mobile genetic elements and therefore likely inactive, which we confirmed experimentally (Supplementary Figs. 3 and 4).

### Domain-level phylogenetic incongruence reveals rapid NRP diversification by recombination

In the absence of recombination, the phylogeny of all other domains should be congruent with the TTE-domain phylogeny, because these domains would always have been inherited together (vertically and horizontally) since the LCA of all GxpS. In our analysis below, we searched for deviations from this reference phylogeny in the trees of GxpS’ other domains. Our purpose was twofold: First to discover recombination events that led to the emergence of new NRP. And second, to untangle the complex effects these recombinations had on GxpS’ phylogeny to finally derive a fully resolved history of how, when, and why new NRP evolved.

We first inferred a phylogeny of all GxpS A-domains. On this tree, the five A-domains of each GxpS form separate clades (Extended Data Fig. 4a). Each clade descends from one of five ancestral A-domains that existed in the LCA of all GxpS. There are several instances of A-domains that are not located in the clade that we would expect them to be in based on their position in the linear sequence of GxpS. For example, instead of grouping with other A3 domains, the A3 domain of the GxpS of *Xenorhabdus* sp. PB62.4 is sister to its own A2 domain (Fig. 2a). This implies that the A2 domain overwrote the A3 domain in an ancestor of *Xenorhabdus* sp. PB62.4. A sequence similarity analysis revealed that the entire A-domain was overwritten (Fig. 2b and Extended Data Table 1, No. 33). The A3 domain usually incorporates phenylalanine, whereas the A2 domain usually incorporates leucine. We therefore expected this recombination altered the structure of the NRP. Through heterologous expression of this GxpS and HPLC-MS analysis, we were able to confirm that this synthetase solely incorporates leucine instead of phenylalanine in the third position (Fig. 2c).

**Fig. 2:**
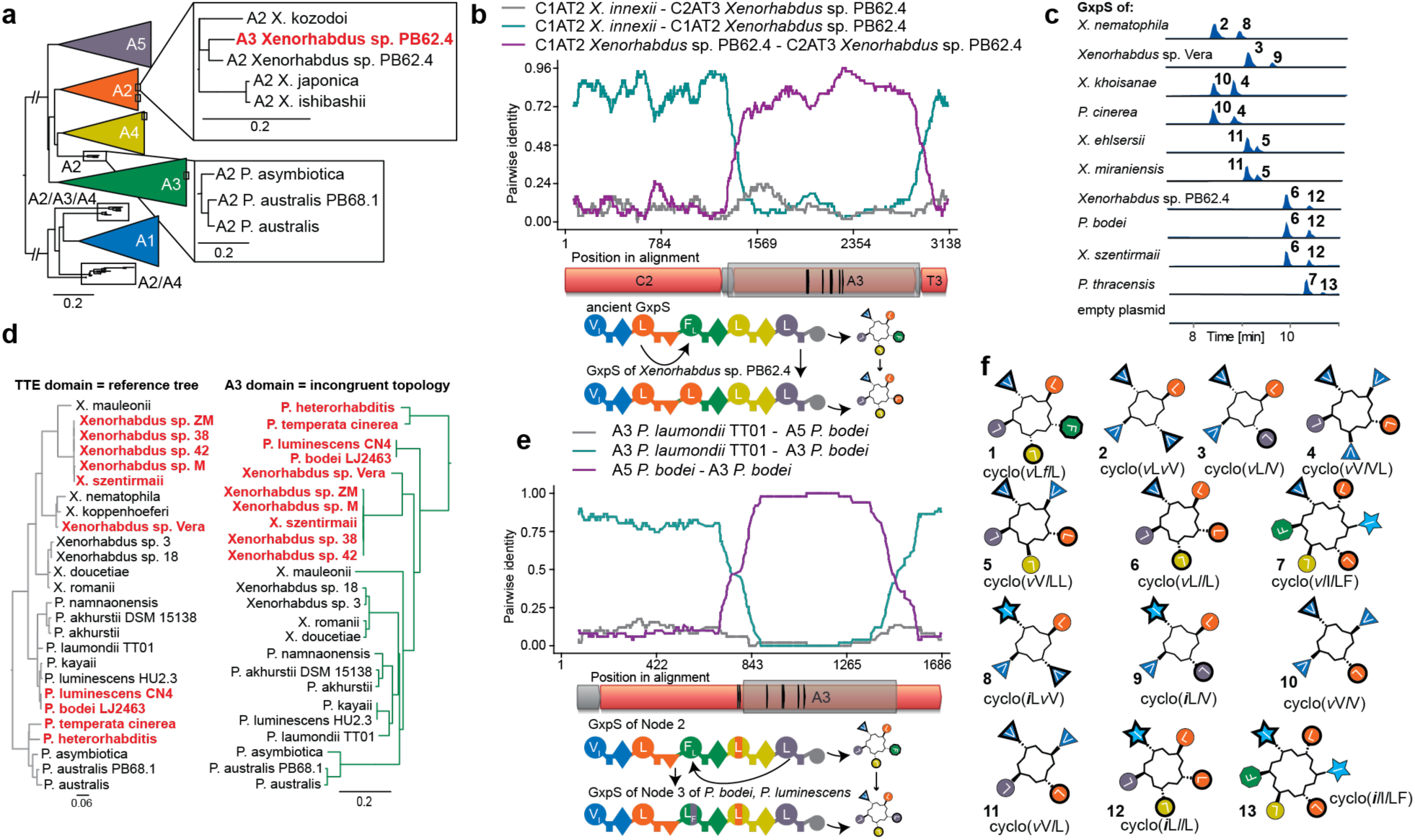
Phylogenetic identification of new GameXPeptides. (**a**) Obvious recombination events on the A-domain tree of GxpS. All first, second, third, fourth and fifth A-domains form separate monophyletic clades. The A3 domain of the GxpS of *Xenorhabdus* sp. PB62.4 is sister to its own A2 domain. The A2 domains of the GxpS from *P. asymbiotica* sit outside of the A2 domain and beside the A4 domain clade. (**b**) Sequence similarity analysis of the recombination event in the third module of GxpS from *Xenorhabdus* sp. PB62.4. Sequence similarity analysis for variable positions on the third module of the GxpS from *Xenorhabdus* sp. PB62.4 and an unrecombined second CAT-tridomain of a GxpS from another *Xenorhabdus* (cyan); its own second module (purple), with which it recombined; and a comparison between an unrecombined second module and the third module from *Xenorhabdus* sp. PB62.4 (grey). Here the entire A3 domain was replaced by its own A2 domain. The module structure is shown below, with specificity encoding sites^30^ in the A-domain indicated as black stripes. Bottom: Cartoon representation of the recombination event, with A-and C-domains colored as in Fig. 1a. (**c**) HPLC-MS measurements of culture extracts from *E. coli* expressing newly predicted GxpS. Extracted ion chromatograms of the first two main peptides are shown. (**d**) Comparison of the TTE-domain tree with the third A-domain clade. The third A-domain clade is highly incongruent with the TTE-domain tree. (**e**) Sequence similarity analysis of the recombination event of the third A-domain of GxpS from *P. bodei*. Labelling scheme as in B. Cyan compares the recombined sequence with an unrecombined relative, purple compares the recombined sequence with a descendant of the recombination donor, and grey compares recombination donor to unrecombined relative. Here a fifth A-domain has recombined into the middle of the third A-domain of the same synthetase, leading to a shift from main peptide **1** resulting in **6** becoming the main peptide. Specificity-encoding sites^30^ are annotated in the A-domain in black strokes. (**f**) Cartoon structures of all new GXP predicted via phylogenetics and confirmed through HPLC-MS measurements.

In total we found 15 instances in which A-domains appear to have recombined, based on their anomalous position in clades of A-domains that do not match their position in the linear sequence of GxpS. We observed similar patterns for the TC-domain phylogeny: Some TC-domains sit inside the^L^C_L_ clade but outside the GxpS TC-domain clade. This particular pattern implies that they likely recombined with C-domains from other NRPS. The fourth TC-domain of the canonical GxpS is a dual C/E-domain (Fig. 1a), but we find two cases where C4 domains of other GxpS sit in the^L^C_L_ clade (Extended Data Fig. 4b). This suggests that recombination has changed these C-domains from dual C/E-domains towards^L^C_L_-domains.

We next used more detailed congruence analysis between our A-domain tree and the TTE-domain phylogeny to identify more recombinations. We noticed that the A3-domain’s phylogeny was very incongruent with the TTE-domain tree (Fig. 2d). This suggested there is conflicting phylogenetic signal from the unrecombined and recombined parts within the A3 domain resulting in a topology that reflects the history of neither unrecombined nor the sequence that underwent recombination.

To test this idea, we applied a phylogenetic Hidden Markov Model analysis^25^ to a separate alignment containing A3 and A5 domains as an outgroup. Briefly, this approach infers two topologies that together best describe the dataset. This revealed that this alignment is indeed described by at least two different phylogenetic histories (Supplementary Fig. 5): one that is mostly congruent with the TTE-domain phylogeny and therefore represents the unrecombined sequence (Supplementary Fig. 6), and another, spanning from the middle until almost the end of the A-domain, that underwent recombination with leucine-incorporating A-domains (Supplementary Fig. 7). The phylogeny of only the recombined stretch clearly reveals that some A3 domains position in the leucine-incorporating A5 clade. Using this approach, we iteratively disentangled a complex history of small recombinations in the A3 clade, each time using sequence similarity analysis to identify which segments recombined, and then phylogenies of those segments to identify the donor clade (Fig. 2e). In some cases, this was complicated by the fact that recombinations can occur in segments that had already recombined previously. Each recombination event we solved brought a portion of the A3 clade back into congruence with the TTE phylogeny when the recombined stretch was excluded. Where parts of the A3 phylogeny then remained incongruent, we searched for more recombination events (Supplementary Fig. 8 and Extended Data Table 1, No. 24 and 34). Overall, we discovered five additional NRP-altering recombination events that involved segments too small to move them into entirely different A-domain clades on our full A-domain phylogeny (Extended Data Table 1, No. 2, 5, 24, 34, 42).

We used similar approaches to disentangle how shorter and longer variants of GxpS evolved on our tree. The four-modular XtpS evolved via recombination with a foreign NRPS, whose A-domain recombined with the flanking regions of the GxpS’ A4 and A5 domain that led to the excision of A4TC4A5 domains (Extended Data Table 1, No. 7). The origin of the six-modular Gamehexpeptide synthetase (HexS) involved a TCA deletion, an ATC duplication as well as an ATC insertion (Extended Data Table 1, No. 11-19). Overall, our congruence and sequence similarity methods allowed us to identify 9 unique variants of GxpS that we predicted to have a product spectrum different from the canonical ortholog (Extended Data Fig. 5). Based on the predicted specificities and side-products of these NRPS, we predicted at least 12 new main products and dozens of new side products. We then tested these predictions by heterologously expressing representatives of all synthetases we predicted to produce variant NRP in *E. coli* (Fig. 2c and f). We analyzed their products using HPLC-MS and co-injection of chemically synthesized versions of the predicted peptide. In addition, we performed feeding experiments with isotopically labelled amino acids to confirm the stereochemistry of newly discovered GXP. In these experiments we detected a total diversity of 49 compounds (Supplementary Tables 1 and 2), virtually all of which were consistent with our phylogenetic predictions (Supplementary Figs. 9-46). We also performed PAPA and mPAPA feeding experiments for different types of GxpS and found that all synthetases that can incorporate phenylalanine can also incorporate PAPA and mPAPA. In almost all cases, the PAPA-or mPAPA-incorporating version was the dominant NRP under our feeding conditions. For the *Xenorhabdus* species that do not have the PAPA operon but still have a phenylalanine-incorporating A3 domain, GXPA is the main product (Supplementary Figs. 47-50).

The only synthetases whose experimentally determined products defied our phylogenetic predictions were a few five-modular NRPS that produced cyclic tetrapeptides as their main products (Fig. 2c and f). We reasoned that this may result from module-skipping, a well-known phenomenon to produce smaller peptides^26^, though what makes this case unusual is the fact that cyclic tetrapeptides are extremely rare in nature^27^. In the absence of other obvious changes, we suspected that these synthetases prematurely cyclize their product using an^L^C_L_-domain, which replaced their ancestral C4 dual C/E-domain via recombination. To test this theory, we introduced loss-of-function mutations in both the fifth T-domain and the TE-domain. This still resulted in the production of cyclic tetrapeptides, consistent with cyclization by the recombined C4-domain. To further confirm that the C4 domain is responsible, we replaced this domain with the^L^C_L_-C2 domain of the same synthetase. This completely repressed the formation of cyclic tetrapeptides and resulted in only cyclic pentapeptides, confirming that the ability to produce cyclic tetrapeptides is encoded in the recombined C4-domain of these synthetases (Extended Data Fig. 6).

Overall, our phylogenetic and biochemical analyses therefore revealed a substantial and thus-far unnoticed peptide diversity in the family of GxpS within only two genera of bacteria (Fig. 2f and Supplementary Tables 1 and 2).

### Resurrection of the last common ancestor of all GxpS reveals highly convergent NRP evolution

Having established the extent of NRP diversity among GxpS, we were now in a position to unravel how this diversity evolved. To do this, we mapped the structure of the main product of every synthetase we characterized onto the TTE-domain phylogeny (Fig. 3a). This revealed that multiple peptides must have evolved several times in parallel, because we found the same structures in distant parts of our tree. But the distribution of different NRP on our TTE-domain tree made it difficult to confidently assign which GXP is ancestral, and which GXP evolved multiple times from this ancestral state.

**Fig. 3:**
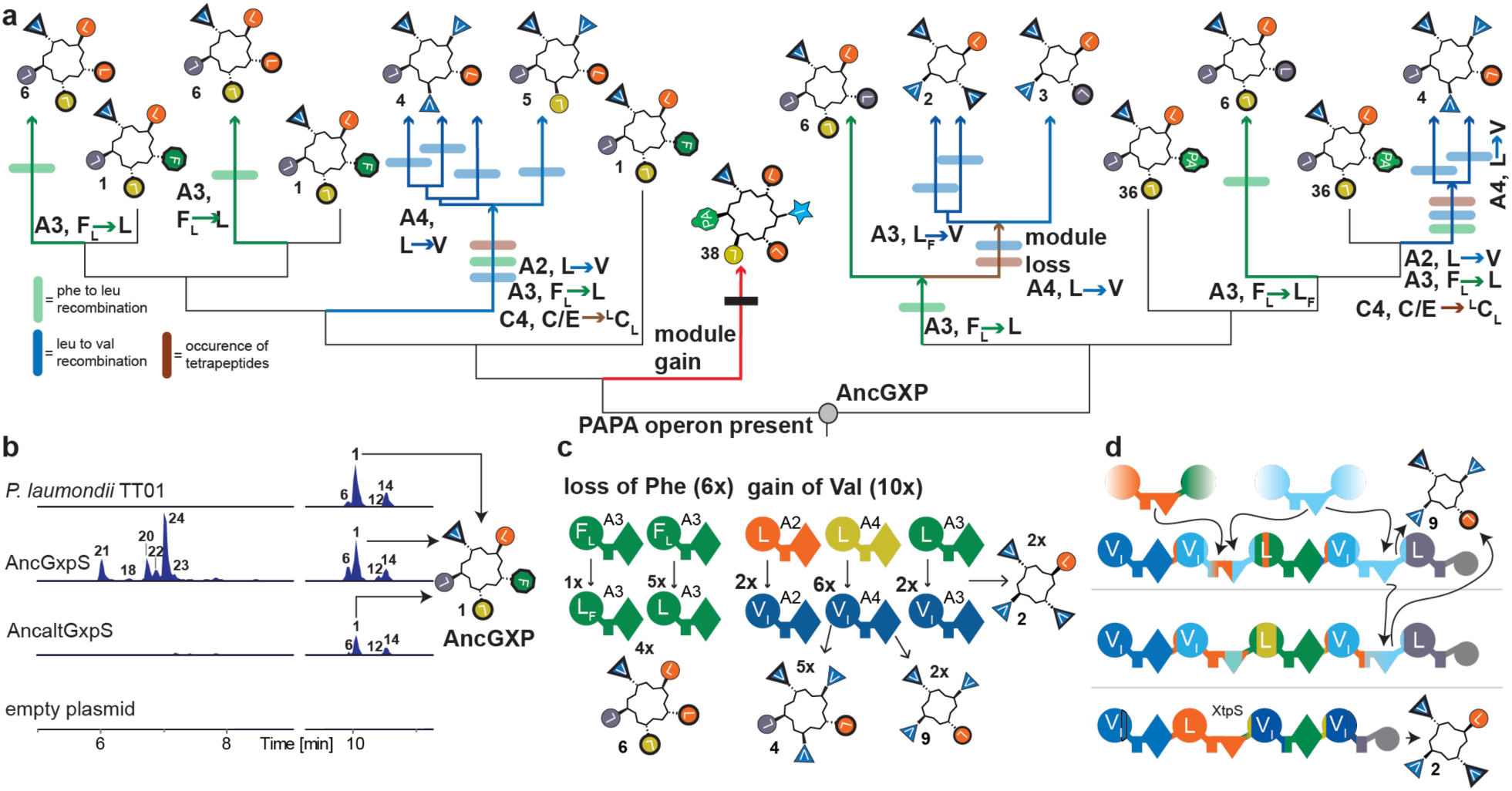
Convergent evolution of novel GXP from an ancestral cyclic pentapeptide. (**a**) A simplified TTE-domain tree with natural products mapped on the tips. Specificity changing recombinations are marked along the relevant branches. Green arrows and strokes indicate recombinations event that changed Phe to Leu on position 3 changing GXPA **1** to GXPC **6** on some branches. Blue arrows and blue strokes indicate Leu to Val recombinations which create GXP with more than one valine. Brown arrows and strokes indicate the emergence of tetrapeptides. Full tree is shown in Extended Data Fig. 8. (**b**) HPLC-MS measurements of AncGxpS and an alternative reconstruction of the same protein. TT01’s GxpS is shown for reference. Peptides 19-24 are linear tri– and tetrapeptides derived from GxpS. (**c**) Convergent evolution of GxpS. Convergent recombination events of particular domains are shown along the top with convergent origins of particular peptides along the bottom. (**d**) Convergent evolution of tetrapeptides. One GxpS type acquired a specific C-domain in its 4^th^ module that enables premature peptide release and cyclization (Extended Data Fig. 6). The second GxpS type acquired the premature releasing C-domain from another GxpS horizontally, making it also produce cyclic tetrapeptides. The third GxpS type lost one full module and also produces cyclic tetrapeptides.

To solve this problem, we used ancestral sequence reconstruction to resurrect the LCA of all GxpS on our tree. Our aim was to resurrect nodes at the base of each A– and TC– and TTE-clade that correspond to the last common ancestor of GxpS on our TTE-domain phylogeny (AncGxpS, Fig. 1c, Extended Data Fig. 4). This was possible for all domains except for A3, where we first had to exclude several recombined sequences to bring the A3 tree into congruence with the TTE-domain phylogeny (Extended Data Fig. 7). We then assembled our separately reconstructed domains back into a complete synthetase, expressed this gene in *E. coli* and used HPLC-MS analysis to determine which product it formed. This revealed that the ancestral product was **1** as the cyclic main product (cyclo(D-Val, L-Leu, D-Phe, D-Leu and L-Leu)) (Fig. 3b). Our reconciliations of the gene histories of the PAPA operon indicated that it was present in the ancestral genome that would have contained AncGxpS (Extended Data Fig. 1 and 3). Consistent with this we found that AncGxpS could incorporate PAPA and mPAPA (Supplementary Figs. 47-50). We verified that these inferences are robust to uncertainty by reconstructing a much less likely version of the same ancestor (AncaltGxpS, see methods) (Supplementary Fig. 51), which differed from AncGxpS at 244 amino acid positions. This confirmed that the ancestral main product of the GxpS family of synthetases was **1** (Fig. 3B).

Having defined the starting point of GxpS’ diversification, we could now polarize in which order the natural products on our tree evolved (Fig. 3a). This revealed a remarkable degree of convergence in the evolution of novel GXP. The ancestral phenylalanine at position three was independently replaced with leucines via recombination in the A3 domain six times. This convergently yielded four NRPS producing **6** as their main peptide (Fig. 3c). Moreover, recombinations introduced A-domains specific for valine 10 times independently into different positions of GxpS. This resulted in five convergent origins of **4**, a GXP with the structure of cyclo(D-Val, L-Val, D-Leu, L-Val and L-Leu) (Fig. 3c). A last and very dramatic example of convergence is the evolution of tetrapeptide-producing synthetases. One instance is the four-modular GxpS, XtpS^23^ (Fig. 3d). The second mechanism is the evolution of five-modular synthetases producing cyclic tetrapeptides we described above. A third instance is another five-modular synthetase, which acquired the early-terminating^L^C_L_ in its fourth position from the other early terminating synthetases via a recombination. Convergence is often caused by strong selection for beneficial functions^28^. Therefore, we can assume that some of the peptide diversity we discovered was fixed by selection.

### Drift and selection fix NRPS recombinations

We next wanted to further investigate what the evolutionary causes of the GXP diversity is. In principle, it could be caused by natural selection for more active NRP, which overcome an acquired resistance or allow new hosts to be killed. Alternatively, much of this diversity may stem from recombinations that are fixed by neutral drift, resulting in structurally diverse, but equally active peptides (Fig. 1b). We reasoned that there might be a particular kind of recombination that could serve as a proxy for the possibility of drift in GxpS. These would be recombinations between A-domains that incorporate the same amino acid, or C-domains that enforce the same stereochemistry. We refer to such recombinations as homotypic and they are synonymous with respect to the NRP – they do not change the NRP structure and might therefore be invisible to selection to a first approximation. This is analogous to the assumption that synonymous substitutions in a codon triplet can serve as an approximation of the neutral DNA substitution rate in genes^29^. Heterotypic recombinations between A-domains of different specificities, on the other hand, can change the specificity (i.e. they can be non-synonymous on the level of the peptide structure), if they cover a particular portion of the A-domain that encodes specificity^30^ (Fig. 4a). They can also be synonymous if they do not cover this region.

**Fig. 4:**
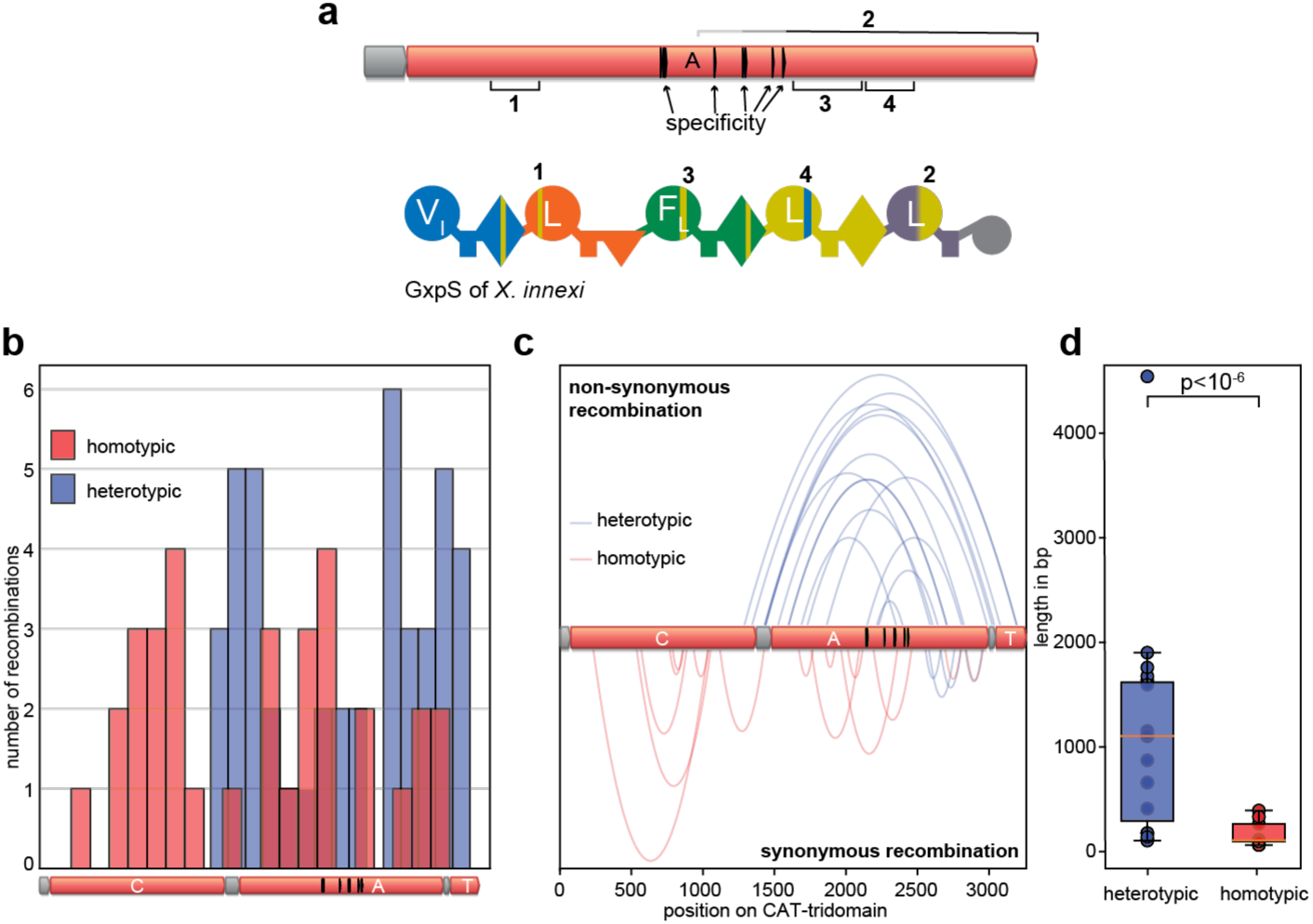
Analysis of homotypic vs. heterotypic recombinations. (**a**) An illustration of a synthetase that underwent several homo– and heterotypic recombinations. Specificity-encoding sites are annotated in the A domain in black strokes^30^. The left recombination border of recombination 2 is not clear and therefore annotated in gray. (**b**) Histogram of recombination breakpoints for homotypic and heterotypic recombinations along the CAT-tridomain. Specificity-encoding sites are annotated in the A-domain in black strokes^30^. (**c**) Parabolas connect the left and right breakpoints of recombinations in our dataset along the CAT-tridomain. Specificity-encoding sites are annotated in the A-domain in black strokes^30^. Only recombinations for which we could identify both breakpoints were used in this analysis. (**d**) Boxplot analysis of the length of homotypic and heterotypic recombination. p-value was calculated using a Wilcoxon rank-sums test.

The A-domain tree of GxpS contains one obvious synonymous recombination (Fig. 2a). The second A-domain, presumably specific for leucine, of *Photorhabdus asymbiotica* sits close to its own fourth A-domain that should also encode leucine. This pattern was caused by a recombination event, in which *P. asymbiotica’*s A2 domain received a fragment from the A4 domain of the same synthetase (Extended Data Table 1, No. 48). Encouraged by this discovery, we looked for signs of more synonymous recombinations on our trees. In this case, we relied mostly on local sequence similarity analyses^31^ and tree incongruence, which guided us to a large number of often very small synonymous recombinations. Overall, we found 27 synonymous recombinations, compared to 23 non-synonymous ones (Extended Data Table 1; Extended Data Fig. 8). These data imply that drift plays a significant role in the evolution of NRPS.

We next sought to use homotypic recombinations as a proxy for the neutral recombination process to test for signs of selection in heterotypic recombinations. To do this, we compared homo– and heterotypic recombinations. Homotypic recombinations occur all over the CAT-tridomain and are more frequent at the middle of the A-domain and in the middle of the C-domain. Conversely, heterotypic recombinations occur all over the AT-domain and are more pronounced in both the CA-Linker and at the end of the A-domain (Fig. 4b and c). Homotypic recombinations are mostly very short, whereas heterotypic ones are on average longer and can span entire domains or modules (Fig. 4c and d). We then aimed to use homotypic recombinations to construct a proxy of what types of recombinations would fix under neutrality through genetic drift. To do this, we examined the fraction of homotypic recombinations in A-domains that span the so-called specificity area^30^ in the middle of the A domain. Specificity never changes through homotypic recombinations, so the fraction that covers the specificity area gives an estimate of how frequently the specificity area should be affected by recombination under neutrality. We then compared this to the fraction of recombinations that span the specificity area in heterotypic recombinations (Supplementary Table 3), which can change specificity. We find a statistically significant excess of heterotypic recombinations that cover the specificity area compared to homotypic ones for all recombinations with clearly identifiable borders (Supplementary Tables 3-5; p<0.05). The test was marginally significant (Supplementary Tables 6, p=0.0587, Fisher’s Exact test) only if we assigned all homotypic recombinations with ambiguous borders to cover the specificity area. These data suggest that rare, specificity-changing recombinations are probably driven to fixation by selection, implying that some of the GXP variants we discovered may confer better activities, allow new prey insects to be killed or enable escape from resistance.

To test these predictions from our analysis of recombination patterns, we next tested several of our GXP variants for their insecticidal activity. We chose four phylogenetically diverse insect model organisms: *Spodoptera exigua*, *Plutella xylostella*, *Tenebrio molitor* and *Frankliniella occidentalis*. We then tested the insecticidal activities of eight different types of GXP against these insects. To cover a wide variety of peptides, we chose **1**, **6**, **36**, **10**, **11**, **4, 5** and **38** (Fig. 5). We fed leaves soaked in solutions of these peptides to starving insect larvae and quantified the fraction of larvae that died after 3 days. We next tested if our data show evidence of species-specific toxicities of our peptides. To do this we fitted 5 models of increasing parameter richness (Supplementary Figs. 52-57) and used the Akaike Information Criterion (AIC) to determine which level of complexity most appropriately explains the data. The AIC chose the most complex model, which assigns each peptide-insect combination to a unique activity. For example, **36** is the most active peptide against *S. exigua,* but one of the least effective peptides against *P. xylostella*. *P. xylostella* is most effectively killed by **4**, which is of medium efficacy in *S. exigua* compared to other peptides. *T. molitor* and *F. occidentalis* were overall less susceptible to GXP, but at least *F. occidentalis* showed clear differences between peptides, with it being killed most effectively by **1**, **6**, and **10**, and least effectively by **4** (Fig. 5).

**Fig. 5:**
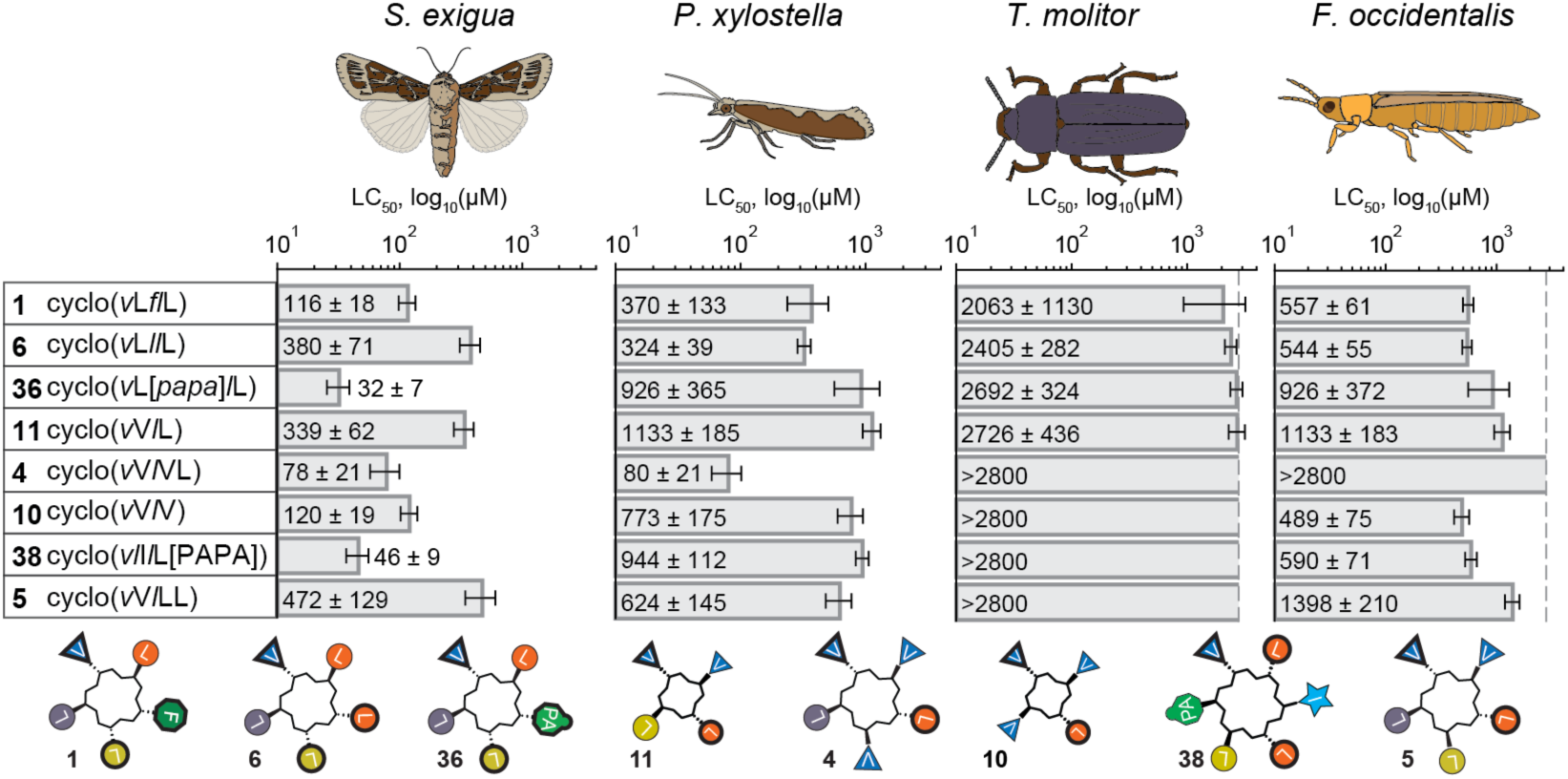
Insecticidal activity of 8 different GXP against four different insects. LC_50_ values for a selection of GXP when tested as insecticides against larvae of *S. exigua*, *P. xylostella*, *T. molitor* and *F. occidentalis*. Errors were calculated using 200 monte-carlo simulations. LC_50_ values above 2800 µM were not reliably determined and are shown as open squares without error bars. Every bar contains the corresponding LC_50_ value. Insects are shown as cartoons of the mature form for illustration; experiments were conducted on caterpillar larvae. Raw kill curves with fits are shown in Supplementary Fig. 56. The sequence of every NRP is written next to the number of the NRP. Capital letters refer to L-amino acids and small italic letters refer to D-amino acids. *p*-amino phenylalanine is shown as PAPA and in brackets.

These data show that structural variation in the GXP family leads to insect-specific activity differences, which could in principle be acted on by selection. At the same time, we find that for each insect there are several different peptides with similar potencies. This shows some degeneracy in the relationship between the NRP’ exact structure and their function as an insecticide, which would allow neutral drift through the space of chemically different but equally active peptides. Together with the large number of synonymous recombinations, this suggests that drift, in addition to positive selection, has plausibly fixed some of the new GXP on our tree.

## Discussion

Our data reveal that the chemical structures of NRPS-encoded NRP are subject to the same complex forces that shape protein sequences: both selection and random genetic drift drive NRP diversification, and therefore not all aspects of their structure can be assumed to be optimal. We show these processes can generate species-specific toxicities over short phylogenetic timescales. This has important implications for the evolutionary dynamics of NRPS clusters that produce antibiotics: resistance to antibiotic NRP evolves very fast, on a timescale of a few generations. Our work implies that changes in the chemical structures of NRP may overcome resistance on faster timescales than previously thought. Based on our discoveries, it is plausible that the phylogenies of well-known antibiotic-producing NRPS clusters also contain many so-far undetected recombination events leading to new peptides that overcome acquired resistances of their targets through modification of their products.

Our study of GxpS’ historical evolution reveals a deep connection between the evolvability of biological molecules and the genetic mechanisms by which new variation in the structure of these molecules is generated. The size of the amino acid alphabet available for NRPS produced peptides is very large, perhaps up to 300 different amino acids^32^. However, most of that alphabet appears to be inaccessible to a given NRPS. This is because recombination rates are very sensitive to local sequence similarity^33^. The possible evolutionary trajectories of an NRP are therefore dictated by which other NRPS domains are both available in the genome and similar enough for productive recombination. In our dataset, most recombination events occurred between modules of GxpS itself, which encode a very limited set of possible amino acids. The degree of convergent evolution we observe in the history of GxpS may in part be explained by the efficacy of new GXP but also by a recombination bias that results from the very few options GxpS has for recombination. By comparison, conventional protein evolution is much less constrained: every position in a normal protein is at most a few point mutations away from every other proteinogenic amino acid, irrespective of which amino acids are used elsewhere in the protein or the genome.

Our work not only highlights the promise of studying natural product evolution, but also its historical limits: The evolutionary mechanism by which NRP evolve erases the record of its own past. Each recombination obliterates a small part of the history of these enzymes, as one sequence and its historical record is overwritten completely by another. Over relatively small historical distances detailed phylogenetic work can still untangle these events. But recombinations have churned through NRPS for probably billions of years, fueling arms races between natural products and their molecular targets as well as the evolution of many other NRP-related functions. The evolutionary record of that ancient history is almost all lost, like tears in a rain of recombination.

## Data and materials availability

All raw data of HPLC-MS measurements, phylogenetic trees, alignments, ancestral sequences, PhyML_Multi and RDP generated files as well as raw bioactivity data will be deposited on Edmond, the Open Research Data Repository of the Max Planck Society and will be released for public access upon publication. Scripts to determine LC_50_ values and to perform statistics will be deposited on GitHub.

## Acknowledgments

We thank the Hochberg and Bode labs for careful reading of the manuscript. SK, TSMG, DVB, ZQ, DS, GKAH and HBB were supported by the Max Planck Society. C.J.B is grateful for the support of a Human Frontiers Science Programme Long Term Fellowship (LT0018/2024-L). NS and YK were supported by a grant (No. 2022R1A2B5B03001792) from National Research Foundation, Korea. SGG was supported by the Human Frontiers Science Programme grant RGP0028.

## Author contributions

SK, GKAH, HBB conceived the project, analyzed the data and planned experiments. SK, CL and ZQ performed molecular work. SK carried out phylogenetics, ancestral sequence reconstruction, recombination analysis and structural elucidation of the peptides. DS designed plasmids to construct the ancestral NRPS. TSMG and SK synthesized the peptides. YK planned the insect assays and NS the corresponding experiments. CJB, GKAH and SK performed curve-fitting for the insect toxicity data, determination of the LC50 values and the statistical tests. SGG extracted the SCG needed for species tree inference. DVB conducted insect assay using *X. doucetiae*. SK, GKAH and HBB wrote the manuscript.

## Competing interests

Authors declare that they have no competing interests.

## Materials and Methods

### Cultivation of strains

Liquid or solid low salt LB medium (pH 7.5, 10 g/L tryptone, 5 g/L yeast extract and 5 g/L NaCl) was used to culture *E. coli* DH10B::*mtaA* cells (Supplementary Table 7). Selection markers were either kanamycin (50 µg/ml), ampicillin, chloramphenicol (34 µg/ml) or gentamycin (20 µg/ml). 1 % (w/v) agar were added to create solid media. The cultivation temperature was 37 °C. For peptide production, cultivation temperature was 22 °C.

### Cloning of biosynthetic gene clusters and NRPS modules

Genomic DNA (gDNA) was extracted from *Xenorhabdus* and *Photorhabdus* (XP) as listed in Supplementary Table 7 using the Monarch® Genomic DNA Purification Kit (NEB). The gDNA was taken as a template for PCR amplification. PCR was conducted using the Q5® High-Fidelity DNA Polymerase (NEB). The generated oligonucleotides and their size are depicted in Supplementary Table 8. Also documented in Supplementary Table 9 were already cloned plasmids that were also used as a PCR template. Monarch® DNA Gel Extraction Kit (NEB) were used to gel extract the PCR products. These purified PCR products were used as a substrate for Gibson cloning using the NEBuilder® HiFi DNA Assembly Cloning Kit (NEB).

The ancestral GxpS domains were ordered from Genscript Biotech Corp. We bought 5 plasmids. Plasmid 1 – 4 contains ATC-tridomain 1 – ATC-tridomain 4 of *ancgxpS*, respectively. Plasmid 5 contained ATTE-tridomains of *ancgxpS*. Every natural occurring *bsaI*-sites was removed within the NRPS sequence. We added to plasmid 1 – 5 suitable *bsaI*-sites to clone the full *ancgxpS* gene into pMY-e1-0000_GG to create pSK37 and pSK43 (Supplementary Table 9). For Golden Gate the NEBridge® Golden Gate Assembly Kit (BsaI-HF®v2) was used.

### Heterologous expression of NRPS constructs and HPLC-MS analysis

Each plasmid was transformed into *E. coli* DH10B::*mtaA*. These cells were then grown overnight in LB medium containing the required antibiotics (50 µg/ml kanamycin, 34 µg/ml chloramphenicol, 100 µg/ml ampicillin, 20 µg/ml gentamicin). 10 ml XPP media^34^ that were supplemented with the required antibiotics, and 1 % of the overnight culture was added. The cells were first incubated for 3 hours at 37°C and then incubated 69 hours at 22 °C. For nonribosomal peptide (NRP) extraction, 100 ul of the production culture were mixed to 900 ul methanol and shaken for 5 min. Then, the culture was centrifuged for 20 min at 15871 g. The supernatant was transferred to HPLC-MS tubes. UltiMate 3000 system (Thermo Fisher) coupled to an AmaZonX mass spectrometer (Bruker) with an ACQUITY UPLC BEH C18 column (130 Å, 2.1 mm × 100 mm, 1.7-mm particle size, Waters) at a flow rate of 0.4 ml min−1 (5–95% acetonitrile/water with 0.1% formic acid, vol/vol, 16 min, UV detection wavelength 190–800 nm) was used for HPLC-UV-MS analysis. HPLC-UV-HRMS analysis was conducted on an ESI ion-trap mass spectrometer (timsTOF fleX MALDI 2, Bruker). ESI-MS spectra were detected in positive-ion-mode with the mass range from *m/z* 100-2000, and ultraviolet (UV) signal at 190 – 800 nm. DataAnalysis 5.3 software (Bruker) was used to evaluate the data.

### Confirmation of the absolute amino acid configurartion in NRP

To determine the incorporation of leucine, valine and phenylalanine and their respective stereochemistry, *E. coli* DH10B^penta^ was cultured with XPP media that lacks these amino acids. 2 mM of the deuterated versions of leucine, valine and phenylalanine were added to the culture media. *E. coli* DH10B^penta^ was co-transformed with two plasmids containing either a plasmid containing a *mtaA* gene and another plasmid carrying the *gxpS* gene. Cell culturing and HPLC-MS measurements were done as described above but the culture volume was 5 ml. Determination of the absolute amino acid composition was done as described previously^18,23^. Also, co-injection with synthetic and natural produced compound was done to confirm the structure of GXP (Supplementary Tables 1 and 2 and 11). For Fig. 2c and f: we determined the two highest peaks as our main peptides. If three peaks had an equal height, we omitted the third one for simplicity and treated this peak as a side product. Our main text figures therefore somewhat understate the true diversity of products.

### Chemical synthesis of GameXPeptides

GameXPeptide derivatives were synthesized through solid-phase peptide synthesis of the linear precursor and subsequent cyclization in solution. The linear peptide precursor was synthesized using the peptide synthesizer Liberty Prime 2.0 from CEM with a high-throughput loader (HT12) and preloaded 2-chlorotrityl resin (100-200 mesh). For coupling, the respective Fmoc-amino acid solution (1 mL), N,N’-diisopropylcarbodiimide (DIC, 1 mL), and 2-cyano-2-(hydroxyimino)acetate/N,N’-diisopropylethylamine (Oxyma/DIPEA, 1.5 mL) were combined in the resin-containing reaction vessel and left to react for 60 minutes at room temperature with agitation through a nitrogen flow. After two washing steps with dimethylformamide (DMF), the Fmoc group was deprotected with pyrrolidine (0.75 mL) for 10 minutes at room temperature. Afterward, the resin was washed five times with N-methylpyrrolidone (NMP), DMF, and dichloromethane (DCM) and dried. Peptide cleavage of the resin was performed by incubating the resin with hexafluoroisopropanol (HFIP) in DCM (1:4 v/v, 6 mL/100 µmol) for approximately two hours until the color faded. The peptide-containing solution was collected in a round-bottom flask, and the resin was washed twice with the cleavage solution. After the solvent was evaporated under reduced pressure, the solid residue was washed twice with DCM (via solvation and evaporation) to obtain the linear peptide as a colorless solid.

Subsequent head-to-tail cyclization was performed in solution. The linear peptide was dissolved in DMF to a concentration of 1 mM. After adding DIPEA (4 eq) and HATU (4 eq) and gentle shaking for a few seconds, the colorless solution turned yellow-orange, indicating successful cyclization. The cyclization of tetrapeptides was carried out similarly but with twice the amount of DIPEA (8 eq) and an overnight incubation. Samples were taken before and after the reaction and analyzed by LC-MS to validate and quantify the cyclization reactions. The solvent was then removed under reduced pressure (water bath temperature of 60 °C).

The synthesized peptides were purified via filtration using solid-phase extraction (SPE) Strata C_18_-E (55 μm, 70 Å, 200 mg/ 6 mL) cartridges from Phenomenex. The following protocol was performed (using the SPE cartridges as a filtration tool) (Supplementary Table 10). The purified peptide was transferred using a spatula and methanol. All remaining solvents were removed under reduced pressure to yield the desired peptide as a colorless solid (Supplementary Table 11).

### Testing insecticidal activity of different GXP on insects

The peptides were prepared with dimethylsulfoxide (DMSO) at a concentration of 1000 ppm (1 mg/ml). The stock solutions were stored in the freezer and diluted with DMSO to prepare working solutions. The stock solution was diluted to 1000, 500, 100, 50 and 5 ppm working solution. DMSO and bovine serum albumin (BSA) were purchased from Sigma-Aldrich Korea (Seoul, Korea). An anticoagulant buffer (ACB, pH 4.5) was prepared with 186 mM of NaCl, 17 mM of Na 2 EDTA, and 41 mM of citric acid. Phosphate-buffered saline (PBS) was prepared with 100 mM phosphoric acid and the pH was adjusted to 7.4 using 1 N NaOH.

The oral toxicity of different test compounds was evaluated in *Spodoptera exigua*, *Plutella xylostella*, *Tenebrio molitor*, and *Frankia occidentalis*. Prior to treatment, larvae were starved for 5 h to enhance ingestion of treated diets. Diets were soaked in test compound solutions for 3 min and air-dried on filter paper for ∼5 min to remove excess liquid. This was done for every different working solution. DMSO-treated diets served as negative controls. Ten larvae per treatment group were fed the prepared diets and maintained at 25 ± 2°C. Mortality was recorded at 3 days post-treatment. Each treatment with each 10 different larvae was replicated three times.

### Statistical analysis of the bioactivity data

Sigmoid functions with hill and IC50 values (2 parameters total) were fitted to insect lethality data using numpy^35^, scipy^36^ and matplotlib^37^ libraries in python. To estimate errors for each parameter, a Monte-Carlo procedure was performed: data were resampled 200 times using errors from triplicate measurements and refitted. Standard deviations of the resulting parameter fits were used as the errors. 5 models of increasing complexity were evaluated, and the model with the lowest Akaike Information Criterion (AIC) was chosen. AIC in principle penalizes more parameters, so indicates if additional parameters are justified by the lowered error. AIC parameters were calculated using a log likelihood derived from the least squares fitting residuals which assumes gaussian errors (Supplementary Table 12)^38^:

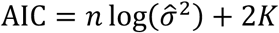

Where:

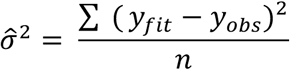

### Collection of sequences

All sequences were collected from the in-house Bode group sequencing collection and from The National Center for Biotechnology Information (NCBI) RefSeq^39^.

### Natural products produced during the infection of *Galleria* with *Steinernema disprepesi* carrying

#### Xenorhabdus doucetiae

A total of 60 *Galleria mellonella* instar larvae were disinfected using 70% alcohol and placed separately in 1.5-ml Eppendorf tubes with filter paper, on which 50 IJs of *Steinernema diaprepesi* were placed. The insects were maintained at a temperature of 24°C during the incubation process. Three larvae were collected at 12-hour intervals until the 120-hour mark, at which point 30 microliters of hemolymph were extracted and stored at −20°C for further analysis. Following the completion of the sampling procedure, each hemolymph sample was subjected to a dilution process using a 1:1 (v/v) ratio of MeOH. This was followed by a centrifugation step at 13,000 rpm for 20 minutes. Subsequently, 50 μl of the clear supernatant was transferred to vials for further analysis. The extracts were analyzed using a TimsTOF (Bruker) instrument with a gradient solution comprising solvent A (ddH_2_O + 0.1% FA) and solvent B (ACN + 0.1% FA), at a flow rate of 0.4 ml/min, with a linear gradient from 5% to 95% of solvent B over a duration of 14 minutes. The data analysis was conducted using Bruker Compass DataAnalysis 5.3.

### Determination of A– and TC-domain phylogenetic unit

215 ATC-tridomain sequences were extracted from 54 GxpS sequences from *Xenorhabdus* and *Photorhabdus* using local BLAST with the first ATC tridomain from GxpS of *Photorhabdus laumondii* TT01 as query. The ATC sequences were aligned using mafft v7.490 ––globalpair option^40^ and trimming was done using trimAl 1.2 using –resoverlap 0.8 and –seqoverlap 75^41^. Then PhyML_Multi^25^ was run with both the Hidden Markov Model (HMM) and the Mixture Model (MM) that together best fit the alignment. First, the best fit model was determined using ModelFinder implemented in IQ-TREE^42^. Afterwards, PhyML_Multi was run with JTT^43^ as the best fit model with a 4-category gamma distribution of among site rate-variation. PhyML_Multi uses PhyML v2.4.4 for tree inference^44^. The site-wise log-likelihood values from tree 2 were deducted from those from tree 1 and then plotted. To determine the phylogenetic border between the A– and TC-domain, we used the outputs of the HMM and MM analysis and applied the Viterbi and Forward-Backward algorithms that is implemented in PhyML_Multi (Supplementary Fig. 58).

The mixtures across sites and trees (MAST)^45^ analysis was done twice using the same alignment as for the PhyML_Multi analysis. First, we used for the MAST analysis the trees generated by PhyML_Multi. Then, we used separated inferred trees from an A (including A1 – A4) and C-domain (including C1 – C4) alignment that we used as an input for our analysis. For the first MAST analysis, we used again JTT as the best fit model with a 4-category gamma distribution of among site rate-variation. JTT with a 5-category free rate distribution of among site variation was used for the second analysis. In both cases, log likelihood values of the A and C-domain tree were deducted and plotted (Supplementary Fig. 59).

### Phylogenetics and ancestral sequence reconstruction

Before every round of tree inference, model finder implemented in IQ-TREE was used to determine the best fit model of every nucleotide or amino acid alignment. For ASR, we used the tree topology we obtained from amino acid alignments, but used nucleotide sequences to infer ancestral sequences in order to preserve the codon usage from *Xenorhabdus* and *Photorhabdus* to ensure such a long gene is translated appropriately. To do this, we used the topology from trees inferred from protein sequences and reoptimized optimized branch-lengths and substitution parameters using a corresponding nucleotide alignment. Ancestral sequences and posterior probabilities of ancestral state were inferred using the –ASR option in IQ-TREE.

For the GxpS A-domain tree in Extended Data Fig. 4a, 269 A-domain sequences consisting of nucleotides with the preceding C-A-Linker region were extracted from all 54 GxpS sequences to construct the phylogeny. These sequences were codon-aligned using the translation align option implemented in AliView^46^. Gaps were carefully trimmed manually. Maximum likelihood tree inference was done using IQ-TREE multicore version 2.0.7^47^ with TVMe as the best fit model with a 6-category free-rate model (R6) for among-site rate variation. The A-domain nucleotide alignment was translated to an amino acid alignment. This amino acid alignment was used to infer another tree to with JTT as the best fit model with a 6-category free rate distribution of among site variation. Afterwards the branch-lengths of the protein A-domain tree were optimized using the nucleotide A-domain alignment applying the –te option in IQ-TREE. The A-domain tree was rooted using paralogous A-domains specific for valine as the outgroup.

For the GxpS TC-domain tree in Extended Data Fig. 4b; the same procedure was done using 215 TC-domain nucleotide sequences including the preceding A-T-Linker. For nucleotide tree inference, TVM was the best fit model with empirical state frequency observed from the data and a 10-category free rate distribution of among site variation. JTT was the best fit model with a 5-category free rate distribution of among site variation for the protein tree. The TC-domain tree was rooted using the paralogous^L^C_L_-domains as the outgroup.

161 nucleotide sequences of phenylalanine and leucine incorporating A-domains of GxpS, KolS^22^ and PhpS^33^ were extracted and codon aligned using the translation align option implemented in AliView version 1.28. Gaps were carefully removed manually. TVM with empirical state frequency observed from the data and a 5-category free rate distribution of among site variation were used to infer the nucleotide tree. The best fit model of the translated protein alignment was the use of freedom to search for invariable sites with a 4-category gamma distribution of among site rate-variation. The branch length of the protein tree was optimized with the nucleotide alignment using the –te option in IQ-TREE. The tree was rooted using the paralogous leucine incorporating A-domains as the outgroup.

54 nucleotide sequences of the TTE-domain of GxpS and 3 nucleotide sequences of the TTE-domain of KolS^22^ (that was used as the outgroup) including the prior A-T-Linker were extracted and codon aligned using the translation align option implemented in AliView. Gaps in the alignment were carefully trimmed. TVM with empirical state frequency observed from the data, the freedom to search for invariable sites with a 4-category gamma distribution of among site rate-variation were used to infer the TTE-domain tree. The best fit model of the translated protein tree was JTT with a 4-category gamma distribution of among site rate-variation. Branch length optimization of the protein tree with the nucleotide alignment was done using the –te option in IQ-TREE.

We used the last T-domain including the TE-domain because it added more sites to our reference (gene) tree. This led to better agreement between species tree and gene tree, as well as less incongruent to A– and TC-domains overall when compared with this tree.

Nucleotide extraction was done using AliView on the whole GxpS alignment. Manually, A, TC or TTE-domains were selected and extracted.

For all trees we computed support values using ultrafast bootstraps approximation (UFBoot)^48^ together with SH-like approximate likelihood ratio test (SH-aLRT)^49^ with 1000 bootstrap replicates as implemented in IQ-TREE.

### Species tree inference

The XP species tree was inferred with GToTree v1.7.00^50^, using the prepackaged single-copy gene-set for proteobacteria consisting of 119 target genes. In order to predict genes on input genomes provided as fasta files, prodigal v.2.6.3 was used^51^. This software package identifies the target genes in all genomes with HMMER3 v3.3.2^52^. Afterwards, it aligns the identified target genes with muscle v5.1^53^ and then trims the alignment using trimal v.1.4.rev15^41^. Concatenation was done with FastTree2 v2.1.11^54^ before the species tree was inferred. GToTree workflow then used TaxonKit to connect full lineages to taxonomic IDs^55^. The dataset included all sequenced XP genomes from the Bode in house dataset. Additional XP genomes were downloaded from NCBI. As outgroup, SCGs from few gammaproteobacteria were used. JTT was used as the best fit model as determined by ModeFinder implemented in IQ-TREE with empirical state frequency observed from the data and a 5-category free rate distribution of among site variation. Also, a second tree was inferred, that contained *Pseudomonas* species. Here, the best fit model was JTT with empirical state frequency observed from the data and an 8-category free rate distribution of among site variation.

### Reconciliation

Reconciliation was done using notung v.3.0-24^24^. As input, we used the TTE-domain tree as our gene tree and reconciled it with our species tree. Notung uses the Edge Weight Threshold of 0.15, a Duplication Cost of 1.5, a Transfer Cost of 3, a Co-Divergence Cost of, and a Loss Cost of 1 as parameters.

### Phylogenetic analysis of the PAPA operon

A BLAST analysis on every gene of the PAPA operon was done to sample orthologous sequences. Every PAPA protein sequence collection was aligned individually using using mafft –globalpair option. Gaps were manually trimmed. Maximum likelihood tree inference was done for every gene using the best fit model as determined by ModelFinder implemented in IQ-TREE. Branch support was determined using 1000 bootstrap iterations applying UFBoot and the SH-aLRT.

To visualize the distribution of the PAPA operon among different XP species genofig v.1.1.1^56^ was used. The genomic area that contained the PAPA operon were taken as gene bank file to feed the software genofig.

### Detection of recombination in NRPS

To get a rough estimate of NRPS recombination, we used HMM approach implemented in PhyML-Multi. We used the A-domain nucleotide alignment as input and translated it into amino acid sequences. We then used PhyML_Multi to find the two best trees that best describe the alignment. JTT was chosen as the best fit model as done for resurrection. Since PhyML_Multi does not contain free rate categories, we chose as rate heterogeneity across sites a gamma model with 4 rate-categories. We further determined the breakpoints by choosing the Viterbi and forward-backward algorithm. PhyML_Multi could roughly detect the unrecombined and recombined tree. However, this alignment contains many recombinations, which cannot be described by only two trees, making it hard to find exact breakpoints using this approach. For exact breakpoint determination, we used Recombination Detection Program v. 5.23 (RDP5)^30^.

To detect recombination within ATC-domains the Recombination Detection Program^30^ v. 5.23 (RDP5) was used. We tested putative recombination within our A– and TC-domain alignment used for ASR. Further, we also aligned several further sequences with different lengths or taxa where we suspected other hidden recombination events. In our analysis, we used the default options in RDP. That included the algorithms of RDP^57^, GENECONV^58^, MaxChi^59^, BootScan^60^ and SiScan^61^. To improve recombination detection, in some cases also the algorithms of Chimaera^62^ and 3Seq^63^ were used. The highest acceptable p-value was 0.05. For RDP a window size of 30 nucleotides was used. To confirm recombination events, we always inferred a maximum likelihood tree of the unrecombined and the recombined area of the sequence alignment. For maximum likelihood tree inference, we used IQ-TREE and ModelFinder to determine the best fit model. UF-Boot and SH-aLRT were used to determine the branch support. For very short fragments, we used nucleotides and for longer ones in most cases amino acid. This is always listed in the figure legend for the corresponding recombination.

Detecting in particular synonymous recombinations is very difficult, because we cannot produce experimental verification of these events the way we can for non-synonymous ones. It is therefore likely that we missed potentially many synonymous recombinations. It is also possible that we included false positives, and the frequency of such false assignments is not known to us. A further complication is that recombinations frequently happen in areas that have already recombined. This makes finding recombinations a recursive and manual process that does not follow a fixed set of steps. What gives us confidence is that excluding recombined areas in most cases returns our phylogenies to congruence with the TTE-domain phylogeny.

Underneath every RDP plot, we showed the corresponding domain that was affected by recombination. In order to visualize the domain, we used Geneious Prime® 2023.2.1.

### Incomplete lineage sorting (ILS) in our dataset

To construct a resolved history of recombination events, we assigned each of the 50 recombination events we discovered to a branch on our TE-domain phylogeny Supplementary Fig. 60 and Extended Data Fig. 8. For virtually all recombination events, we could identify a single branch over which the recombinations occurred, which was then inherited by all descendants of that branch. But in one case this was clearly not possible (Supplementary Fig. 65). This recombination event inserted parts of the fifth A-domain that is specific for leucine into the 3^rd^ domain, diminishing its ability to incorporate Phe. However, placing this recombination event on the branch leading to all species that share it includes one species, *Xenorhabdus maulenoii,* in which this recombination seemingly never took place. There are two possible explanations for this: Either this recombination event occurred twice independently. We judge this to be extremely unlikely, as the donor and borders of this recombination are inferred to be identical between all GxpS that carry it. More likely, this pattern is caused by incomplete lineage sorting. This occurs if a polymorphism persists for long enough for speciation to occur without fixation of either the ancestral or derived allele (here the unrecombined and recombined sequences in the A3 domain, respectively). In this case, *X. maulenoii* would then have fixed the ancestral, unrecombined allele, whereas the recombined allele was fixed twice independently. This would imply that there is long lasting standing variation in the structure of GXP in populations of XP. We illustrated the process of ILS in Supplementary Figs. 65. The alternative to this interpretation is to assume independent recombination events with identical borders, or that an already recombined fragment was horizontally transferred and recombined into other synthetases with borders outside the original recombination borders. We can produce no evidence for this latter case and therefore chose incomplete lineage sorting as our working hypothesis for these cases.

### Statistical analysis of recombination patterns

The length between homotypic and heterotypic recombinations were analyzed and plotted on a boxplot (Fig. 4d). To test if homotypic and heterotypic recombinations are significantly different a Wilcoxon rank-sums test was used. Wilcoxon rank-sums test was done using the python package scipy.stats^36^. The raw data of all recombinations and their breakpoints is summarized in Supplementary Figs. 107 and 108. The modular structure of a CAT-tridomain was illustrated with Geneious Prime® 2023.2.1.

To determine if most recombinations are shaped by selection or drift, the number of homotypic and heterotypic recombinations that targeted the specificity pocket were compared to homotypic and heterotypic recombinations that did not target the specificity pocket^64^. We used a Fisher’s Exact Test to test if the distribution between columns and rows is random. Fisher’s Exact Test was used on the graph pad web browser application. We ignored C-domain recombinations because all C-domain recombinations are homotypic. C-domain recombinations that are heterotypic do not target the C-domain instead they target the A– or T-domain.

There are cases where we are not certain if some recombination has occurred several times or if they are a result of ILS. Similarly, for some recombination events, we were not sure whether they covered the specificity pocket, because one of the recombination breakpoints could not be detected. To verify that our conclusions are robust to this uncertainty, we created two additional tables that did and did not count these ambiguous cases. For Supplementary Tables 3–6, we used the Fisher’s exact test for significance testing. For 3 out of 4 tables, the test was significant. Only when all ambiguous cases were classified such that they are least consistent with a difference between homotypic and heterotypic recombinations was the test not significant above the 0.05 level, and it remained very close to significance. Our data therefore favour the interpration that heterotypic recombinations are different from our proxy of the neutral process, though some ambiguity remains because of the difficulty in correctly identifying all recombination breakpoints.

---

**Extended Data. Fig. 1:**
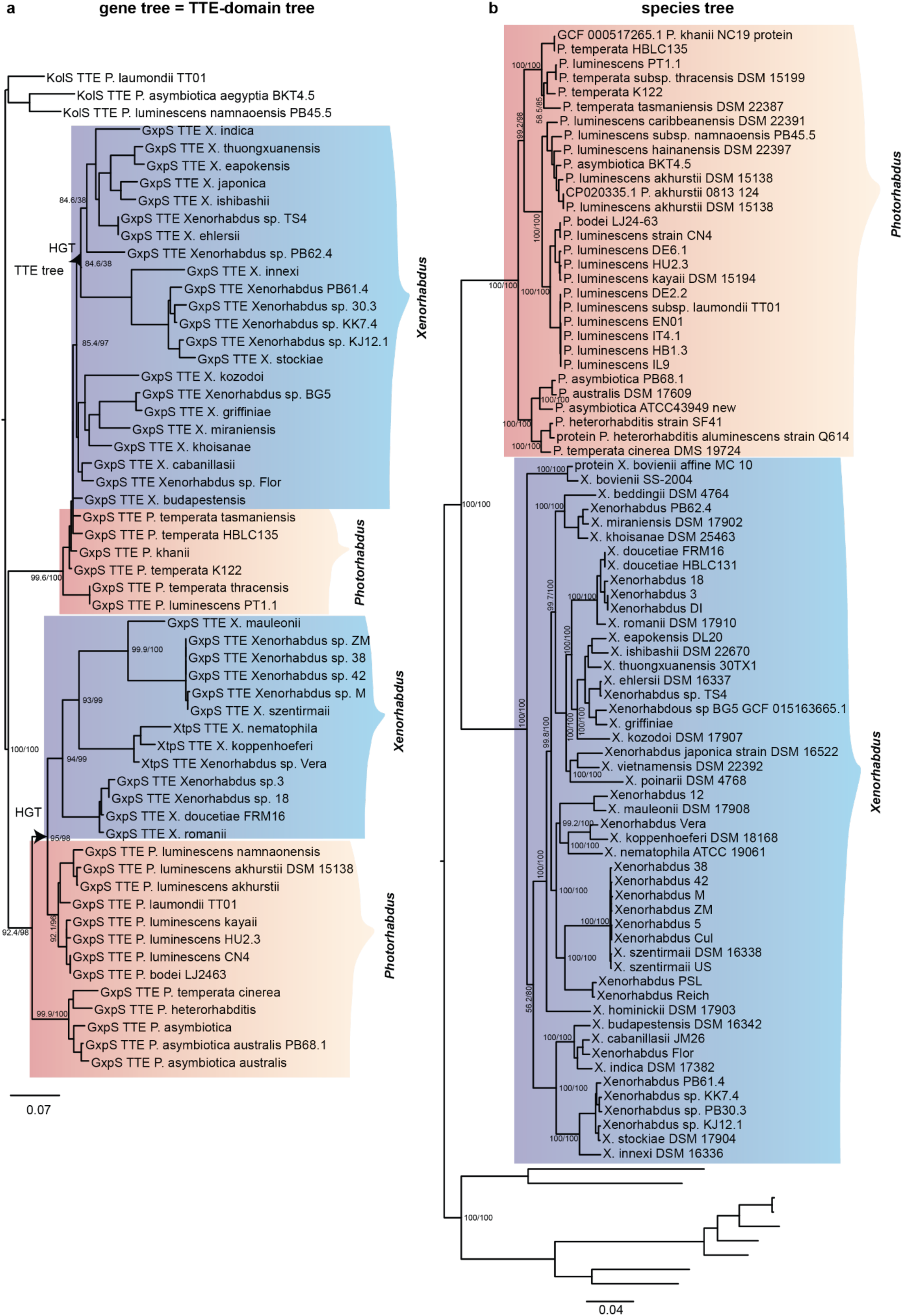
Species tree and gene tree of *Xenorhabdus* and *Photorhabdus*. SH-alrt and UF-Boots are annotated at major clades. (**a**) An expanded view of the TTE-domain tree. Here, XP are not monophyletic and *Xenorhabdus* branch inside *Photorhabdus* twice. (**b**) The species tree was inferred from a concatenation of 119 SCGs. *Xenorhabdus* and *Photorhabdus* form a monophyletic group, respectively. Other gammaproteobacteria were taken as the outgroup.

**Extended Data. Fig. 2:**
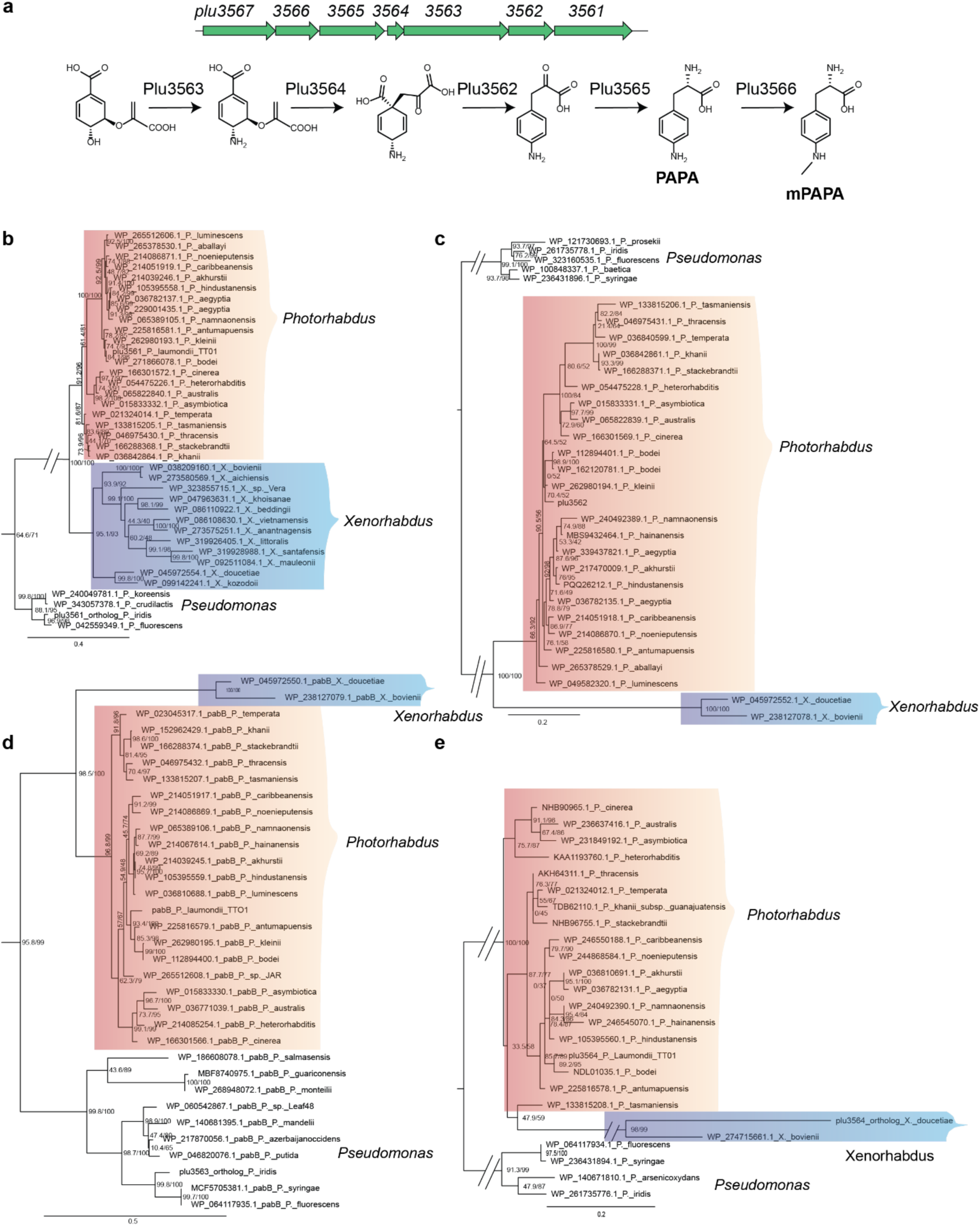
Maximum-likelihood protein trees of four different genes from the PAPA operon. (**a**) Biosynthetic pathway of the PAPA-operon as shown previously. (**b**) *plu3561*, (**c**) *plu3562*, (**d**) *plu3563* (*pabB*), (**e**) *plu3564* from the PAPA operon. The closest outgroup are PAPA operon genes from *Pseudomonas* species.

**Extended Data. Fig. 3:**
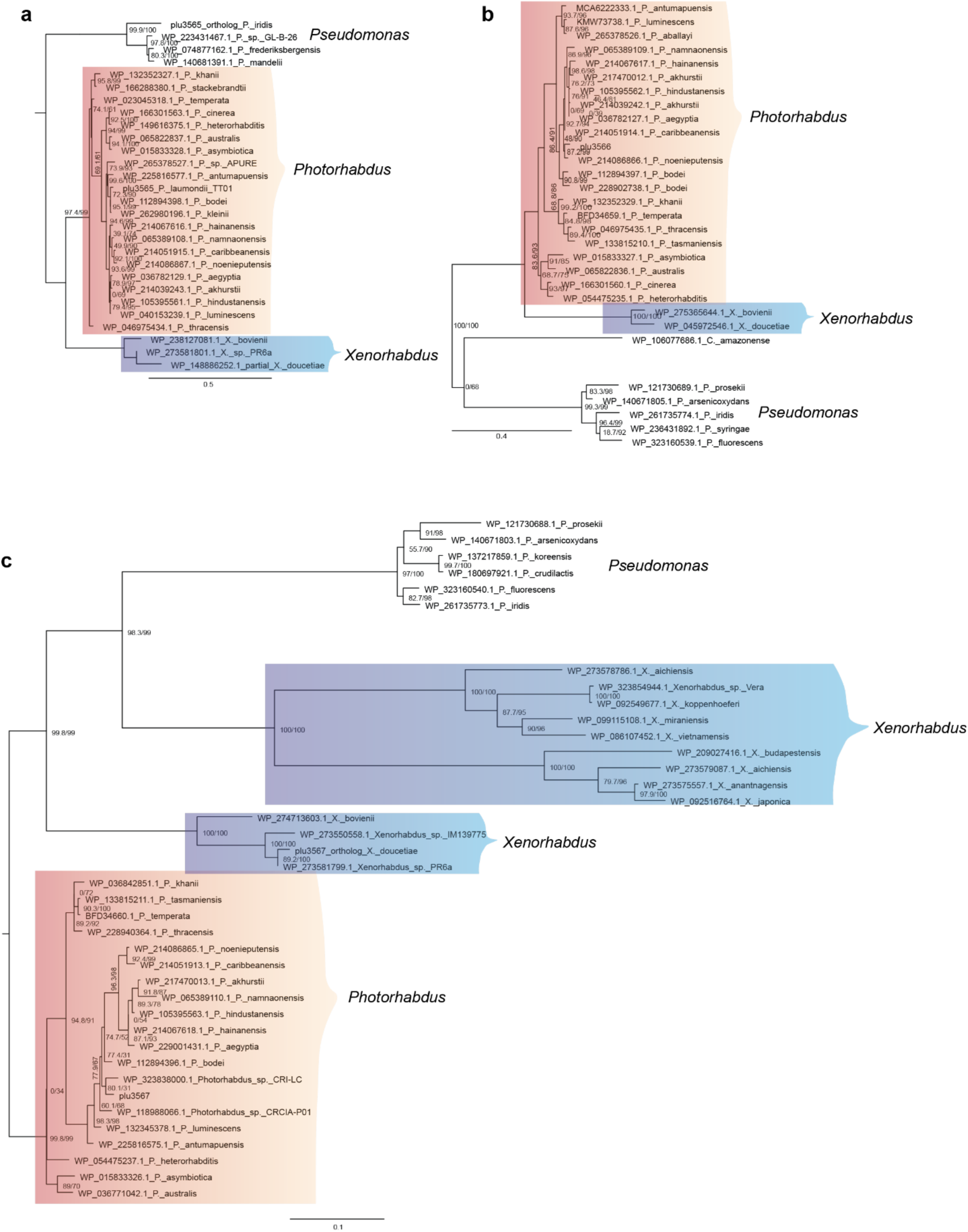
Maximum-likelihood protein trees of three remaining genes from the PAPA operon. (**a**) *plu3565*, (**b**) *plu3566*, (**c**) *plu3567* from the PAPA operon. The closest outgroup are PAPA operon genes from *Pseudomonas* species.

**Extended Data. Fig. 4:**
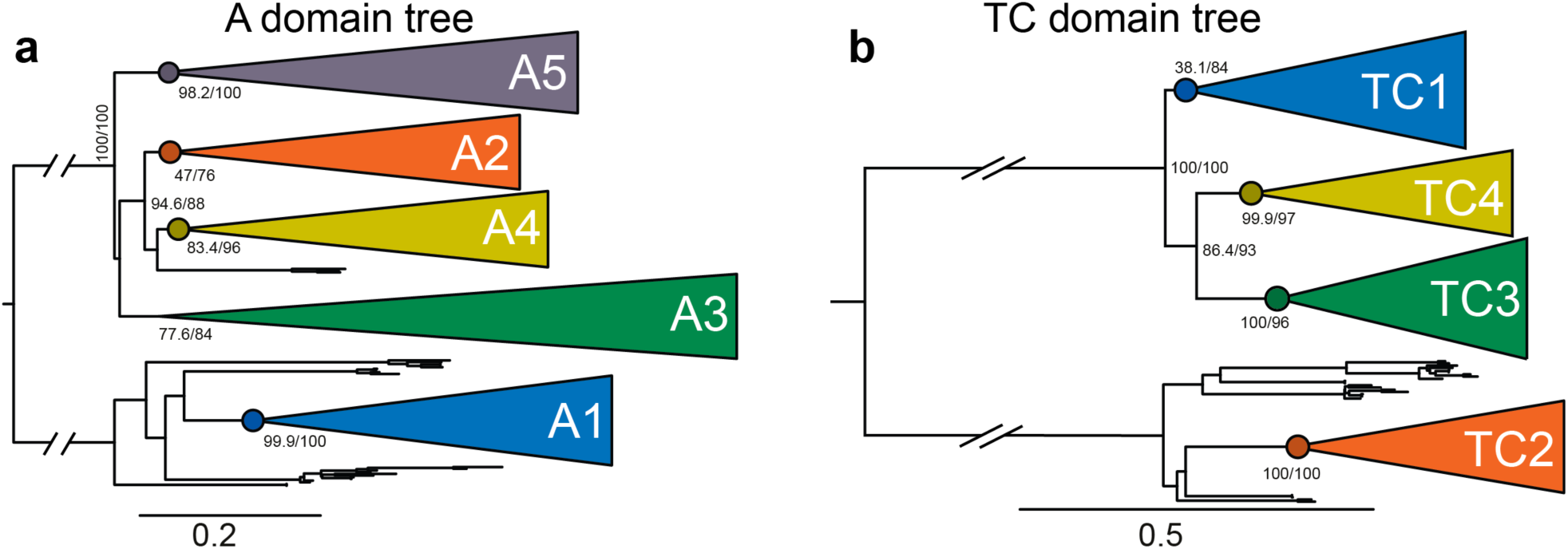
A phylogenetic tree of all A– and all TC-domains of all GxpS from XP. (**a**) Phylogenetic tree of all A-domains from all GxpS from XP. All first, second, third, fourth and fifth A-domains form a monophyletic group. (**b**) Phylogenetic tree of all TC-domains from GxpS from XP. All first, second, third, fourth and fifth TC-domains form a monophyletic group. (**a+b**) Nodes used for ASR are shown as dots. The color code of the differenT-domain clades was the same as for the GxpS diagram displayed in Fig. 1a. SH-aLRT and UF-Boot values are shown at every GxpS domain clade.

**Extended Data. Fig. 5:**
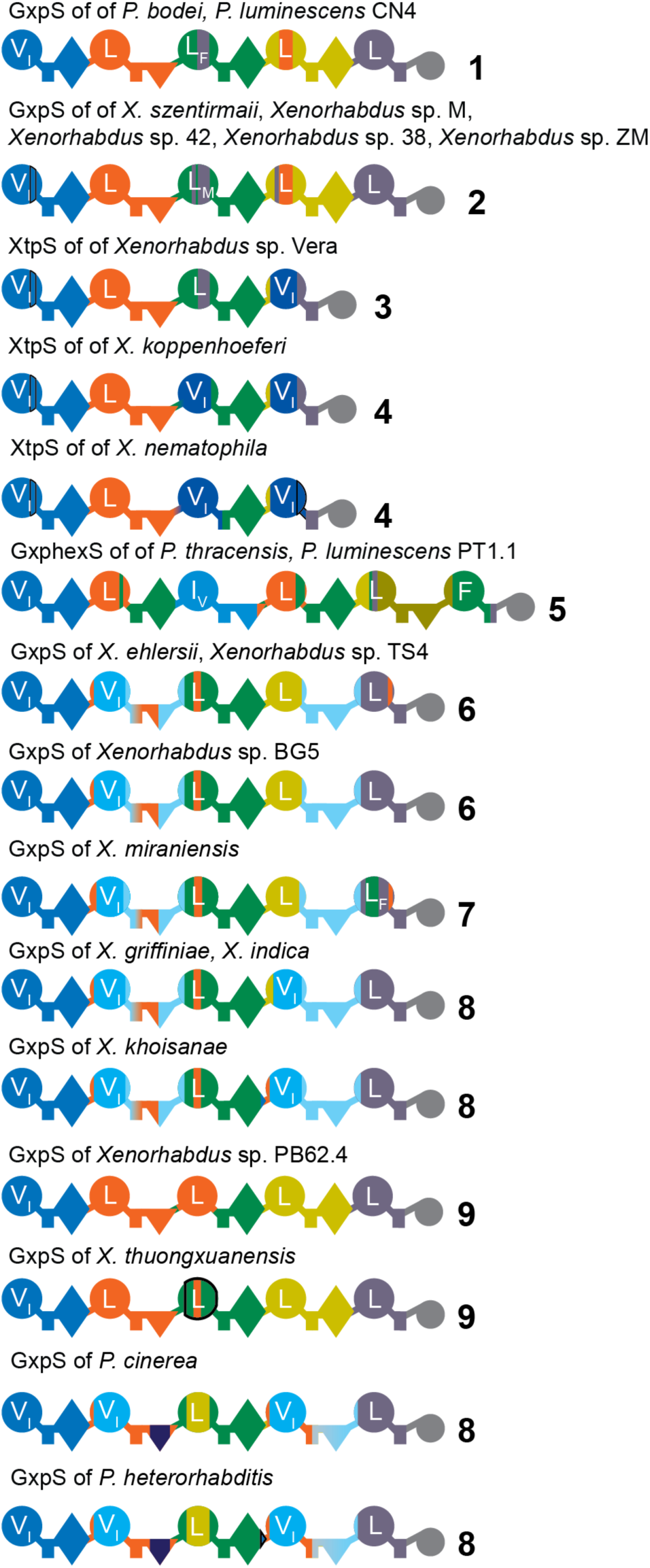
All GxpS that have changed through non-synonymous recombinations. 23 of 54 species have experienced non-synonymous recombinations in their *gxpS*. In total there are 9 new variants of GxpS with different specificity as the canonical GxpS from *P. laumondii* TT01. The different variants GxpS structure are numbered 1-9.

**Extended Data. Fig. 6:**
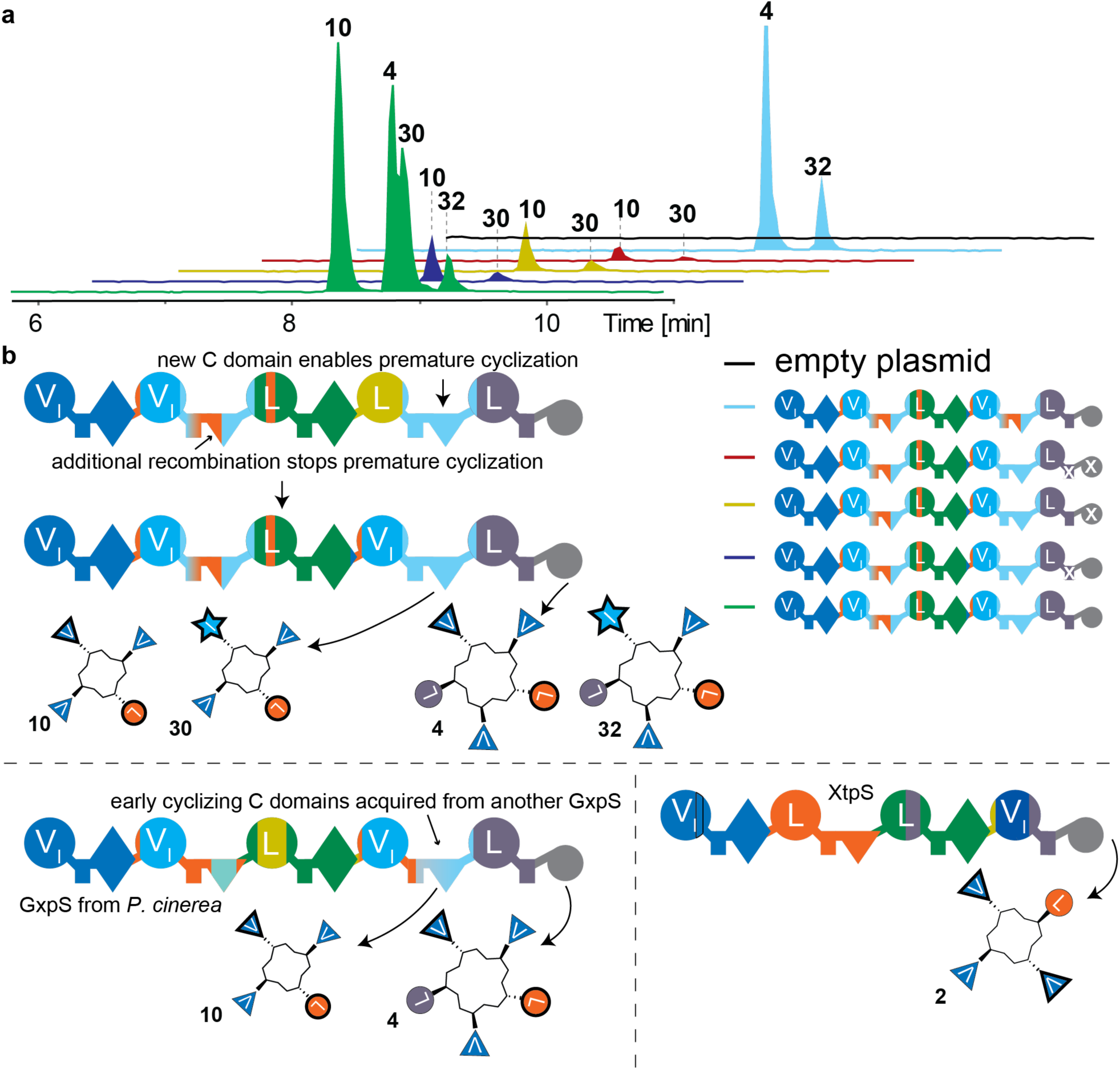
Premature peptide release is mediated by a novel C-domain. (**a**) Retention time of the cyclic pentapeptides (**4**, **32**) and cyclic tetrapeptides (**10**, **30**) produced by the GxpS of *X. khoisanae*. Solely tetrapeptide production was detected for the GxpS mutant where the 5^th^ T-domain and the TE-domain was knocked out. Solely pentapeptide production was detected when the GxpS C2 domain was replaced with its own C4 domain. All GxpS mutants of *X. khoisanae* were heterologously expressed in *E. coli* DH10B::*mtaA*. As negative control, the empty pCOLA plasmid was used. HPLC-MS was done to measure natural product production. (**b**) A GxpS diagram of the GxpS of *X. khoisanae* that explains the evolution of this GxpS and why this GxpS produces cyclic penta– and tetrapeptides. The introduction of a new C-domain in the second and fifth position enabled premature release of cyclic peptides. Also shown is the GxpS of *P. cinerea* that produces the same NP as the GxpS of *X. khoisanae*. It acquired the early cyclizing C-domain horizontally from *X. khoisanae*. XtpS is displayed that produces tetrapeptides due to its four modular structure.

**Extended Data. Fig. 7.**
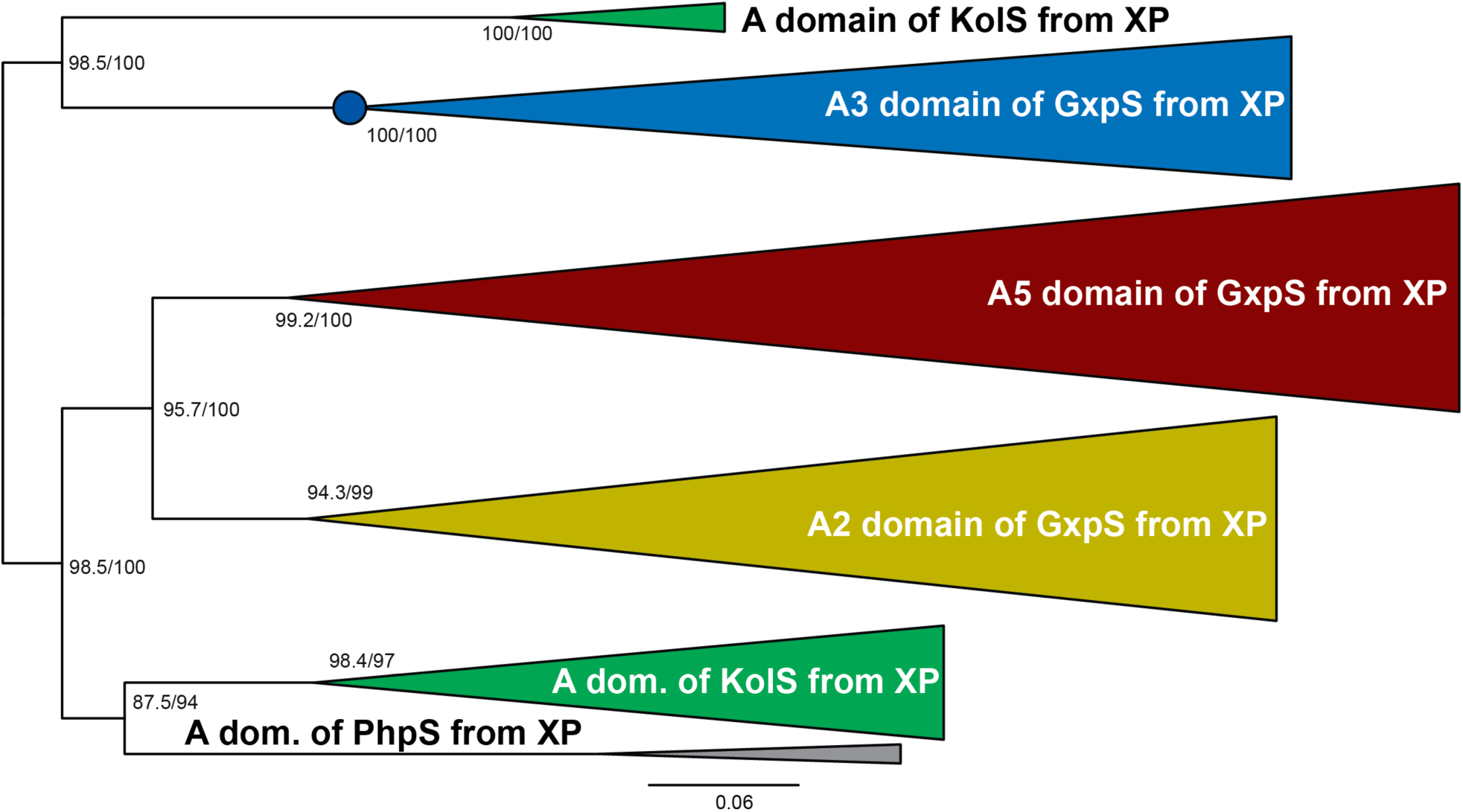
An A-domain maximum likelihood nucleotide tree containing A2, A3 and A5 domains from GxpS. It also contains A-domains from KolS^22^ and PhpS^34^. This tree contains two main clades. One clade that contains the A3 domain of GxpS and A-domains of KolS that are specific for phenylalanine or tyrosine. The other clade contains all A-domains specific for leucine. All A3 domains of GxpS that had been affected by non-synonymous recombination were excluded from this tree inference. This tree was used to calculate the last common ancestor of all A3 domains of GxpS. SH-aLRT and ultrafast bootstrap values are shown at every domain clade.

**Extended Data. Fig. 8:**
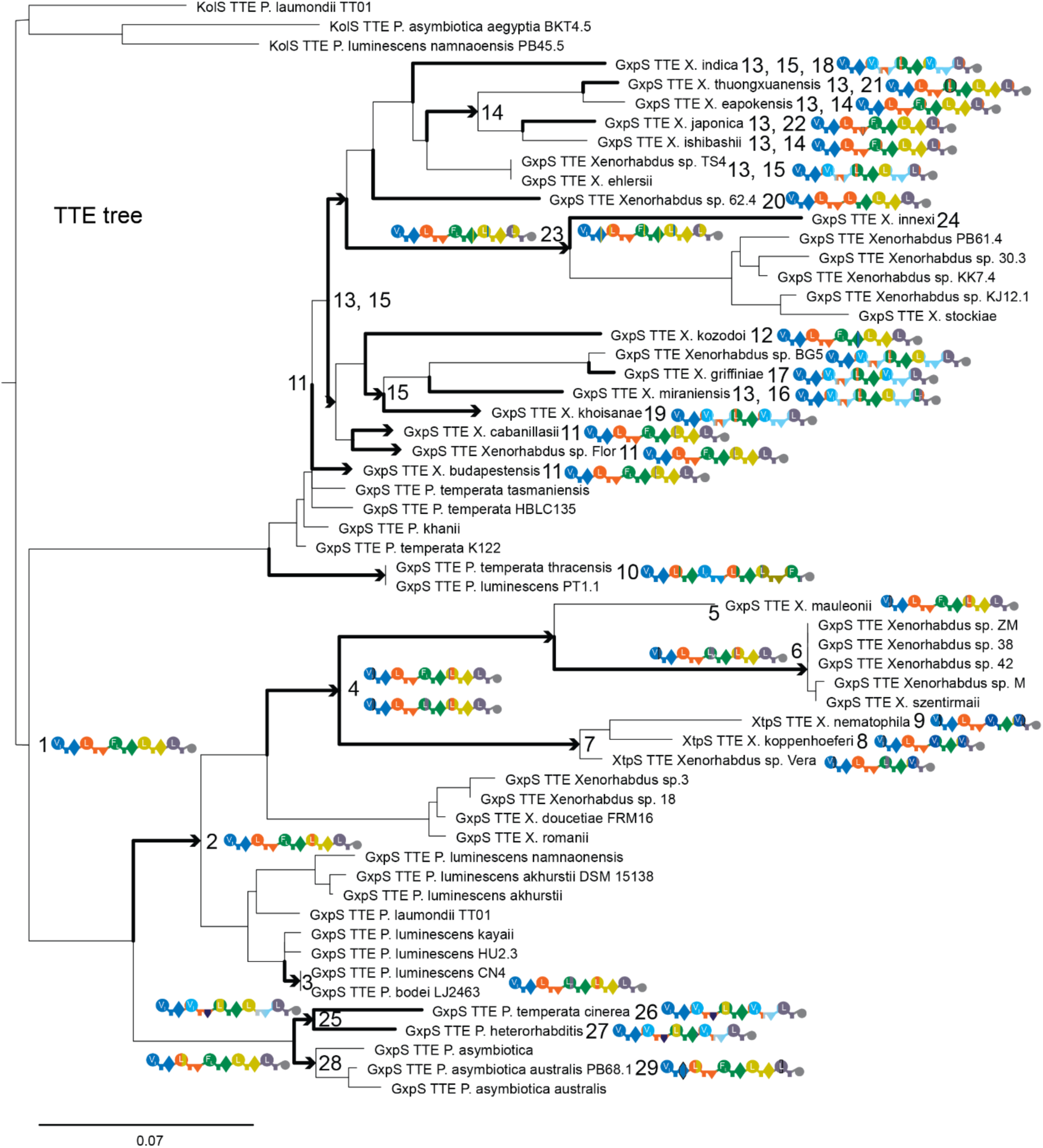
The TTE-domain tree of all GxpS from *Xenorhabdus* and *Photorhabdus*. All GxpS that have experienced recombinations are mapped on this tree and at the node where they emerged. The branches in arrow shape indicate the time point where a specific recombination occurred. The nodes are arbitrarily numbered from past to present and a more detailed explanation for every recombination is shown in Extended Data Table 1.

**Extended Data Table 1:**
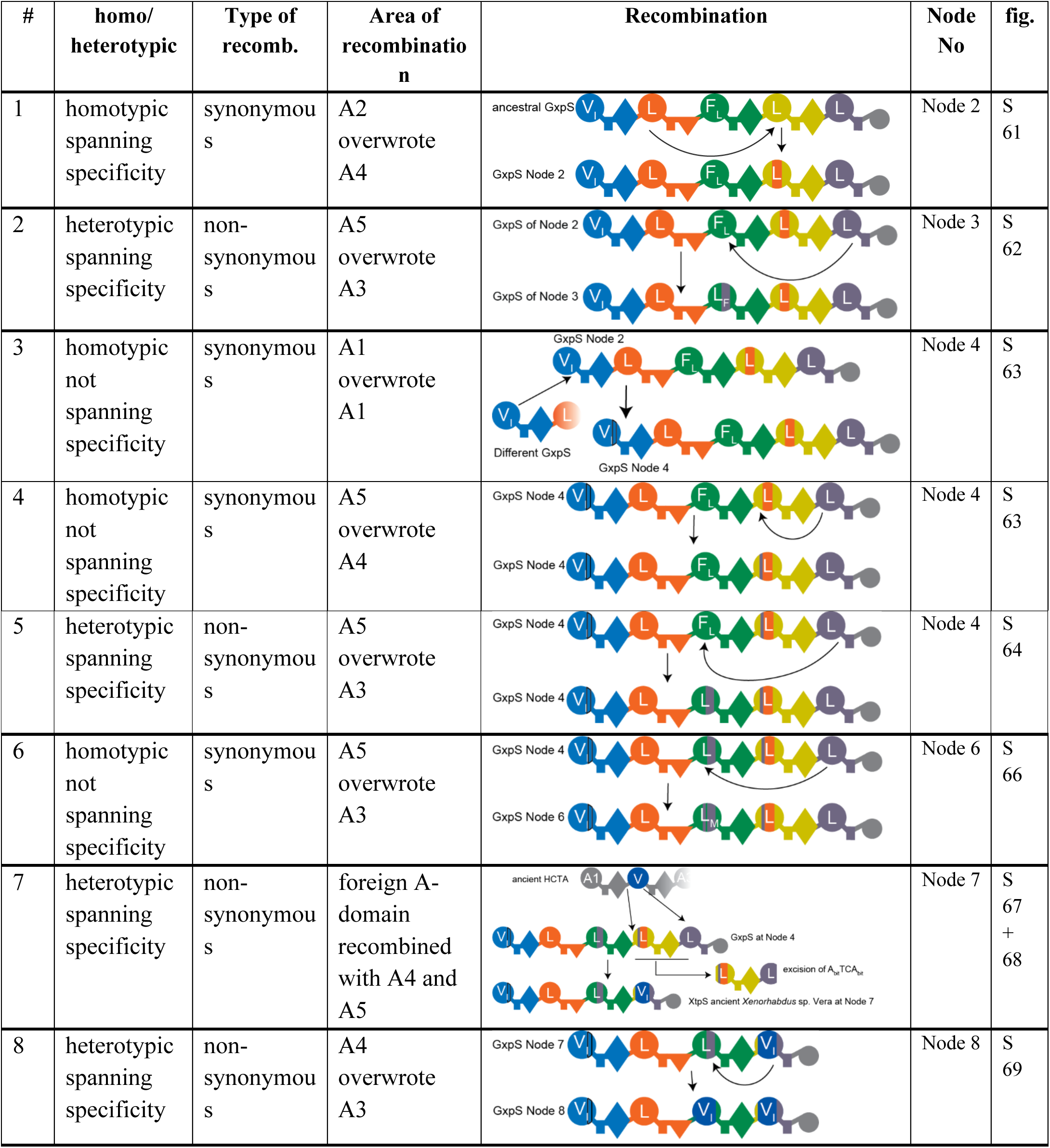
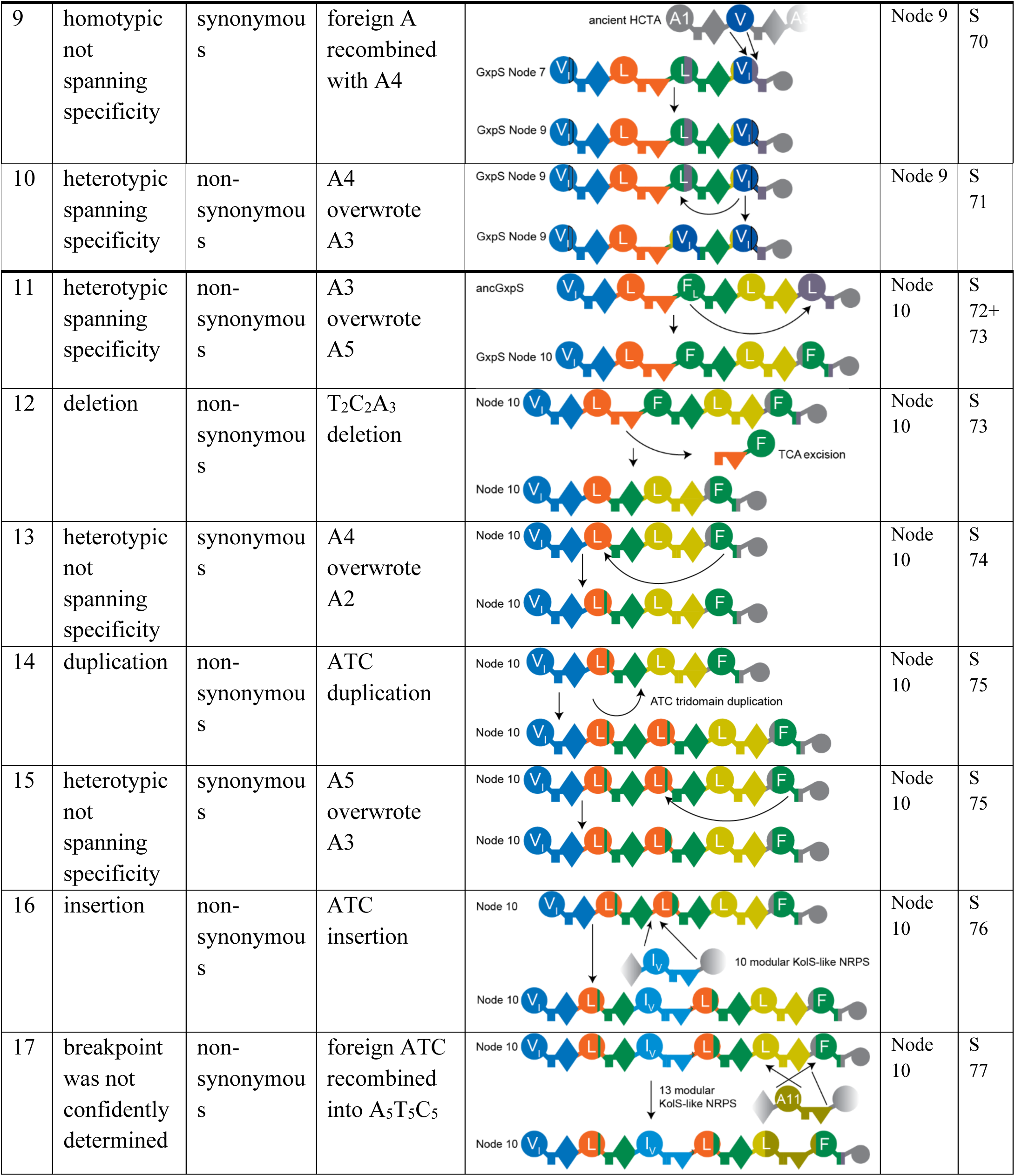
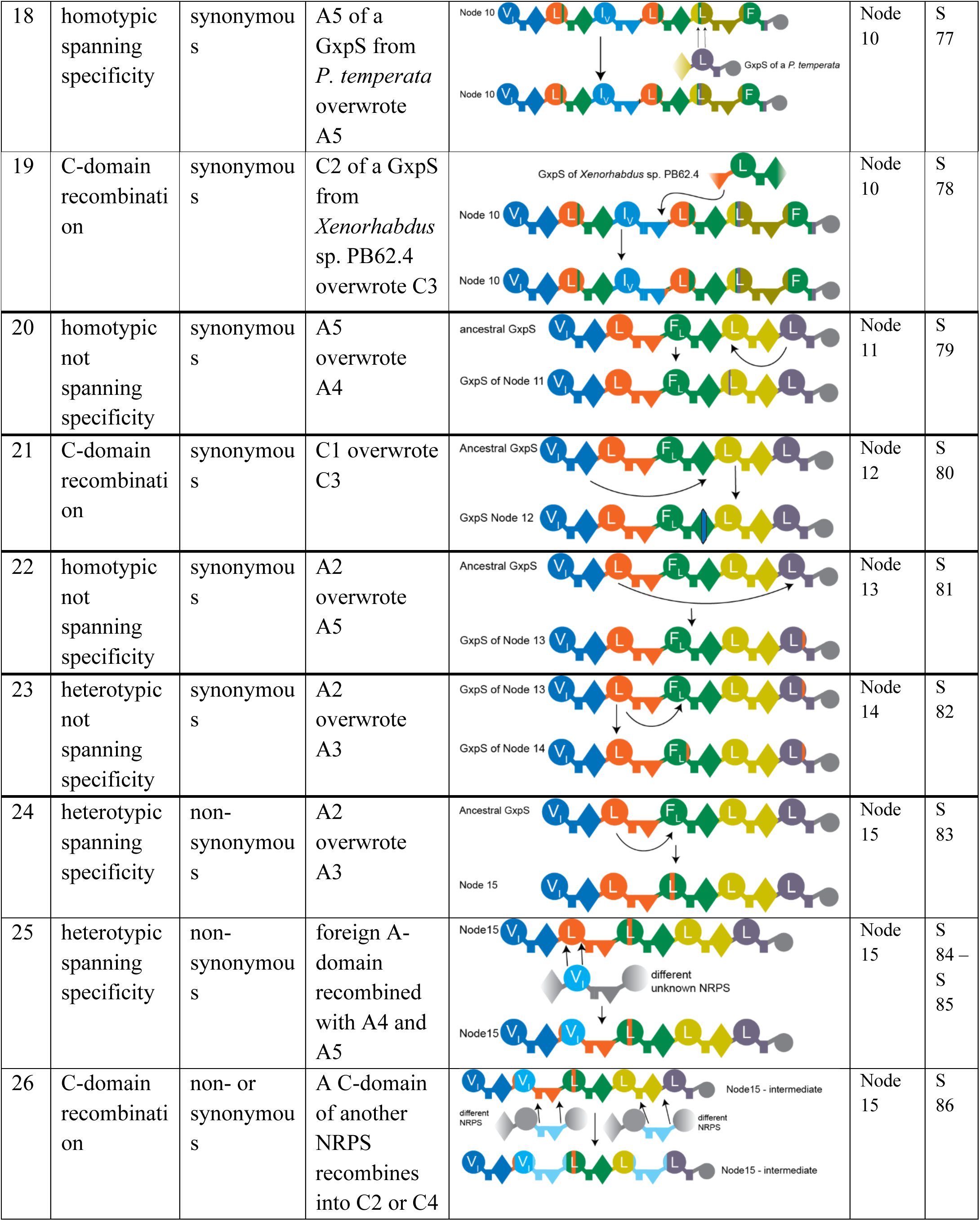
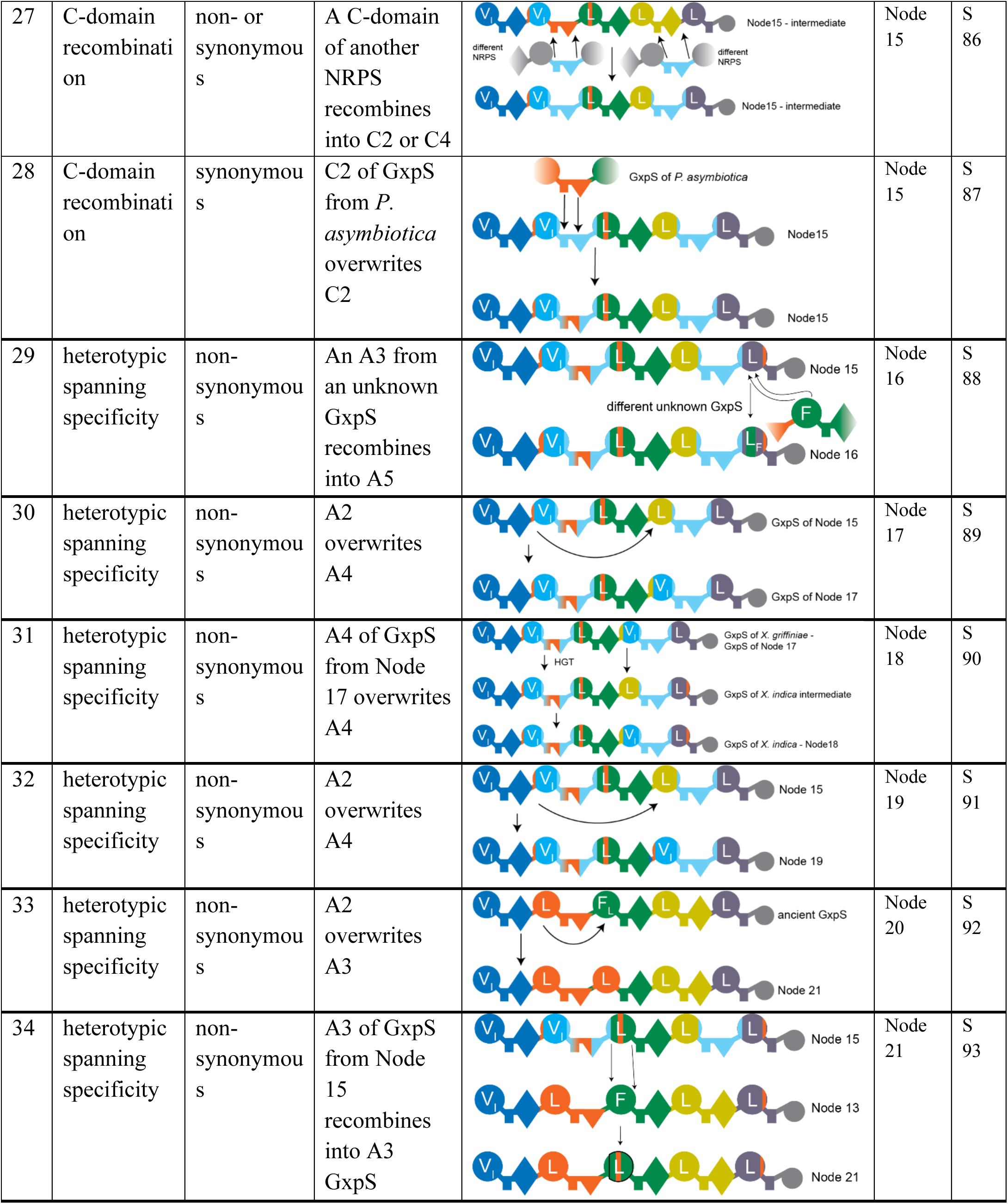
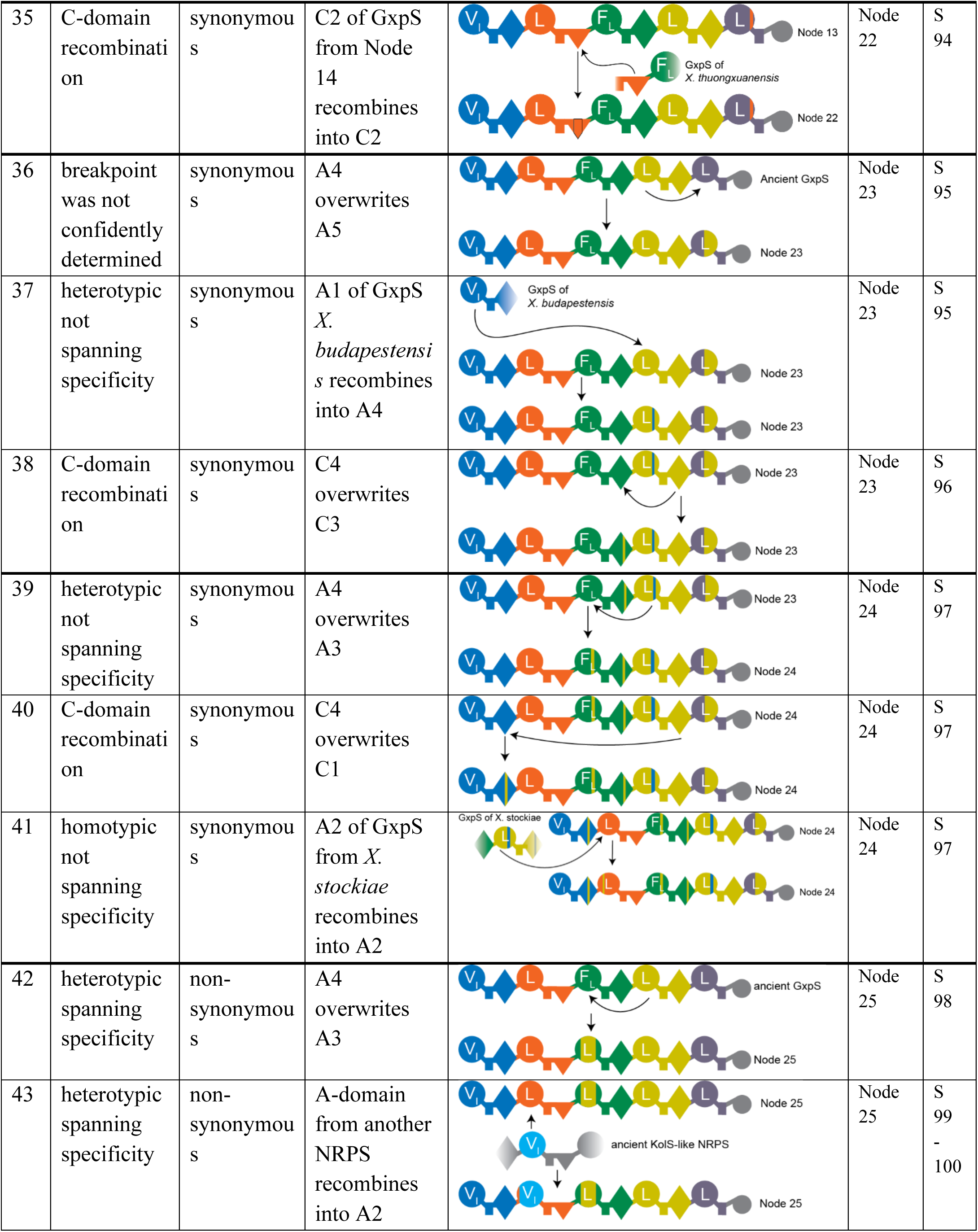
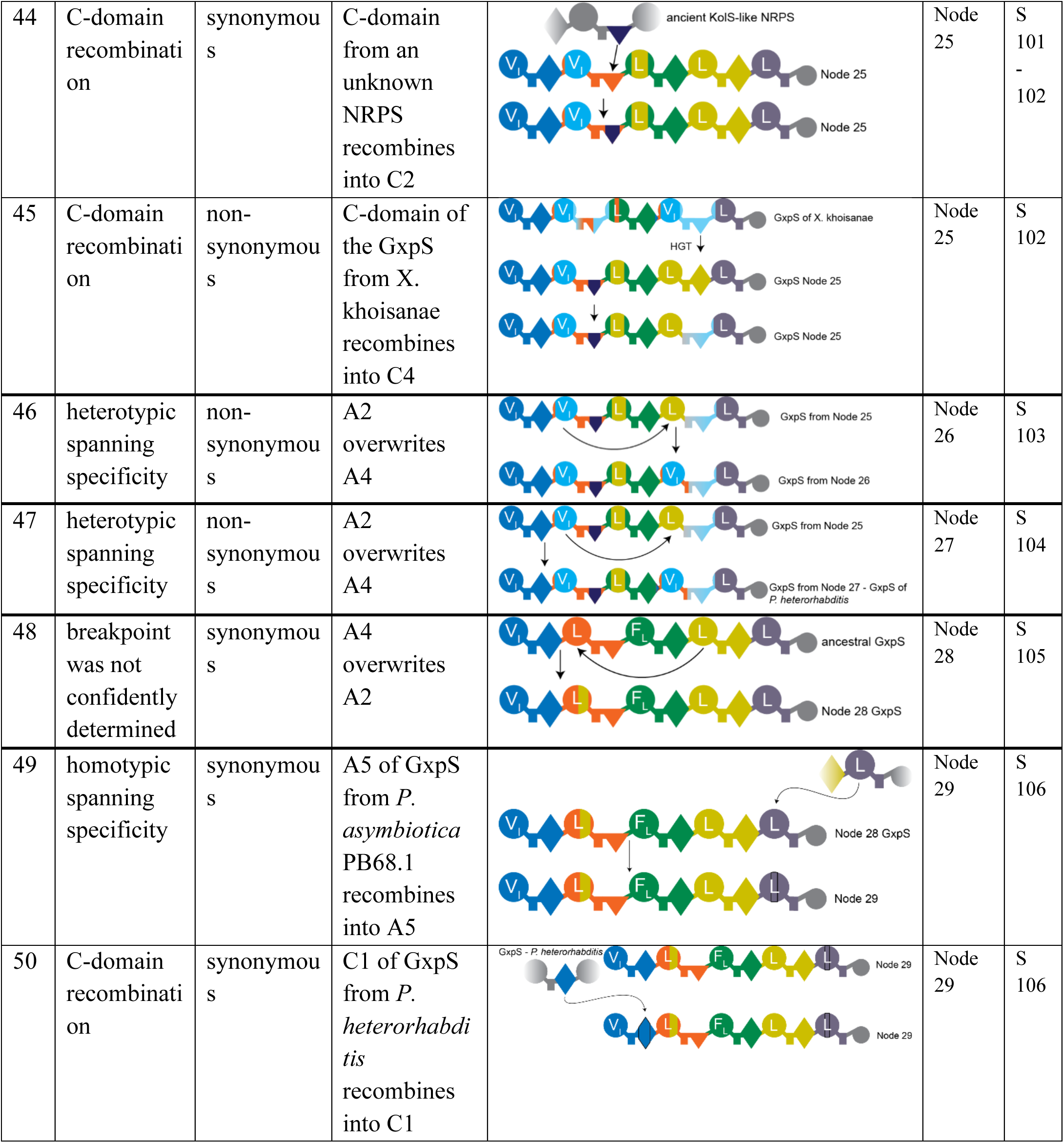
Summary of all recombinations.

## Notes

### Competing Interest Statement

The authors have declared no competing interest.

## References

1 Newman, D. J. & Cragg, G. M. Natural Products as Sources of New Drugs over the Nearly Four Decades from 01/1981 to 09/2019. J Nat Prod 83, 770–803 (2020). 10.1021/acs.jnatprod.9b01285

2 Baunach, M., Chowdhury, S., Stallforth, P. & Dittmann, E. The Landscape of Recombination Events That Create Nonribosomal Peptide Diversity. Mol Biol Evol 38, 2116–2130 (2021). 10.1093/molbev/msab015

3 Schierup, M. H. & Hein, J. Consequences of recombination on traditional phylogenetic analysis. Genetics 156, 879–891 (2000). 10.1093/genetics/156.2.879

4 Shi, Y. M. et al. Global analysis of biosynthetic gene clusters reveals conserved and unique natural products in entomopathogenic nematode-symbiotic bacteria. Nat Chem 14, 701–712 (2022). 10.1038/s41557-022-00923-2

5 Sussmuth, R. D. & Mainz, A. Nonribosomal Peptide Synthesis-Principles and Prospects. Angew Chem Int Ed Engl 56, 3770–3821 (2017). 10.1002/anie.201609079

6 Fischbach, M. A., Walsh, C. T. & Clardy, J. The evolution of gene collectives: How natural selection drives chemical innovation. Proc Natl Acad Sci U S A 105, 4601–4608 (2008). 10.1073/pnas.0709132105

7 Wang, Z. et al. A naturally inspired antibiotic to target multidrug-resistant pathogens. Nature 601, 606–611 (2022). 10.1038/s41586-021-04264-x

8 Kronheim, S. et al. A chemical defence against phage infection. Nature 564, 283–286 (2018). 10.1038/s41586-018-0767-x

9 Chevrette, M. G. et al. Evolutionary dynamics of natural product biosynthesis in bacteria. Nat Prod Rep 37, 566–599 (2020). 10.1039/c9np00048h

10 Waglechner, N., McArthur, A. G. & Wright, G. D. Phylogenetic reconciliation reveals the natural history of glycopeptide antibiotic biosynthesis and resistance. Nat Microbiol 4, 1862–1871 (2019). 10.1038/s41564-019-0531-5

11 Culp, E. J. et al. Evolution-guided discovery of antibiotics that inhibit peptidoglycan remodelling. Nature 578, 582–587 (2020). 10.1038/s41586-020-1990-9

12 Wang, Z., Koirala, B., Hernandez, Y., Zimmerman, M. & Brady, S. F. Bioinformatic prospecting and synthesis of a bifunctional lipopeptide antibiotic that evades resistance. Science 376, 991–996 (2022). 10.1126/science.abn4213

13 Xu, M. et al. Phylogeny-Informed Synthetic Biology Reveals Unprecedented Structural Novelty in Type V Glycopeptide Antibiotics. ACS Cent Sci 8, 615–626 (2022). 10.1021/acscentsci.1c01389

14 Shi, Y. M. & Bode, H. B. Chemical language and warfare of bacterial natural products in bacteria-nematode-insect interactions. Nat Prod Rep 35, 309–335 (2018). 10.1039/c7np00054e

15 Bozhuyuk, K. A. J. et al. Natural Products from Photorhabdus and Other Entomopathogenic Bacteria. Curr Top Microbiol Immunol 402, 55–79 (2017). 10.1007/82_2016_24

16 Nollmann, F. I. et al. Insect-specific production of new GameXPeptides in photorhabdus luminescens TTO1, widespread natural products in entomopathogenic bacteria. Chembiochem 16, 205–208 (2015). 10.1002/cbic.201402603

17 Bian, X., Plaza, A., Yan, F., Zhang, Y. & Muller, R. Rational and efficient site-directed mutagenesis of adenylation domain alters relative yields of luminmide derivatives in vivo. Biotechnol Bioeng 112, 1343–1353 (2015). 10.1002/bit.25560

18 Bode, H. B. et al. Determination of the absolute configuration of peptide natural products by using stable isotope labeling and mass spectrometry. Chemistry 18, 2342–2348 (2012). 10.1002/chem.201103479

19 Waterfield, N. R., Ciche, T. & Clarke, D. Photorhabdus and a host of hosts. Annu Rev Microbiol 63, 557–574 (2009). 10.1146/annurev.micro.091208.073507

20 Hedrick, P. W. Conservation genetics: where are we now? Trends Ecol Evol 16, 629–636 (2001). Doi 10.1016/S0169-5347(01)02282-0

21 Medema, M. H., Cimermancic, P., Sali, A., Takano, E. & Fischbach, M. A. A systematic computational analysis of biosynthetic gene cluster evolution: lessons for engineering biosynthesis. PLoS Comput Biol 10, e1004016 (2014). 10.1371/journal.pcbi.1004016

22 Bode, H. B. et al. Structure Elucidation and Activity of Kolossin A, the D-/L-Pentadecapeptide Product of a Giant Nonribosomal Peptide Synthetase. Angew Chem Int Ed Engl 54, 10352–10355 (2015). 10.1002/anie.201502835

23 Kegler, C. et al. Rapid determination of the amino acid configuration of xenotetrapeptide. Chembiochem 15, 826–828 (2014). 10.1002/cbic.201300602

24 Darby, C. A., Stolzer, M., Ropp, P. J., Barker, D. & Durand, D. Xenolog classification. Bioinformatics 33, 640–649 (2017). 10.1093/bioinformatics/btw686

25 Boussau, B., Gueguen, L. & Gouy, M. A mixture model and a hidden markov model to simultaneously detect recombination breakpoints and reconstruct phylogenies. Evol Bioinform Online 5, 67–79 (2009). 10.4137/ebo.s2242

26 Shishido, T. K. et al. Simultaneous Production of Anabaenopeptins and Namalides by the Cyanobacterium Nostoc sp. CENA543. ACS Chem Biol 12, 2746–2755 (2017). 10.1021/acschembio.7b00570

27 Pye, C. R., Bertin, M. J., Lokey, R. S., Gerwick, W. H. & Linington, R. G. Retrospective analysis of natural products provides insights for future discovery trends. Proc Natl Acad Sci U S A 114, 5601–5606 (2017). 10.1073/pnas.1614680114

28 Storz, J. F. Causes of molecular convergence and parallelism in protein evolution. Nat Rev Genet 17, 239–250 (2016). 10.1038/nrg.2016.11

29 Kimura, M. Evolutionary rate at the molecular level. Nature 217, 624–626 (1968). 10.1038/217624a0

30 Stachelhaus, T., Mootz, H. D. & Marahiel, M. A. The specificity-conferring code of adenylation domains in nonribosomal peptide synthetases. Chem Biol 6, 493–505 (1999). 10.1016/S1074-5521(99)80082-9

31 Martin, D. P. et al. RDP5: a computer program for analyzing recombination in, and removing signals of recombination from, nucleotide sequence datasets. Virus Evol 7, veaa087 (2021). 10.1093/ve/veaa087

32 Marahiel, M. A., Stachelhaus, T. & Mootz, H. D. Modular Peptide Synthetases Involved in Nonribosomal Peptide Synthesis. Chem Rev 97, 2651–2674 (1997). 10.1021/cr960029e

33 Chen, J. M., Cooper, D. N., Chuzhanova, N., Ferec, C. & Patrinos, G. P. Gene conversion: mechanisms, evolution and human disease. Nat Rev Genet 8, 762–775 (2007). 10.1038/nrg2193

## References

34 Zhao, L. & Bode, H. B. Production of a photohexapeptide library from entomopathogenic Photorhabdus asymbiotica PB68.1. Org Biomol Chem 17, 7858–7862 (2019). 10.1039/c9ob01489f

35 Bode, E. et al. Promoter Activation in Deltahfq Mutants as an Efficient Tool for Specialized Metabolite Production Enabling Direct Bioactivity Testing. Angew Chem Int Ed Engl 58, 18957–18963 (2019). 10.1002/anie.201910563

36 Harris, C. R. et al. Array programming with NumPy. Nature 585, 357–362 (2020). 10.1038/s41586-020-2649-2

37 Virtanen, P. et al. SciPy 1.0: fundamental algorithms for scientific computing in Python. Nat Methods 17, 261–272 (2020). 10.1038/s41592-019-0686-2

38 Hunter, J. D. Matplotlib: A 2D graphics environment. Computing in science & engineering 9, 90–95 (2007).

39 Burnham, K. P. & Anderson, D. R. Model selection and multimodel inference: a practical information-theoretic approach. (Springer, 2002).

40 Goldfarb, T. et al. NCBI RefSeq: reference sequence standards through 25 years of curation and annotation. Nucleic Acids Res 53, D243–D257 (2025). 10.1093/nar/gkae1038

41 Katoh, K. & Standley, D. M. MAFFT multiple sequence alignment software version 7: improvements in performance and usability. Mol Biol Evol 30, 772–780 (2013). 10.1093/molbev/mst010

42 Capella-Gutierrez, S., Silla-Martinez, J. M. & Gabaldon, T. trimAl: a tool for automated alignment trimming in large-scale phylogenetic analyses. Bioinformatics 25, 1972–1973 (2009). 10.1093/bioinformatics/btp348

43 Kalyaanamoorthy, S., Minh, B. Q., Wong, T. K. F., von Haeseler, A. & Jermiin, L. S. ModelFinder: fast model selection for accurate phylogenetic estimates. Nat Methods 14, 587–589 (2017). 10.1038/nmeth.4285

44 Jones, D. T., Taylor, W. R. & Thornton, J. M. The rapid generation of mutation data matrices from protein sequences. Comput Appl Biosci 8, 275–282 (1992). 10.1093/bioinformatics/8.3.275

45 Guindon, S. & Gascuel, O. A simple, fast, and accurate algorithm to estimate large phylogenies by maximum likelihood. Syst Biol 52, 696–704 (2003). 10.1080/10635150390235520

46 Wong, T. K. F. et al. MAST: Phylogenetic Inference with Mixtures Across Sites and Trees. Syst Biol 73, 375–391 (2024). 10.1093/sysbio/syae008

47. Larsson, A. AliView: a fast and lightweight alignment viewer and editor for large datasets. Bioinformatics 30, 3276-3278 (2014). 10.1093/bioinformatics/btu531

48 Minh, B. Q. et al. IQ-TREE 2: New Models and Efficient Methods for Phylogenetic Inference in the Genomic Era. Mol Biol Evol 37, 1530–1534 (2020). 10.1093/molbev/msaa015

49 Hoang, D. T., Chernomor, O., von Haeseler, A., Minh, B. Q. & Vinh, L. S. UFBoot2: Improving the Ultrafast Bootstrap Approximation. Mol Biol Evol 35, 518–522 (2018). 10.1093/molbev/msx281

50 Guindon, S. et al. New algorithms and methods to estimate maximum-likelihood phylogenies: assessing the performance of PhyML 3.0. Syst Biol 59, 307–321 (2010). 10.1093/sysbio/syq010

51 Lee, M. D. GToTree: a user-friendly workflow for phylogenomics. Bioinformatics 35, 4162–4164 (2019). 10.1093/bioinformatics/btz188

52 Hyatt, D. et al. Prodigal: prokaryotic gene recognition and translation initiation site identification. BMC Bioinformatics 11, 119 (2010). 10.1186/1471-2105-11-119

53 Eddy, S. R. Accelerated Profile HMM Searches. PLoS Comput Biol 7, e1002195 (2011). 10.1371/journal.pcbi.1002195

54 Edgar, R. C. Muscle5: High-accuracy alignment ensembles enable unbiased assessments of sequence homology and phylogeny. Nat Commun 13, 6968 (2022). 10.1038/s41467-022-34630-w

55 Price, M. N., Dehal, P. S. & Arkin, A. P. FastTree 2––approximately maximum-likelihood trees for large alignments. PLoS One 5, e9490 (2010). 10.1371/journal.pone.0009490

56 Shen, W. & Ren, H. TaxonKit: A practical and efficient NCBI taxonomy toolkit. J Genet Genomics 48, 844–850 (2021). 10.1016/j.jgg.2021.03.006

57 Branger, M. & Leclercq, S. O. GenoFig: a user-friendly application for the visualization and comparison of genomic regions. Bioinformatics 40 (2024). 10.1093/bioinformatics/btae372

58 Martin, D. & Rybicki, E. RDP: detection of recombination amongst aligned sequences. Bioinformatics 16, 562–563 (2000). 10.1093/bioinformatics/16.6.562

59 Padidam, M., Sawyer, S. & Fauquet, C. M. Possible emergence of new geminiviruses by frequent recombination. Virology 265, 218–225 (1999). 10.1006/viro.1999.0056

60 Smith, J. M. Analyzing the mosaic structure of genes. J Mol Evol 34, 126–129 (1992). 10.1007/BF00182389

61 Martin, D. P., Posada, D., Crandall, K. A. & Williamson, C. A modified bootscan algorithm for automated identification of recombinant sequences and recombination breakpoints. AIDS Res Hum Retroviruses 21, 98–102 (2005). 10.1089/aid.2005.21.98

62 Gibbs, M. J., Armstrong, J. S. & Gibbs, A. J. Sister-scanning: a Monte Carlo procedure for assessing signals in recombinant sequences. Bioinformatics 16, 573–582 (2000). 10.1093/bioinformatics/16.7.573

63 Posada, D. & Crandall, K. A. Evaluation of methods for detecting recombination from DNA sequences: computer simulations. Proc Natl Acad Sci U S A 98, 13757–13762 (2001). 10.1073/pnas.241370698

64 Lam, H. M., Ratmann, O. & Boni, M. F. Improved Algorithmic Complexity for the 3SEQ Recombination Detection Algorithm. Mol Biol Evol 35, 247–251 (2018). 10.1093/molbev/msx263

65 Conti, E., Stachelhaus, T., Marahiel, M. A. & Brick, P. Structural basis for the activation of phenylalanine in the non-ribosomal biosynthesis of gramicidin S. EMBO J 16, 4174–4183 (1997). 10.1093/emboj/16.14.4174

