## Supplementary Material for "Drift, selection and convergence in the evolution of a nonribosomal peptide"

Affiliations:

[3]School of Life Sciences and Engineering, Major in Plant Medicals, Gyeongkuk National University, Andong 36729, Korea

**Table S1. Detected compounds in this work.** Small, italic letters indicate D-amino acids; ***i*** = D-*allo*-Ile; *p*-aminophenylalanine = PAPA and its methylated version as mPAPA and both are put in brackets.

| Peptide | Sequence | MS exp. [M+H]$^+$ | MS calc. [M+H]$^+$ | Molecular ion formular | □ppm | Reference |
|---|---|---|---|---|---|---|
| **1** GXPA | cyclo(*v*L*fl*L) | 586.3967 | 586.3963 | $C_{32}H_{51}N_5O_5$ | 0.7 | [1] |
| **2** XTPA | cyclo(*v*L*v*V) | 411.2958 | 411.2965 | $C_{21}H_{38}N_4O_4$ | -1.7 | [2] |
| **3** XTPD | cyclo(*v*L*l*V) | 425.3116 | 425.3122 | $C_{22}H_{40}N_4O_4$ | -1.4 | This work |
| **4** GXPR | cyclo(*v*V*l*VL) | 524.3811 | 524.3806 | $C_{27}H_{49}N_5O_6$ | 0.9 | This work |
| **5** GXPN | cyclo(*v*V*l*LL) | 538.3964 | 538.3962 | $C_{28}H_{51}N_5O_5$ | 0.3 | This work |
| **6** GXPC | cyclo(*v*L*ll*L) | 552.4131 | 552.4119 | $C_{29}H_{53}N_5O_5$ | 0.3 | [1] |
| **7** HEXA | cyclo(*vl*I*l*LF) | 699.4802 | 699.4803 | $C_{38}H_{62}N_6O_6$ | -0.1 | This work |
| **8** XTPB | cyclo(***i***L*v*V) | 425.3115 | 425.3122 | $C_{22}H_{40}N_4O_4$ | -1.6 | This work |
| **9** XTPE | cyclo(***i***L*l*V) | 439.3268 | 439.3278 | $C_{23}H_{42}N_4O_4$ | -2.2 | This work |
| **10** XTPJ | cyclo(*v*V*l*V) | 411.2966 | 411.2965 | $C_{21}H_{38}N_4O_4$ | 0.2 | This work |
| **11** XTPG | cyclo(*v*V*l*L) | 425.3123 | 425.3122 | $C_{22}H_{40}N_4O_4$ | -0.2 | This work |
| **12** GXPD | cyclo(***i***L*ll*L) | 566.4258 | 566.4275 | $C_{30}H_{55}N_5O_5$ | 3.0 | [3] |
| **13** HEXB | cyclo(***i****l*I*l*LF) | 713.4950 | 713.4960 | $C_{39}H_{64}N_6O_6$ | 1.4 | This work |
| **14** GXPB | cyclo(***i***L*fl*L) | 600.4110 | 600.4119 | $C_{33}H_{53}N_5O_5$ | 1.5 | [3] |
| **15** GXPL | cyclo(*v*L*vl*L) | 538.3951 | 538.3962 | $C_{28}H_{51}N_5O_5$ | 2.0 | [3] |
| **16** GXPK | cyclo(*v*L*ml*L) | 570.3675 | 570.3684 | $C_{28}H_{51}N_5O_5S$ | -1.6 | [3] |
| **17** GXPM | cyclo(*v*L*yl*L) | 602.3911 | 602.3912 | $C_{32}H_{51}N_5O_6$ | 0.2 | [3] |

| | | | | | | |
|---|---|---|---|---|---|---|
| **18**, **19** | L*l*L, *ll*L | 358.2701 | 358.2700 | $C_{18}H_{35}N_3O_4$ | 0.3 | This work |
| **20**, **21** | F*l*L, *fl*L | 392.2543 | 392.2543 | $C_{21}H_{33}N_3O_4$ | 0 | This work |
| **22** | L*ll*L | 471.3541 | 471.3541 | $C_{24}H_{46}N_4O_5$ | 0 | This work |
| **23** | *v*L*l*L | 491.3228 | 491.3228 | $C_{26}H_{42}N_4O_5$ | 0 | This work |
| **24** | L*fl*L | 505.3388 | 505.3384 | $C_{27}H_{44}N_4O_5$ | 0.8 | This work |
| **25** XTPH | cyclo(***i***V*l*L) | 439.3280 | 439.3279 | $C_{23}H_{42}N_4O_4$ | 0.2 | This work |
| **26** XTPI | cyclo(***i***I*l*L) | 453.3438 | 453.3435 | $C_{24}H_{44}N_4O_4$ | 0.7 | This work |
| **27** GXPO | cyclo(***i***V*l*LL) | 552.4118 | 552.4119 | $C_{29}H_{53}N_5O_5$ | -0.2 | This work |
| **28** GXPP | cyclo(***i***I*l*LL) | 566.4271 | 566.4276 | $C_{30}H_{55}N_5O_5$ | -0.9 | This work |
| **29** GXPQ | cyclo(*v*V*l*LF) | 572.3810 | 572.3810 | $C_{31}H_{49}N_5O_5$ | 0 | This work |
| **30** XTPK | cyclo(***i***V*l*V) | 425.3124 | 425.3122 | $C_{22}H_{40}N_4O_4$ | 0.5 | This work |
| **31** XTPL | cyclo(***i***I*l*V) or cyclo(***i***V*l*I) | 439.3281 | 439.3279 | $C_{23}H_{42}N_4O_4$ | 0.5 | This work |
| **32** GXPS | cyclo(***i***V*l*VL) | 538.3967 | 538.3963 | $C_{28}H_{51}N_5O_5$ | 0.7 | This work |
| **33** HEXC | cyclo(*vl*V*l*LF) | 685.4642 | 685.4647 | $C_{37}H_{60}N_6O_6$ | -0.7 | This work |
| **34** XTPC | cyclo(***i***L*v*I) | 439.3280 | 439.3279 | $C_{23}H_{42}N_4O_4$ | 0.2 | This work |
| **35** XTPF | cyclo(*v*L*m*V) | 443.2688 | 443.2687 | $C_{21}H_{38}N_4O_4S$ | 0.2 | This work |
| **36** GXPE | cyclo(*v*L[*papa*]*l*L) | 601.4065 | 601.4071 | $C_{32}H_{52}N_6O_5$ | -1.0 | [4] |
| **37** GXPF | cyclo(***i***L[*papa*]*l*L) | 615.4224 | 615.4228 | $C_{33}H_{54}N_6O_5$ | -0.6 | This work |
| **38** HEXD | cyclo(*vl*I*l*L[PAPA]) | 714.4909 | 714.4913 | $C_{38}H_{63}N_6O_5$ | -0.5 | This work |
| **39** HEXE | cyclo(*il*I*l*L[PAPA]) | 728.5057 | 728.5069 | $C_{39}H_{65}N_7O_6$ | -1.6 | This work |

| | | | | | | |
|---|---|---|---|---|---|---|
| **42** GXPG | cyclo(*v*L[*mpapa*]*l*L) | 615.4213 | 615.4228 | $C_{33}H_{54}N_6O_5$ | -1.9 | [4] |
| **43** GXPH | cyclo(***i***L[m*papa*]*l*L) | 629.4368 | 629.4385 | $C_{34}H_{56}N_6O_5$ | -2.7 | This work |
| **47** HEXF | cyclo(*vl*I*l*L[mPAPA]) | 728.5045 | 728.5069 | $C_{39}H_{65}N_7O_6$ | -3.3 | This work |
| **48** HEXG | cyclo(***i****l*I*l*L[mPAPA[) | 742.5259 | 742.5226 | $C_{40}H_{67}N_7O_6$ | 4.4 | This work |

**Table S2. Further detected compounds in this work.** These compounds are not naturally occurring compounds. However, these compounds artificially are produced after addition of PAPA. Small, italic letters indicate D-amino acids; **i** = D-*allo*-Ile; *p*-aminophenylalanine = PAPA and its methylated version as mPAPA and both are put in brackets.

| **Peptide** | **Sequence** | **MS exp. [M+H]$^+$** | **MS calc. [M+H]$^+$** | **Molecular ion formular** | **Δppm** | **Reference** |
|---|---|---|---|---|---|---|
| **40** XTPM | cyclo(*v*V[*papa*]L) | 474.3058 | 474.3075 | $C_{25}H_{39}N_5O_4$ | -3.6 | This work |
| **41** GXPT | cyclo(*v*V*l*L[PAPA]) | 587.3916 | 587.3915 | $C_{31}H_{50}N_6O_5$ | 0.2 | This work |
| **44** XTPN | cyclo(*v*V[*mpapa*]L) | 488.3219 | 488.3231 | $C_{26}H_{41}N_5O_4$ | -2.5 | This work |
| **45** XTPO | cyclo(*v*V[*mpapa*]V) | 474.3068 | 474.3075 | $C_{25}H_{39}N_5O_4$ | -1.4 | This work |
| **46** XTPP | cyclo(***i***V[*mpapa*]V) | 488.3212 | 488.3231 | $C_{26}H_{41}N_5O_4$ | -3.9 | This work |
| **49** GXPU | cyclo(*v*V*l*L[mPAPA]) | 601.4054 | 601.4071 | $C_{32}H_{52}N_6O_5$ | -2.8 | This work |

**Table S3. Selection test no. 1 summarizes homo- and heterotypic recombinations.** Recombinations are divided into recombinations that span or do not span the specificity conferring area. The recombination that has changed phenylalanine to leucine at node number 4, could also have occurred at Node number 6 and 7 using two times the same borders. This scenario would not include ILS. The recombination at Node 13 includes all its descendants on the TTE-domain tree. The only exception is *X. miraniensis*. Likely, a HGT occurred and lead to this recombination. An alternative hypothesis would be ILS that causes this tree topology. In this table, we counted these two recombinations additionally. A two-tailed Fisher's exact test has been conducted. The test, that considers the relationship between columns and rows, is statistically significant. The two-tailed p-value equals 0.0053.

| recombinations | **spanning specificity** | **not spanning specificity** |
|---|---|---|
| **homotypic** | 3<br>No. 1, 18, 49 | 8<br>No. 3, 4, 6, 9, 20, 2x22, 41 |
| **heterotypic** | 19<br>No. 2, 2x5, 7, 8, 10, 11, 24, 25, 29, 30, 31, 32, 33, 34, 42, 43, 46, 47 | 5<br>No. 13, 15, 23, 37, 39 |

**Table S4. Selection test no. 2 that summarizes homo- and heterotypic recombinations** Recombinations are divided into recombinations that span or do not span the specificity conferring area. Here, recombination at Node 4 and Node 13 is interpreted to lead to a polymorphism within the bacterial population that causes this tree topology. A two-tailed Fisher's exact test has been conducted. The test, that considers the relationship between columns and rows, is statistically significant. The two-tailed p-value equals 0.0164.

| recombinations | **spanning specificity** | **not spanning specificity** |
|---|---|---|
| **homotypic** | 3<br>No. 1, 18, 49 | 7<br>No. 3, 4, 6, 9, 20, 22, 41 |
| **heterotypic** | 18<br>No. 2, 5, 7, 8, 10, 11, 24, 25, 29, 30, 31, 32, 33, 34, 42, 43, 46, 47 | 5<br>No. 13, 15, 23, 37, 39 |

**Table S5. Selection test no. 3 summarizes homo- and heterotypic recombinations.** Recombinations are divided into recombinations that span or do not span the specificity conferring area. The recombination that has changed phenylalanine to leucine at node number 4, could also have occurred at Node number 6 and 7 using two times the same borders. This scenario would not include ILS. The recombination at Node 13 includes all its descendants on the TTE-domain tree. The only exception is *X. miraniensis*. Likely, a HGT occurred and lead to this recombination. An alternative hypothesis would be ILS that causes this tree topology. In this table, we counted these two recombinations additionally. We also counted recombination no. 36 and 48 even though we are not really certain if the touched the specificity area. A two-tailed Fisher's exact test was conducted. The two-tailed p-value equals 0.0282.

| recombinations | **spanning specificity** | **not spanning specificity** |
|---|---|---|
| **homotypic** | 5<br>No. 1, 18, 36, 48, 49 | 8<br>No. 3, 4, 6, 9, 20, 2x22, 41 |
| **heterotypic** | 19<br>No. 2, 2x5, 7, 8, 10, 11, 24, 25, 29, 30, 31, 32, 33, 34, 42, 43, 46, 47 | 5<br>No. 13, 15, 23, 37, 39 |

**Table S6. Selection test no. 4 that summarizes homo- and heterotypic recombinations** Recombinations are divided into recombinations that span or do not span the specificity conferring area. Here, recombination at Node 4 and Node 13 is interpreted to lead to a polymorphism within the bacterial population that causes this tree topology. We also counted recombination no. 36 and 48 even though we are not certain if these cover the specificity area. A two-tailed Fisher's exact test was conducted. The two-tailed p-value equals 0.0587.

| recombinations | **spanning specificity** | **not spanning specificity** |
|---|---|---|
| **homotypic** | 5<br>No. 1, 18, 36, 48, 49 | 7<br>No. 3, 4, 6, 9, 20, 22, 41 |
| **heterotypic** | 18<br>No. 2, 5, 7, 8, 10, 11, 24, 25, 29, 30, 31, 32, 33, 34, 42, 43, 46, 47 | 5<br>No. 13, 15, 23, 37, 39 |

**Table S7**. Strains used in this work.

| **Strain** | **Genotype/ NRPS** | **Reference** |
| --- | --- | --- |
| *E. coli* DH10B::*mtaA* | F_mcrA (*mrr-hsd*RMS-*mcr*BC), 80*lac*ZΔ, M15, Δ*lac*X74 *rec*A1 *end*A1 *ara*D 139Δ (*ara*, *leu*)7697 *gal*U *gal*K λ *rps*L (*Strr*) *nupG* / -, *mtaA* from pCK_*mtaA*Δ*entD* / - | [5] |
| *E. coli* DH10B$^{penta}$ | DH10B Δ*ilvE*, Δ*tyrB*, Δ*aspC*, Δ*avtA*, Δ*yfbQ*; FRT scar | [2] |
| *P. laumondii* TT01 | WT (*gxpS*) | DSMZ |
| *P. namnaonensis* PB 45.5 | WT (*gxpS*) | [6] |
| *P. tasmaniensis* DSM 22387 | WT (*gxpS*) | DSMZ |
| *P. bodei* LJ24-63 | WT (*gxpS*) | [6] |
| *P. cinerea* DMS 19724 | WT (*gxpS*) | DSMZ |
| *P. thracensis* DSM 15199 | WT (*gxpS*) | DSMZ |
| *X. doucetiae* FRM16 | WT (*gxpS*) | [7] |
| *X. innexi* DSM 16336 | WT (*gxpS*) | DSMZ |
| *Xenorhabdus* sp. PB62.4 | WT (*gxpS*) | This lab |
| *X. szentirmaii* DSM 16338 | WT (*gxpS*) | DSMZ |
| X. ehlersii DSM 16337 | WT (*gxpS*) | DSMZ |
| *X. miraniensis* DSM 17902 | WT (*gxpS*) | DSMZ |
| *X. khoisanae* DSM 25463 | WT (*gxpS*) | DSMZ |
| *Xenorhabdus* sp. Vera | WT (*xtpS*)<br>Genetic information was retrieved from NCBI [8] and the nucleotide sequence was synthesized by Genscript Biotech Corp. | [9] |

**Table S8.** Primer and templates used in this work to generate plasmids. Bp length of PCR products are shown below the template.

| Plasmids | Oligonucleotides | Sequence (5' to 3') | Template Product size in bp |
|---|---|---|---|
| pSK4 – *gxpS* of *P. namnaonensis* in pCOLA | oSK1 | TTGGGCTAACAGGAGGAATTCCATGAAAGATAGCATGGCTAAAAAGG | *P. namnaonensis* gDNA 15654 |
| | oSK2 | CAGCGGTGGCAGCAGCCTAGGTTAATTACAGCGCCTCCGCTTCAC | |
| | pCOLA-FW | TAATTAACCTAGGCTGCTGC | pMY-e1-0000 3020 |
| | pCOLA-RV | CATGGAATTCCTCCTGTTAGC | |
| pSK3 – *gxpS* of *X. ehlersii* in pCOLA | oSK3 | GCTAACAGGAGGAATTCCATGAAAGACAGTATTACCAGCAAGG | *X. ehlersii* gDNA 15480 |
| | oSK4 | CAGCGGTGGCAGCAGCCTAGGTTAATTACACCACCTCCGCTTCAC | |
| | pCOLA-FW | TAATTAACCTAGGCTGCTGC | pMY-e1-0000 3020 |
| | pCOLA-RV | CATGGAATTCCTCCTGTTAGC | |
| pSK5 – *gxpS* of *X. innexi* in pCOLA | oSK7 | GCTAACAGGAGGAATTCCATGAAAGATAGTATTACCAGTAAAGGA | *X. innexi* gDNA 15720 |
| | oSK8 | GGCAGCAGCCTAGGTTAATTACACCGCTTCTGTTTCATACG | |
| | pCOLA-FW | TAATTAACCTAGGCTGCTGC | pMY-e1-0000 3020 |
| | pCOLA-RV | CATGGAATTCCTCCTGTTAGC | |
| pMY-e1-0000_GG – pCOLA vector with BsaI sites and *sacB* cassette | oSK9 | TAACCTAGGCTGCTGCCACC | pMY-e1-0000 3016 |
| | pCOLA-RV | CATGGAATTCCTCCTGTTAGC | |
| | oSK21_FW | TTTTTTTGGGCTAACAGGAGGAATTCCATGAGAGACCGTAGCCAATCGCGCGGGTTTGTTACTG | pAR18 |
| | oSK22_R | ATTGCTCAGCGGTGGCAGCAGCCTAGGTTACTCAAGAGACCGTAGCCTTATTTGTTAACTGTTAATTGTCCTTGTTCAAGG | |
| pSK12 – *gxpS* of *X. khoisana* | oSK10 | GCTAACAGGAGGAATTCCATGAAAGACAGTATTACCGGCAAG | *X. khoisanae* gDNA 15534 |
| | oSK11 | GGCAGCAGCCTAGGTTAATTAATTTAACGCCATACTTAAGGTATTAGC | |

| | | | |
|---|---|---|---|
| *e* in pCOLA | pCOLA-FW | TAATTAACCTAGGCTGCTGC | pMY-e1-0000 3020 |
| | pCOLA-RV | CATGGAATTCCTCCTGTTAGC | |
| pSK6 – *gxpS* of *P. cinerea* in pCOLA | oSK12 | GCTAACAGGAGGAATTCCATGAAAGAGAGTATCGCT AAAAAGGG | *P. cinerea* gDNA 15477 |
| | oSK13 | GGGATTCAAATAAGGTGGTCAG | |
| | oSK14 | CTGACCACCTTATTTGAATCCC | |
| | oSK15 | GGCAGCAGCCTAGGTTAATTACAGCGCTTCTGCCTCA CAATT | |
| | pCOLA-FW | TAATTAACCTAGGCTGCTGC | pMY-e1-0000 3020 |
| | pCOLA-RV | CATGGAATTCCTCCTGTTAGC | |
| pSK8 – *gxpS* of *P. thracensis* in pCOLA | oSK16 | GCTAACAGGAGGAATTCCATGAAAGACAGTATTACC AGAAAGGA | *P. thracensis* gDNA 8252 |
| | oSK17 | CGTATCGCTTCAGCAAAGGC | |
| | oSK18 | GCCTTTGCTGAAGCGATACG | *P. thracensis* gDNA 10437 |
| | oSK19 | GGCAGCAGCCTAGGTTAATTACACCGCCTCCGCTTCA C | |
| | pCOLA-FW | TAATTAACCTAGGCTGCTGC | pMY-e1-0000 3020 |
| | pCOLA-RV | CATGGAATTCCTCCTGTTAGC | |
| pSK7 – *gxpS* of *Xenorhabdus* sp. PB62.4 in pCOLA | oSK20 | GCTAACAGGAGGAATTCCATGAAAGACAGTATTACC AGCAAGG | *Xenorhabdus* sp. PB62.4 gDNA 5083 |
| | oSK21 | ATCTATCCCTTCCCTCATTGCG | |
| | oSK22 | CGCAATGAGGGAAGGGATAGAT | *Xenorhabdus* sp. PB62.4 gDNA 10539 |
| | oSK23 | GGCAGCAGCCTAGGTTAATTACACCGCCTCTGCTTCA C | |
| | pCOLA-FW | TAATTAACCTAGGCTGCTGC | pMY-e1-0000 3020 |
| | pCOLA-RV | CATGGAATTCCTCCTGTTAGC | |
| pSK32 – corancA 3 in pUC57-BsaI- | oSK57 | GCGAATGCATCTAGATATACGGTCTCTCTGAACGGA AGTTATTGCTGGAAACCT | corancA3 1593 |
| | oSK58 | CTTGGTAAACCTGACGGGCAAA | |
| | oSK54 | CAAAAAAGCGGTTAGCTCCTTCGGT | pUC57-BsaI-free-ATC3 1092 |
| | oSK55 | AGAGACCGTATATCTAGATGCATTCGC | |
| | oSK56 | TTTGCCCGTCAGGTTTACCAAG | |

| | | | |
|---|---|---|---|
| free-ATC3 | oSK27 | CCGAAGGAGCTAACCGCTTTTTGC | pUC57-BsaI-free-ATC3 3340 |
| pSK33 – corancA2 in pUC57-BsaI-free-ATC2 | oSK59 | CACGACTATCGATATTTGTCATCC | pUC57-BsaI-free-ATC2 5901 |
| | oSK60 | ACTGATTGTTGAGGATTGTTCAC | |
| pSK34 – corancaltA3 in pUC57-BsaI-free-ancaltATC3 | oSK54 | CAAAAAAGCGGTTAGCTCCTTCGGT | pUC57-BsaI-free-ancaltATC3 1092 |
| | oSK61 | GAGGAGAGACCATCTAGATGCATT | |
| | oSK62 | AACGGTAAATTGGATCGCCGGGCGTTGCCAGCGCCTGATCAGAATGCCTTT | pUC57-BsaI-free-ancaltATC3 3325 |
| | oSK27 | CCGAAGGAGCTAACCGCTTTTTGC | |
| | oSK63 | AGATGGTCTCTCCTCAACAACCGGTCACGGCCATCGATATTTGTCATCCGCTGAGCGGACATTATTGCTG | corancaltA3 1562 |
| | oSK64 | ATCCAATTTACCGTTCGGCGTC | |
| pSK35 – *gxpS* of *P. bodei* in pCOLA | oSK65 | TTTTTTTGGGCTAACAGGAGGAATTCCATGAAAGATAGCATAGCTAAAAAGGAAATTATCT | *P. bodei* gDNA 6071 |
| | oSK66 | GGTATACCAATGACAACATCATCCTGCC | |
| | oSK67 | GGCAGGATGATGTTGTCATTGGTATACC | *P. bodei* gDNA 9680 |
| | oSK68 | CTCAGCGGTGGCAGCAGCCTAGGTTAATTACAGCGCCTCCGCCTCA | |
| | pCOLA-FW | TAATTAACCTAGGCTGCTGC | pMY-e1-0000 3020 |
| | pCOLA-RV | CATGGAATTCCTCCTGTTAGC | |
| pCL05 - *gxpS* of *X. khoisanae* containing a knocked out T5 domain in pCOLA | oCL28 | TTGGGTGGACACGCACTGCTTGCTATGCGGATGATAAACCTTG | pSK12 1904 |
| | oCL04 | CGATAATGTCGGGCAATCAGGTGC | |
| | oCL03 | GCACCTGATTGCCCGACATTATCG | pSK12 8441 |
| | oCL31 | ACCCTGTCCTCCTCTTCGAGCAAGT | |
| | oCL30 | TTGCTCGAAGAGGAGGACAGGGTTGTTGGTTATC | pSK12 8265 |
| | oCL29 | TGCGTGTCCACCCAACGCAAAGAAATTGTCATACCGGCC | |

| | | | |
|---|---|---|---|
| pSK1 – *gxpS* of *X. szentirmaii* in pCOLA | oSK73 | TTTTTTTGGGCTAACAGGAGGAATTCCATGAAAGATAGCAAGGTTGCTAAAAAGGG | *X. szentirmaii* gDNA 8905 |
| | oSK74 | CGCTGTTCATTACCTCCAATGTGG | |
| | oSK75 | CCACATTGGAGGTAATGAACAGCG | *X. szentirmaii* gDNA 6878 |
| | oSK76 | CTCAGCGGTGGCAGCAGCCTAGGTTAATTACAGCACTTCCTCCTCACAATTCA | |
| | pCOLA-FW | TAATTAACCTAGGCTGCTGC | pMY-e1-0000 3020 |
| | pCOLA-RV | CATGGAATTCCTCCTGTTAGC | |
| pSK44 - *gxpS* of *X. khoisanae* containing a knocked out TE-domain in pCOLA | oSK95 | CGATACCTGTGCGGCCCTTC | ΔTE-domain of *X. khoisanae* 814 |
| | oSK96 | AACGCCATACTTAAGGTATTAGCCA | |
| | oSK99 | CACCTGATTGCCCGACATTATCG | pSK12 2132 |
| | oSK100 | TTGCCGGTAATACTGTCTTTCAT | |
| | oSK101 | ATGAAAGACAGTATTACCGGCAAGG | pSK12 6332 |
| | oCL31 | ACCCTGTCCTCCTCTTCGAGCAA | |
| | oSK102 | TTGCTCGAAGAGGAGGACAGGGTTGTTGGTTATC | pSK12 8424 |
| | oSK103 | GAAGGGCCGCACAGGTATCG | |
| | oSK97 | TGGCTAATACCTTAAGTATGGCGTT | pSK12 961 |
| | oSK98 | CGATAATGTCGGGCAATCAGGTG | |
| pSK45 - *gxpS* of *X. khoisanae* containing a knocked out T5 and TE-domain in pCOLA | oSK95 | CGATACCTGTGCGGCCCTTC | ΔTE-domain of *X. khoisanae* 814 |
| | oSK96 | AACGCCATACTTAAGGTATTAGCCA | |
| | oSK99 | CACCTGATTGCCCGACATTATCG | pCL05 2132 |
| | oSK100 | TTGCCGGTAATACTGTCTTTCAT | |
| | oSK101 | ATGAAAGACAGTATTACCGGCAAGG | pCL05 6332 |
| | oCL31 | ACCCTGTCCTCCTCTTCGAGCAA | |
| | oSK102 | TTGCTCGAAGAGGAGGACAGGGTTGTTGGTTATC | pCL05 8424 |
| | oSK103 | GAAGGGCCGCACAGGTATCG | |
| | oSK97 | TGGCTAATACCTTAAGTATGGCGTT | pCL05 961 |
| | oSK98 | CGATAATGTCGGGCAATCAGGTG | |
| pSK47 - *gxpS* of *X. khoisanae* | oCL03_2 | GCACCTGATTGCCCGACATTATC | pSK12 8441 |
| | oCL31_2 | ACCCTGTCCTCCTCTTCGAGC | |
| | oCL30_2 | TTGCTCGAAGAGGAGGACAGGG | pSK12 5069 |
| | oSK104 | CCCAACGCGAAGAAGTTATCATGT | |
| | oSK105 | TAGTCGACATGATAACTTCTTCGCGTTGGGTGGTCATTCGTTGTTGGCGA | pSK12 |

| | | | |
|---|---|---|---|
| containing its C2 domain also in its 4th module in pCOLA | oSK106 | TTCAGAATAGGCTGTTTCAGTCGCATTCCGGGTTTCCAGCAACAACCGGC | 1579 |
| | oSK107 | CGGAATGCGACTGAAACAGCCTATTC | pSK12 3581 |
| | oCL04 | CGATAATGTCGGGCAATCAGGTGC | |

**Table S9.** Plasmids and corresponding NRPS used in this work.

| Plasmids | Genotype | Reference |
|---|---|---|
| pCK_0433 pCOLA | ori ColA, *kan*$^R$, *araC*- $P_{BAD}$,, tacI, I-SceI, I-CeuI | [10] |
| pSK4 | ori ColA, *kan*$^R$, *araC*- $P_{BAD}$, *gxpS P. luminescens namnaonensis* | This work |
| pSK3 | ori ColA, *kan*$^R$, *araC*- $P_{BAD}$, *gxpS X. ehlersii* | This work |
| pSK5 | ori ColA, *kan*$^R$, *araC*- $P_{BAD}$, gxpS *X. innexi* | This work |
| pUC57-BsaI-free-ATC1 | *E. coli* cloning vector, ori ColE1, *amp*$^R$, suitable *bsaI*-sites for Golden Gate assembly, ancATC1 of *ancgxpS* | Genscript Biotech Corp |
| pUC57-BsaI-free-ATC2 | *E. coli* cloning vector, ori ColE1, *amp*$^R$, suitable *bsaI*-sites for Golden Gate assembly, ancATC2 of *ancgxpS* | Genscript Biotech Corp |
| pUC57-BsaI-free-ATC3 | *E. coli* cloning vector, ori ColE1, *amp*$^R$, suitable *bsaI*-sites for Golden Gate assembly, ancATC3 of *ancgxpS* | Genscript Biotech Corp |
| pUC57-BsaI-free-ATC4 | *E. coli* cloning vector, ori ColE1, *amp*$^R$, suitable *bsaI*-sites for Golden Gate assembly, ancATC4 of *ancgxpS* | Genscript Biotech Corp |
| pUC57-BsaI-free-ATTE | *E. coli* cloning vector, ori ColE1, *amp*$^R$, suitable *bsaI*-sites for Golden Gate assembly, ancATTE of *ancgxpS* | Genscript Biotech Corp |
| pMY-e1-0000_GG | pMY-e1-0000 plasmid with suitable *bsaI* sites for Golden Gate assembly, *sacB* cassette as second selection marker | This work |
| pSK12 | ori ColA, *kan*$^R$, *araC*- $P_{BAD}$, *gxpS X. khoisanae* | This work |
| pSK6 | ori ColA, *kan*$^R$, *araC*- $P_{BAD}$, *gxpS P. cinerea* | This work |
| pSK8 | ori ColA, *kan*$^R$, *araC*- $P_{BAD}$, *gxpS P. thracensis* | This work |
| pSK7 | ori ColA, *kan*$^R$, *araC*- $P_{BAD}$, *gxpS Xenorhabdus* sp. PB 62.4 | This work |

| | | |
|---|---|---|
| pUC57-BsaI-free-ancaltATC1 | *E. coli* cloning vector, ori ColE1, *amp*$^R$, suitable *bsaI*-sites for Golden Gate assembly, ancaltATC1 of *ancaltgxpS* | Genscript Biotech Corp |
| pUC57-BsaI-free-ancaltATC1 | *E. coli* cloning vector, ori ColE1, *amp*$^R$, suitable *bsaI*-sites for Golden Gate assembly, ancaltATC2 of *ancaltgxpS* | Genscript Biotech Corp |
| pUC57-BsaI-free-ancaltATC1 | *E. coli* cloning vector, ori ColE1, *amp*$^R$, suitable *bsaI*-sites for Golden Gate assembly, ancaltATC3 of *ancaltgxpS* | Genscript Biotech Corp |
| pUC57-BsaI-free-ancaltATC1 | *E. coli* cloning vector, ori ColE1, *amp*$^R$, suitable *bsaI*-sites for Golden Gate assembly, ancaltATC4 of *ancaltgxpS* | Genscript Biotech Corp |
| pUC57-BsaI-free-ancaltATC1 | *E. coli* cloning vector, ori ColE1, *amp*$^R$, suitable *bsaI*-sites for Golden Gate assembly, ancaltATTE of *ancaltgxpS* | Genscript Biotech Corp |
| pSK35 | ori ColA, *kan*$^R$, *araC- $P_{BAD}$, gxpS P. bodei* | This work |
| pSK1 | ori ColA, *kan*$^R$, *araC- $P_{BAD}$, gxpS X. szentirmaii* | This work |
| pUC57-BsaI-free-Vera, ATC1 | *E. coli* cloning vector, ori ColE1, *amp*$^R$, suitable *bsaI*-sites for Golden Gate assembly, ATC1 of *Xenorhabdus* sp. Vera | Genscript Biotech Corp |
| pUC57-BsaI-free-Vera, ATC2 | *E. coli* cloning vector, ori ColE1, *amp*$^R$, suitable *bsaI*-sites for Golden Gate assembly, ATC2 of *Xenorhabdus* sp. Vera | Genscript Biotech Corp |
| pUC57-BsaI-free-Vera, ATC3 | *E. coli* cloning vector, ori ColE1, *amp*$^R$, suitable *bsaI*-sites for Golden Gate assembly, ATC3 of *Xenorhabdus* sp. Vera | Genscript Biotech Corp |
| pUC57-BsaI-free-Vera, ATTE | *E. coli* cloning vector, ori ColE1, *amp*$^R$, suitable *bsaI*-sites for Golden Gate assembly, ATTE of *Xenorhabdus* sp. Vera | Genscript Biotech Corp |
| pSK42 | ori ColA, *kan*$^R$, *araC- $P_{BAD}$ xtpS Xenorhabdus* sp. Vera | This work |
| pSK32 | *E. coli* cloning vector, ori ColE1, *amp*$^R$, suitable *bsaI*-sites for Golden Gate assembly, corancATC3 | This work |
| pSK33 | *E. coli* cloning vector, ori ColE1, *amp*$^R$, suitable *bsaI*-sites for Golden Gate assembly, corancATC2 | This work |
| pSK37 | ori ColA, *kan*$^R$, *araC- $P_{BAD}$*, *ancgxpS* | This work |
| pSK34 | *E. coli* cloning vector, ori ColE1, *amp*$^R$, suitable *bsaI*-sites for Golden Gate assembly, corancaltATC3 | This work |
| pSK43 | ori ColA, *kan*$^R$, *araC- $P_{BAD}$*, ancalt*gxpS* | This work |

| | | |
|---|---|---|
| pCOLA *ara gxpS tacI* JW | ori ColA, *kan*$^R$, *araC*- $P_{BAD}$, *tacI gxpS P. laumondii* TT01 | 11 |
| pCOLA *ara xtpS tacI* JW | ori ColA, *kan*$^R$, *araC*- $P_{BAD}$, *tacI gxpS X. nematophila* ATCC 19061 | 11 |
| pACYC *araP gxpS X. doucetiae* | *cm*$^R$, ori p15A, *araC*- $P_{BAD}$, *X. doucetiae* FRM16 | This work |
| pACYC *araP gxpS X. miraniensis* | *cm*$^R$, ori p15A, *araC*- $P_{BAD}$, *X. miraniensis* DSM 17902 | This work |
| pCL05 | ori ColA, *kan*$^R$, *araC*- $P_{BAD}$, *gxpS X. khoisanae* ΔT5 domain | This work |
| pSK44 | ori ColA, *kan*$^R$, *araC*- $P_{BAD}$, *gxpS X. khoisanae* ΔTE-domain | This work |
| pSK45 | ori ColA, *kan*$^R$, *araC*- $P_{BAD}$, *gxpS X. khoisanae* ΔT5 domain ΔTE-domain | This work |
| pSK47 | ori ColA, *kan*$^R$, *araC*- $P_{BAD}$, *gxpS X. khoisanae*_$A_1T_1C/E_1A_2T_2C_2A_3T_3C/E_3$ $A_4T_2C_2$ $A_5T_5$TTE | This work |
| pCK0403 | *cm*$^R$, ori p15A, *araC*- $P_{BAD}$, *mtaA* | 2 |
| pCK0413 | *kan*$^R$, ColA p15A, *araC*- $P_{BAD}$, *mtaA* | 12 |

**Table S10.** Purification steps of GXP using a SPE Strata.

| Step | Volume | Solvent | Comments |
|---|---|---|---|
| Column conditioning | 2x2.5 mL | MeOH | |
| Equilibration | 3x1.5 mL | DCM | |
| Sample loading | - | Sample/DCM suspension | 50 µmol peptide |
| Wash | Min. 5 mL | DCM | |
| Dry | - | - | To complete dryness |
| Wash | 3x2 mL | dd$H_2O$ + 0.1% FA (4°C) | |
| Wash | 3x2 mL | 1:1 MeOH/ dd$H_2O$ + 0.1% FA (4°C) | |
| Wash | 3x2 mL | MeOH + 0.1% FA (4°C) | Sample should be colorless |

**Table S11.** GameXPeptides that were synthesized by solid-phase-peptide synthesis. Compounds **1**, **36** and **7** were synthesized by WUXI AppTec.

| No. | No. AAs | Sequence | Linearization position | Peptide amount [mg] |
|---|---|---|---|---|
| **1** | 5 | cyclo (D-Val/L-Leu/D-Phe/D-Leu/L-Leu) | 5-1 | 111 |
| **36** | 5 | cyclo (D-Val/L-Leu/D-*p*-aminophenylalanine/D-Leu/L-Leu) | 5-1 | 83 |
| **43** | 5 | cyclo (D-Val/L-Leu/D-monomethylated-*p*-aminophenylalanine/D-Leu/L-Leu) | 5-1 | 11 |
| **6** | 5 | cyclo (D-Val/L-Leu/D-Leu/D-Leu/L-Leu) | 5-1 | 7.7 |
| **11** | 4 | cyclo (D-Val/L-Val/D-Leu/L-Leu) | 4-1 | 29.5 |
| **4** | 4 | cyclo (D-Val/L-Val/D-Leu/L-Val/L-Leu) | 5-1 | 53 |
| **10** | 4 | cyclo (D-Val/L-Val/D-Leu/L-Val) | 4-1 | 20.9 |
| **38** | 6 | cyclo (D-Val/D-Leu/L-Ile/D-Leu/L-Leu/L-*p*-aminophenylalanine) | 6-1 | 33.2 |
| **7** | 6 | cyclo (D-Val/D-Leu/L-Ile/D-Leu/L-Leu/L-Phe) | 6-1 | 60.4 |
| **2** | 4 | cyclo (D-Val/L-Leu/D-Val/L-Val) | 4-1 | 20.5 |
| **5** | 4 | cyclo (D-Val/L-Val/D-Leu/L-Leu/L-Leu) | 5-1 | 56.2 |
| **3** | 4 | cyclo (D-Val/L-Leu/D-Leu/L-Val) | 2-3 | 30.9 |
| **13** | 6 | cyclo (D-*allo*-Ile/D-Leu/L-Ile/D-Leu/L-Leu/L-Phe) | 6-1 | 26.5 |
| **8** | 4 | cyclo (D-*allo*-Ile/L-Leu/D-Val/L-Val) | 2-3 | 33.3 |
| **9** | 4 | cyclo (D-*allo*-Ile/L-Leu/D-Leu/L-Val) | 2-3 | 32.5 |

| | | | | |
|---|---|---|---|---|
| **14** | 5 | cyclo (D-*allo*-Ile/L-Leu/D-Phe/D-Leu/L-Leu) | 5-1 | 7.7 |
| **12** | 5 | cyclo (D-*allo*-Ile/L-Leu/D-Leu/D-Leu/L-Leu) | 5-1 | 50.6 |
| **16** | 5 | cyclo (D-Val/L-Leu/D-Met/D-Leu/L-Leu) | 5-1 | 25.1 |

**Table S12.** Five different models that were used to describe the best fit to our bioactivity data. AIC determined that model 5 fits best our data.

| **Model** | **Description** | **Number of Parameters** | $\chi^2$ | **AIC** | **Ref. Supplementary Figs.** |
|---|---|---|---|---|---|
| 1 | 1 sigmoid for all | 2 | 2266 | 1707 | S52 |
| 2 | 1 sigmoid for each peptide | 16 | 2067 | 1723 | S53 |
| 3 | 1 sigmoid for each species | 8 | 805 | 1498 | S54 |
| 4 | 1 sigmoid for each peptide + correction factor for each species | 20 | 634 | 1483 | S55 |
| 5 | 1 sigmoid for each peptide and species combination | 64 | 391 | 1290 | S56 |

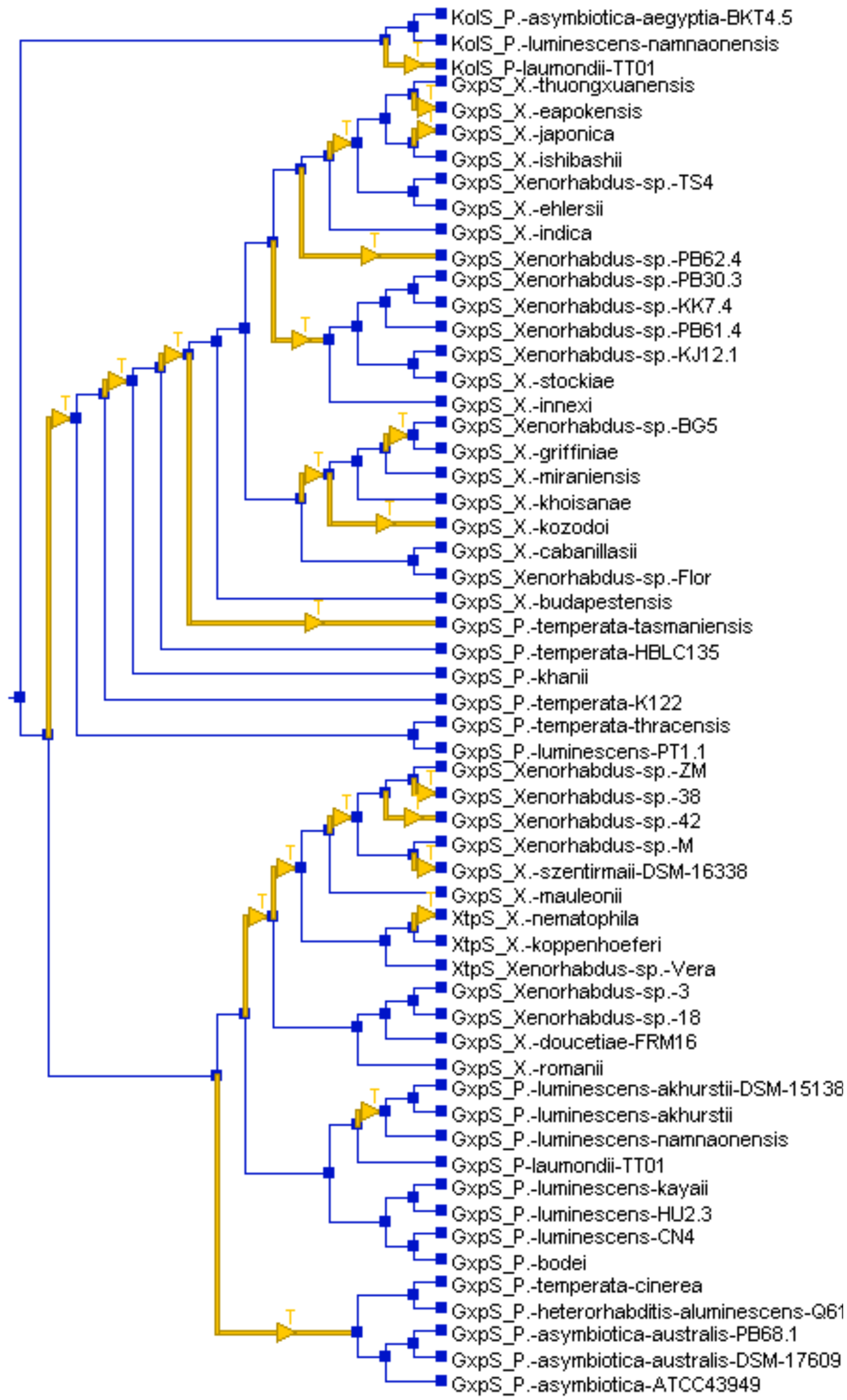


**Figure S1. Reconciliation analysis of the TTE-domain tree of GxpS from *Xenorhabdus* and *Photorhabdus*.** TTE-domain phylogeny with inferred horizontal transfers mapped onto branches as yellow arrows labelled T. Losses are not shown.

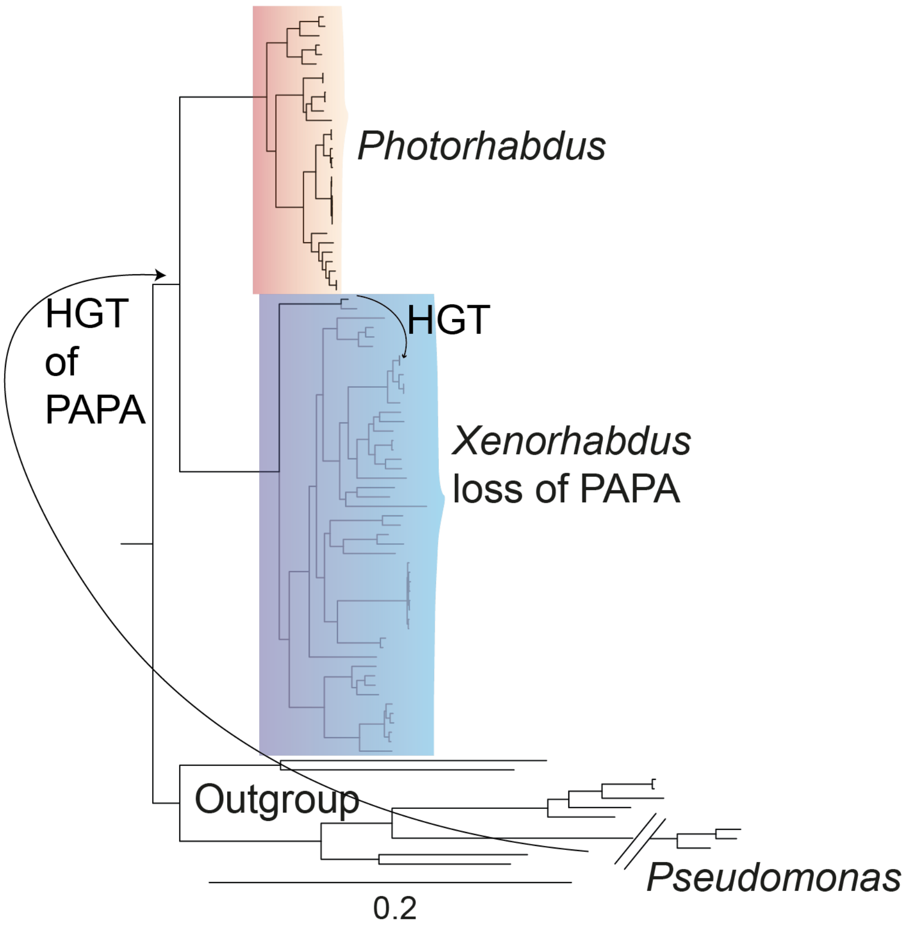


**Figure S2. A species tree from *Xenorhabdus* and *Photorhabdus* includes also three *Pseudomonas* species.** As outgroup other gammaproteobacteria were used. Branches within the XP clade that are black still contain the PAPA operon. Branches in gray lost it. Indicated is the potential HGT from *Pseudomonas* towards the LCA of XP. Also indicated is the HGT of the PAPA operon within the *Xenorhabdus* clade.

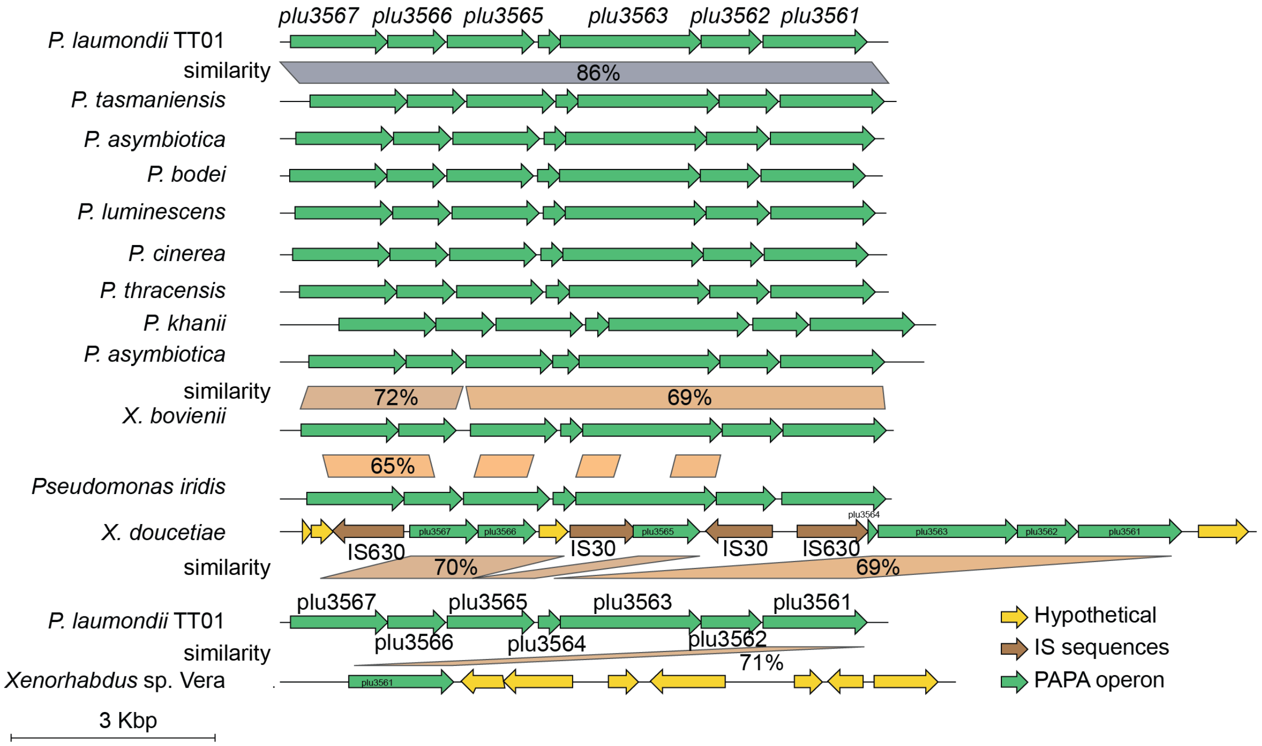


**Figure S3. Representative examples of PAPA operons from various species of *Photorhabdus*.** In the *Xenorhabdus* genus only *X. bovienii* affine MC10 has still a functional PAPA operon (other *X. bovienii* strains do not have this operon anymore). Non-functional PAPA operons from *X. doucetiae* and *Xenorhabdus* sp. Vera are also shown.

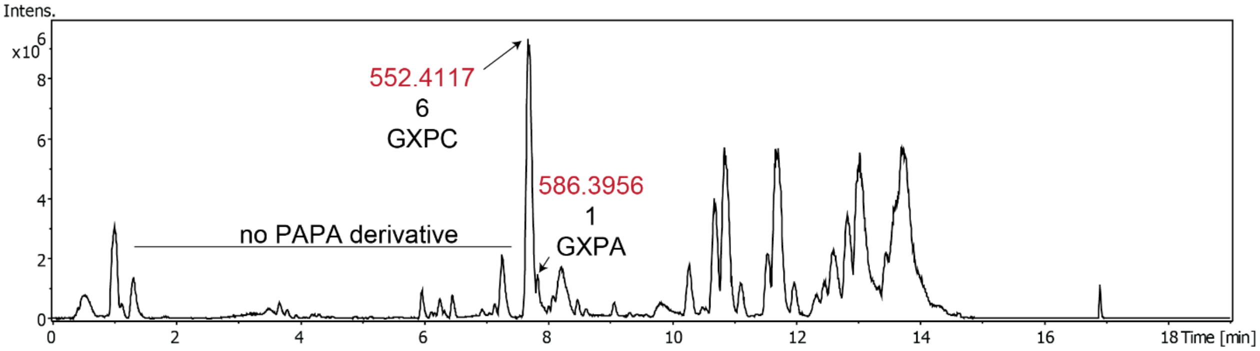


**Figure S4. *X. doucetiae* cultured in *G. mellonella* for 72 h.** Base peak chromatogram of natural products extracted with methanol from the insect hemolymph. Peaks corresponding to GXP are labelled. No GameXPeptides incorporating PAPA were detected.

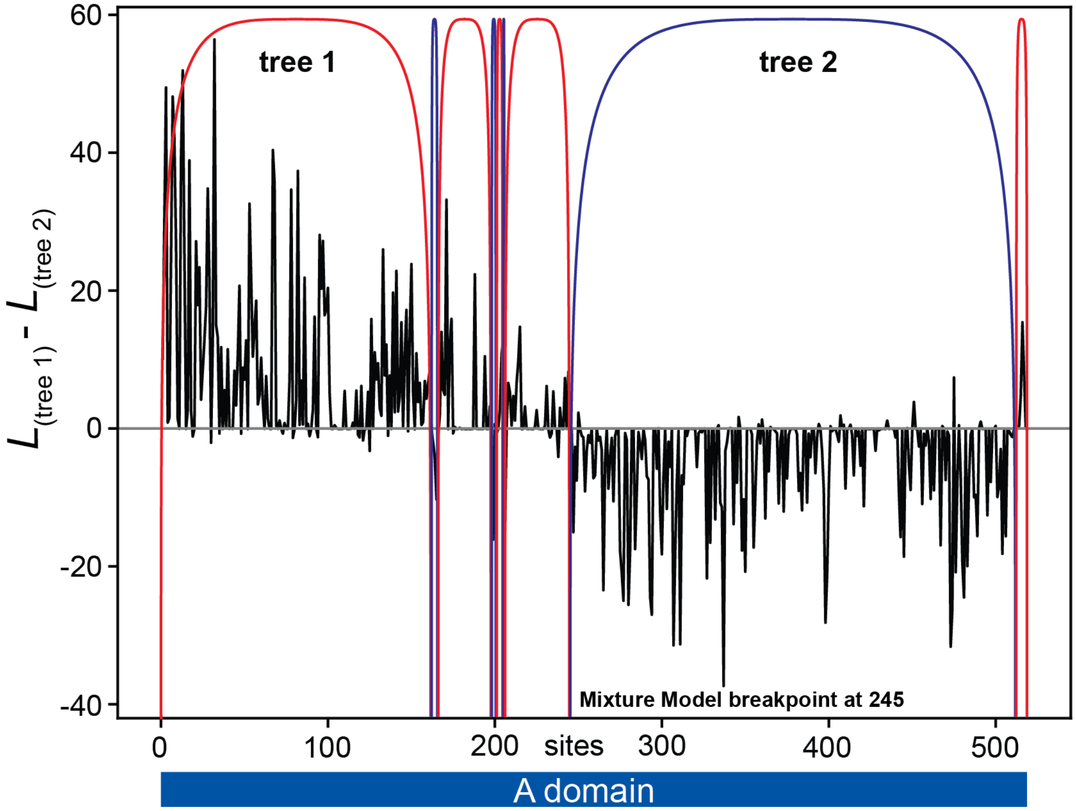


**Figure S5**. **HMM analysis of A3 and A5 domains of GxpS to find recombinations within A3 domains.** Plot shows the difference in the site-wise log-likelihoods of two phylogenetic trees that together best describe the alignment using a phylogenetic hidden Markov model. Sites with positive values are better described by tree 1, sites with negative numbers are better described by tree 2. Alignment was then partitioned using the Viterbi and the back-to-back algorithm. Arches indicated partitions into these two histories where red arches correspond to tree 1 and blue arches to tree 2. Partitioning using the MM analysis pointed to site 245.

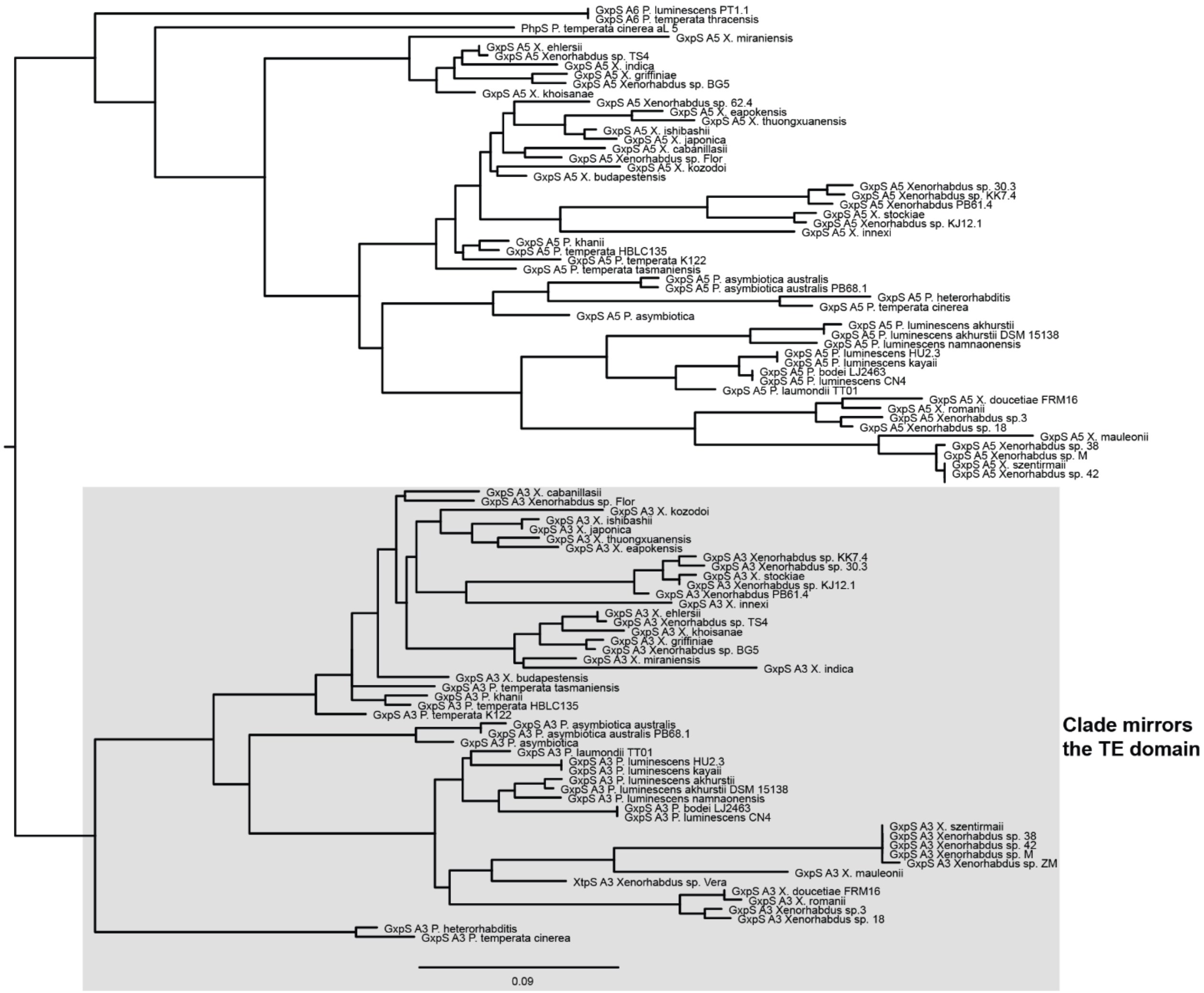


**Figure S6. Tree 1 of the HMM analysis of the A3 and A5 alignment**. All A3 domains form a monophyletic group. A5 domains form an outgroup to the A3 clade. Tree 1 is mostly congruent with the TTE-domain.

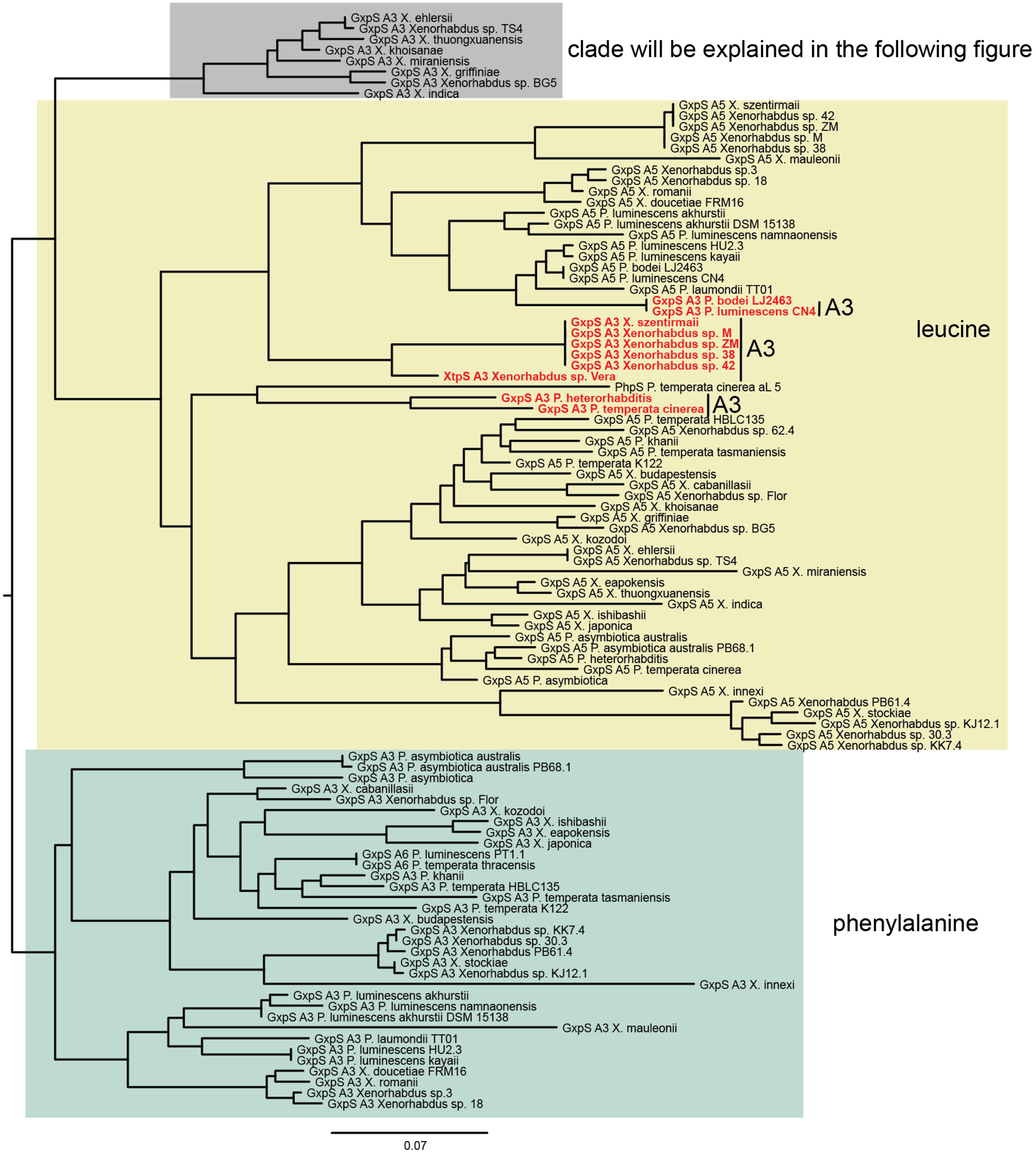


**Figure S7. Tree 2 of the HMM analysis of the A3 and A5 alignment**. This tree separates mostly leucine (in yellow) and phenylalanine (in green) A-domains. Some A3 domains are found in the leucine clade and are colored in red. A small clade (colored in gray) of A3-domains sits neither in the phenylalanine nor in the leucine A-domain and is shown in gray.

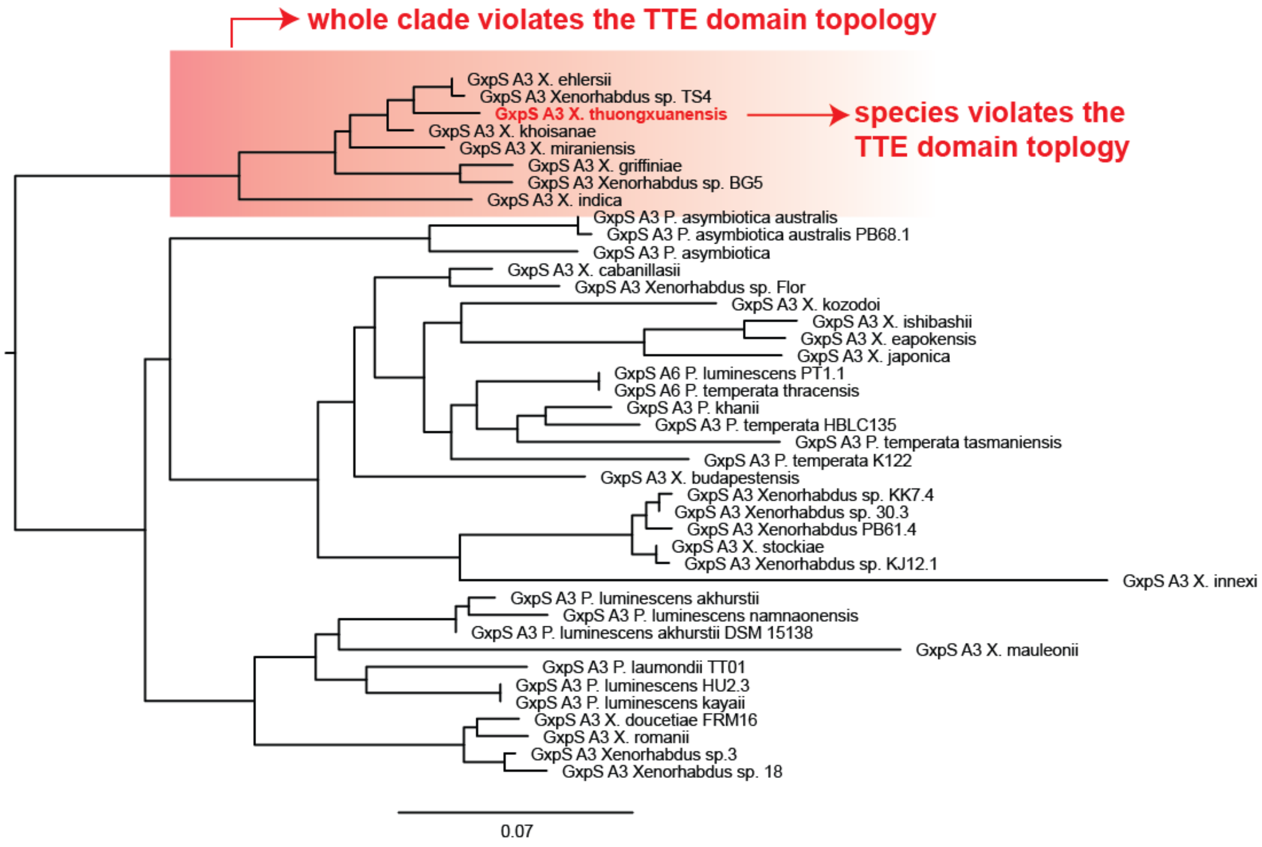


**Figure S8. Tree 2 of the HMM analysis of the A3 and A5 alignment showing more hidden recombination.** A clade of tree 2 of the HMM analysis from the A3 and A5 alignment sits in an anomalous position. 8 species such as *Xenorhabdus ehlersii* sit outside of the A3 clade. The A3 domain of *Xenorhabdus thuongxuanensis* branches inside this clade and sits far from its usual position in the TTE-domain phylogeny and therefore indicates a further recombination.

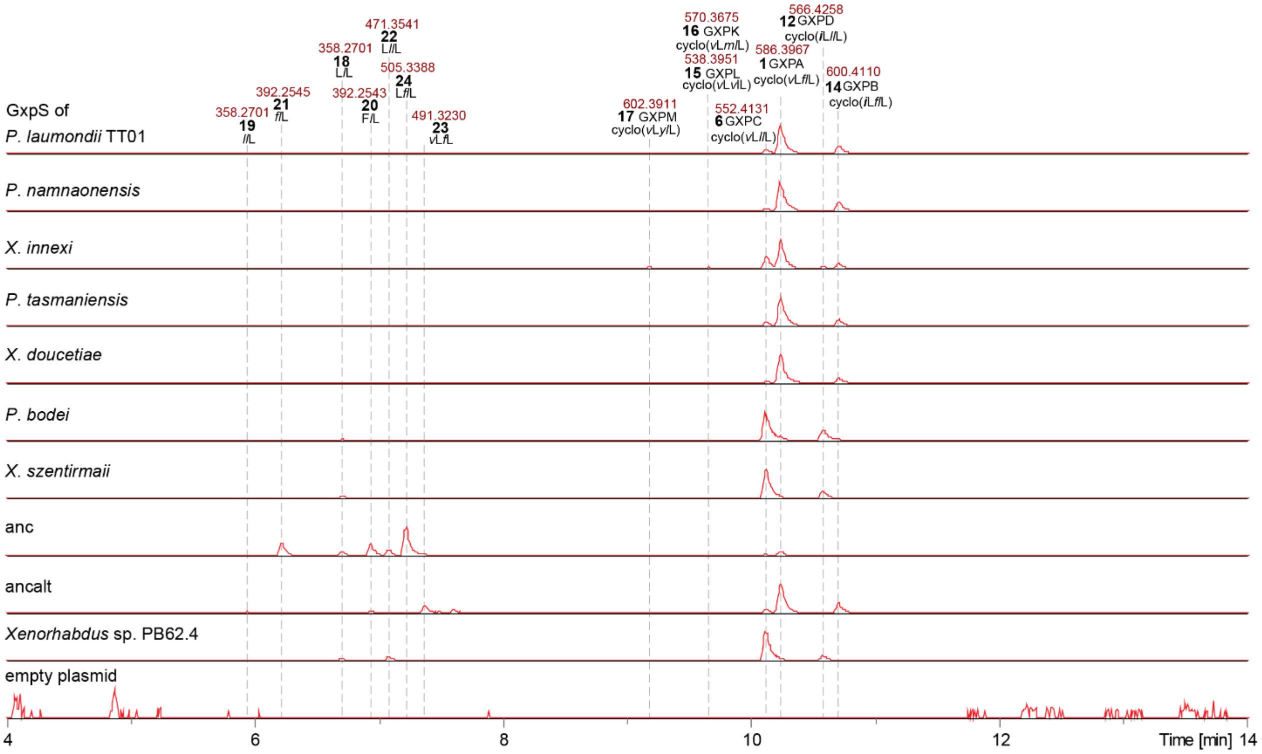


**Figure S9. Extracted ion chromatogram of all main peaks from AncGxpS, all non-recombined GxpS and GxpS that lost phenylalanine specificity in their A3 domain.** Shown are the ionized high-resolution mass and the corresponding peptide sequence. Small, italic letters indicate D-amino acids; ***i*** = D-*allo*-Ile.

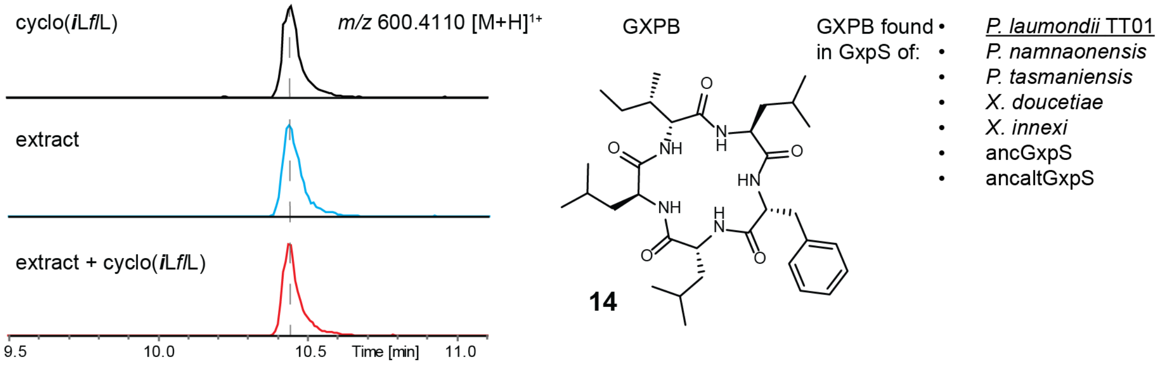


**Figure S10**. **Comparison of retention times of synthetic (black), natural (blue) and co-injected (red) GameXPeptide B (14)**. Small, italic letters indicate D-amino acids; ***i*** = D-*allo*-Ile. The structure of GXPB is shown and is cyclo(***i***L*f*/L). GXPB was measured in a culture extract of *E. coli* DH10B$^{penta}$ carrying *gxpS* of *P. laumondii* TT01 and pCK0403.

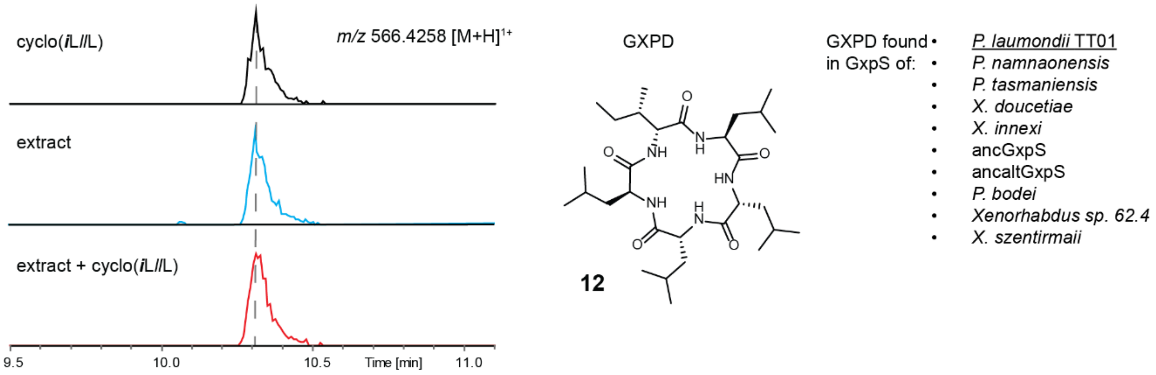


**Figure S11. Comparison of retention times of synthetic (black), natural (blue) and co-injected (red) GameXPeptide D (12).** Small, italic letters indicate D-amino acids; ***i*** = D-*allo*-Ile. The structure of GXPD is shown and is cyclo(***i***L*l*L). GXPD was measured in a culture extract of *E. coli* DH10B$^{penta}$ carrying *gxpS* of *P. laumondii* TT01 and pCK0403.

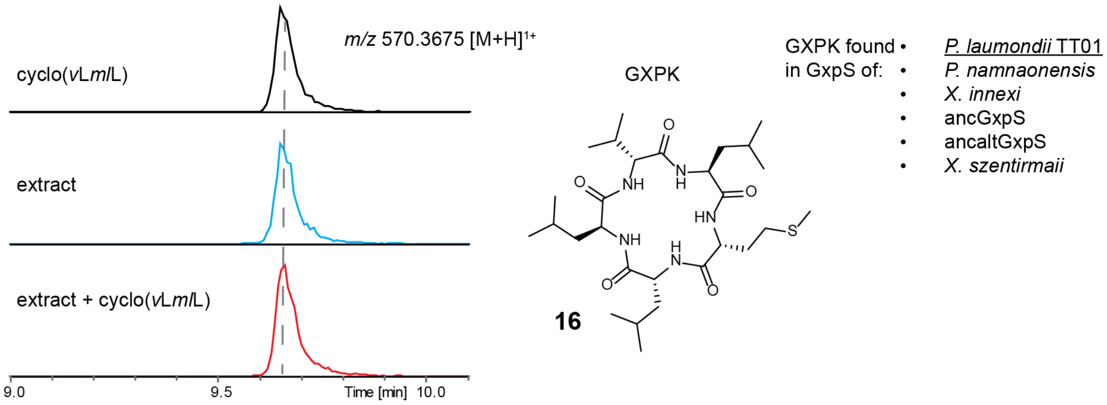


**Figure S12. Comparison of retention times of synthetic (black), natural (blue) and co-injected (red) GameXPeptide K (16).** Co-injection of synthetic GXPK (black), natural GXPK (blue) and co-injected GXPKs (red). Small, italic letters indicate D-amino acids. The structure of GXPK is shown and is cyclo(*v*L*ml*L). GXPK was measured in a culture extract of *E. coli* DH10B$^{penta}$ carrying *gxpS* of *P. laumondii* TT01 and pCK0403.

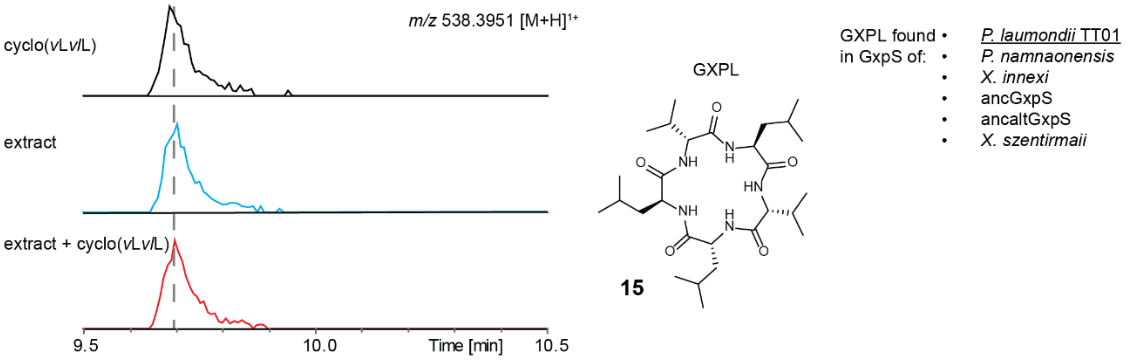


**Figure S13. Comparison of retention times of synthetic (black), natural (blue) and co-injected (red) GameXPeptide L (15).** Small, italic letters indicate D-amino acids. The structure of GXPL is shown and is cyclo(*v*L*vl*L). GXPL was measured in a culture extract of *E. coli* DH10B$^{penta}$ carrying *gxpS* of *P. laumondii* TT01 and pCK0403.

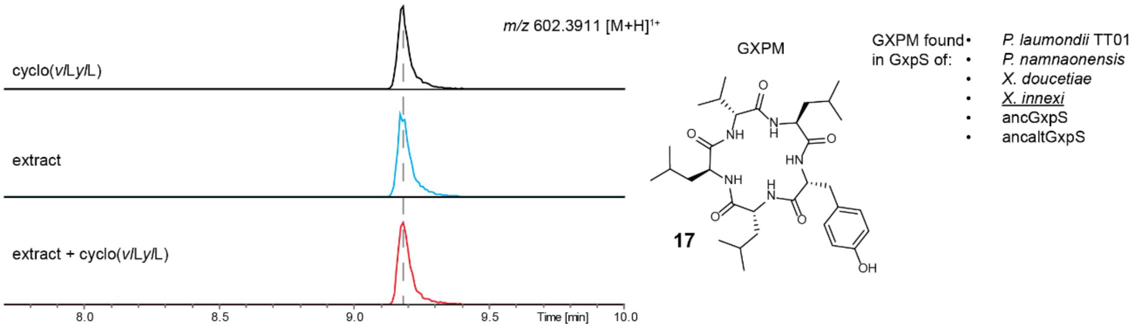


**Figure S14. Comparison of retention times of synthetic (black), natural (blue) and co-injected (red) GameXPeptide D (17).** Small, italic letters indicate D-amino acids. The structure of GXPM is shown and is cyclo(*v*L*yl*L). GXPM was measured in a culture extract of *E. coli* DH10B$^{penta}$ carrying *gxpS* of *X. innexi* and pCK0403.

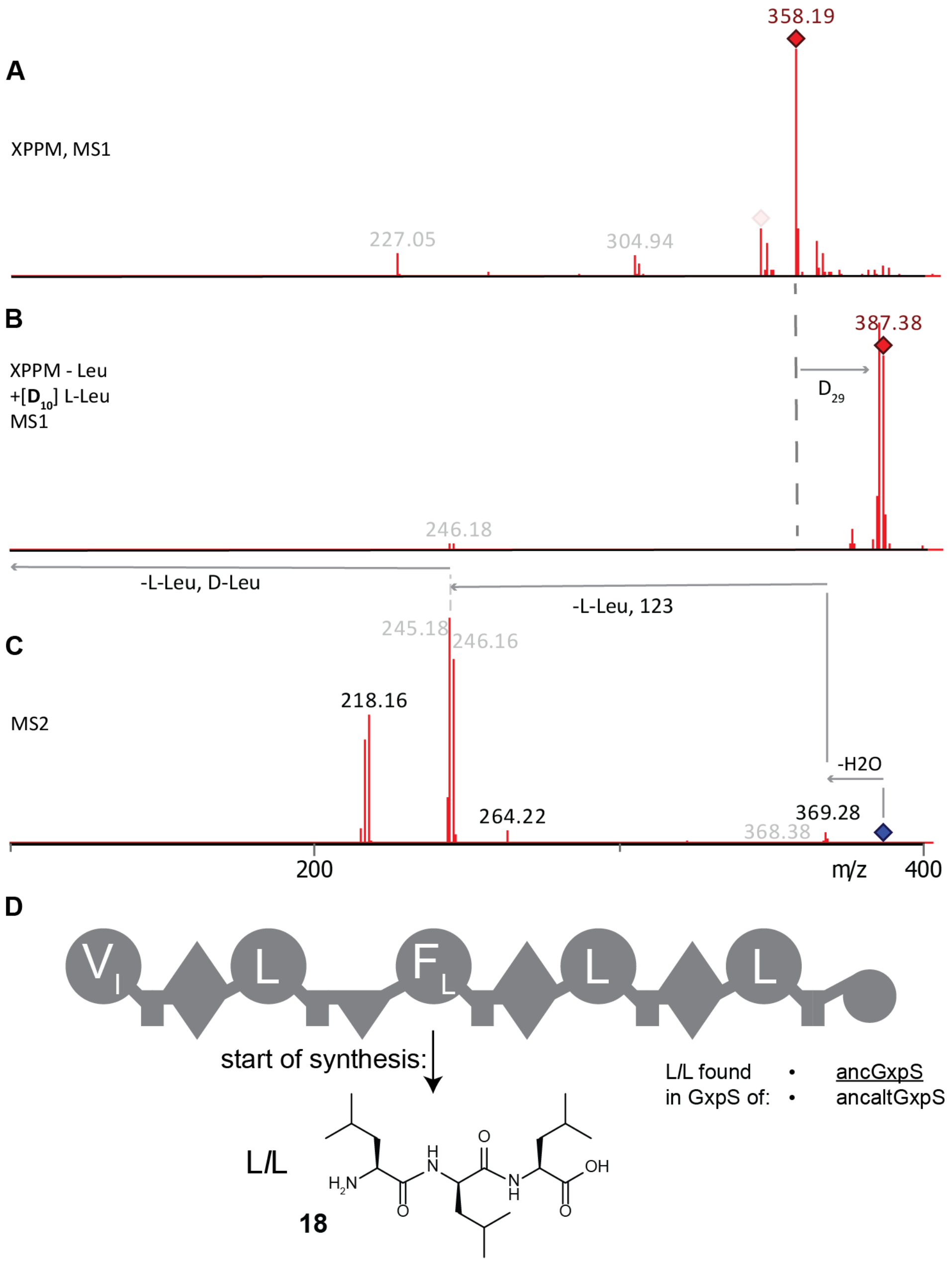


**Figure S15. Structural elucidation of the Tripeptide L*l*L of ancestral GxpS (18).** *ancgxpS* was heterologously expressed in *E. coli* DH10B[penta] together with pCK0403 and the produced natural products were extracted from the culture extract with methanol. (**A**) Mass of Tripeptide L*l*L measured by HPLC-MS. Expression was done in XPPM. (**B**) Mass of Tripeptide L*l*L measured by HPLC-MS. Expression was done in XPPM without leucine supplemented with deuterated leucine. (**C**) MS2 to elucidate the structure of tripeptide L*l*L measured by HPLC-MS. Expression was done in XPPM without leucine supplemented with deuterated leucine. (**D**) Diagram of ancestral GxpS. The start of the biosynthesis of the tripeptide starts at the third A-domain and is indicated by an arrow. Small, italic letters indicate D-amino acids.

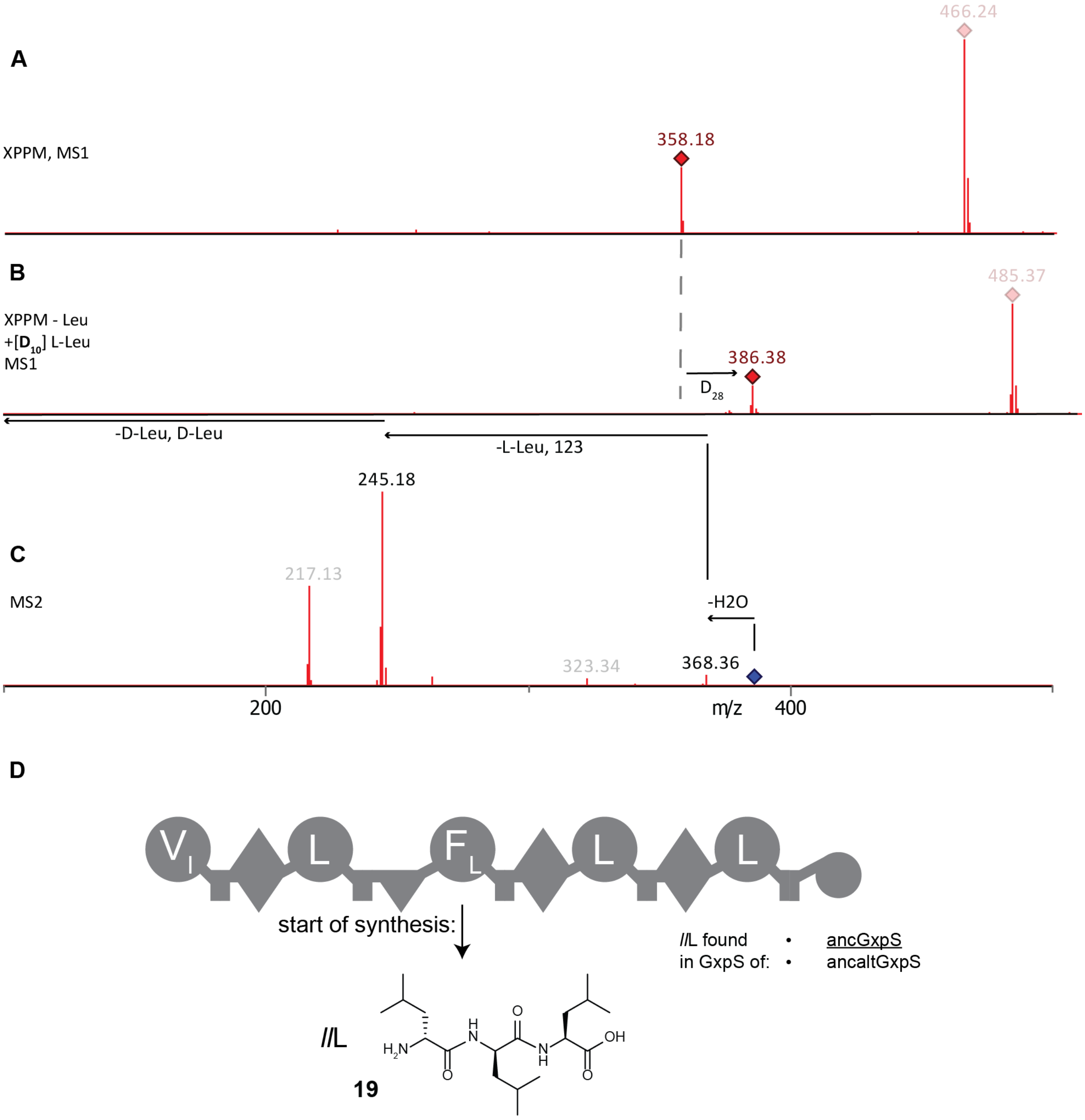


**Figure S16. Structural elucidation of the Tripeptide *ll*L of theancestral GxpS (19).** *ancgxpS* was heterologously expressed in *E. coli* DH10B$^{penta}$ together with pCK0403 and the produced natural products were extracted with methanol. (**A**) Mass of Tripeptide *ll*L measured by HPLC-MS. Expression was done in XPPM. (**B**) Mass of Tripeptide *ll*L measured by HPLC-MS. Expression was done in XPPM without leucine supplemented with deuterated leucine. (**C**) MS2 to elucidate the structure of tripeptide *ll*L. *E. coli* was cultured in XPPM without leucine supplemented with deuterated leucine. (**D**) Diagram of ancestral GxpS. The start of the biosynthesis of the tripeptide starts at the third A-domain and is indicated by an arrow. Small, italic letters indicate D-amino acids.

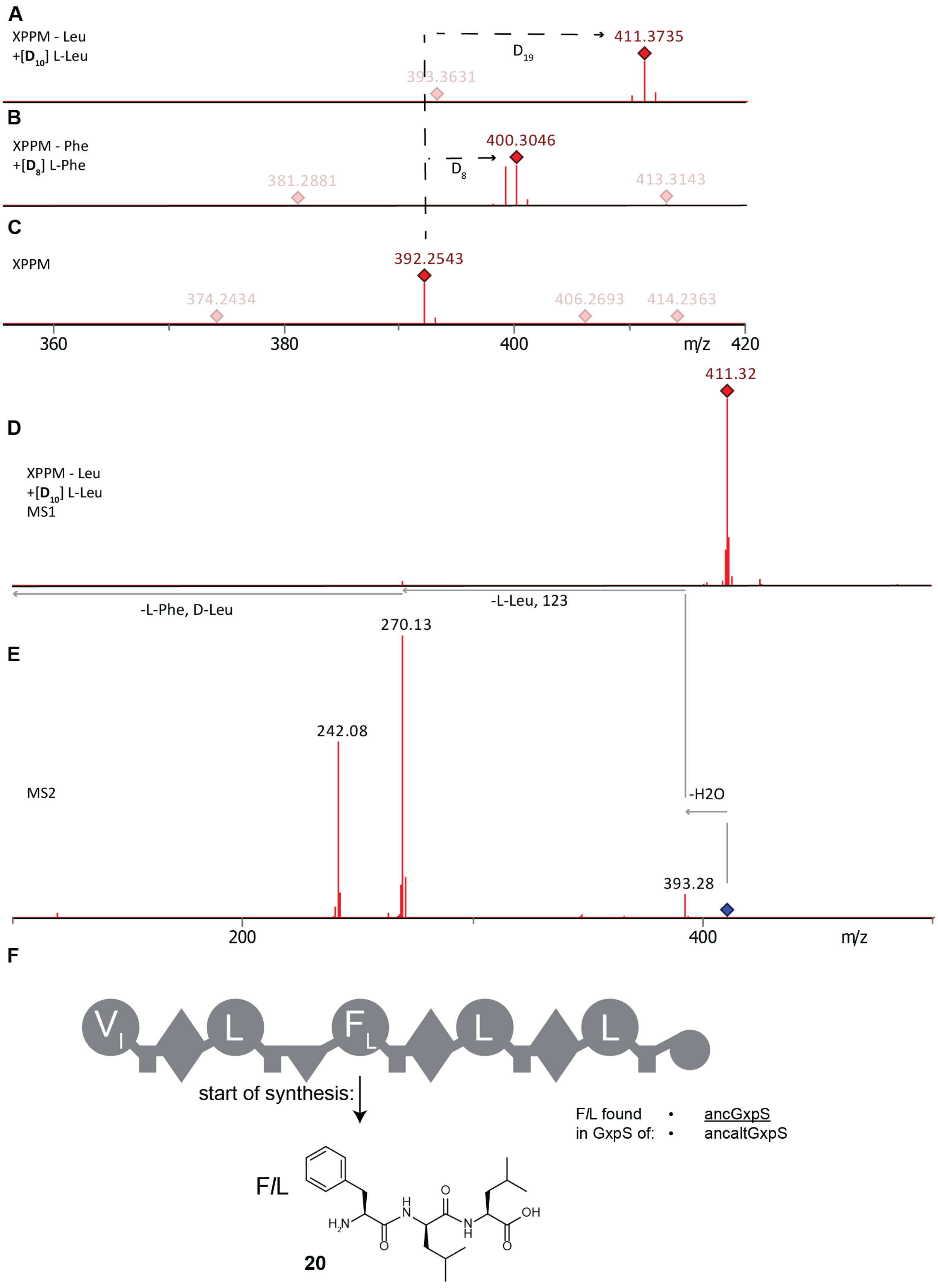
A
XPPM - Leu
+[D10] L-Leu
411.3735
D19
393.3631
B
XPPM - Phe
+[D8] L-Phe
400.3046
D8
381.2881
413.3143
C
XPPM
392.2543
374.2434
406.2693
414.2363
360
380
400
m/z
420
411.32
D
XPPM - Leu
+[D10] L-Leu
MS1
-L-Phe, D-Leu
-L-Leu, 123
270.13
E
242.08
MS2
-H2O
393.28
200
400
m/z
F
V
L
F
L
L
start of synthesis:
F/L found
in GxpS of:
ancGxpS
ancaltGxpS
F/L
H2N
H
N
O
N
H
OH
O
20

**Figure S17. Structural elucidation of the tripeptide F*l*L of the ancestral GxpS (20).** *ancgxpS* was heterologously expressed in *E. coli* DH10B$^{penta}$ together with pCK0403 and the produced natural products were extracted with methanol. (**A**) High-resolution mass of tripeptide F*l*L measured by HPLC-HRMS. Expression was done in XPPM without leucine supplemented with deuterated leucine. (**B**) High-resolution mass of tripeptide F*l*L measured by HPLC-HRMS. Expression was done in XPPM without phenylalanine supplemented with deuterated phenylalanine. (**C**) High-resolution mass of tripeptide F*l*L measured by HPLC-HRMS. Expression was done in XPPM. (**D**) Same as (A) but measured with HPLC-MS. (**E**) MS2 to elucidate the structure of tripeptide F*l*L measured by HPLC-MS. Expression was done in XPPM without leucine supplemented with deuterated leucine. (**F**) Diagram of ancestral GxpS. The start of the biosynthesis of the tripeptide starts at the third A-domain and is indicated by an arrow. Small, italic letters indicate D-amino acids.

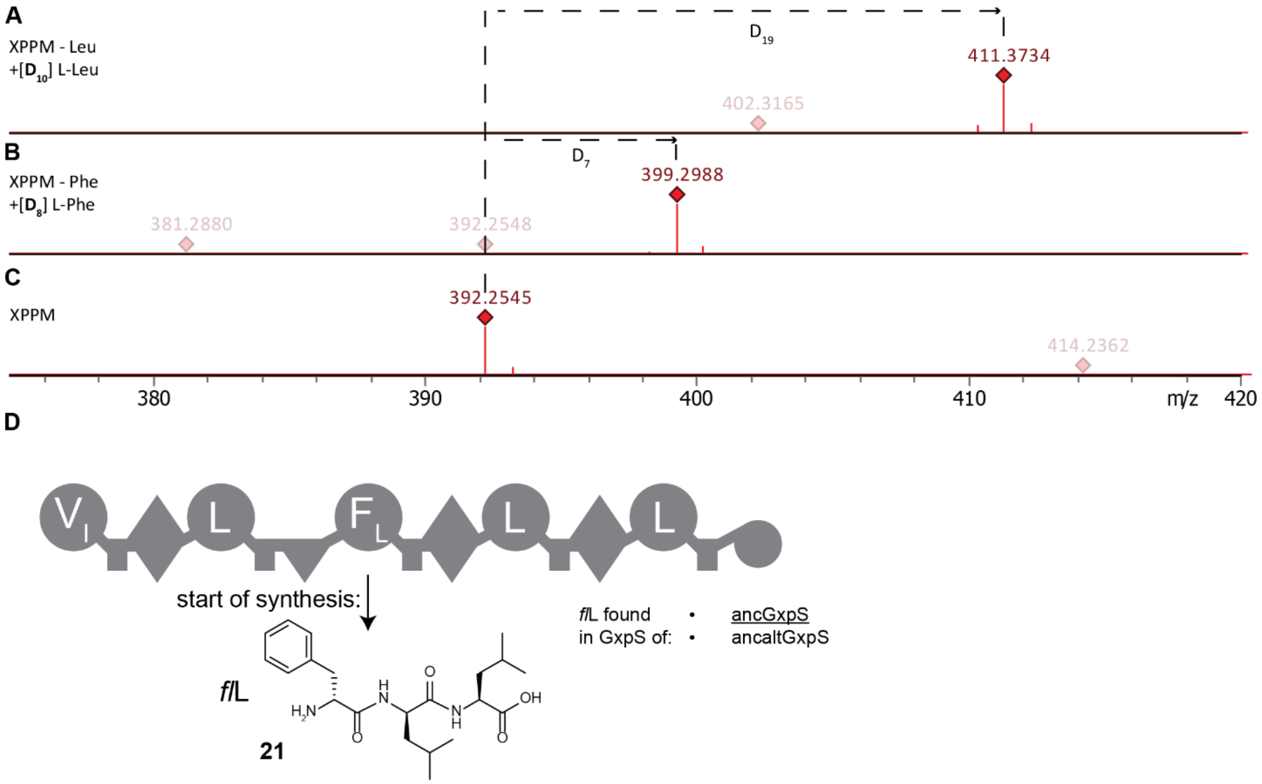


**Figure S18. Structural elucidation of the Tripeptide *fl*L of the ancestral GxpS (21).** *ancgxpS* was heterologously expressed in *E. coli* DH10B$^{penta}$ together with pCK0403 and the produced natural products were extracted with methanol. (**A**) High-resolution mass of tripeptide *fl*L measured by HPLC-HRMS. Expression was done in XPPM without leucine supplemented with deuterated leucine. (**B**) High-resolution mass of tripeptide *fl*L measured by HPLC-HRMS. Expression was done in XPPM without phenylalanine supplemented with deuterated phenylalanine. (**C**) High-resolution mass of Tripeptide *fl*L measured by HPLC-MS. Expression was done in XPPM. (**D**) Diagram of ancestral GxpS. The start of the biosynthesis of the tripeptide starts at the third A-domain and is indicated by an arrow. Small, italic letters indicate D-amino acids.

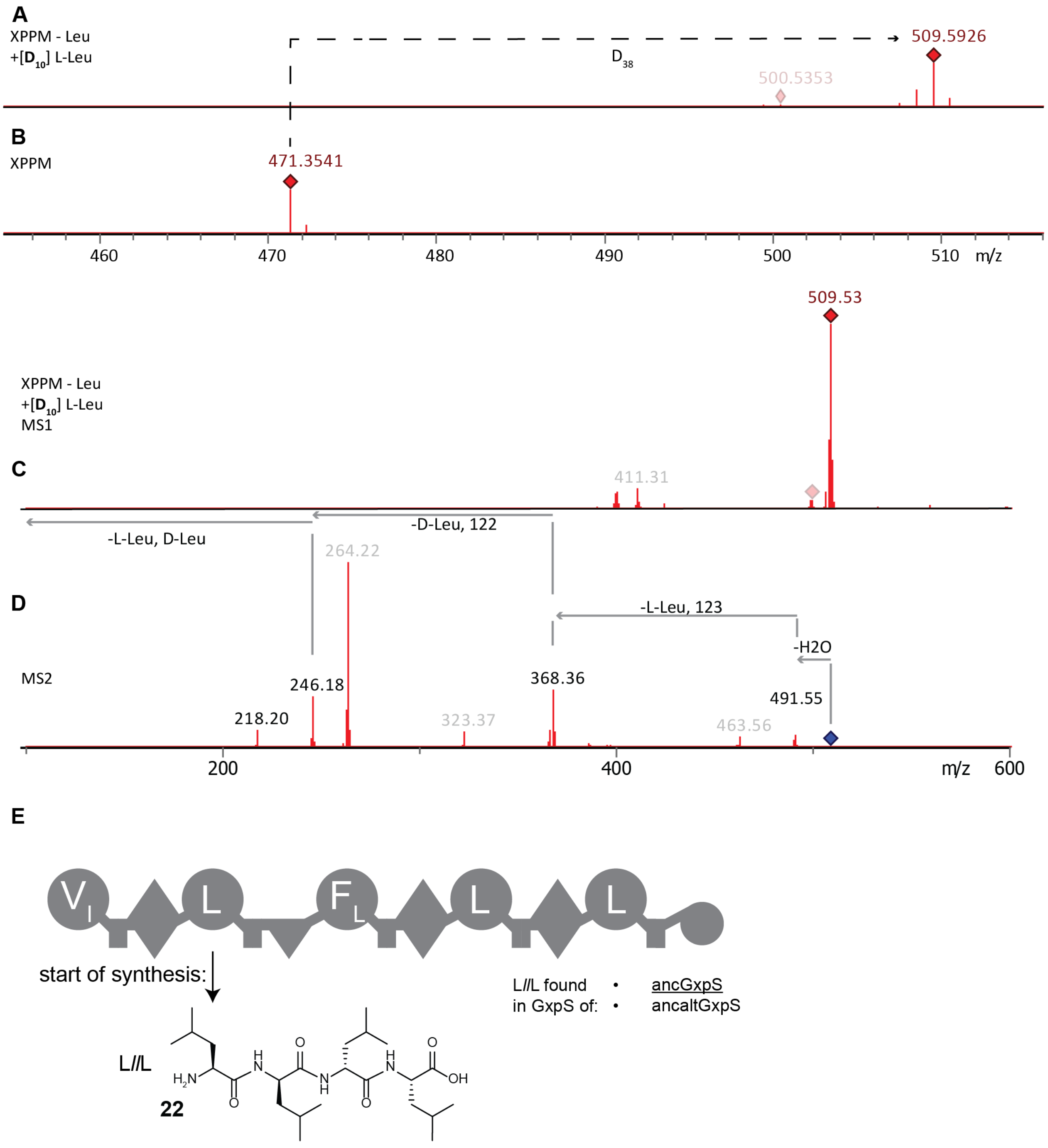


**Figure S19. Structural elucidation of the tetrapeptide L*l*L of the ancestral GxpS (22).** *ancgxpS* was heterologously expressed in *E. coli* DH10B$^{penta}$ together with pCK0403 and the produced natural products were extracted with methanol. (**A**) High-resolution mass of tetrapeptide L*l*L measured by HPLC-HRMS. Expression was done in XPPM without leucine supplemented with deuterated leucine (**B**) High-resolution mass of tetrapeptide L*l*L measured by HPLC-HRMS. Expression was done in XPPM. (**C**) Mass of tetrapeptide L*l*L measured by HPLC-MS. Expression was done in XPPM without leucine supplemented with deuterated leucine. (**D**) MS2 to elucidate the structure of tripeptide L*l*L measured by HPLC-MS. Expression was done in XPPM without leucine supplemented with deuterated leucine. (**E**) Diagram of ancestral GxpS. The start of the biosynthesis of the tetrapeptide starts at the second A-domain and is indicated by an arrow. Small, italic letters indicate D-amino acids.

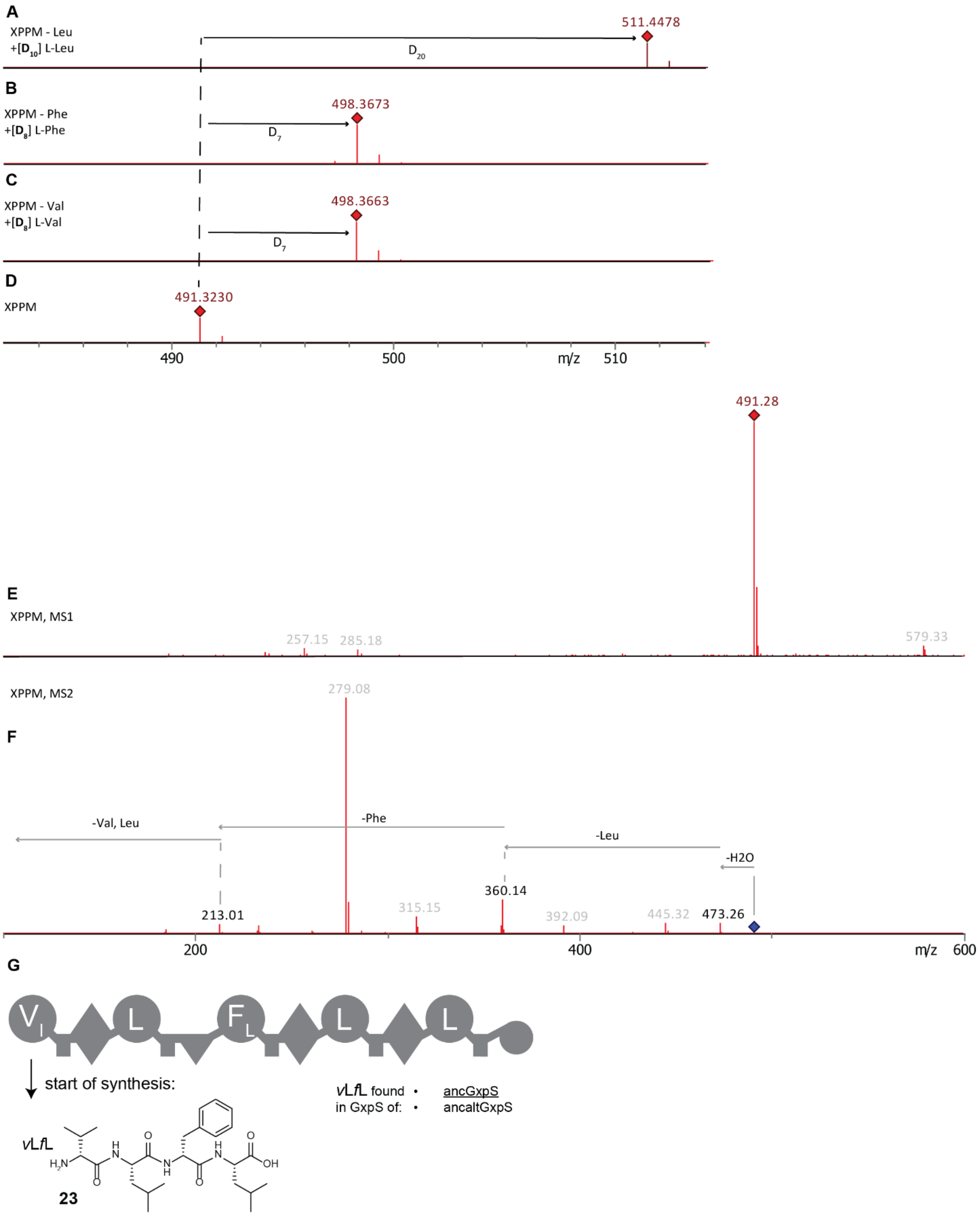


**Figure S20. Structural elucidation of the tetrapeptide *v*L*f*L of the ancestral GxpS (23).** *ancgxpS* was heterologously expressed in *E. coli* DH10B$^{penta}$ together with pCK0403 and the produced natural products were extracted with methanol. (**A**) High-resolution mass of tetrapeptide *v*L*f*L measured by

HPLC-HRMS. Expression was done in XPPM without leucine supplemented with deuterated leucine. (**B**) High-resolution mass of tetrapeptide *v*L*f*L measured by HPLC-HRMS. Expression was done in XPPM without phenylalanine supplemented with deuterated phenylalanine. (**C**) High-resolution mass of tetrapeptide *v*L*f*L measured by HPLC-HRMS. Expression was done in XPPM without valine supplemented with deuterated valine. (**D**) High-resolution mass of tetrapeptide *v*L*f*L measured by HPLC-HRMS. Expression was done in XPPM. (**E**) Mass of tetrapeptide *v*L*f*L measured by HPLC-MS. Expression was done in XPPM. (**F**) MS2 to elucidate the structure of tripeptide *v*L*f*L measured by HPLC-MS. Expression was done in XPPM. (**G**) Diagram of ancestral GxpS. The start of the biosynthesis of the tetrapeptide starts at the second A-domain and is indicated by an arrow. Small, italic letters indicate D-amino acids.

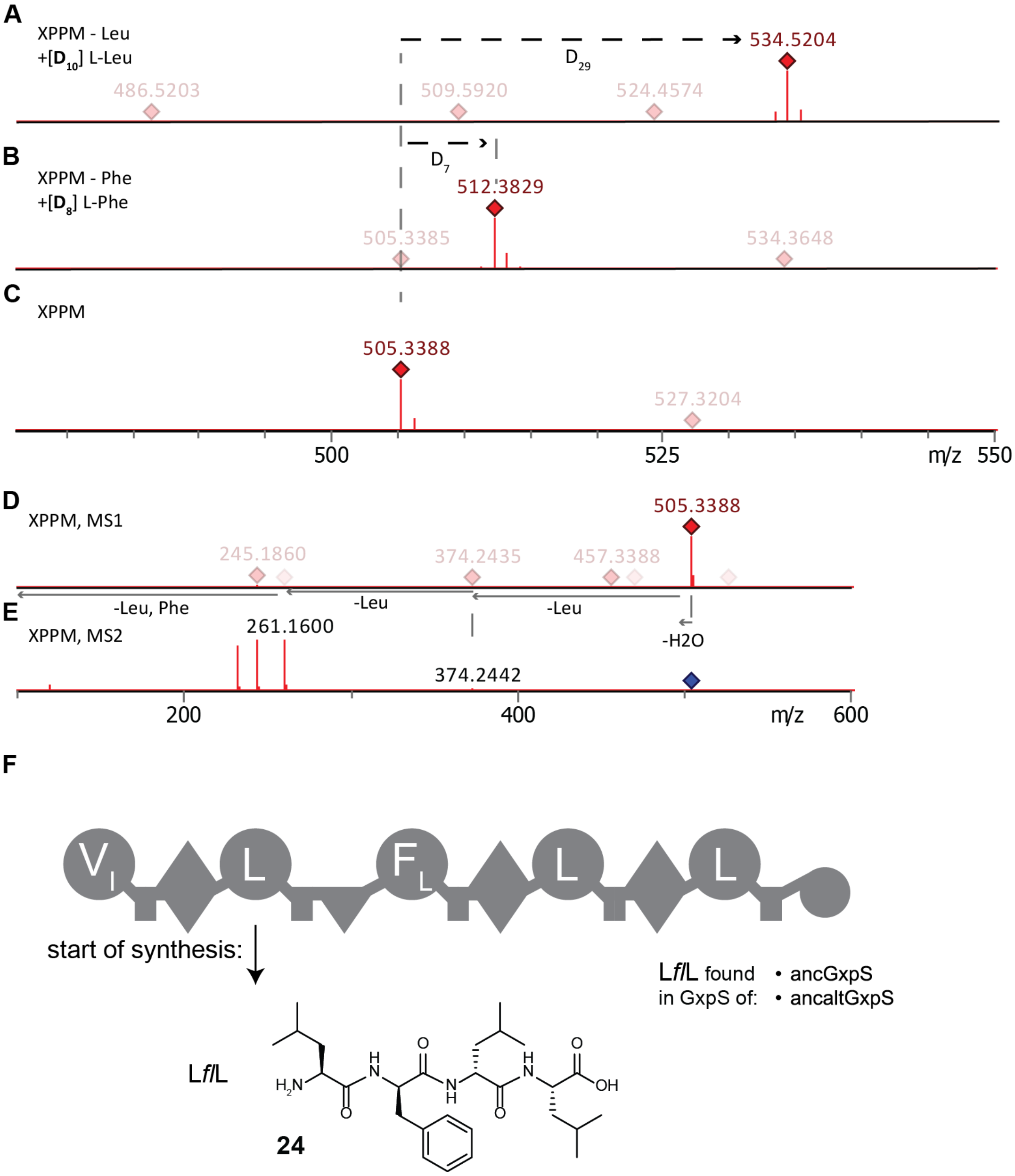


**Figure S21. Structural elucidation of the tetrapeptide L*f*L of the ancestral GxpS (24).** *ancgxpS* was heterologously expressed in *E. coli* DH10B$^{penta}$ together with pCK0403 and the produced natural products were extracted with methanol. (**A**) High resolution mass of tetrapeptide L*f*L measured by HPLC-HRMS. Expression was done in XPPM without leucine supplemented with deuterated leucine. (**B**) Mass of tetrapeptide L*f*L measured by HPLC-MS. Expression was done in XPPM without phenylalanine supplemented with deuterated phenylalanine. (**C**) High resolution mass of tetrapeptide L*f*L measured by HPLC-HRMS. Expression was done in XPPM. (**D**) Same as (C). (**E**) MS2 to elucidate the structure of tetrapeptide L*f*L measured by HPLC-HRMS. Expression was done in XPPM. (**F**) Diagram of ancestral GxpS. Small, italic letters indicate D-amino acids. The start of the biosynthesis of the tetrapeptide starts at the second A-domain and is indicated by an arrow.

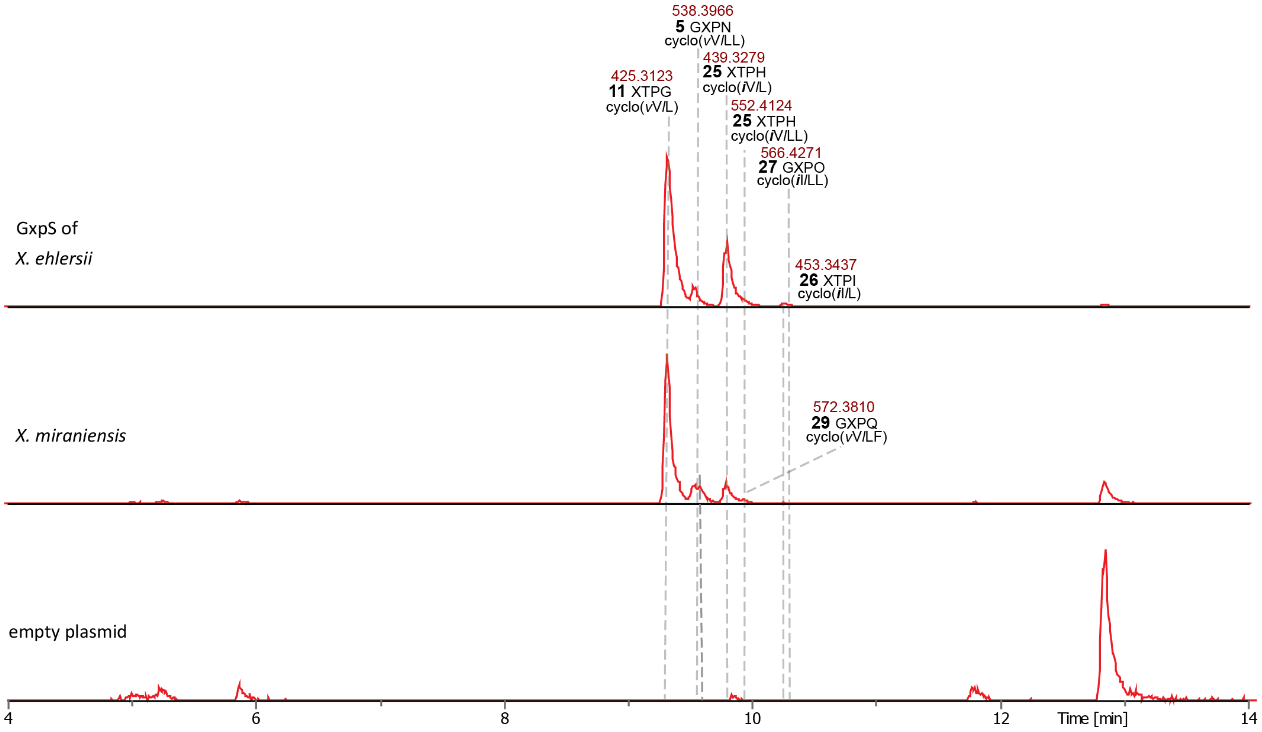


**Figure S22. EIC of all main peaks of GxpS from *X. ehlersii* and *X. miraniensis*.** Shown is the ionized high-resolution mass and the corresponding peptide. Small, italic letters indicate D-amino acids; ***i*** = D-*allo*-Ile.

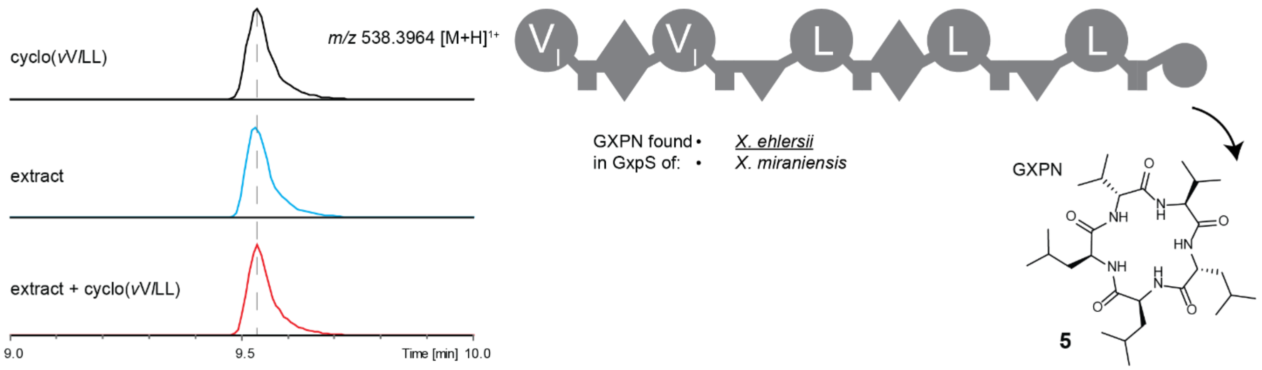


**Figure S23. Comparison of retention times of synthetic (black), natural (blue) and co-injected (red) GameXPeptide N (5)**. Small, italic letters indicate D-amino acids. GXPN was measured in a culture extract of *E. coli* DH10B$^{penta}$ carrying GxpS of *X. ehlersii* and pCK0403.

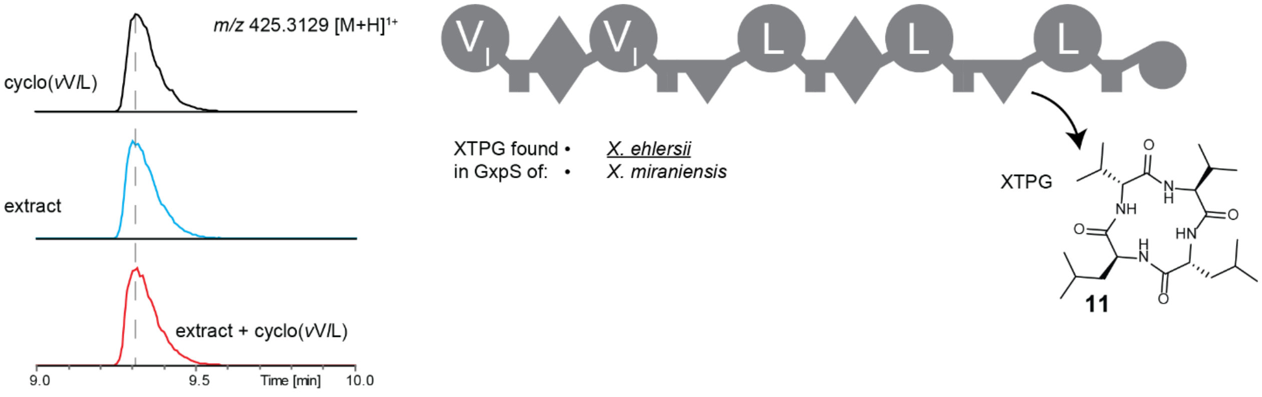


**Figure S24. Comparison of retention times of synthetic (black), natural (blue) and co-injected (red) XTPG (11)**. Small, italic letters indicate D-amino acids. XTPG was measured in a culture extract of *E. coli* DH10B$^{penta}$ carrying *gxpS* of *X. ehlersii* and pCK0403.

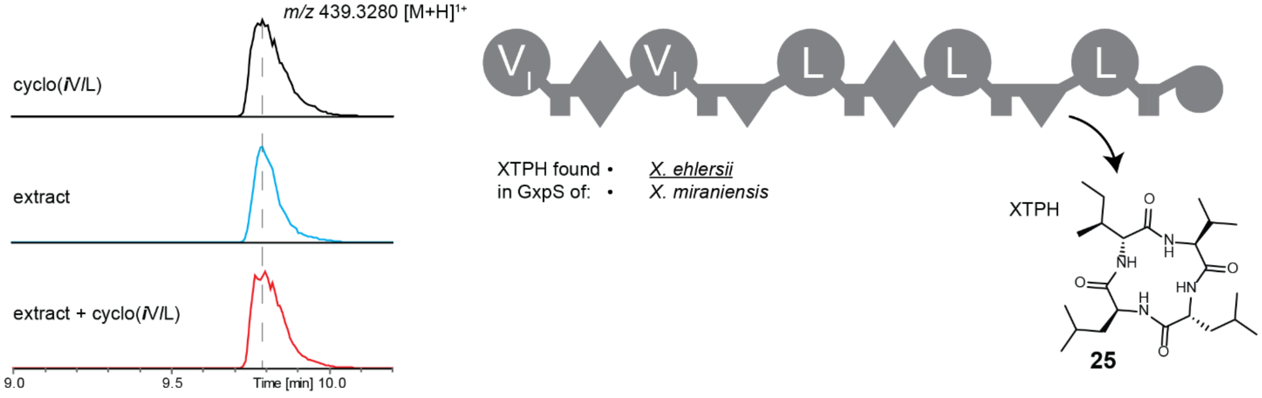


**Figure S25. Comparison of retention times of synthetic (black), natural (blue) and co-injected (red) XTPH (25)**. Small, italic letters indicate D-amino acids; ***i*** = D-*allo*-Ile. XTPH was measured in a culture extract of *E. coli* DH10B$^{penta}$ carrying *gxpS* of *X. ehlersii* and pCK0403.

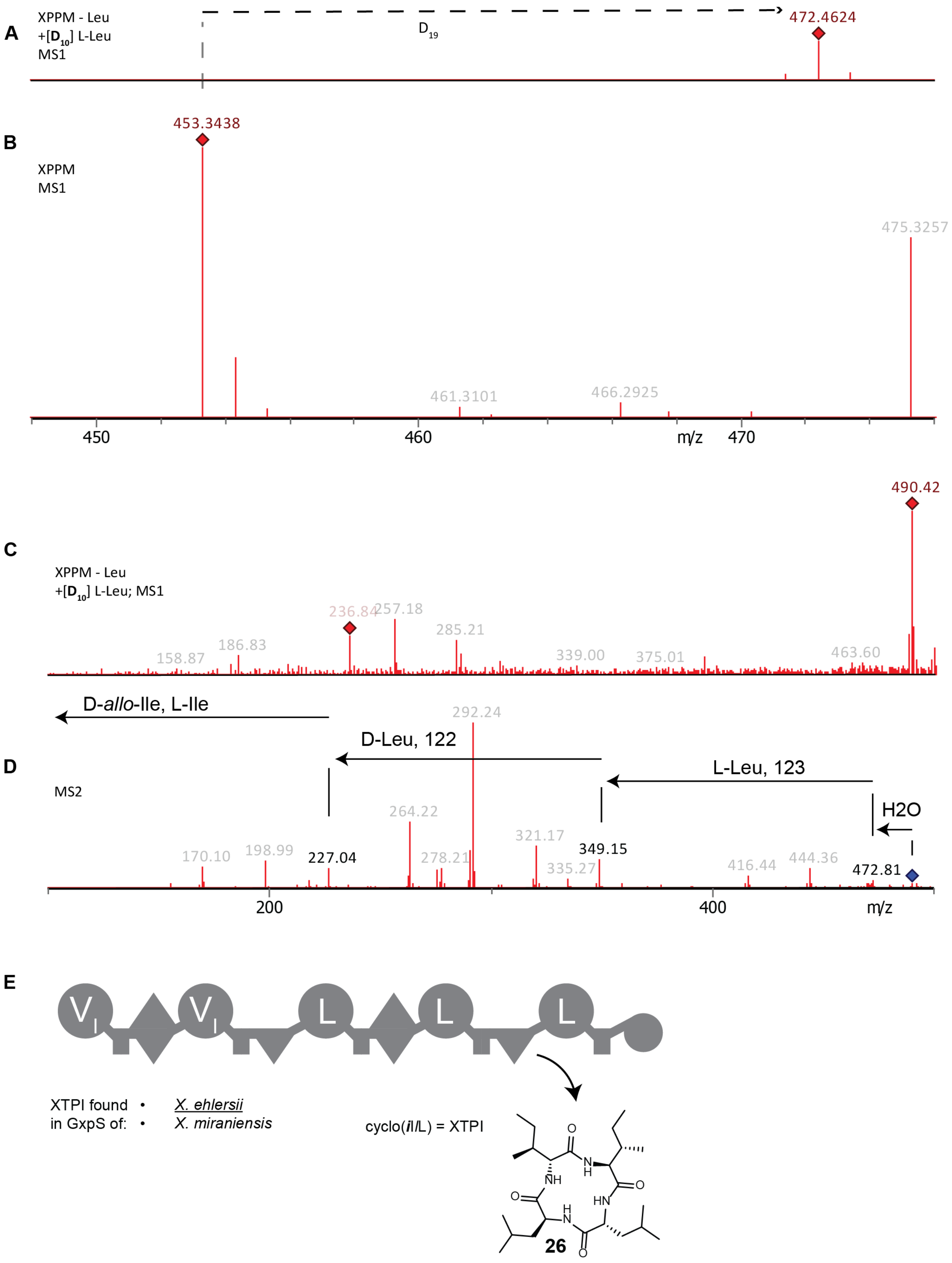


**Figure S26. Structural elucidation of the tetrapeptide XTPI (26) of the GxpS of *X. ehlersii*.** The *gxpS* of *X. ehlersii* was heterologously expressed in *E. coli* DH10B$^{penta}$ together with pCK0403 and the produced natural products were extracted with methanol. (**A**) High resolution mass of XTPI measured by HPLC-HRMS. Expression was done in XPPM without leucine supplemented with deuterated leucine. (**B**) High resolution mass of XTPI measured by HPLC-HRMS. Expression was done in XPPM. (**C**) Mass of XTPI measured by HPLC-MS. Expression was done in XPPM without leucine supplemented with deuterated leucine. (**D**) MS2 to elucidate the structure of XTPI measured by HPLC-MS. Expression was done in XPPM without leucine supplemented with deuterated leucine. (**E**) Diagram of GxpS from *X. ehlersii*. The peptide is released after the last C-domain as indicated by an arrow. Small, italic letters indicate D-amino acids; ***i*** = D-*allo*-Ile.

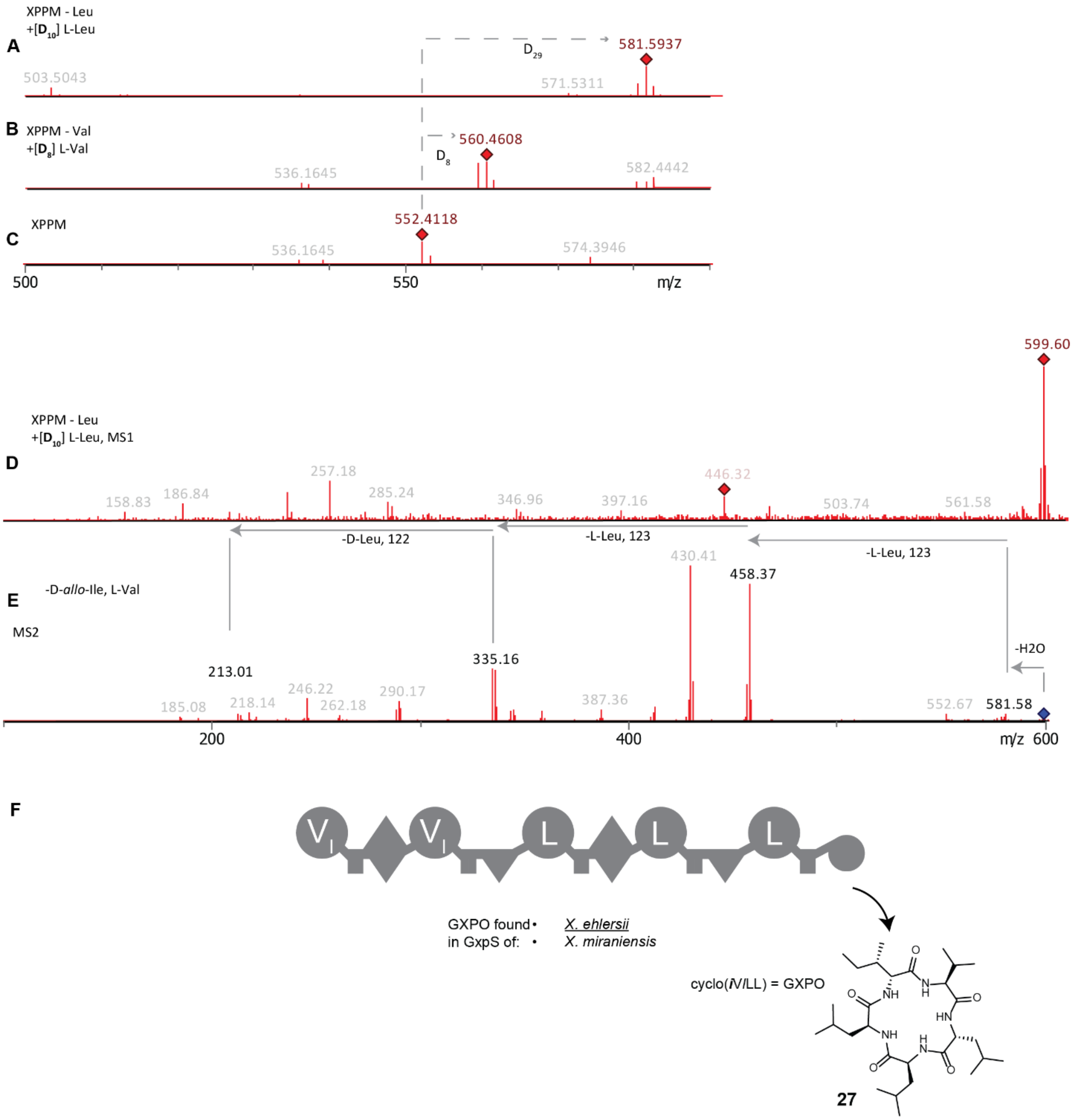


**Figure S27. Structural elucidation of the GXPO (27) of the GxpS of *X. ehlersii*.** *The gxpS of* X. ehlersii was heterologously expressed in *E. coli* DH10B$^{penta}$ together with pCK0403 and the produced natural products were extracted with methanol. (**A**) High-resolution mass of GXPO measured by HPLC-HRMS. Expression was done in XPPM without leucine supplemented with deuterated leucine. (**B**) High-resolution mass of GXPO measured by HPLC-HRMS. Expression was done in XPPM without phenylalanine supplemented with deuterated phenylalanine. (**C**) High-resolution mass of GXPO measured by HPLC-HRMS. Expression was done in XPPM. (**D**) Mass of linear GXPO measured by HPLC-MS. Expression was done in XPPM without leucine supplemented with deuterated leucine. (**E**) MS2 to elucidate the structure of linear GXPO measured by HPLC-MS. Expression was done in XPPM without leucine supplemented with deuterated leucine. (**F**) Diagram of GxpS from *X. ehlersii*. Shown is the release of the cyclic pentapeptide GXPO. Small, italic letters indicate D-amino acids; ***i*** = D-*allo*-Ile.

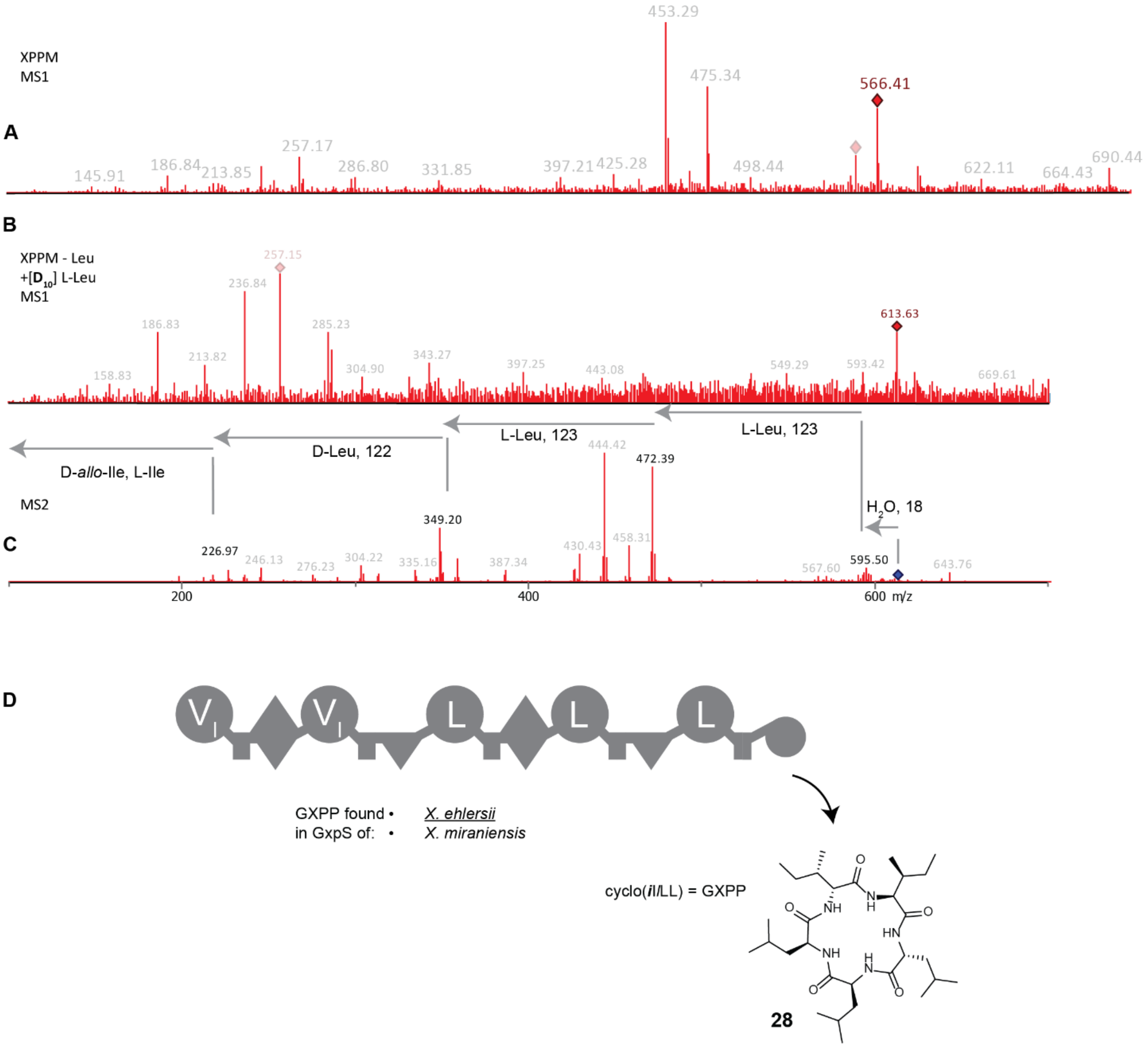


**Figure S28. Structural elucidation of GXPP (28) of the GxpS of *X. ehlersii*.** The *gxpS* of *X. ehlersii* was heterologously expressed in *E. coli* DH10B$^{penta}$ together with pCK0403 and the produced natural products were extracted with methanol. (**A**) Mass of GXPP measured by HPLC-MS. Expression was done in XPPM. (**B**) Mass of linear GXPP measured by HPLC-MS. Expression was done in XPPM without leucine supplemented with deuterated leucine. (**C**) MS2 of linear GXPP to elucidate the structure of the cyclic GXPP measured by HPLC-MS. Expression was done in XPPM without leucine supplemented with deuterated leucine. (**D**) Diagram of GxpS from *X. ehlersii*. Shown is the release of the cyclic pentapeptide GXPP. Small, italic letters indicate D-amino acids; ***i*** = D-*allo*-Ile.

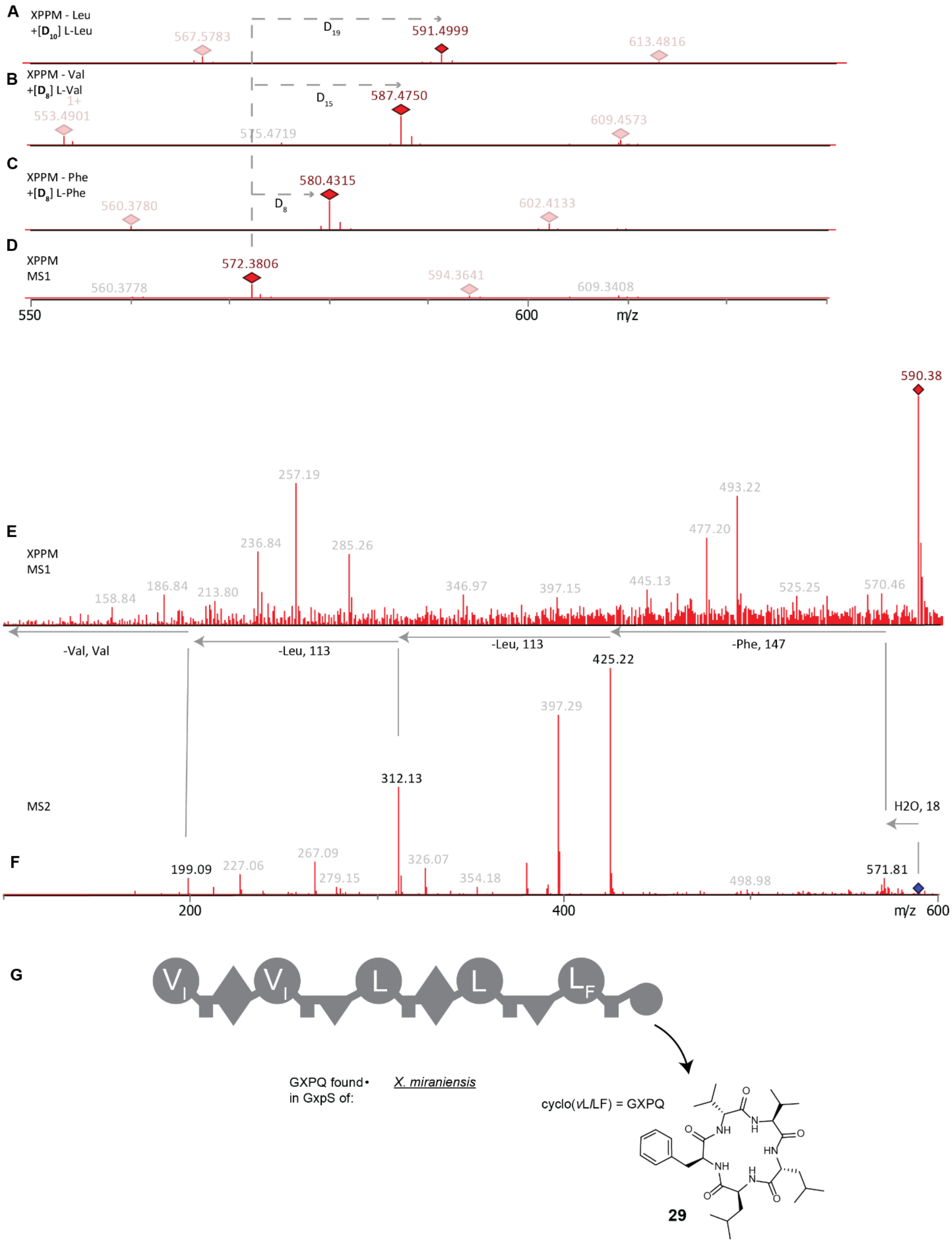


**Figure S29. Structural elucidation of GXPQ (29) of the GxpS of *X. miraniensis*.** *The gxpS of X. miraniensis* was heterologously expressed in *E. coli* DH10B$^{penta}$ together with pCK0413 and the produced natural products were extracted with methanol. (**A**) High resolution mass of GXPQ measured by HPLC-HRMS. Expression was done in XPPM without leucine supplemented with deuterated leucine. (**B**) High resolution mass of GXPQ measured by HPLC-HRMS. Expression was done in XPPM without valine supplemented with deuterated valine. (**C**) High resolution mass of GXPQ measured by HPLC-HRMS. Expression was done in XPPM without phenylalanine supplemented with deuterated phenylalanine. (**D**) High resolution mass of GXPQ measured by HPLC-HRMS. Expression was done in XPPM. (**E**) Mass of linearized GXPQ measured by HPLC-MS. Expression was done in XPPM. (**F**) MS2 of linear GXPQ to elucidate the structure of the cyclic GXPQ measured by HPLC-MS. Expression was done in XPPM. (**G**) Diagram of GxpS from *X. miraniensis*. Shown is the release of the cyclic pentapeptide GXPQ. Small, italic letters indicate D-amino acids.

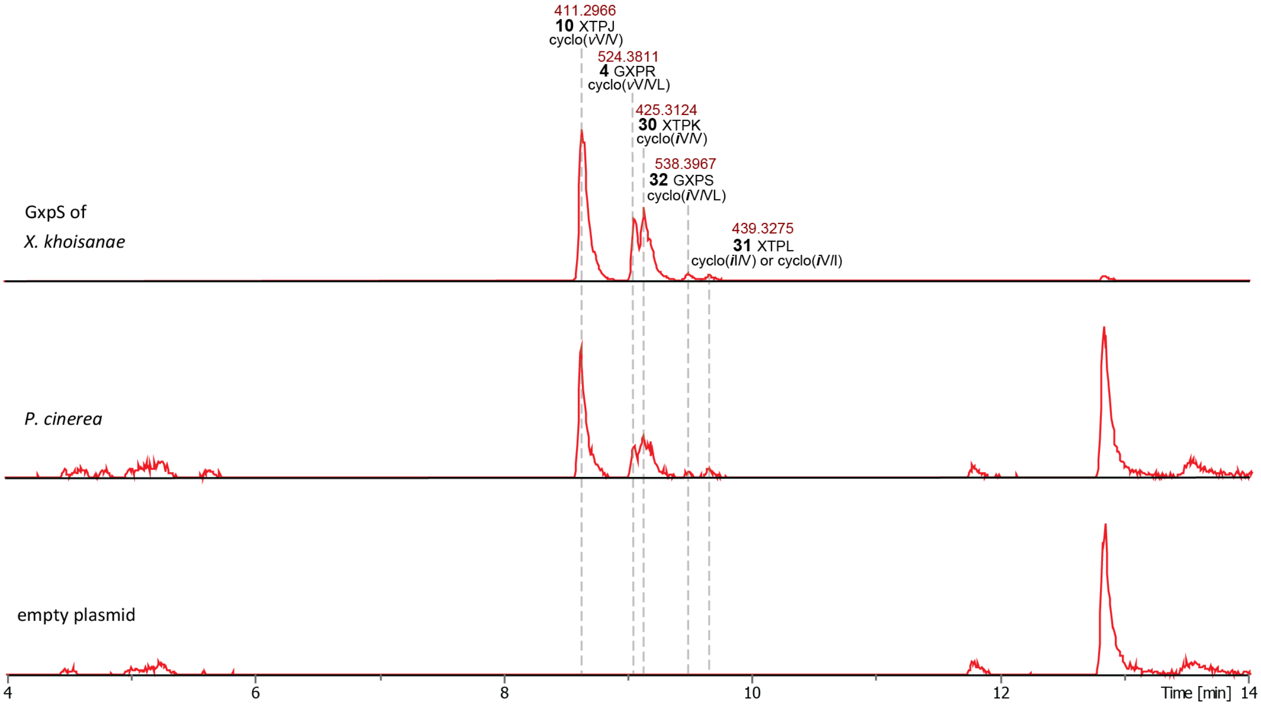


**Figure S30. EIC of all main peaks from GxpS from *X. khoisanae* and *P. cinerea*.** Shown is the ionized high-resolution mass and the corresponding peptide. Small, italic letters indicate D-amino acids; ***i*** = D-*allo*-Ile.

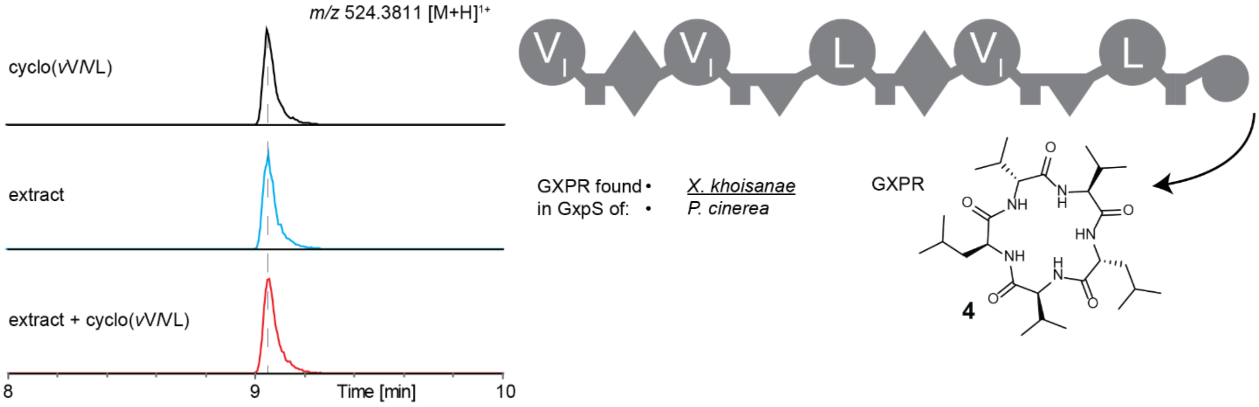


**Figure S31. Comparison of retention times of synthetic (black), natural (blue) and co-injected (red) GXPR (4)**. Small, italic letters indicate D-amino acids. GXPR (cyclo(*v*V*l*VL)) was measured in a culture extract of *E. coli* DH10B$^{penta}$ carrying *gxpS* of *X. khoisanae* and pCK0403.

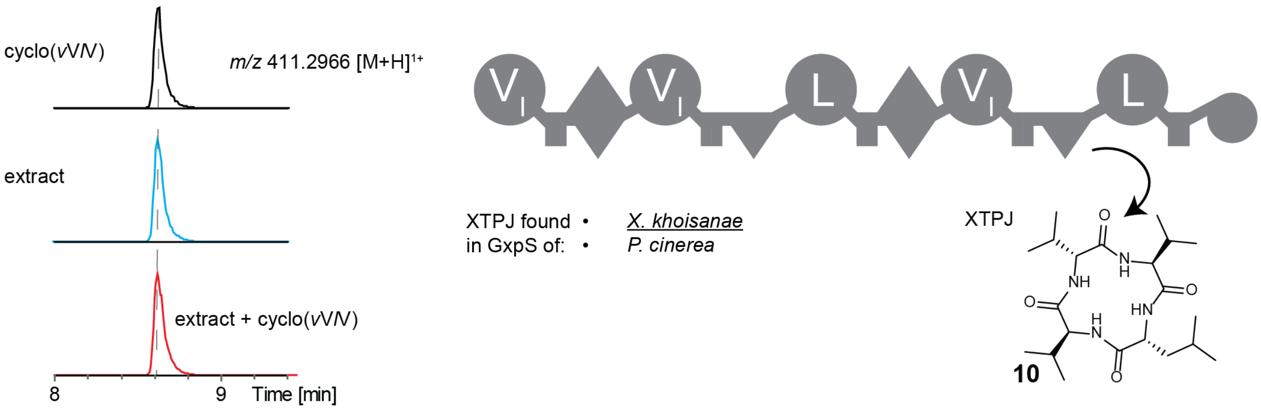


**Figure S32. Comparison of retention times of synthetic (black), natural (blue) and co-injected (red) XTPJ (10)**. Small, italic letters indicate D-amino acids. XTPJ (cyclo(*v*V*l*V)) was measured in a culture extract of *E. coli* DH10B$^{penta}$ carrying *gxpS of X. khoisanae and* pCK0403.

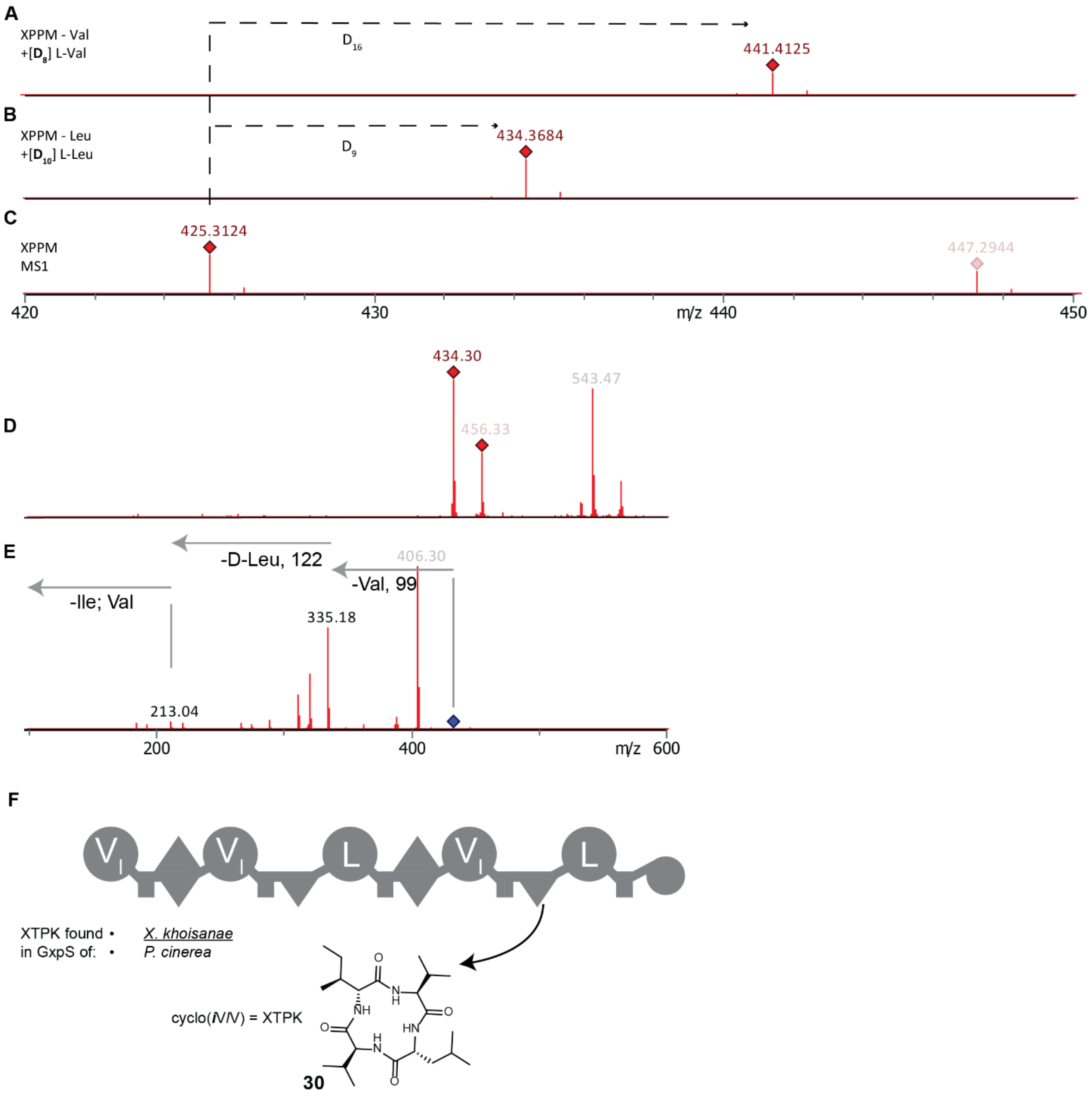


**Figure S33. Structural elucidation of XTPK (30) of the GxpS of *X. khoisanae*.** The *gxpS* of *X. khoisanae* was heterologously expressed in *E. coli* DH10B$^{penta}$ together with pCK0403 and the produced natural products were extracted with methanol. (**A**) High resolution mass of XTPK measured by HPLC-HRMS. Expression was done in XPPM without valine supplemented with deuterated valine. (**B**) High resolution mass of XTPK measured by HPLC-HRMS. Expression was done in XPPM without leucine supplemented with deuterated leucine. (**C**) High resolution mass of XTPK measured by HPLC-HRMS. Expression was done in XPPM. (**D**) Mass of XTPK measured by HPLC-HRMS. Expression was done in XPPM. (**E**) MS2 of XTPK to elucidate the structure of XTPK measured by HPLC-MS cultured in XPPM. (**F**) Diagram of GxpS from *X. khoisanae*. Shown is the release of the cyclic tetrapeptide XTPK at the last C-domain. Small, italic letters indicate D-amino acids; ***i*** = D-*allo*-Ile.

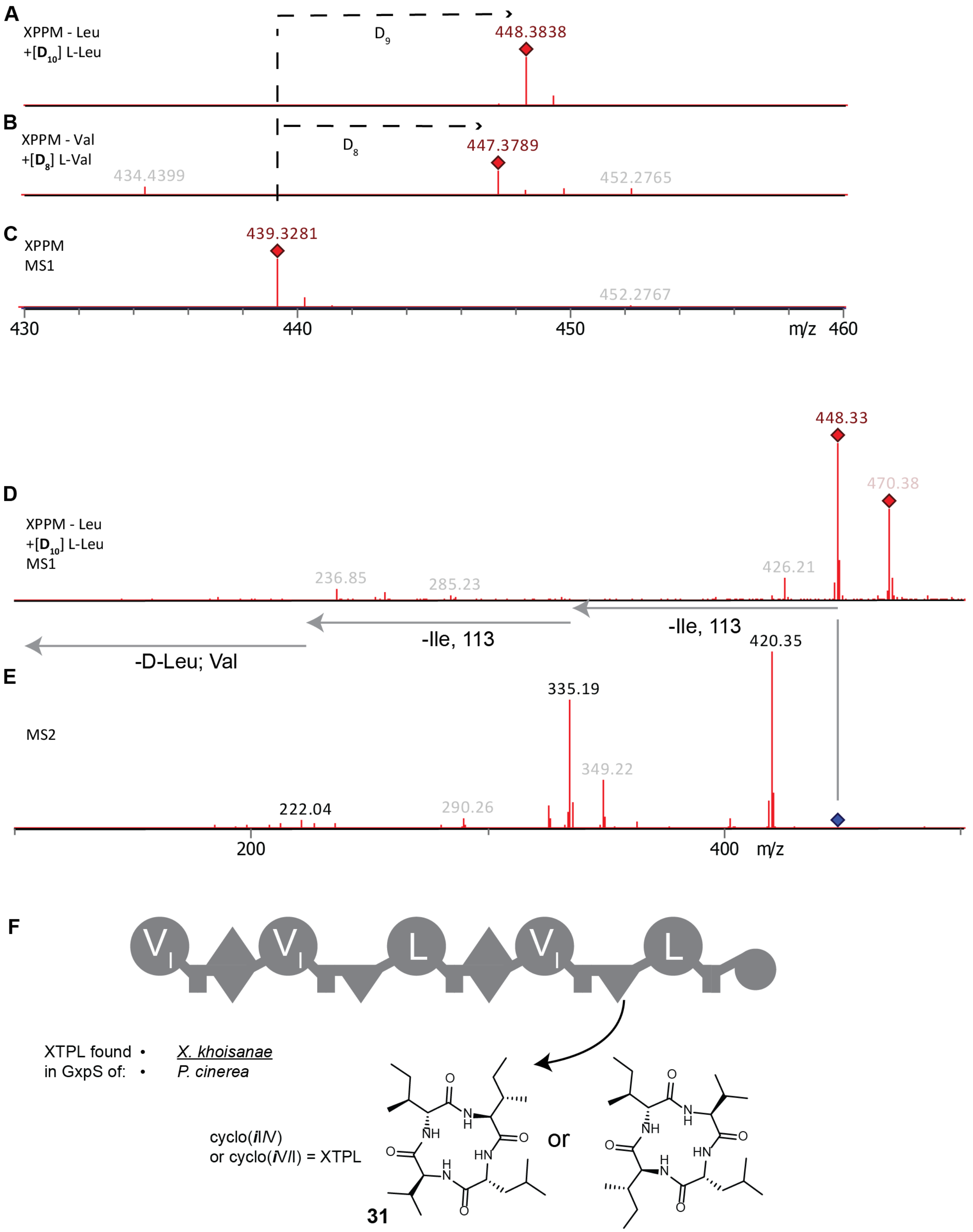


**Figure S34. Structural elucidation of XTPL (31) of the GxpS from *X. khoisanae*.** The *gxpS* of *X. khoisanae* was heterologously expressed in *E. coli* DH10B$^{penta}$ together with pCK0403 and the produced natural products were extracted with methanol. (**A**) High resolution mass of XTPL measured by HPLC-HRMS. Expression was done in XPPM without leucine supplemented with deuterated leucine. (**B**) High resolution mass of XTPL measured by HPLC-HRMS. Expression was done in XPPM without valine supplemented with deuterated valine. (**C**) High resolution mass of XTPL measured by HPLC-HRMS. Expression was done in XPPM. (**D**) Mass of XTPL measured by HPLC-HRMS. Expression was done in XPPM. (**E**) MS2 of XTPL to elucidate the structure of XTPL measured by HPLC-MS. Expression was done in XPPM. (**F**) Diagram of GxpS from *X. khoisanae*. Shown is the release of the cyclic tetrapeptide XTPL at the last C-domain. Small, italic letters indicate D-amino acids; ***i*** = D-*allo*-Ile.

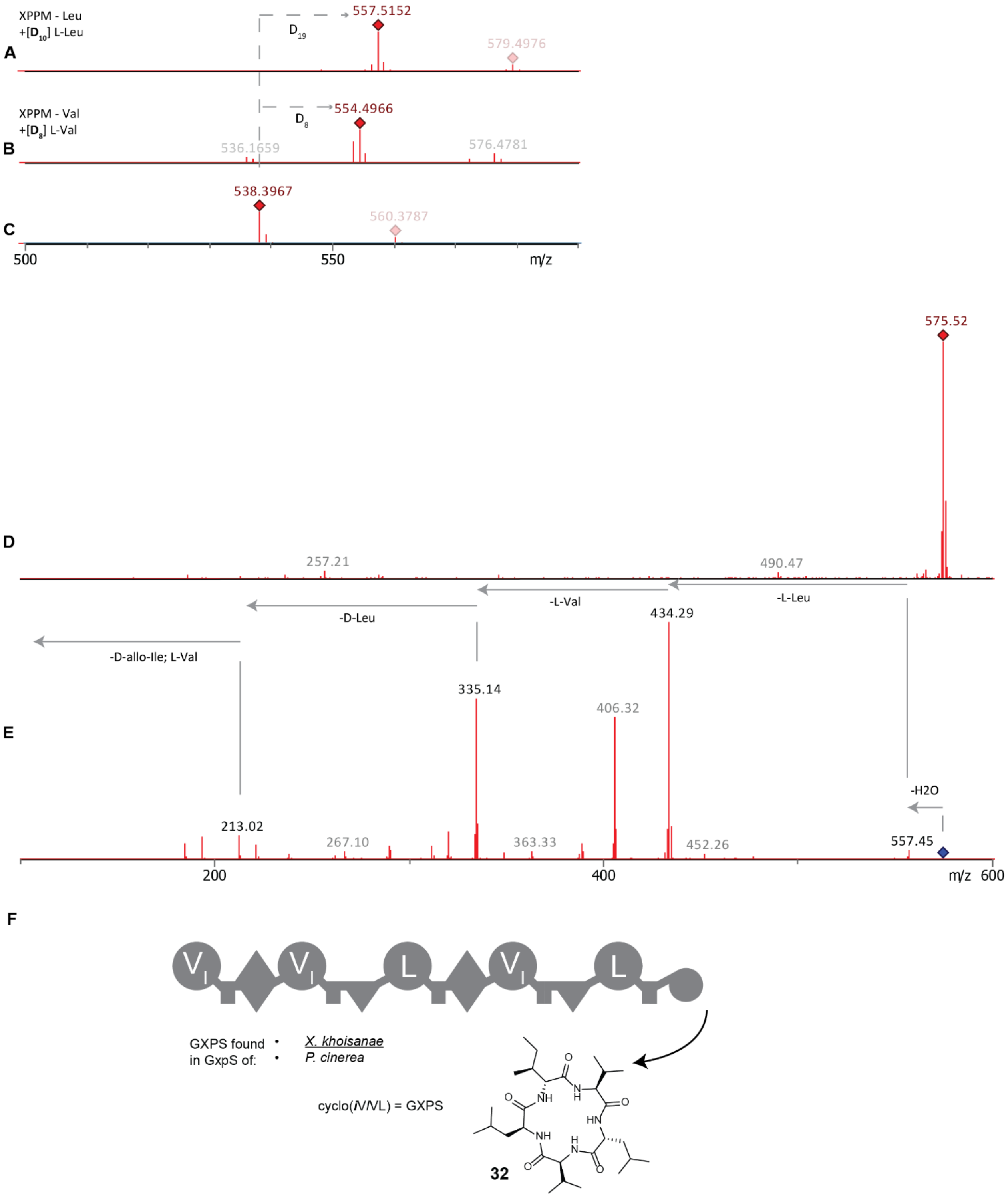


**Figure S35. Structural elucidation of GXPS (32) of the GxpS of *X. khoisanae*.** The *gxpS* of *X. khoisanae* was heterologously expressed in *E. coli* DH10B$^{penta}$ together with pCK0403 and the produced natural products were extracted with methanol. (**A**) High resolution mass of GXPS measured by HPLC-HRMS. Expression was done in XPPM without leucine supplemented with deuterated leucine. (**B**) High resolution mass of GXPS measured by HPLC-HRMS. Expression was done in XPPM without valine supplemented with deuterated valine. (**C**) High resolution mass of GXPS measured by HPLC-HRMS. Expression was done in XPPM. (**D**) Mass of GXPS measured by HPLC-MS. Expression was done in XPPM without leucine supplemented with deuterated leucine. (**E**) MS2 of GXPS to elucidate the structure of GXPS measured by HPLC-MS. Expression was done in XPPM without leucine supplemented with deuterated leucine. (**F**) Diagram of GxpS from *X. khoisanae* producing the pentapeptide GXPS. Small, italic letters indicate D-amino acids; ***i*** = D-*allo*-Ile.

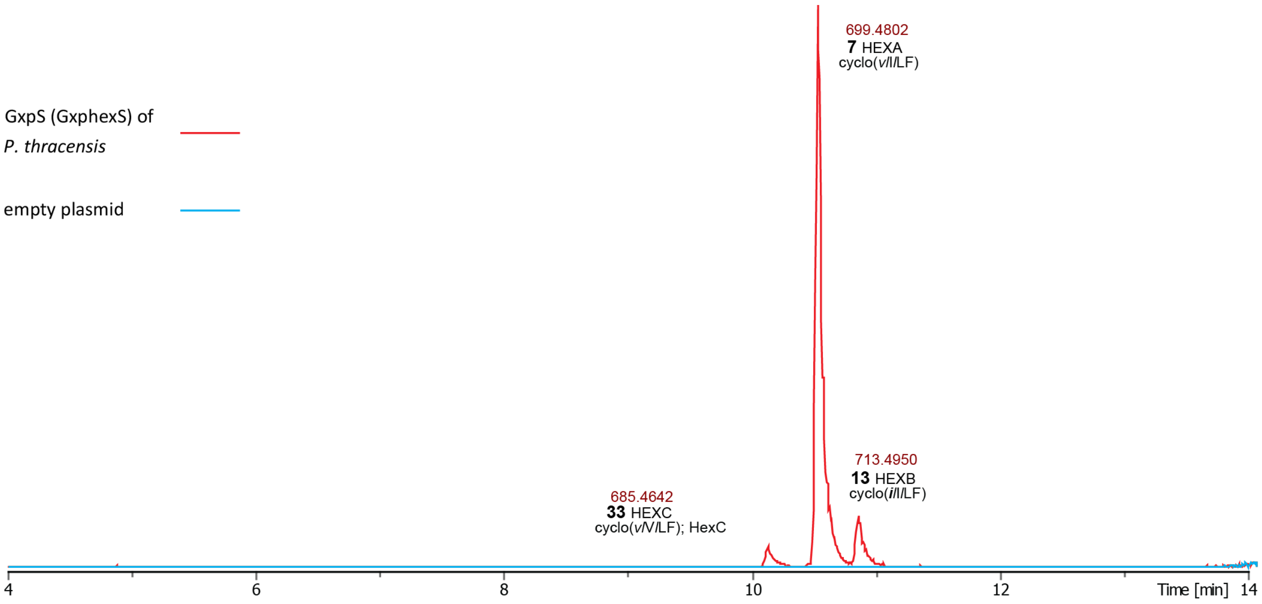


**Figure S36. EIC of all main peaks from the GxpS-derivative HexS from *P. thracensis*.** Shown is the ionized high-resolution mass and the corresponding peptide. Small, italic letters indicate D-amino acids; ***i*** = D-*allo*-Ile.

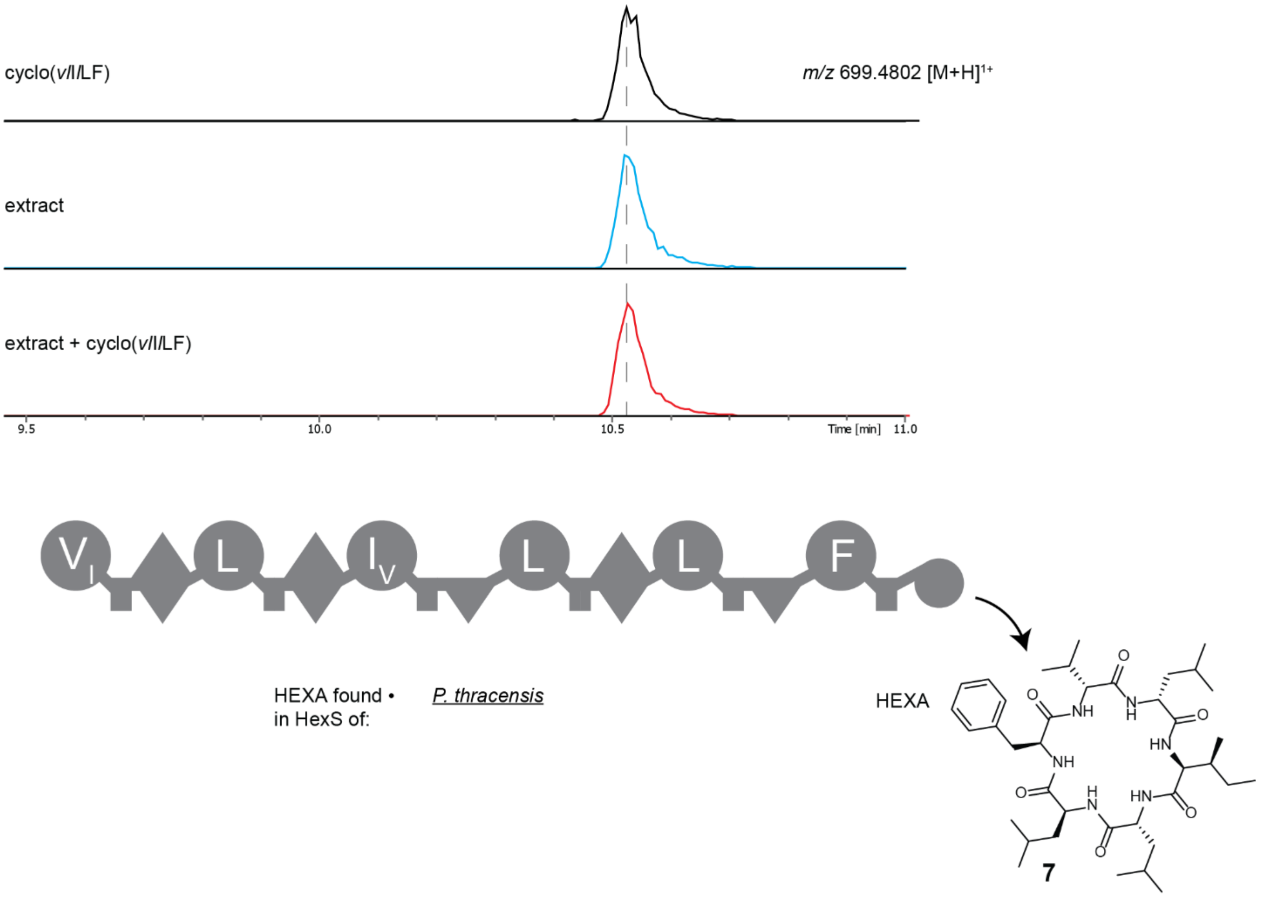


**Figure S37**. **Comparison of retention times of synthetic (black), natural (blue) and co-injected (red) HEXA (7)**. Small, italic letters indicate D-amino acids. HEXA (cyclo(*v*I*l*LF)) was measured in a culture extract of *E. coli* DH10B$^{penta}$ carrying *hexS* of *P. thracensis* and pCK0403.

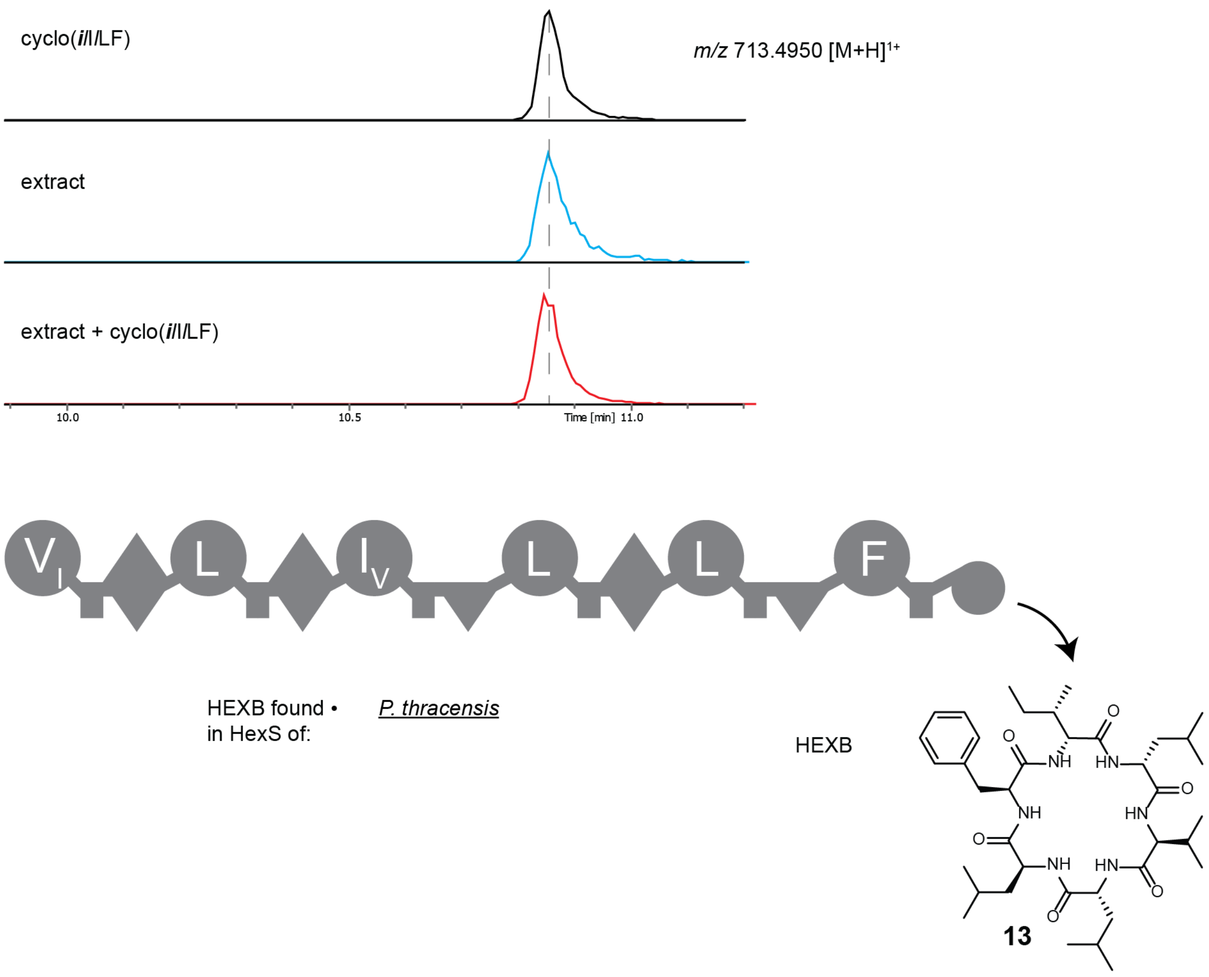


**Figure S38. Comparison of retention times of synthetic (black), natural (blue) and co-injected (red) HEXB (13).** Small, italic letters indicate D-amino acids; ***i*** = D-*allo*-Ile. HEXB (cyclo(***i***I*l*LF)) was measured in a culture extract of *E. coli* DH10B$^{penta}$ carrying GxpS of *P. thracensis* and pCK0403.

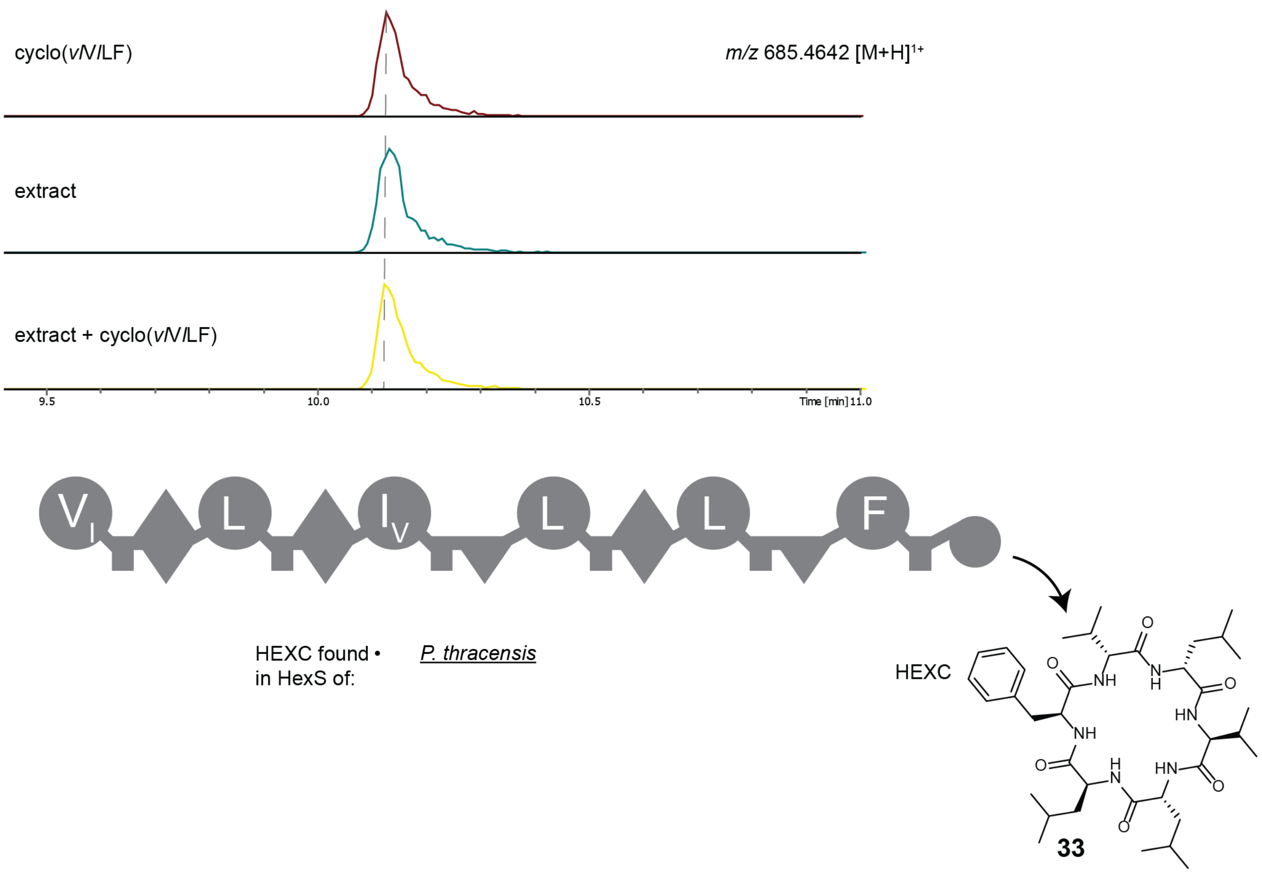


**Figure S39. Comparison of retention times of synthetic (black), natural (blue) and co-injected (red) HEXC (33)**. Small, italic letters indicate D-amino acids. HEXC (cyclo(*vl*V*l*LF)) was measured in a culture extract of *E. coli* DH10B$^{penta}$ carrying *hexS* of *P. thracensis* and pCK0403.

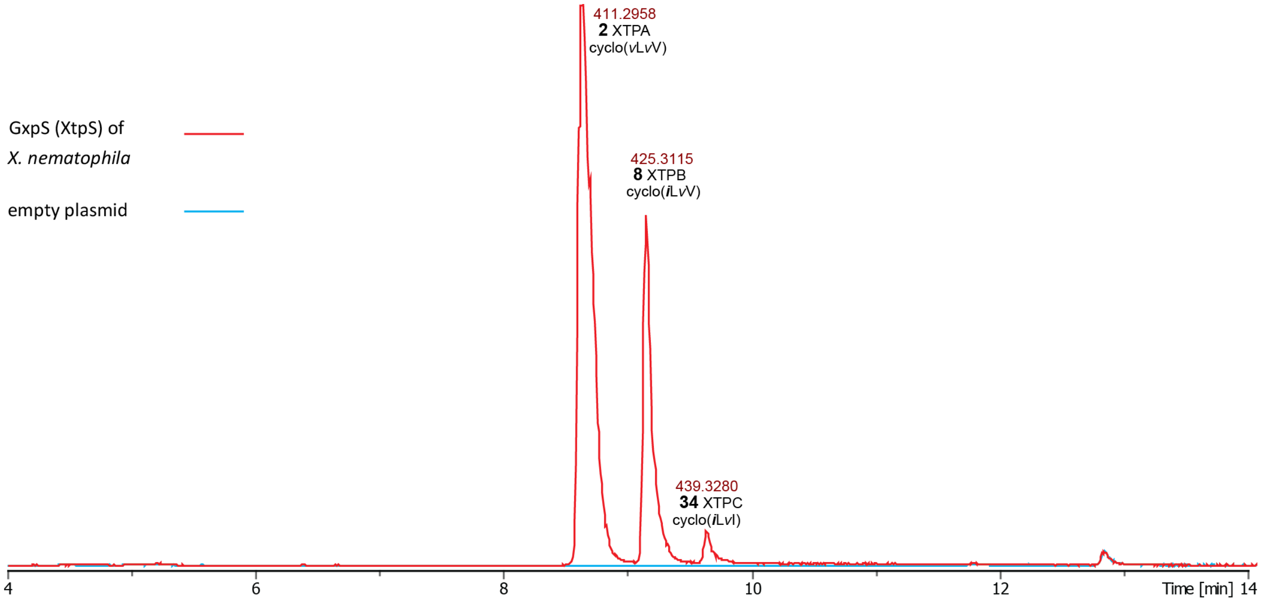


**Figure S40. EIC of all main peaks from the GxpS derivative XtpS from *X. nematophila*.** Shown are the ionized high-resolution mass and the corresponding peptide.

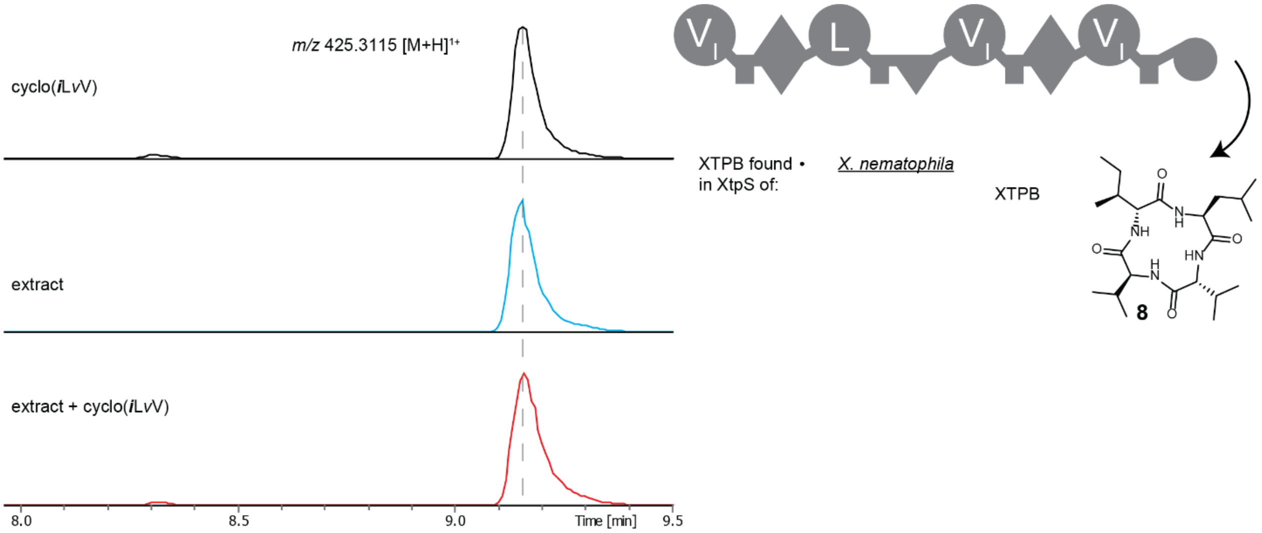


**Figure S41. Comparison of retention times of synthetic (black), natural (blue) and co-injected (red) XTPB (8)**. Small, italic letters indicate D-amino acids; ***i*** = D-*allo*-Ile. XTPB (cyclo(***i***L*v*V)) was measured in a culture extract of *E. coli* DH10B$^{penta}$ carrying *xtpS* of *X. nematophila* and pCK0403.

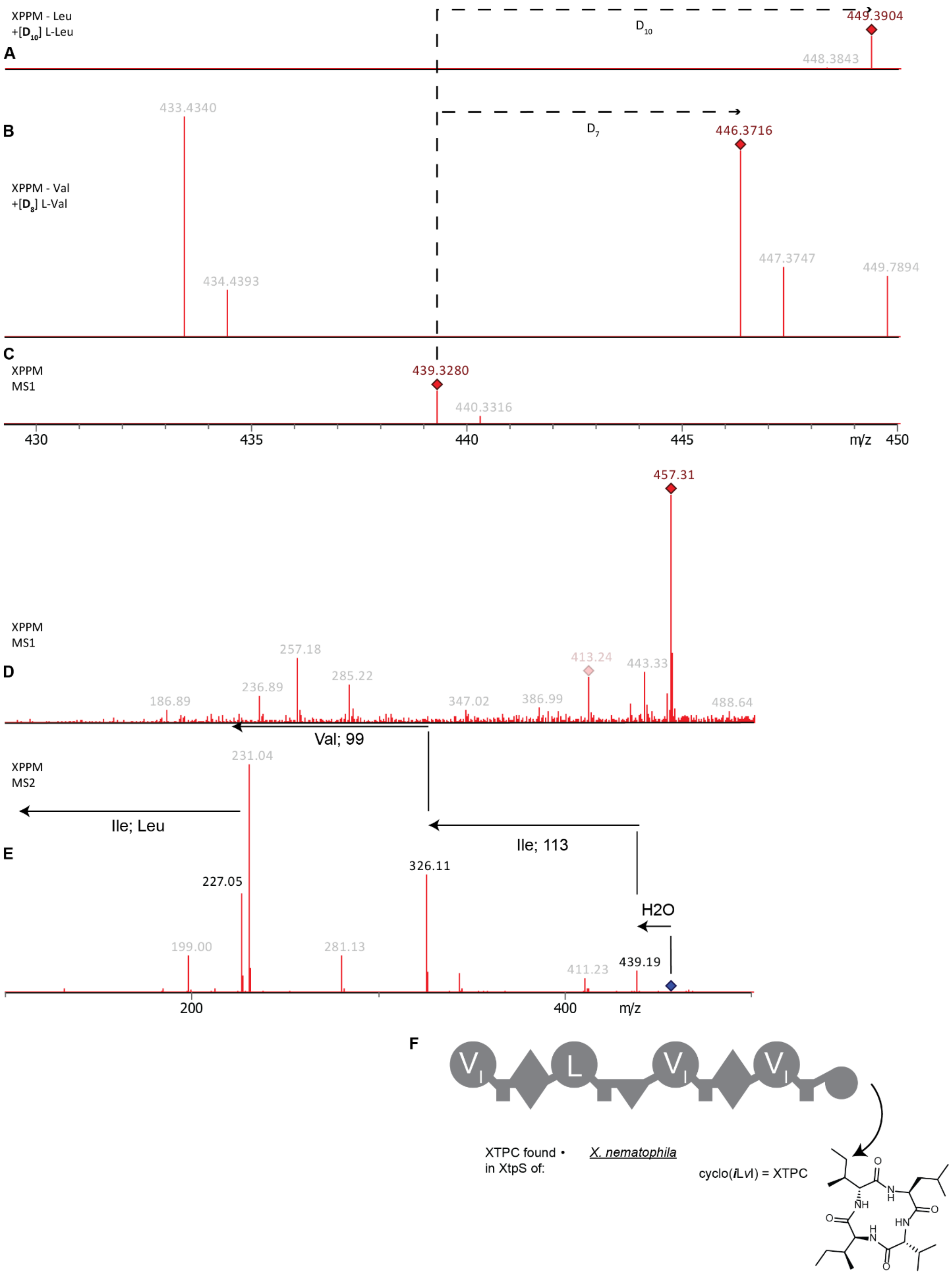

XPPM - Leu
+[D10] L-Leu
A
D10
449.3904
448.3843
B
433.4340
D7
446.3716
XPPM - Val
+[D8] L-Val
447.3747
449.7894
434.4393
C
XPPM
MS1
439.3280
440.3316
430
435
440
445
m/z
450
457.31
XPPM
MS1
257.18
413.24
443.33
D
236.89
285.22
186.89
347.02
386.99
488.64
Val; 99
XPPM
MS2
231.04
Ile; Leu
Ile; 113
E
326.11
227.05
H2O
199.00
281.13
411.23
439.19
200
400
m/z
F
V
L
V
V
XTPC found • in XtpS of:
X. nematophila
cyclo(iLvI) = XTPC


**34**

**Figure S42. Structural elucidation of XTPC (34) of XtpS of *X. nematophila*.** *The xtpS of X. nematophila* was heterologously expressed in *E. coli* DH10B$^{penta}$ together with pCK0403 and the produced natural products were extracted with methanol. (**A**) High resolution mass of XTPC measured by HPLC-HRMS. Expression was done in XPPM without leucine supplemented with deuterated leucine. (**B**) High resolution mass of XTPC measured by HPLC-HRMS. Expression was done in XPPM without valine supplemented with deuterated valine. (**C**) High resolution mass of XTPC measured by HPLC-HRMS. Expression was done in XPPM. (**D**) Mass of linear XTPC measured by HPLC-MS cultured in XPPM. (**E**) MS2 of linear XTPC to elucidate the structure of XTPC measured by HPLC-MS cultured in XPPM. (**F**) Diagram of XtpS from *X. nematophila* producing XTPC.

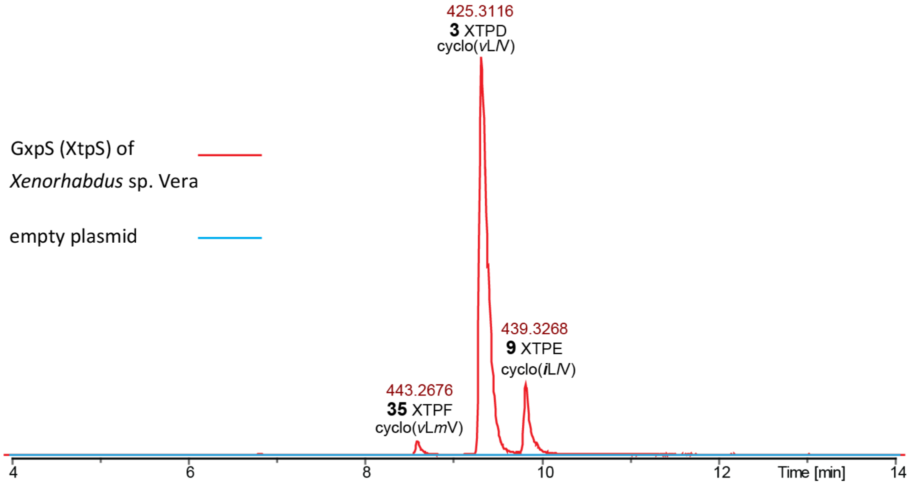


**Figure S43. EIC of all main peaks from the GxpS-derivative XtpS from *Xenorhabdus* sp. Vera.** Shown is the ionized high-resolution mass and the corresponding peptide.

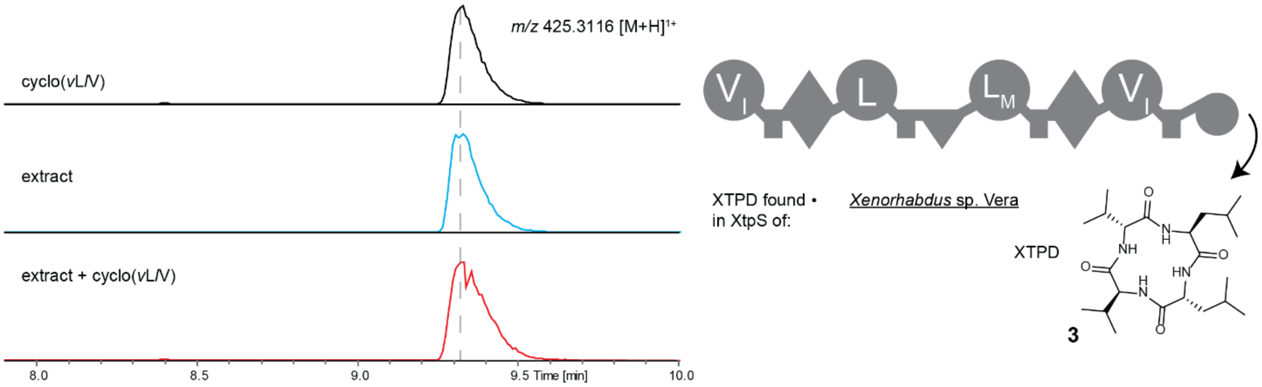


**Figure S44. Comparison of retention times of synthetic (black), natural (blue) and co-injected (red) XTPD (3)**. Small, italic letters indicate D-amino acids. XTPD (cyclo(*v*L*l*V)) was measured in a culture extract of *E. coli* DH10B$^{penta}$ carrying *xtpS* of *Xenorhabdus* sp. Vera and pCK0403.

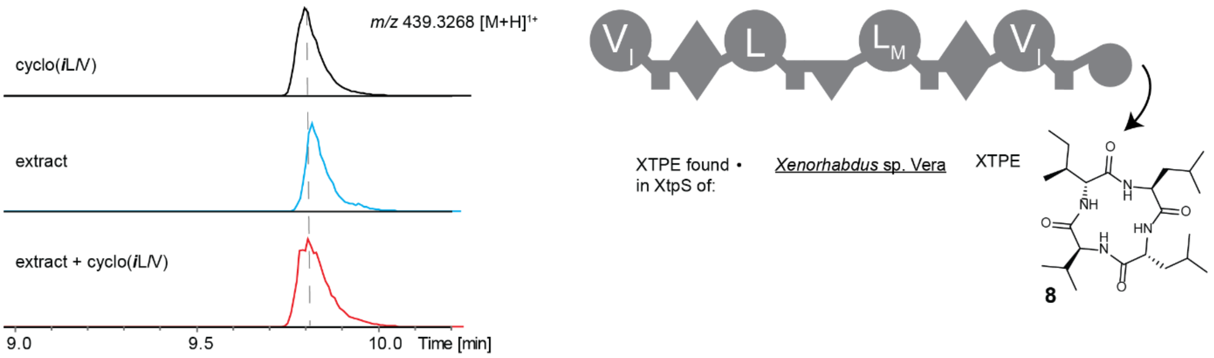


**Figure S45. Comparison of retention times of synthetic (black), natural (blue) and co-injected (red) XTPE (8)**. Small, italic letters indicate D-amino acids. XTPE was measured in a culture extract of *E. coli* DH10B$^{penta}$ carrying *xtpS* of *Xenorhabdus* sp. Vera and pCK0403.

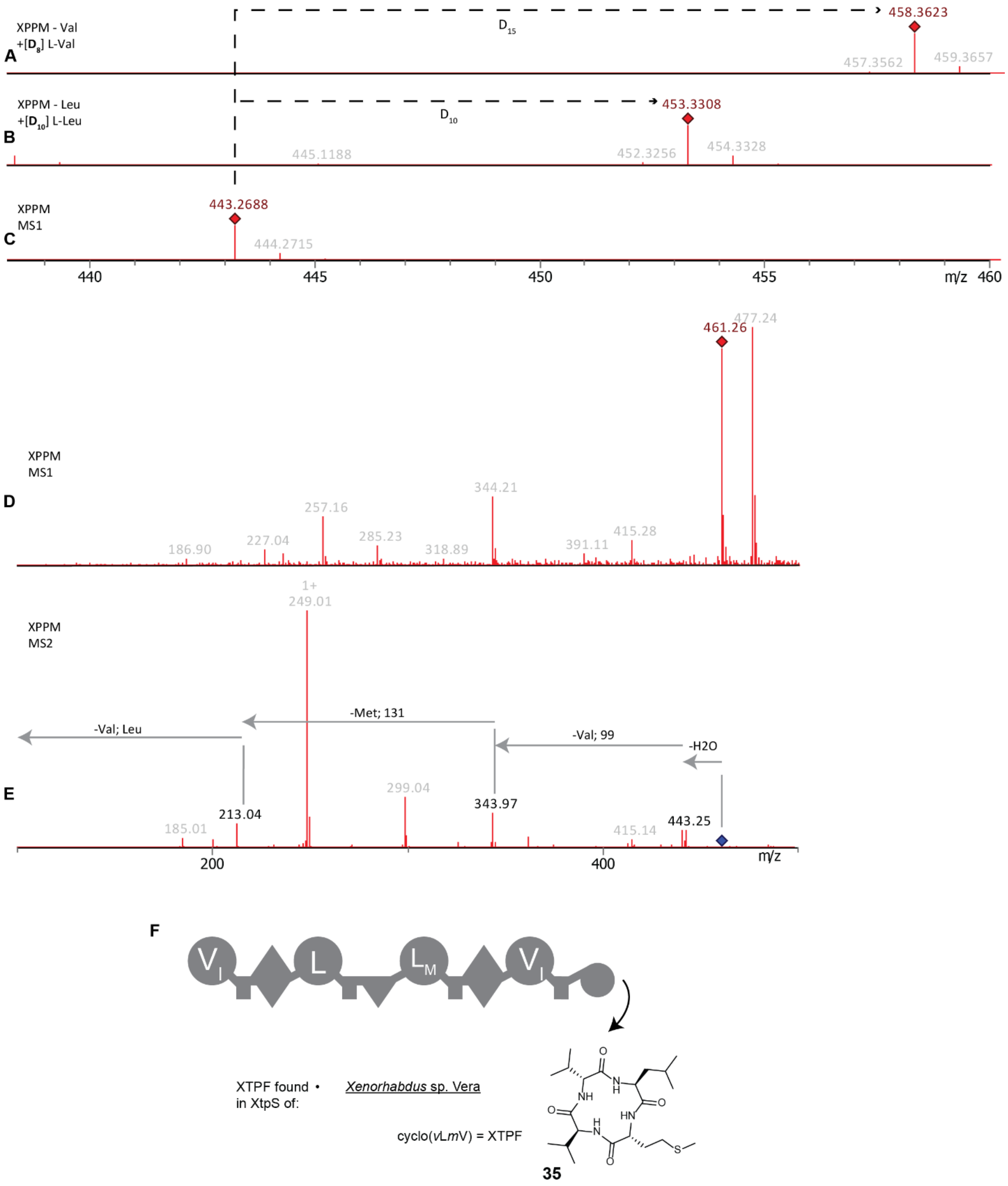


**Figure S46. Structural elucidation of XTPF (35) of XtpS of *Xenorhabdus* sp. Vera.** The *xtpS* of *Xenorhabdus* sp. Vera was heterologously expressed in *E. coli* DH10B$^{penta}$ together with pCK0403 and the produced natural products were extracted with methanol. (**A**) High resolution mass of XTPF (**35**) measured by HPLC-HRMS. Expression was done in XPPM without valine supplemented with deuterated valine. (**B**) High resolution mass of XTPF measured by HPLC-HRMS. Expression was done in XPPM without leucine supplemented with deuterated leucine. (**C**) High resolution mass of XTPF measured by HPLC-HRMS. Expression was done in XPPM. (**D**) Mass of linear XTPF measured by HPLC-MS. Expression was done in XPPM. (**E**) MS2 of linear XTPF to elucidate the structure of XTPF measured by HPLC-MS. Expression was done in XPPM. (**F**) Diagram of XtpS from *Xenorhabdus* sp. Vera producing XTPF.

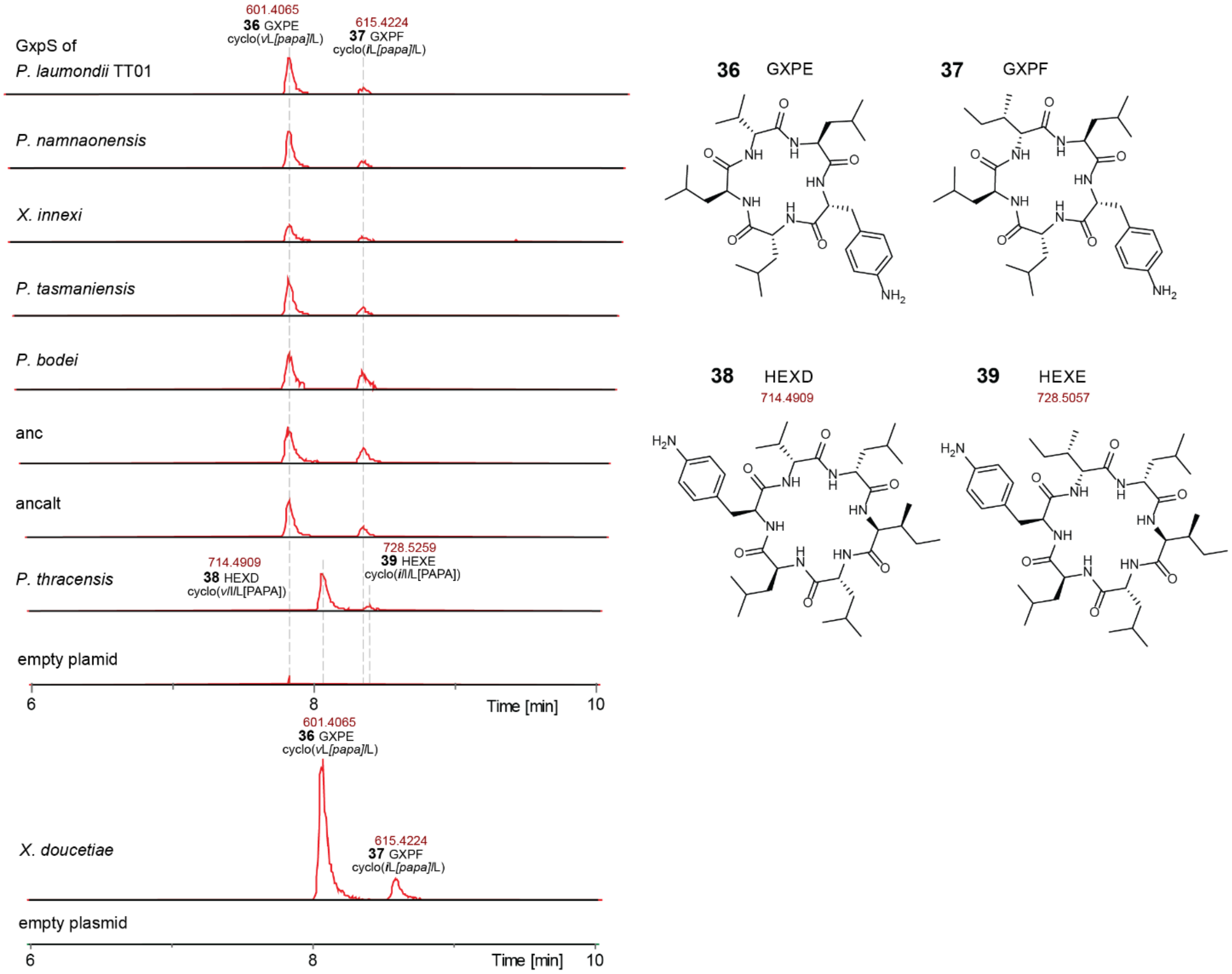


**Figure S47: EIC of all main peaks from different GxpS that can accept PAPA.** Shown are the ionized high-resolution mass and the corresponding peptide. *E. coli* carrying different *gxpS* orthologs and pCK0403 were cultured in XPPM supplemented with 2 mM synthetic PAPA. Natural product production was measured by HPLC-HRMS.

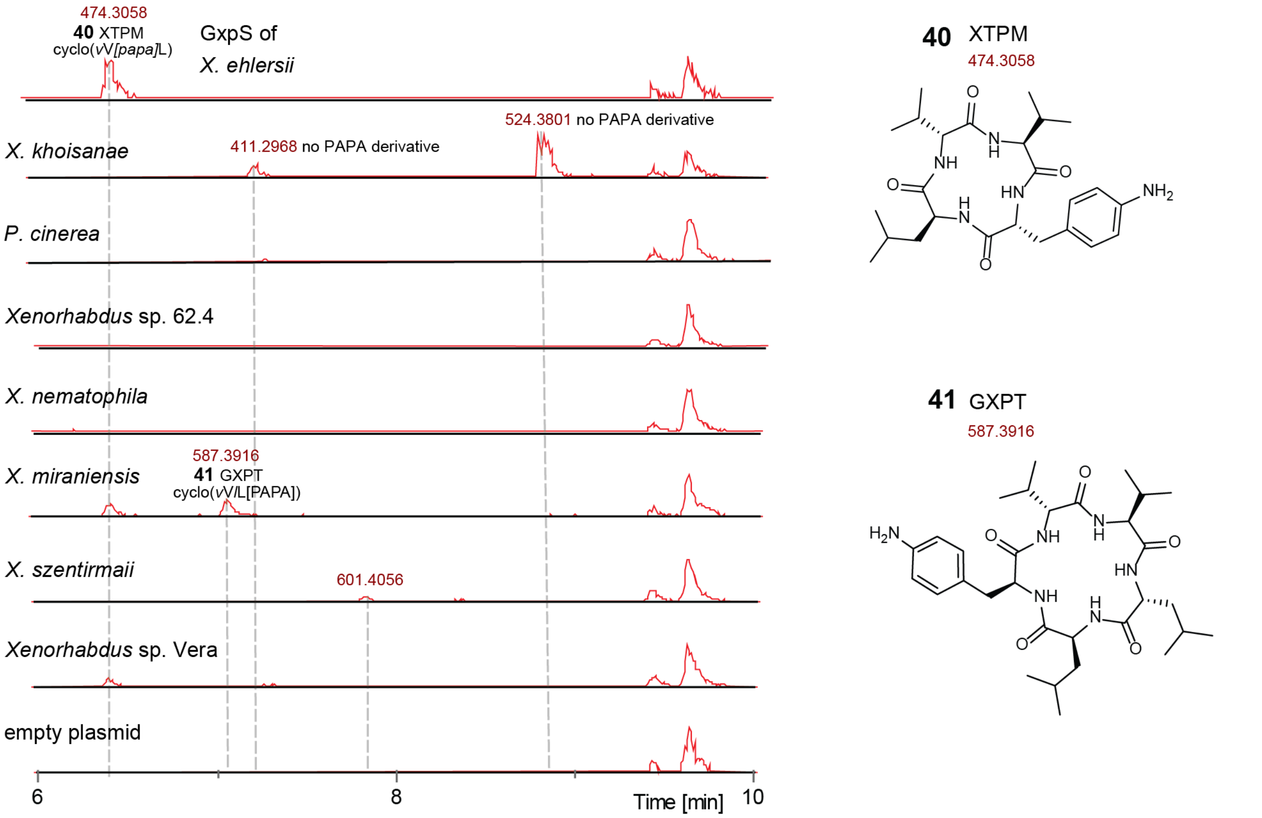


**Figure S48. EIC of all main peaks from different GxpS that are generally not capable of accepting PAPA.** However, two GxpS were still able to produce two GXP (**40**, **41**) with PAPA in small amounts. Shown are the ionized high-resolution mass and the corresponding peptide. *E. coli* carrying different *gxpS* orthologs and pCK0403 were cultured in XPPM supplemented with 2 mM PAPA. Natural product production was measured by HPLC-HRMS. The *E. coli* that carried the *gxpS* of *X. miraniensis* was co-transformed with pCK0413.

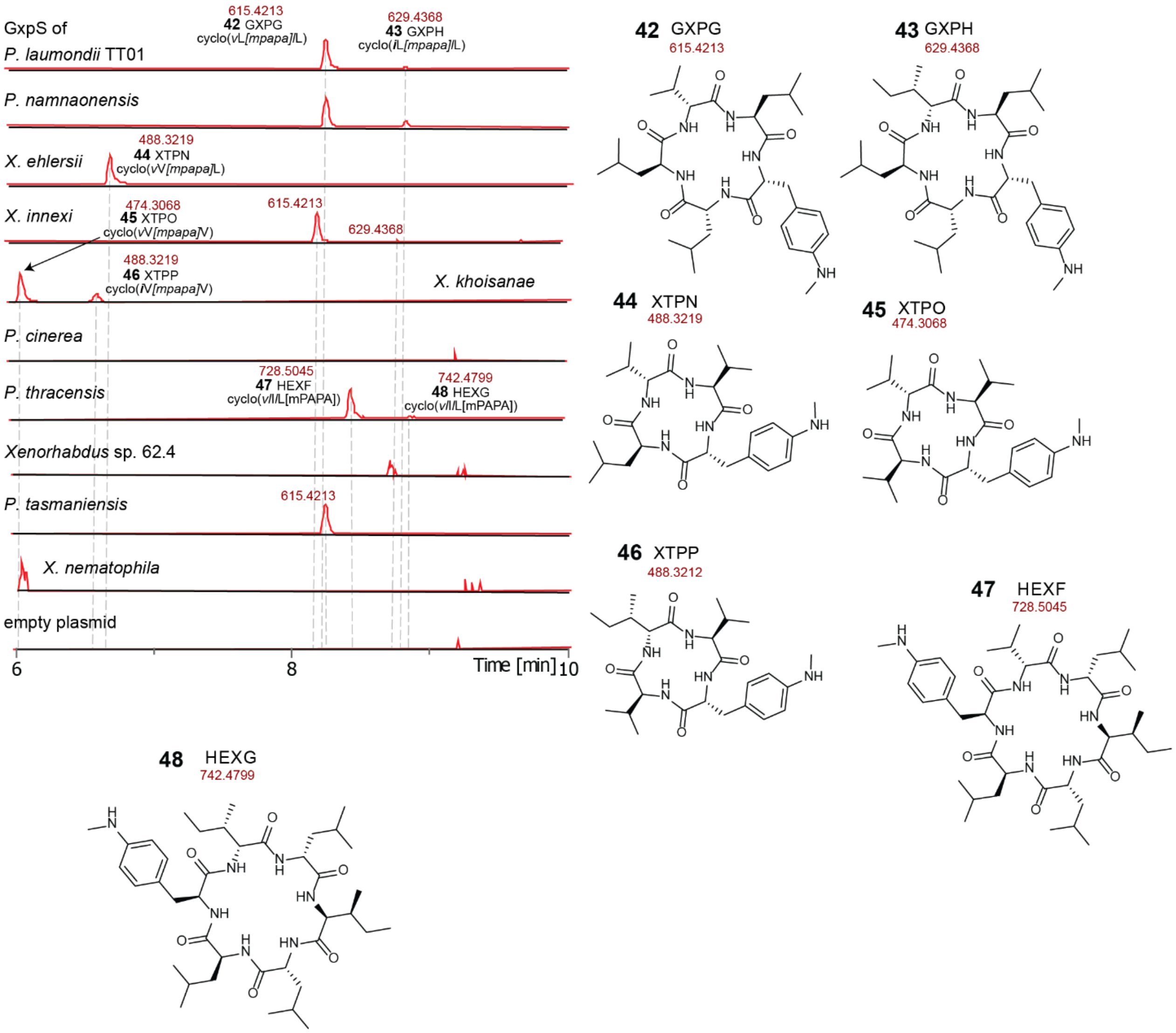


**Figure S49. EIC of all main peaks from different GxpS that are capable of accepting mPAPA.** Shown are the ionized high-resolution mass and the corresponding peptide. *E. colis* carrying different *gxpS* orthologs and pCK0403 were cultured in XPPM supplemented with 2 mM mPAPA. Natural product production was measured by HPLC-HRMS.

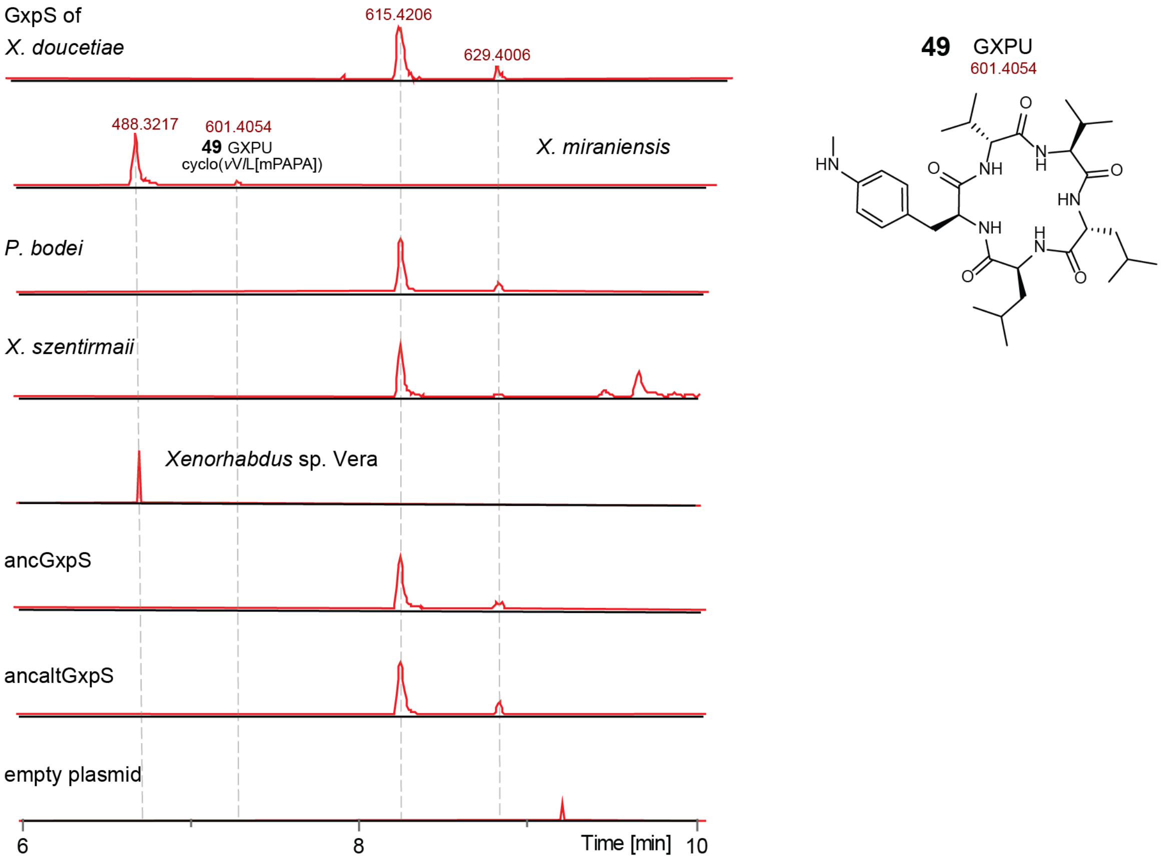


**Figure S50. EIC of all main peaks from different GxpS that are not capable of accepting mPAPA.** Shown is the ionized high-resolution mass and the corresponding peptide. *E. coli* carrying different *gxpS* orthologs and pCK0403 were cultured in XPPM supplemented with 2 mM synthetic mPAPA. Natural product production was measured by HPLC-HRMS. Shown is the mPAPA pentapeptide derivative produced by GxpS from *X. miraniensis*. The *E. coli* that carried the *gxpS* of *X. miraniensis* was co-transformed with pCK0413.

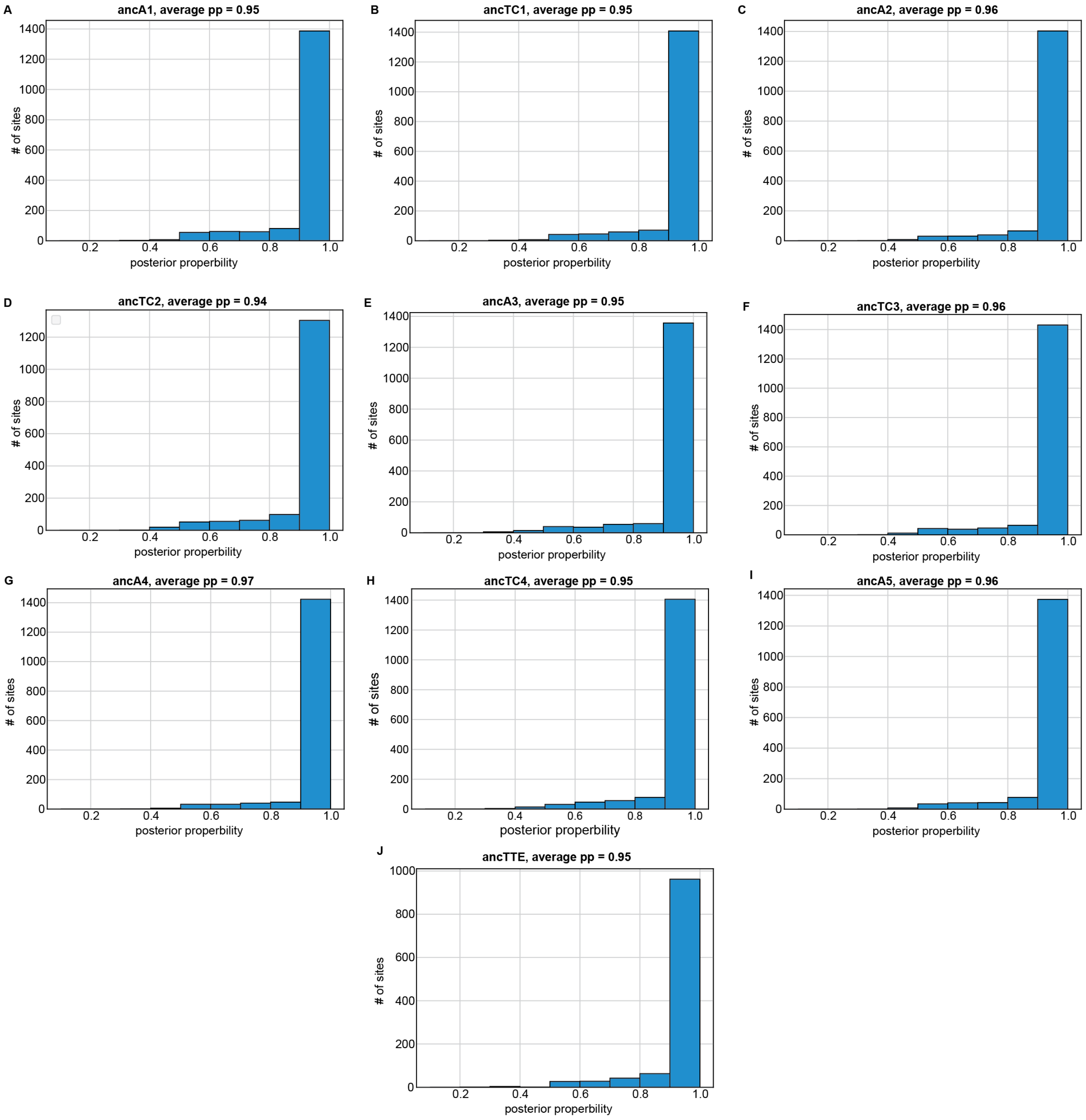


**Figure S51. Histograms of the posterior probability of the most probable state at each site for all individual domains of AncGxpS.**

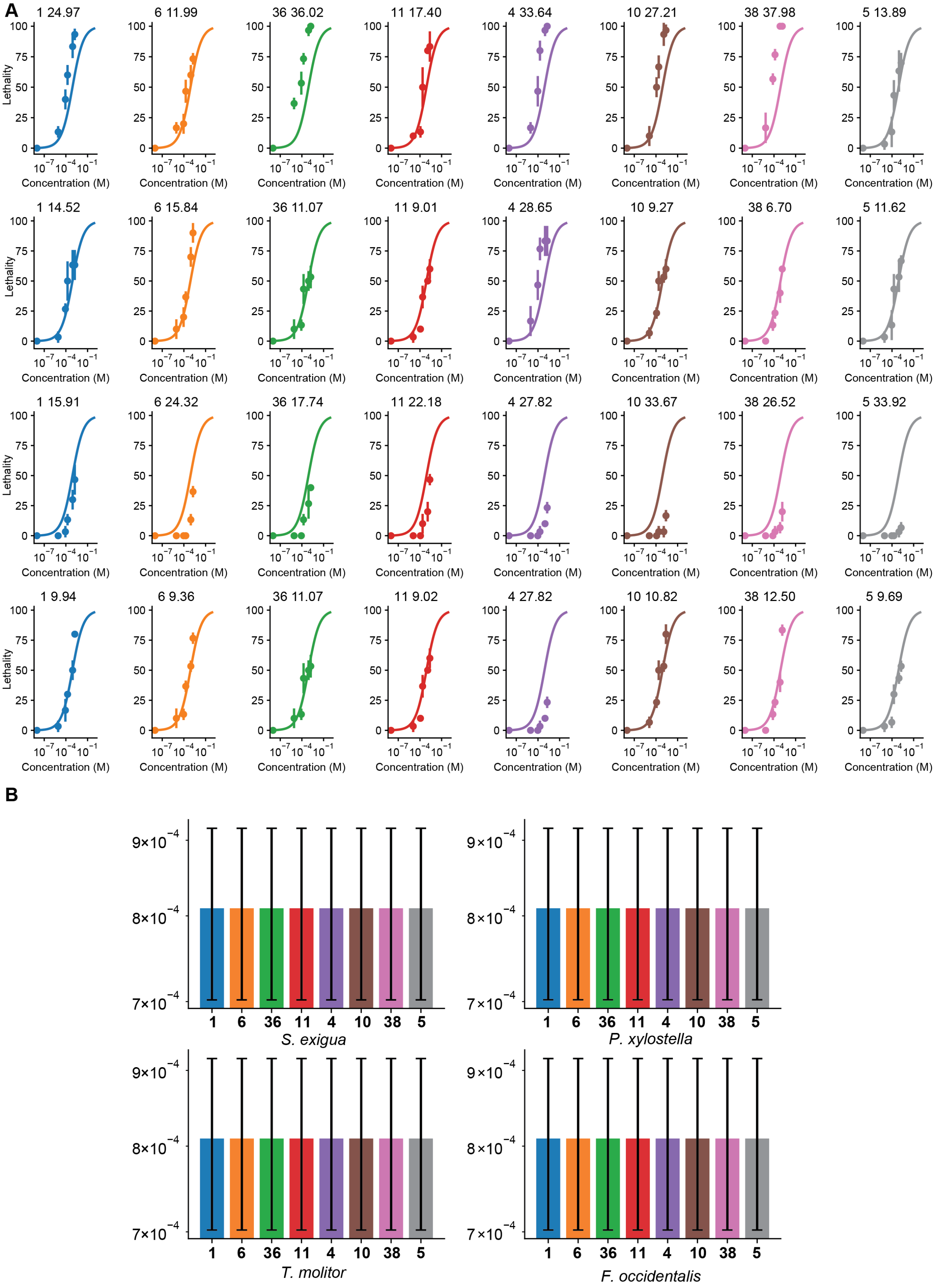
A
1 24.97
6 11.99
36 36.02
11 17.40
4 33.64
10 27.21
38 37.98
5 13.89
1 14.52
6 15.84
36 11.07
11 9.01
4 28.65
10 9.27
38 6.70
5 11.62
1 15.91
6 24.32
36 17.74
11 22.18
4 27.82
10 33.67
38 26.52
5 33.92
1 9.94
6 9.36
36 11.07
11 9.02
4 27.82
10 10.82
38 12.50
5 9.69
Lethality
Concentration (M)
B
S. exigua
P. xylostella
T. molitor
F. occidentalis

**Figure S52. Bioactivity data simulated by model 1.** (**A**) We fitted the GXP-dose-dependent insect lethality to a sigmoid function. We used one sigmoid function for all data, assuming all peptides have the same efficacy across all insects. (**B**) Corresponding $LC_{50}$ values were plotted on bar graphs.

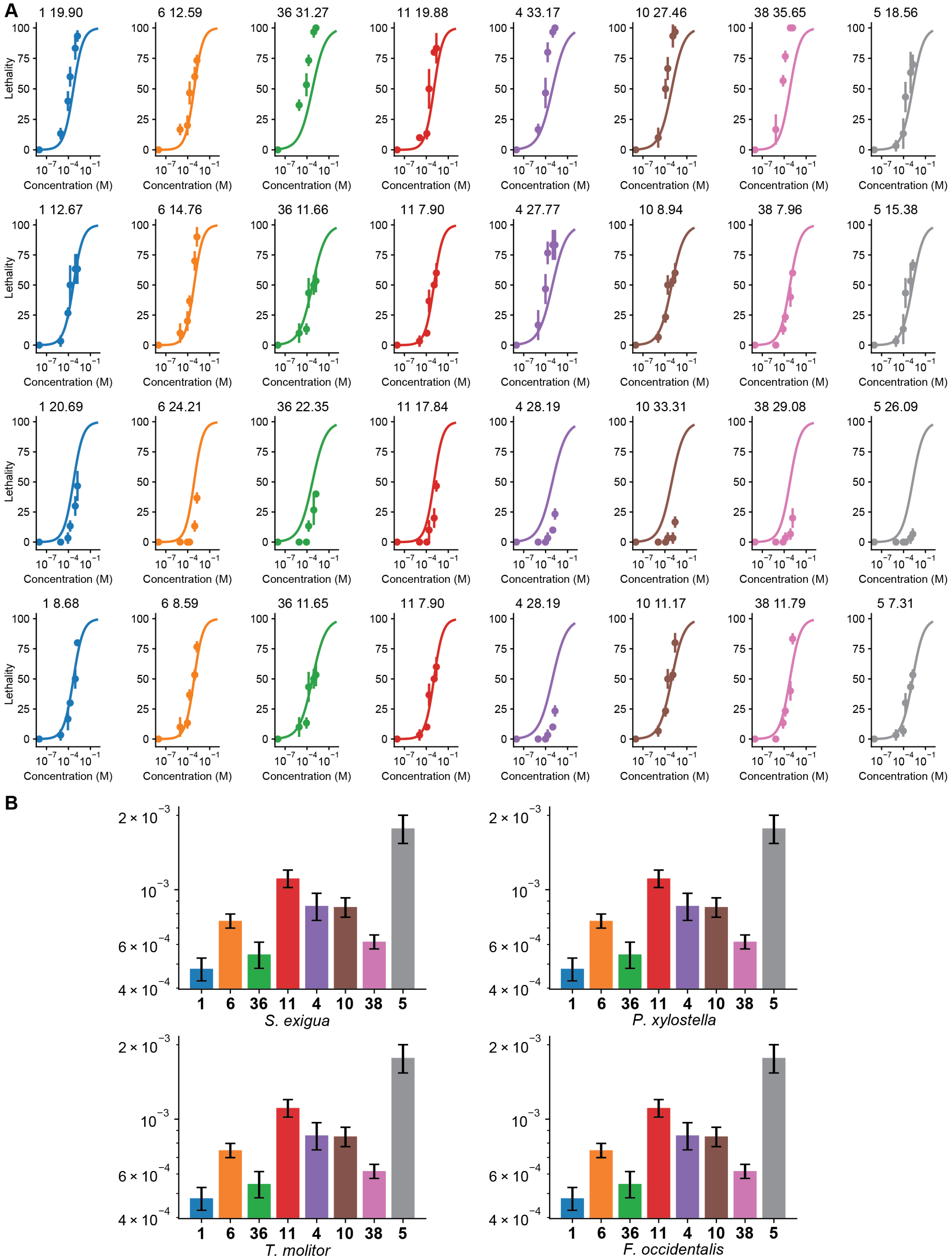
A
1 19.90
6 12.59
36 31.27
11 19.88
4 33.17
10 27.46
38 35.65
5 18.56
1 12.67
6 14.76
36 11.66
11 7.90
4 27.77
10 8.94
38 7.96
5 15.38
1 20.69
6 24.21
36 22.35
11 17.84
4 28.19
10 33.31
38 29.08
5 26.09
1 8.68
6 8.59
36 11.65
11 7.90
4 28.19
10 11.17
38 11.79
5 7.31
Lethality
Concentration (M)
B
1 6 36 11 4 10 38 5
S. exigua
P. xylostella
T. molitor
F. occidentalis

**Figure S53. Bioactivity data simulated by model 2.** (**A**) We fitted the GXP-dose-dependent insect lethality to sigmoid functions. We used one sigmoid function for each peptide, assuming the same efficacy across all insects. (**B**) Corresponding $LC_{50}$ values were plotted on bar graphs.

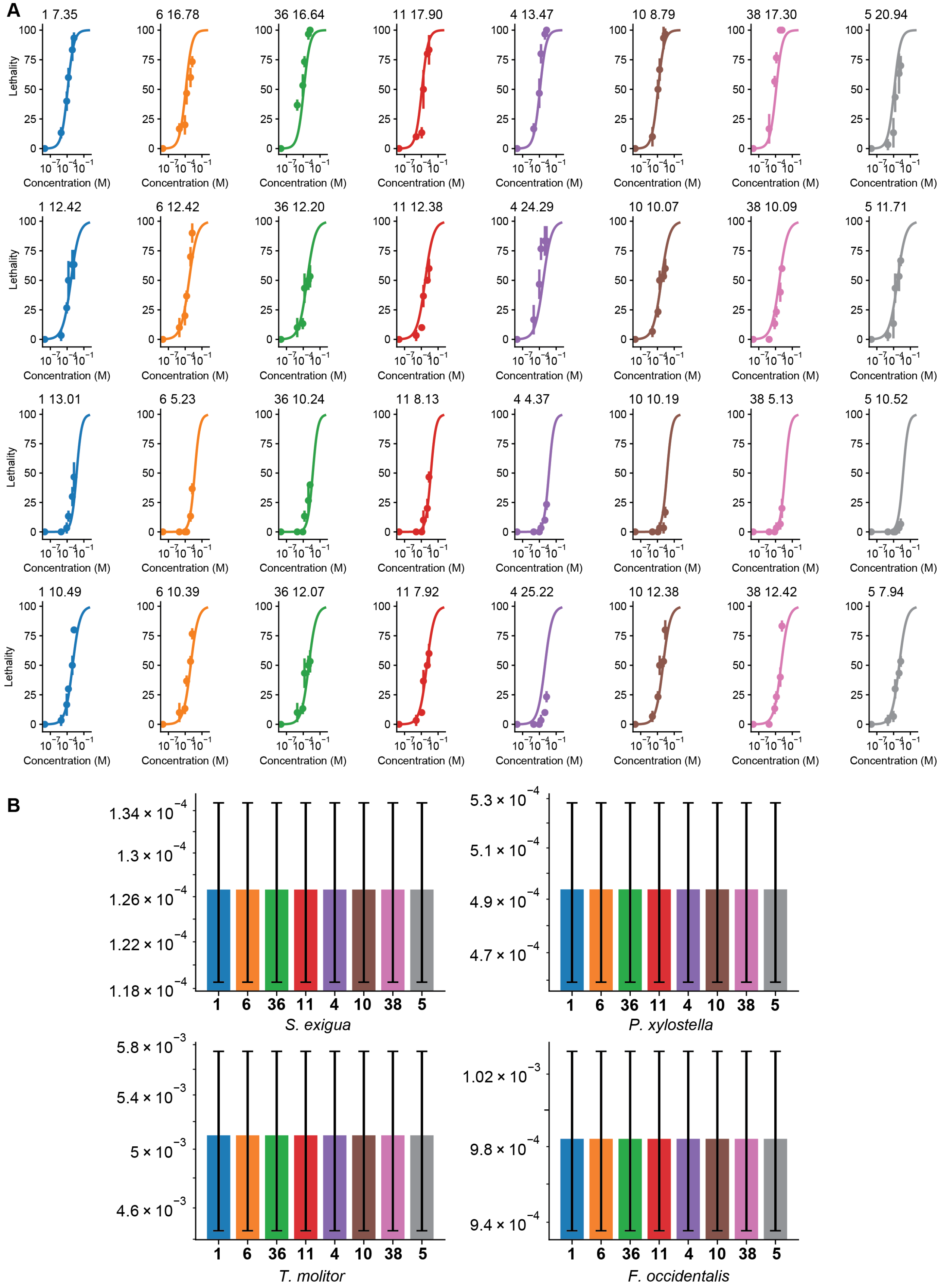
A
1 7.35
6 16.78
36 16.64
11 17.90
4 13.47
10 8.79
38 17.30
5 20.94
1 12.42
6 12.42
36 12.20
11 12.38
4 24.29
10 10.07
38 10.09
5 11.71
1 13.01
6 5.23
36 10.24
11 8.13
4 4.37
10 10.19
38 5.13
5 10.52
1 10.49
6 10.39
36 12.07
11 7.92
4 25.22
10 12.38
38 12.42
5 7.94
Lethality
Concentration (M)
B
S. exigua
P. xylostella
T. molitor
F. occidentalis
1
6
36
11
4
10
38
5

**Figure S54. Bioactivity data simulated by model 3.** (**A**) We fitted the GXP-dose-dependent insect lethality to sigmoid functions. We used one sigmoid function for each insect, assuming the same efficacy across all peptides. (**B**) Corresponding $LC_{50}$ values were plotted on bar graphs.

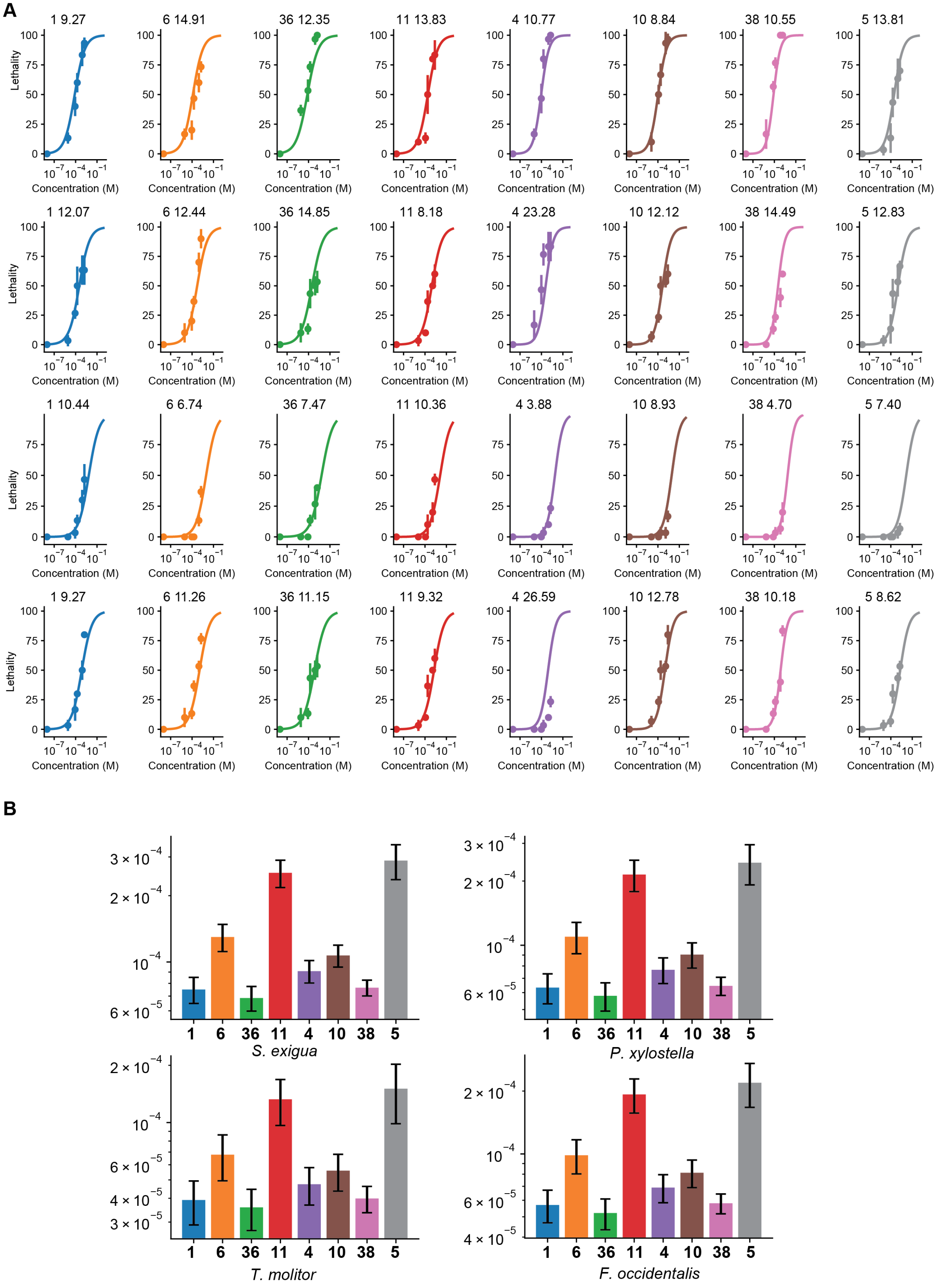
A
1 9.27
6 14.91
36 12.35
11 13.83
4 10.77
10 8.84
38 10.55
5 13.81
1 12.07
6 12.44
36 14.85
11 8.18
4 23.28
10 12.12
38 14.49
5 12.83
1 10.44
6 6.74
36 7.47
11 10.36
4 3.88
10 8.93
38 4.70
5 7.40
1 9.27
6 11.26
36 11.15
11 9.32
4 26.59
10 12.78
38 10.18
5 8.62
Lethality
Concentration (M)
B
S. exigua
P. xylostella
T. molitor
F. occidentalis
1 6 36 11 4 10 38 5

**Figure S55. Bioactivity data simulated by model 4.** (**A**) We fitted the GXP-dose-dependent insect lethality to sigmoid functions. We used one sigmoid function for each peptide and allowed one scaling factor per species. (**B**) Corresponding $LC_{50}$ values were plotted on bar graphs.

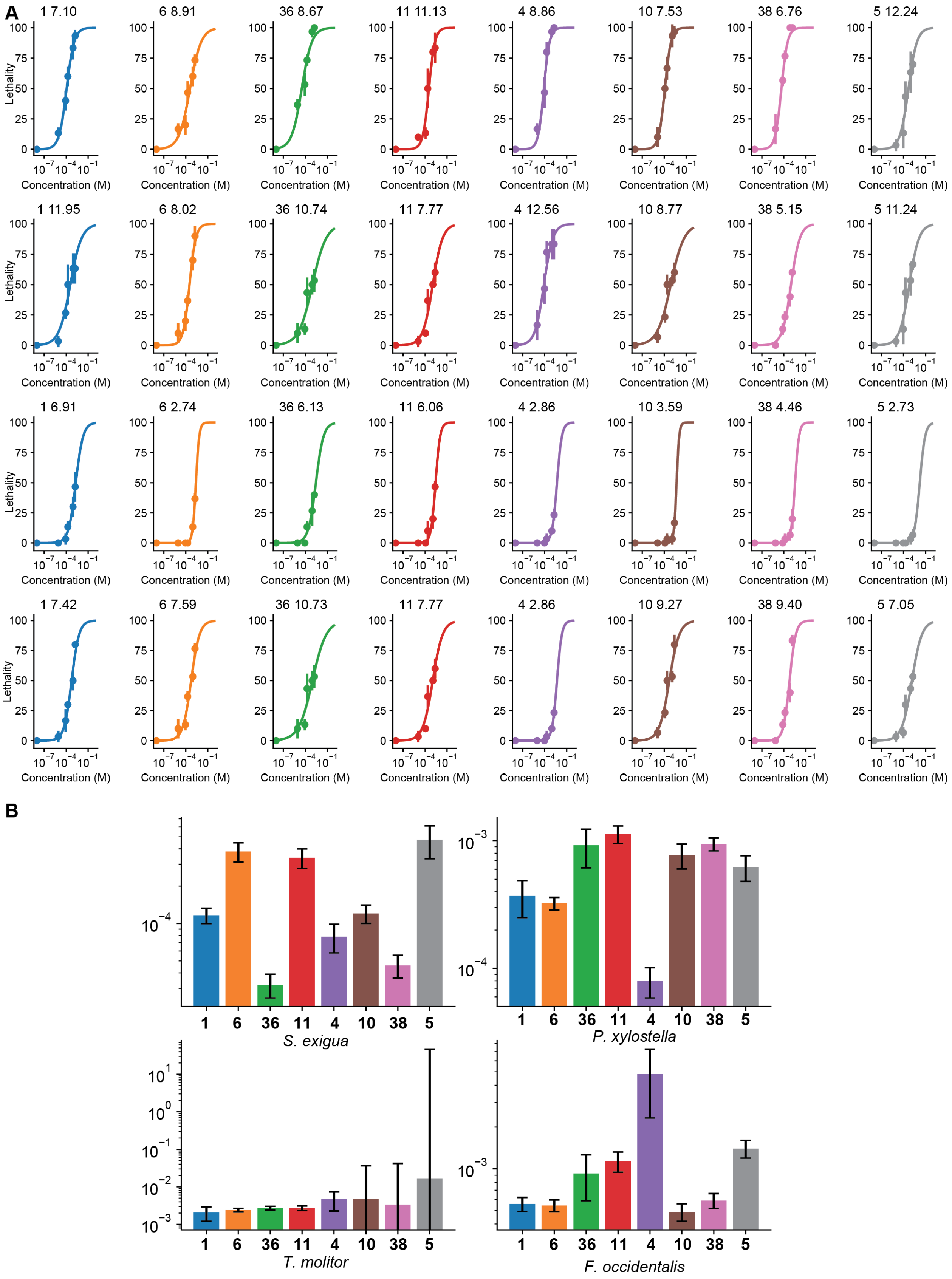
A
1 7.10
6 8.91
36 8.67
11 11.13
4 8.86
10 7.53
38 6.76
5 12.24
1 11.95
6 8.02
36 10.74
11 7.77
4 12.56
10 8.77
38 5.15
5 11.24
1 6.91
6 2.74
36 6.13
11 6.06
4 2.86
10 3.59
38 4.46
5 2.73
1 7.42
6 7.59
36 10.73
11 7.77
4 2.86
10 9.27
38 9.40
5 7.05
Lethality
Concentration (M)
B
S. exigua
P. xylostella
T. molitor
F. occidentalis

**Figure S56. Bioactivity data simulated by model 5.** (**A**) We fitted the GXP-dose-dependent insect lethality to sigmoid functions. We used one sigmoid function for each peptide/species combination. (**B**) Corresponding $LC_{50}$ values were plotted on bar graphs.

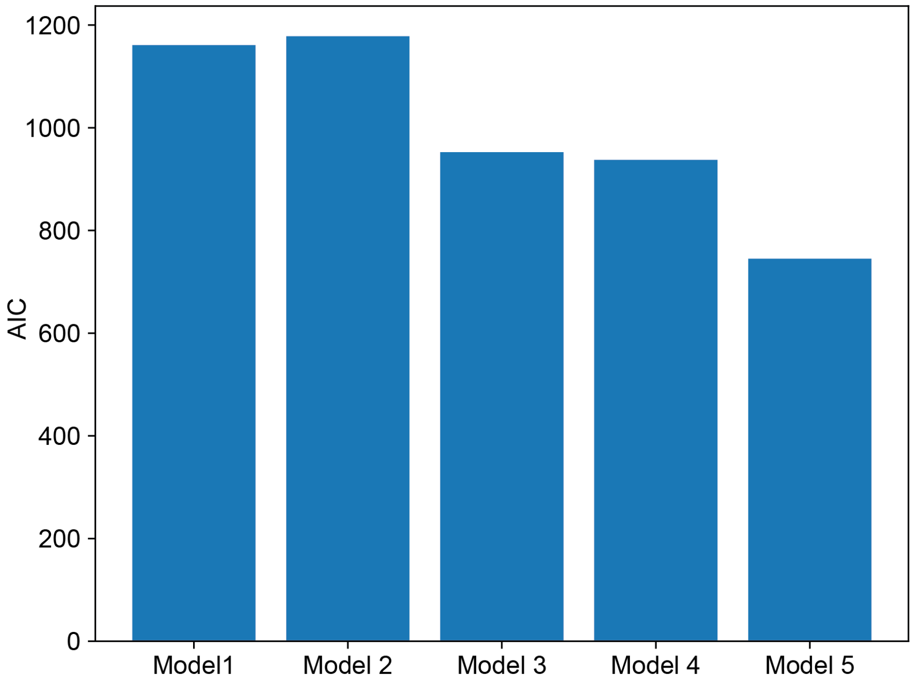


**Figure S57. AIC values for every model of our bioactivity data.**

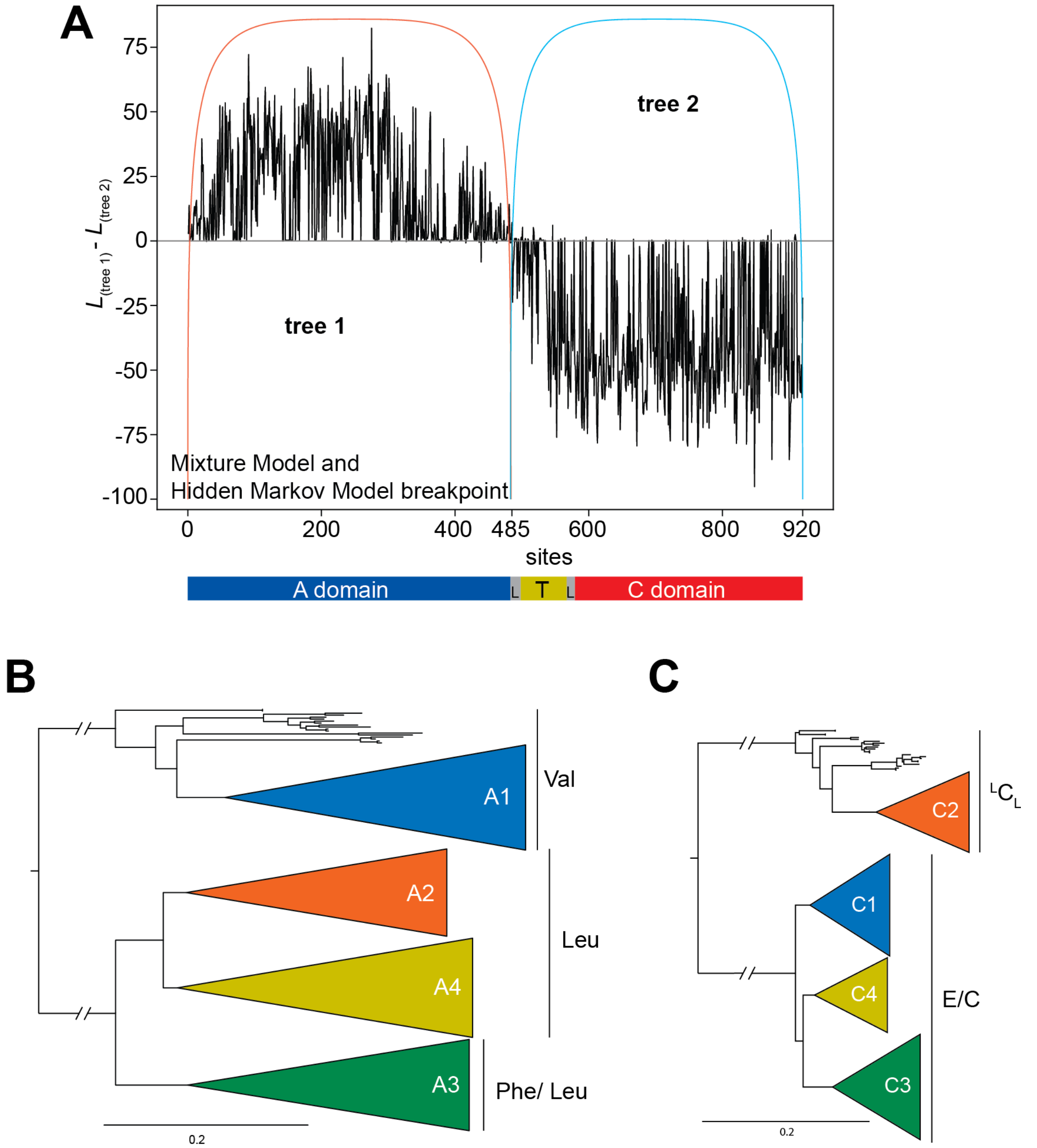


**Figure S58. Hidden Markov Model (HMM) analysis of ATC tridomain protein alignment of GxpS to determine the phylogenetic border of the A and C-domain.** (**A**) The difference of the site-wise log-likelihoods of two phylogenetic trees of ATC tri-domains was plotted that together best describe the alignment using a phylogenetic hidden Markov model. Sites with positive numbers are better described by tree1, sites with negative numbers are better described by tree 2. The Viterbi and Forward-Backward algorithm partition the alignment at the end of the A-domain at site 485. Using a Mixture Model (MM) analysis to partition the alignment also pointed to site 485 as the breakpoint. (**B**) First tree generated from the HMM analysis. The first tree resembles the A-domain tree. The tree falls into clades according to the specificity. There are clades containing A-domains specific for valine, leucine and phenylalanine. (**C**) Second generated tree from the HMM analysis. The second resembles the C-domain tree. The tree falls according to the C-domain type. On clade contains all $^{L}C_{L}$ domains and the other one all dual C/E-domains.

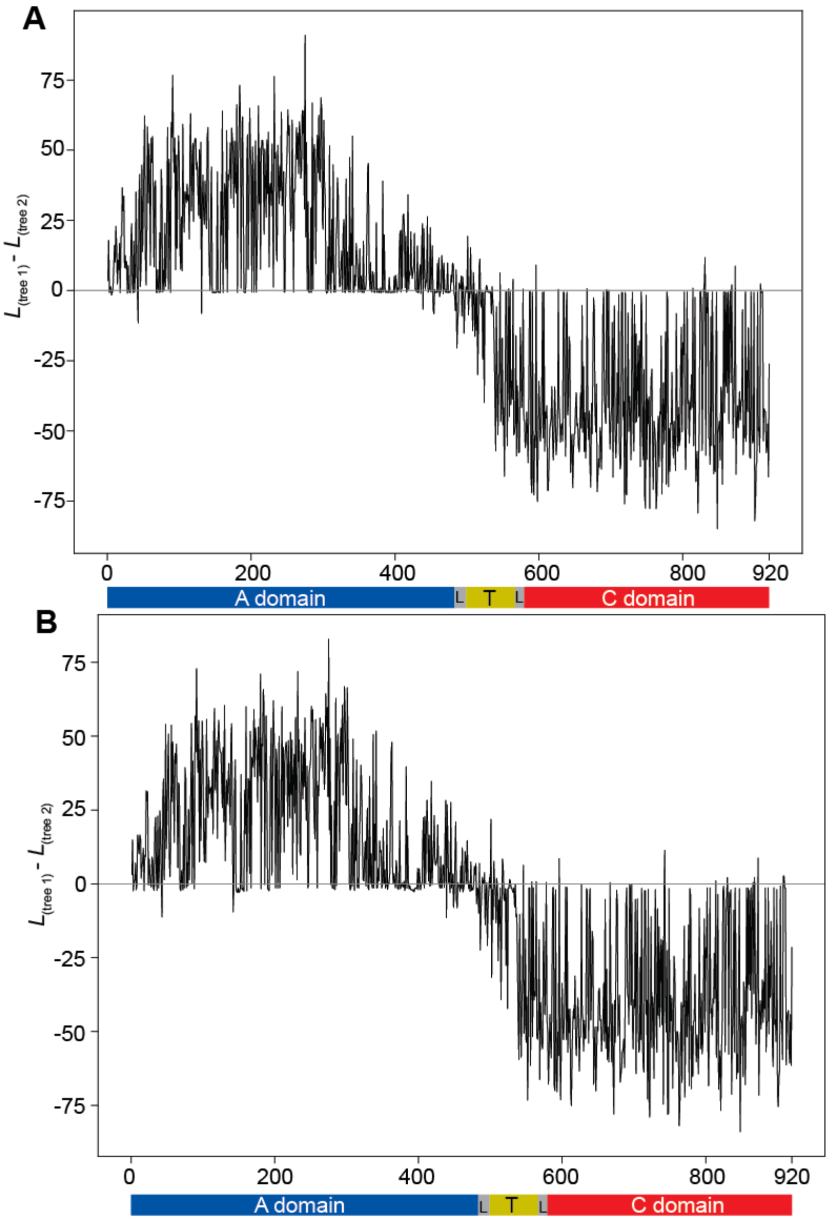


**Figure S59**. **MAST analysis of the ATC tridomain of GxpS to determine the phylogenetic border of the A and C-domain.** As input, the two trees of the HMM and MM analysis were taken. **(A)** The difference of the log likelihood of these two trees along the ATC tri-domain alignment were plotted. Tree 1 is better described by positive numbers and tree 2 is better described by negative numbers. (**B**) MAST analysis of the ATC-tridomain of GxpS to determine the phylogenetic border of the A and C-domain. As input, a separate inferred A and C-domain tree were used. The difference of the site-wise log-likelihood of these two trees along the ATC tri-domain alignment were plotted. Sites with positive numbers are better described by tree 1, sites with negative numbers are better described by tree 2.

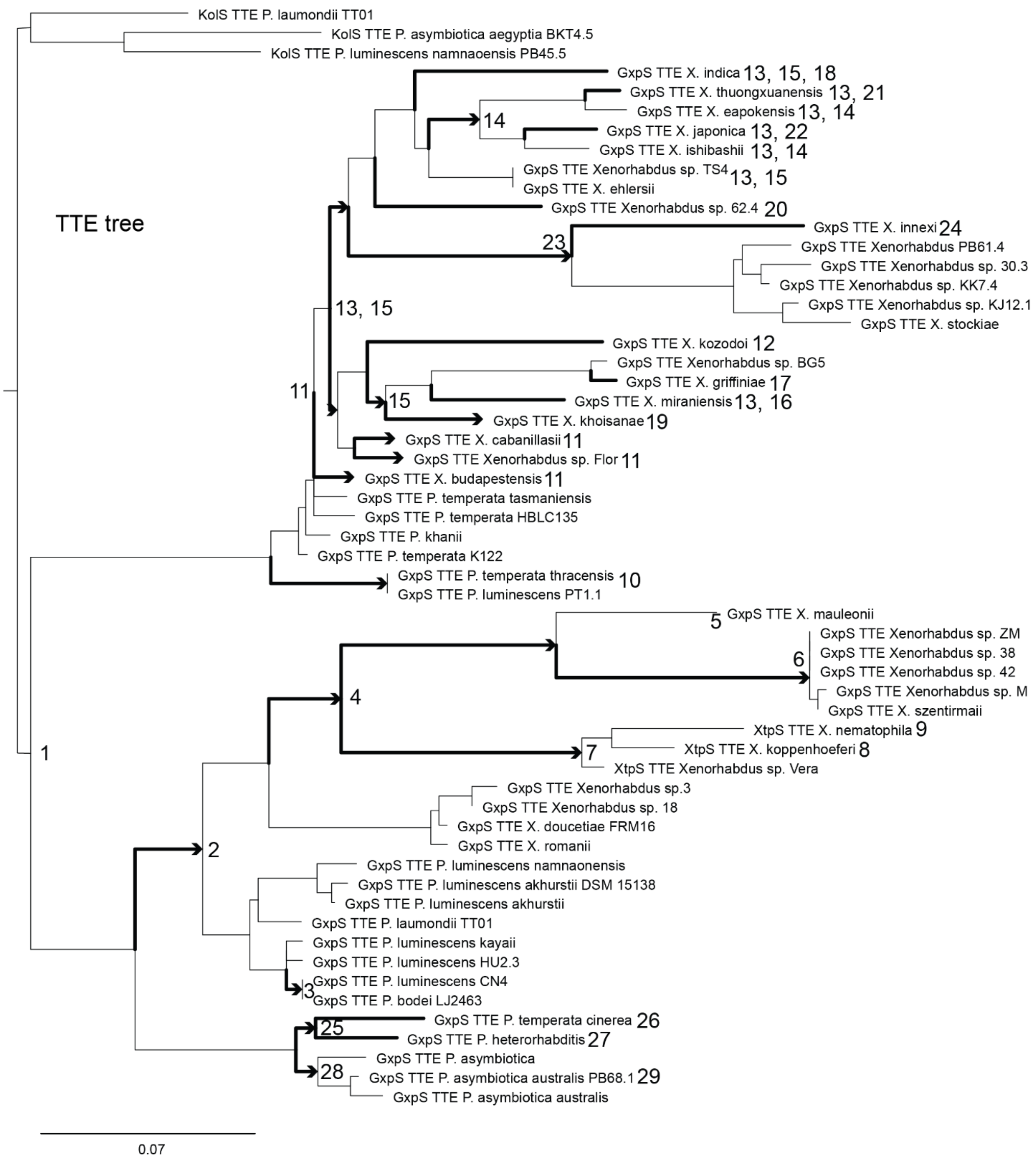


**Figure S60. TTE-domain tree with numbered nodes. Every numbered node corresponds to a GxpS that received at least one recombination.** The branches in arrow shape indicate the time point where a specific recombination occurred.

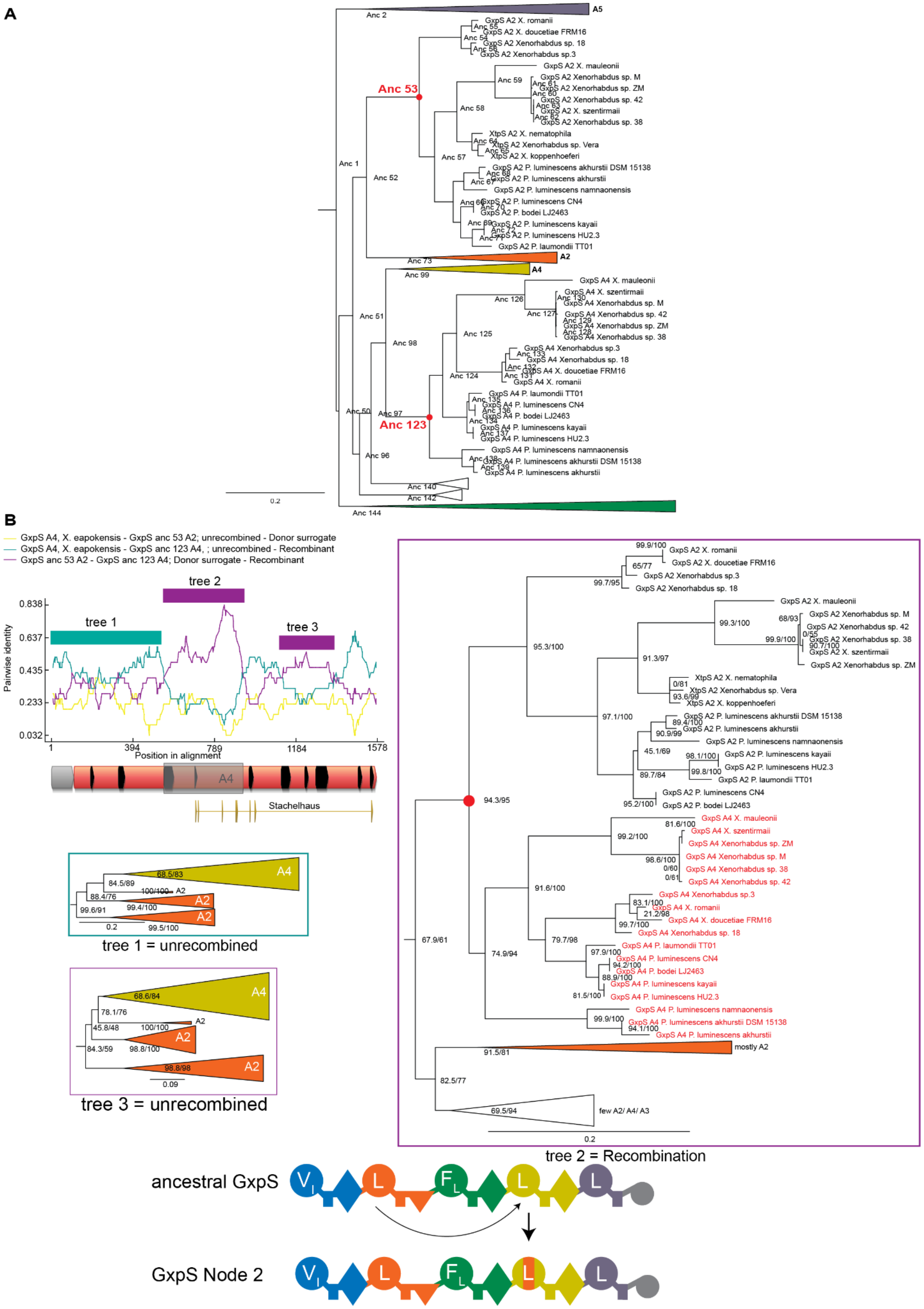
A
A5
A2
A4
Anc 2
Anc 1
Anc 52
Anc 53
Anc 57
Anc 58
Anc 59
Anc 73
Anc 99
Anc 51
Anc 98
Anc 125
Anc 126
Anc 124
Anc 97
Anc 123
Anc 50
Anc 96
Anc 140
Anc 142
Anc 144
GxpS A2 X. romanii
GxpS A2 X. doucetiae FRM16
GxpS A2 Xenorhabdus sp. 18
GxpS A2 Xenorhabdus sp.3
GxpS A2 X. mauleonii
GxpS A2 Xenorhabdus sp. M
GxpS A2 Xenorhabdus sp. ZM
GxpS A2 Xenorhabdus sp. 42
GxpS A2 X. szentirmaii
GxpS A2 Xenorhabdus sp. 38
XtpS A2 X. nematophila
XtpS A2 Xenorhabdus sp. Vera
XtpS A2 X. koppenhoeferi
GxpS A2 P. luminescens akhurstii DSM 15138
GxpS A2 P. luminescens akhurstii
GxpS A2 P. luminescens namnaonensis
GxpS A2 P. luminescens CN4
GxpS A2 P. bodei LJ2463
GxpS A2 P. luminescens kayaii
GxpS A2 P. luminescens HU2.3
GxpS A2 P. laumondii TT01
GxpS A4 X. mauleonii
GxpS A4 X. szentirmaii
GxpS A4 Xenorhabdus sp. M
GxpS A4 Xenorhabdus sp. 42
GxpS A4 Xenorhabdus sp. ZM
GxpS A4 Xenorhabdus sp. 38
GxpS A4 Xenorhabdus sp.3
GxpS A4 Xenorhabdus sp. 18
GxpS A4 X. doucetiae FRM16
GxpS A4 X. romanii
GxpS A4 P. laumondii TT01
GxpS A4 P. luminescens CN4
GxpS A4 P. bodei LJ2463
GxpS A4 P. luminescens kayaii
GxpS A4 P. luminescens HU2.3
GxpS A4 P. luminescens namnaonensis
GxpS A4 P. luminescens akhurstii DSM 15138
GxpS A4 P. luminescens akhurstii
0.2
B
GxpS A4, X. eapokensis - GxpS anc 53 A2; unrecombined - Donor surrogate
GxpS A4, X. eapokensis - GxpS anc 123 A4, ; unrecombined - Recombinant
GxpS anc 53 A2 - GxpS anc 123 A4; Donor surrogate - Recombinant
tree 1
tree 2
tree 3
Pairwise identity
0.838
0.637
0.435
0.233
0.032
1
394
789
1184
1578
Position in alignment
A4
Stachelhaus
tree 1 = unrecombined
tree 3 = unrecombined
tree 2 = Recombination
mostly A2
few A2/ A4/ A3
ancestral GxpS
GxpS Node 2

**Figure S61. Recombination analysis of GxpS at Node 2 on the TTE-domain tree.** (**A**) A trimmed version of the A-domain phylogeny used for ASR. The node 53 and 123 are annotated with red dots. These nodes were used in the subsequent RDP analysis to detect the recombination. (**B**) This recombination was discovered through phylogenetic trees of the middle part of the A-domain, in which some A4 domains always claded inside the A2 clade. Top left: RDP sequence similarity analysis. In this analysis our alignment contained two ancestral A-domains, which should have existed close to the presumed recombination event. This was necessary, because the recombination happened very deep in the GxpS phylogeny such that extant A-domains were not similar enough anymore to the presumed donor and acceptor sequences for RDP to identify them. Recombined refers to an A-domain that carries the recombination (here the ancestral A4 that carries this recombination). Donor surrogate refers to an ancestral A2 domain that is similar to the recombined portion. Unrecombined refers to A-domains that have not experienced this recombination (here the A4 domain domain from *Xeonorhabdus eapokensis*). Pairwise sequence identity refers to the identity across variable sites with a 30 nucleotide sliding window. The cyan line compares the recombined A-domain to its unrecombined relatives. The purple line indicates where recombination has made the recombined A-domain similar to other A-domains. The yellow line serves as a control. Domain architecture of the flanking linker domain (gray) and A-domain (red) are shown under the plot. Black arrows indicate structural motifs in the A-domain [13], orange arrows below denote specificity encoding sites according to Stachelhaus *et* al [14]. A grey box is placed over the recombined area. Tree 1 and Tree 3 show A2 and A4 domain sequences are mostly monophyletic and thus show no recombination. Tree 2 shows the deep recombination event at a deep node (shown as a red dot). All descendant A2 domains show a mirrored subtree with A4 domains that hints recombination.

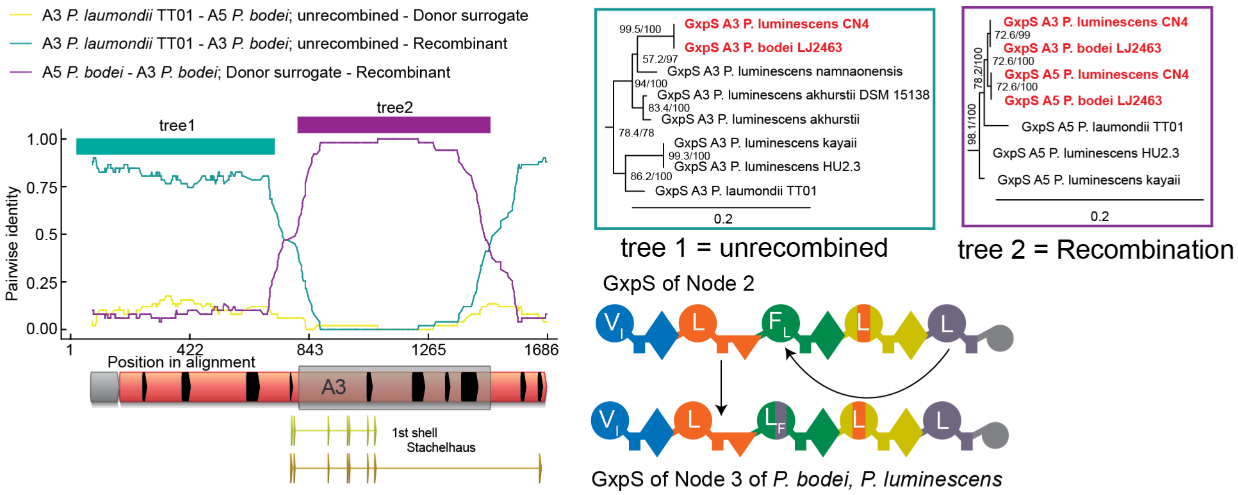


**Figure S62. Recombination analysis of GxpS at Node 3 on the TTE-domain tree.** This recombination was identified through the incongruent tree topology in the A3 clade (Fig. 2d). Left: RDP analysis of recombinant A-domains. Recombined refers to an A-domain (here the A3 domain from *P. bodei*) that carries this recombination. Donor surrogate refers to an extant A domais that is similar to the recombined portion (here the A5 domain from *P. bodei*). Unrecombined refers to A-domains that have not experienced this recombination (here the A5 domain from A3 domain from *P.laumondii*). Pairwise sequence identity refers to the identity across variable sites with a 30 nt sliding window. The cyan line compares the recombined A-domain to its unrecombined relatives. The purple comparison indicates where recombination has made it similar to other A-domains. The yellow line serves as a control. Domain architecture of the flanking linker domain (grey) and A-domain (red) are shown under the plot. Black arrows indicate structural motifs in the A-domain [13], orange arrows below denote specificity encoding sites according to Stachelhaus *et al* [14]. Yellow arrows are the first shell and are 8 residues of the Stachelhaus code. The first D and last K have been excluded in the first shell. A grey box is placed over the recombined area. Middle and right: Phylogenetic trees inferred from amino acid sequences indicated with cyan and purple bars in the RDP plot. In the first tree, the two A-domains are labelled in red clade with other A5 domains. In the second tree, they clade inside A5 domains because of recombination. Bottom right: Schematic representation of this recombination event.

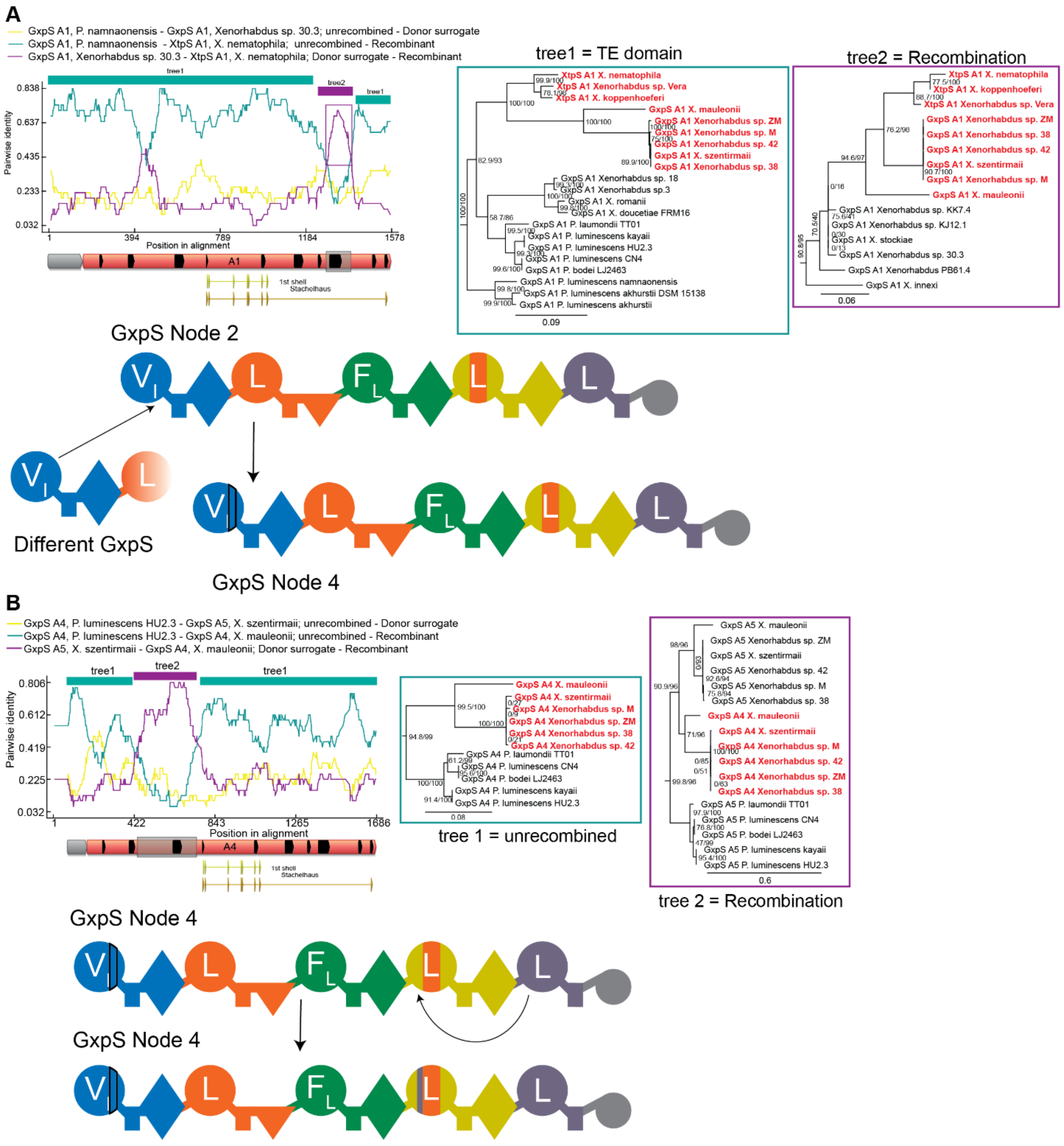


**Figure S63. Two recombinations at Node 4 on the TTE-domain.** These recombinations were discovered using RDP. (**A**) The first recombination that occurred at node 4. RDP5 analysis as in Supplementary Fig. 62, trees inferred from sites indicated on RDP plot were inferred from nucleotides. A small part of the first A-domain was overwritten by an A1 domain from another GxpS. (**B**) Second recombination that occurred at Node 4, with RDP and tree analysis as above. The nucleotide tree of the putative recombined fragment shows a repeated subtree of A4 and A5 domains, indicative of recombination. Recombination occurred between homotypic C-domains and outside the specificity area and was synonymous.

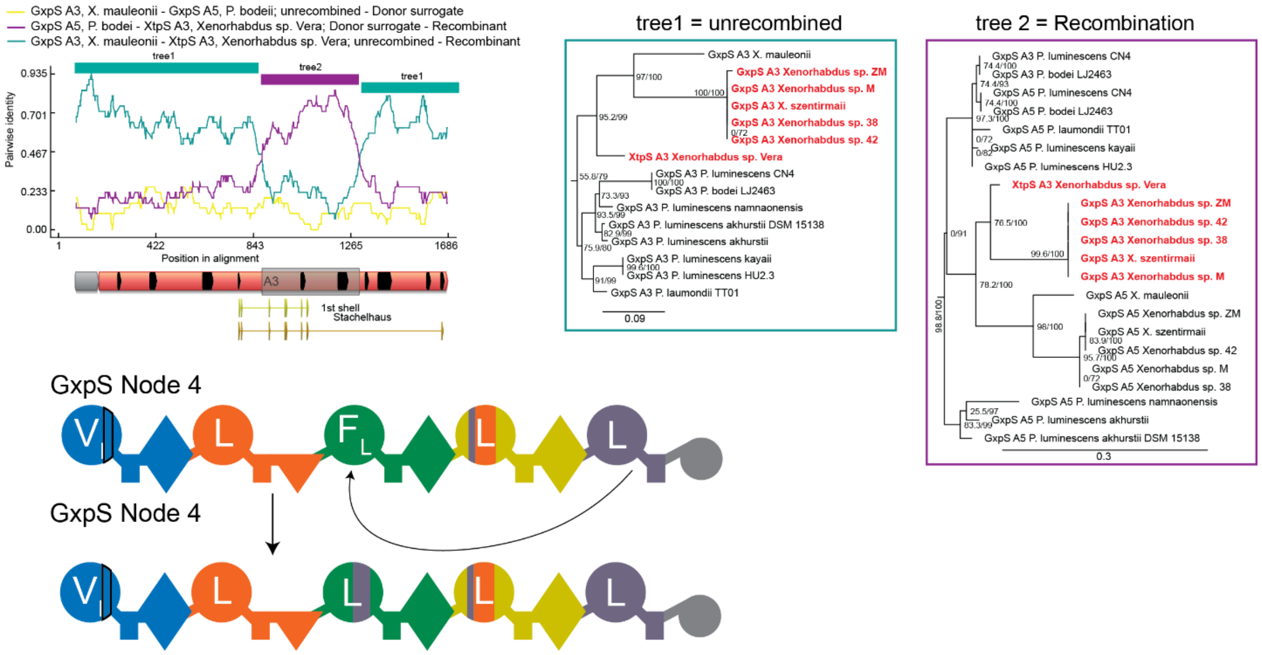

**Figure S64. Third recombination analysis of GxpS at Node 4 on the TTE-domain tree.** This recombination was identified through the incongruent tree topology on the A3 clade (Fig. 2d). RDP and tree analysis as described in Supplementary Fig. 62, trees were inferred from amino acids. The repeated subtree in the second tree suggests GxpS A5 domain has non-synonymously recombined into its own A3 domain and thereby changed the specificity from phenylalanine to leucine.

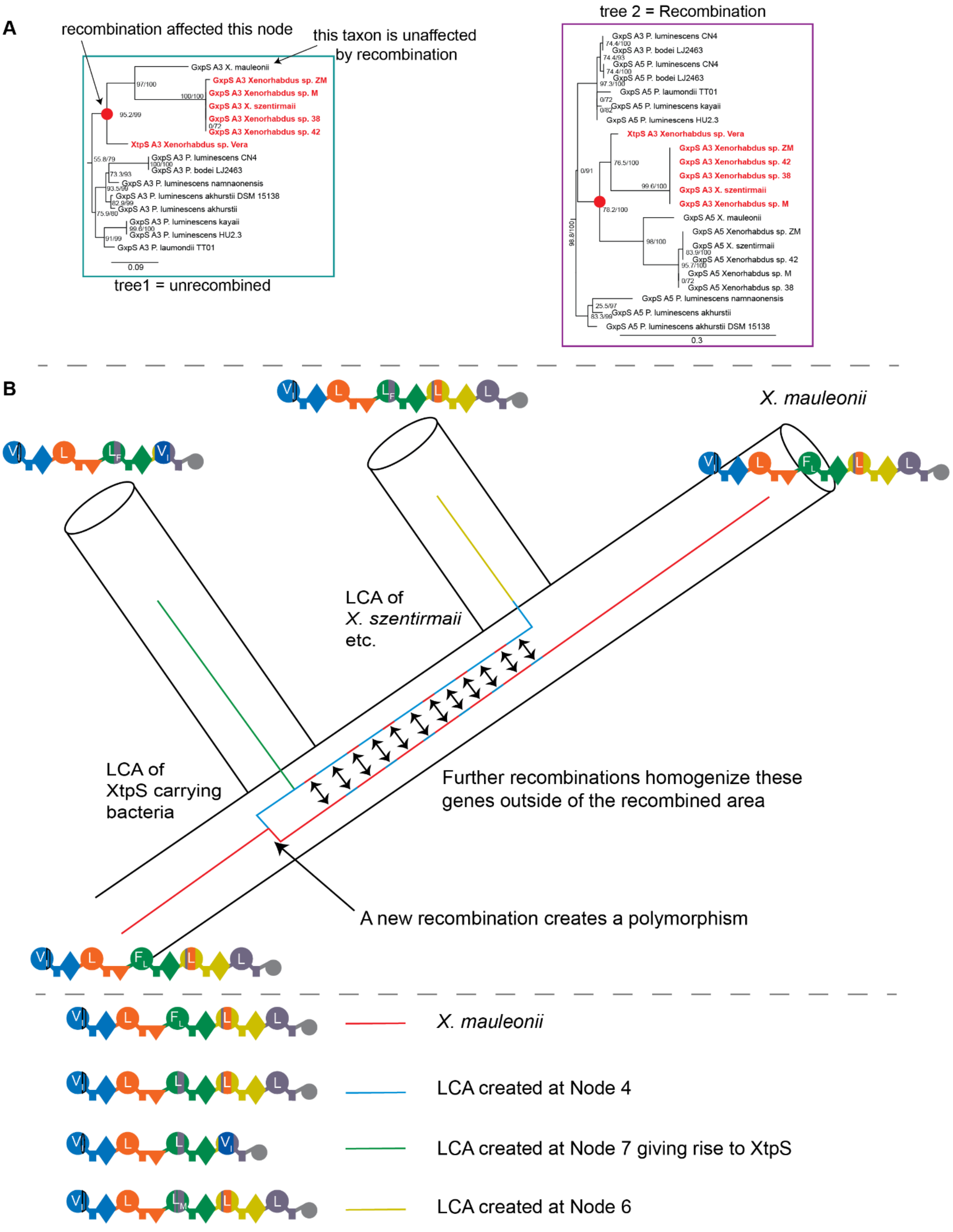


**Figure S65. Incomplete lineage sorting affects NRPS diversity.** (**A**) The unrecombined tree Supplementary Fig. 64, that aligns with the TTE-domain tree, shows the time point of recombination. It shows that all descendants, beside of *X. maulenoii*, were affected by this recombination event.

Tree 2 that shows the recombination, showed which species were affected by this recombination and that *X. mauleonii* is missing. (**B**) A diagram to explain the process of incomplete lineage sorting. The underlying species phylogeny is shown as a series of connected tubes, in which alleles evolve. A recombination created polymorphism in the population (splitting of reed line into blue and red lineages). This polymorphism persists past several speciation events, allowing *X. maulenoii* to inherit the unrecombined allele (red line). The polymorphism also exists in the LCA of XtpS (green line) and in the LCA of species similar to *X. szentirmaii* (yellow line) that carry the recombined A3 domain. To explain why the unrecombined parts of the synthetase still follow the species tree topology, we have to propose that the two alleles exchanged genetic material outside the recombined area and thereby homogenized (red and blue line), s This explains why *X. mauleonii* still carries the unrecombined A3 domain.

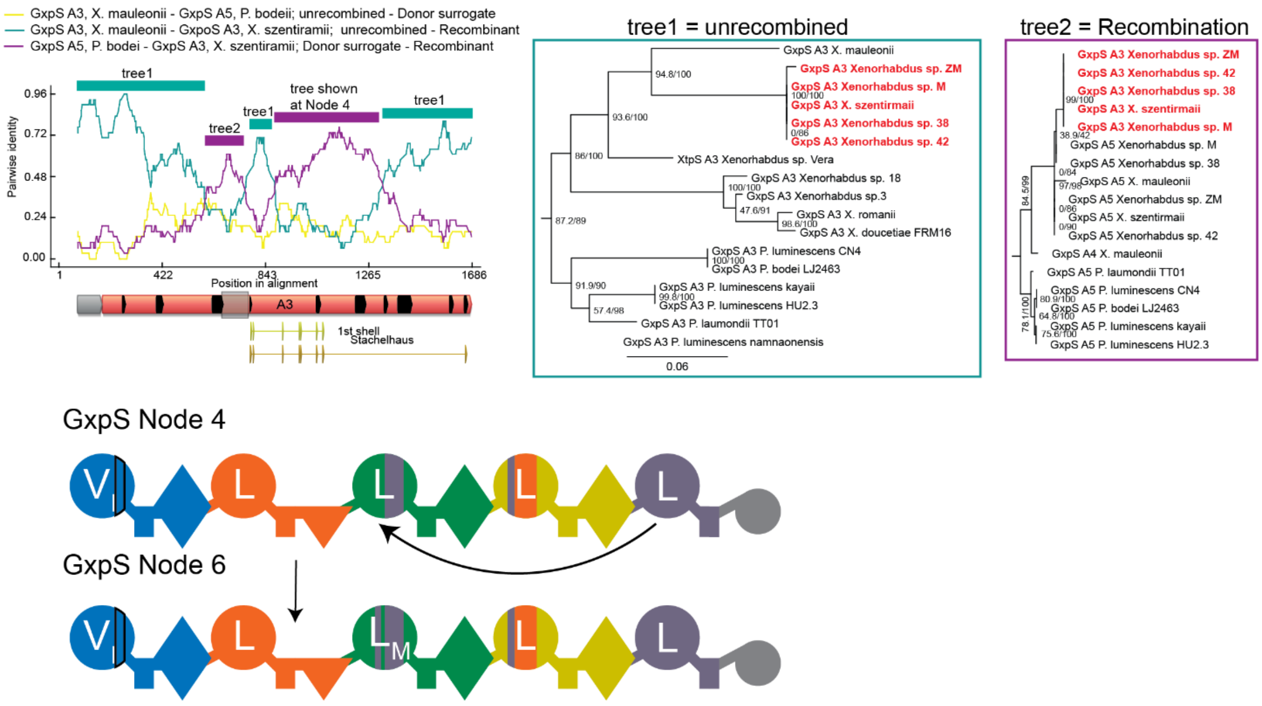


**Figure S66. Recombination analysis of GxpS at Node 6 on the TTE-domain tree.** This recombination was discovered through RDP. RDP and tree analysis as above, tree1 was inferred using amino acids, tree2 using nucleotides. This recombination occurred in a subset of synthetases that gained the recombination described in Supplementary Fig. 64. In this case, another smaller fragment has synonymously recombined into the A3 domain and did not change its specificity. The corresponding nucleotide tree shows that A3 domains are sister to the A5 domains for the same synthetases.

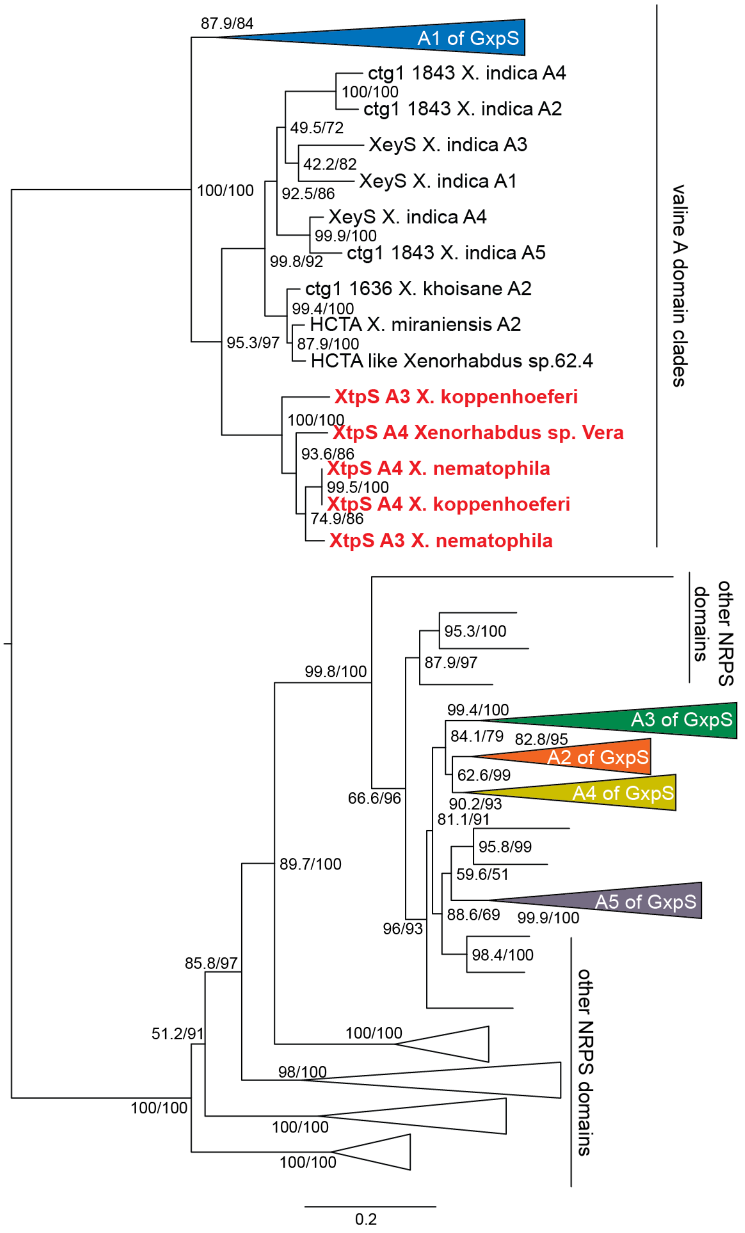


**Figure 67: Discovery of a recombination of A4 and A3 domains from XtpS.** A maximum.-likelihood protein tree of A-domains from all GxpS and some other NRPS that have likely recombined with GxpS. This tree shows that the A4- and A3-domains of XtpS sit in the valine clade but outside of GxpS clades. That indicates that XtpS A-domains have recombined with other NRPS that pulled it from the A4-domains to the valine clade. This would have changed the A4-domain specificity from leucine towards valine. The recombined domains is sister to HCTA [15] synthetases, which are presumably the recombination partner.

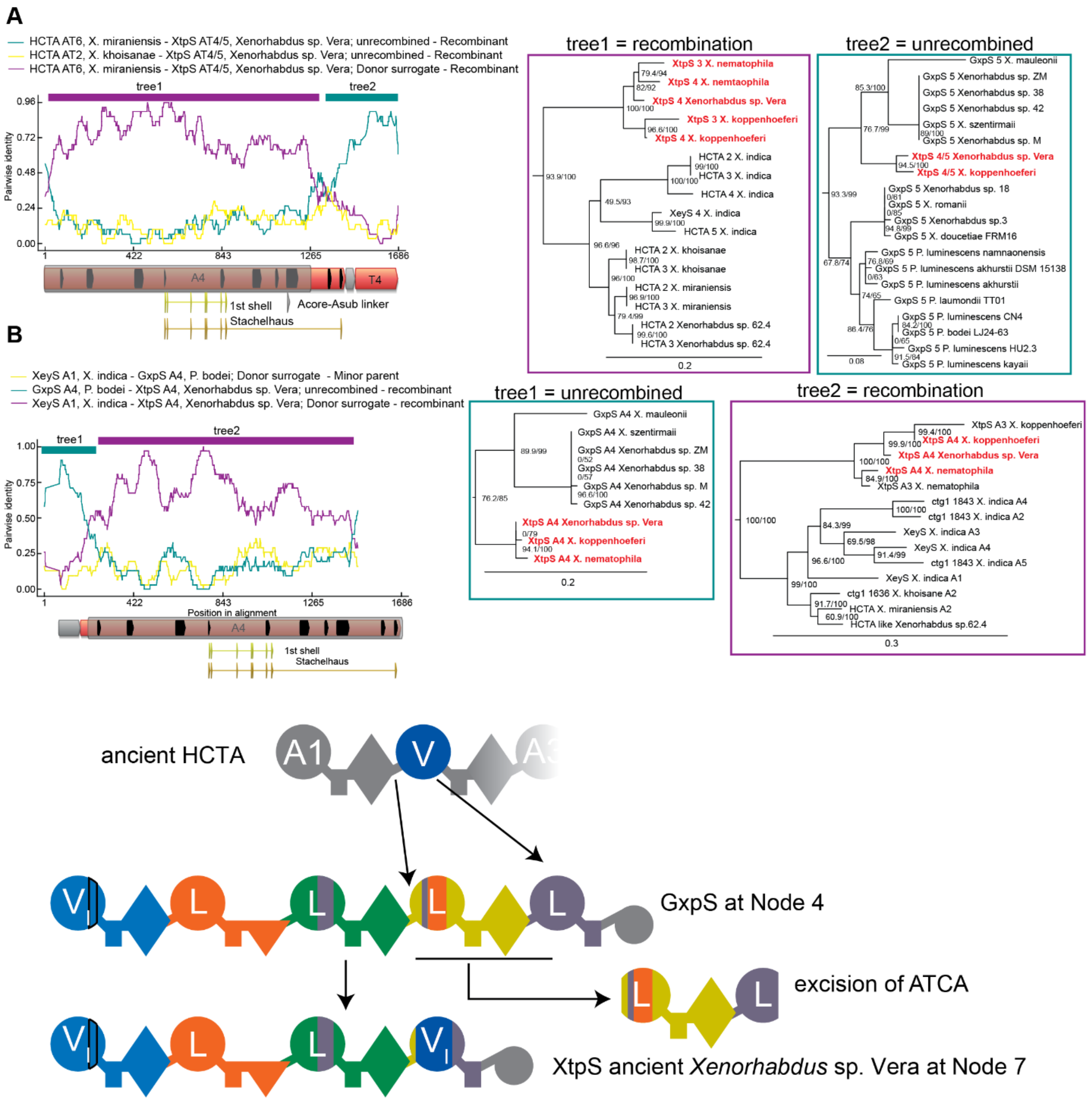


**Figure S68. Recombination analysis at Node 7 on the TTE-domain tree.** The discovery of this recombination is described in the previous figure. This recombination spans an entire module from A4 to A5 (**A**) Identification of the breakpoint in A4. RDP and tree analysis as above, trees were inferred using amino acids. (**B**) Identification of the recombination breakpoint in A5. RDP and tree analysis as above, tree1 was inferred using nucleotides, tree 2 inferred using amino acids. An ancient HCTA [15] NRPS has recombined with parts of the A4 domain and parts of the A5 domain. And thereby excised a whole module. As a result, the valine incorporating HCTA A-domain was inserted in between the ancestral part of A4 and the ancestral part of A5. The result is a four modular GxpS that produces cyclic tetrapeptides.

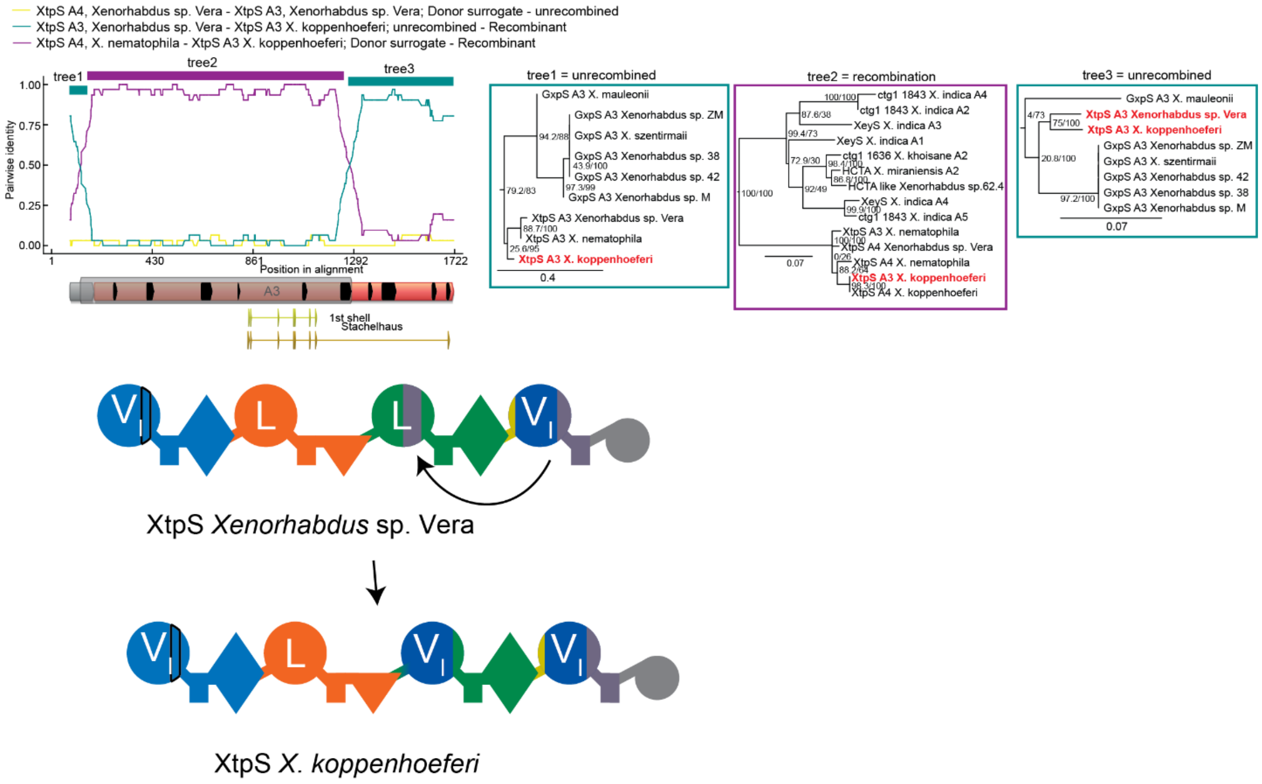


**Figure S69. Recombination analysis of Node 8.** The discovery of this recombination is described in Supplementary Fig. 67. RDP and tree analysis as above, tree 1 was inferred using nucleotides, tree 2 and 3 using amino acids. On tree 2 the A3 domain from *X. koppenhoeferi* is sister to its own A4 domain, indicating a heterotypic, non-synonymous recombination that changed the A3-domain specificity from leucine to valine.

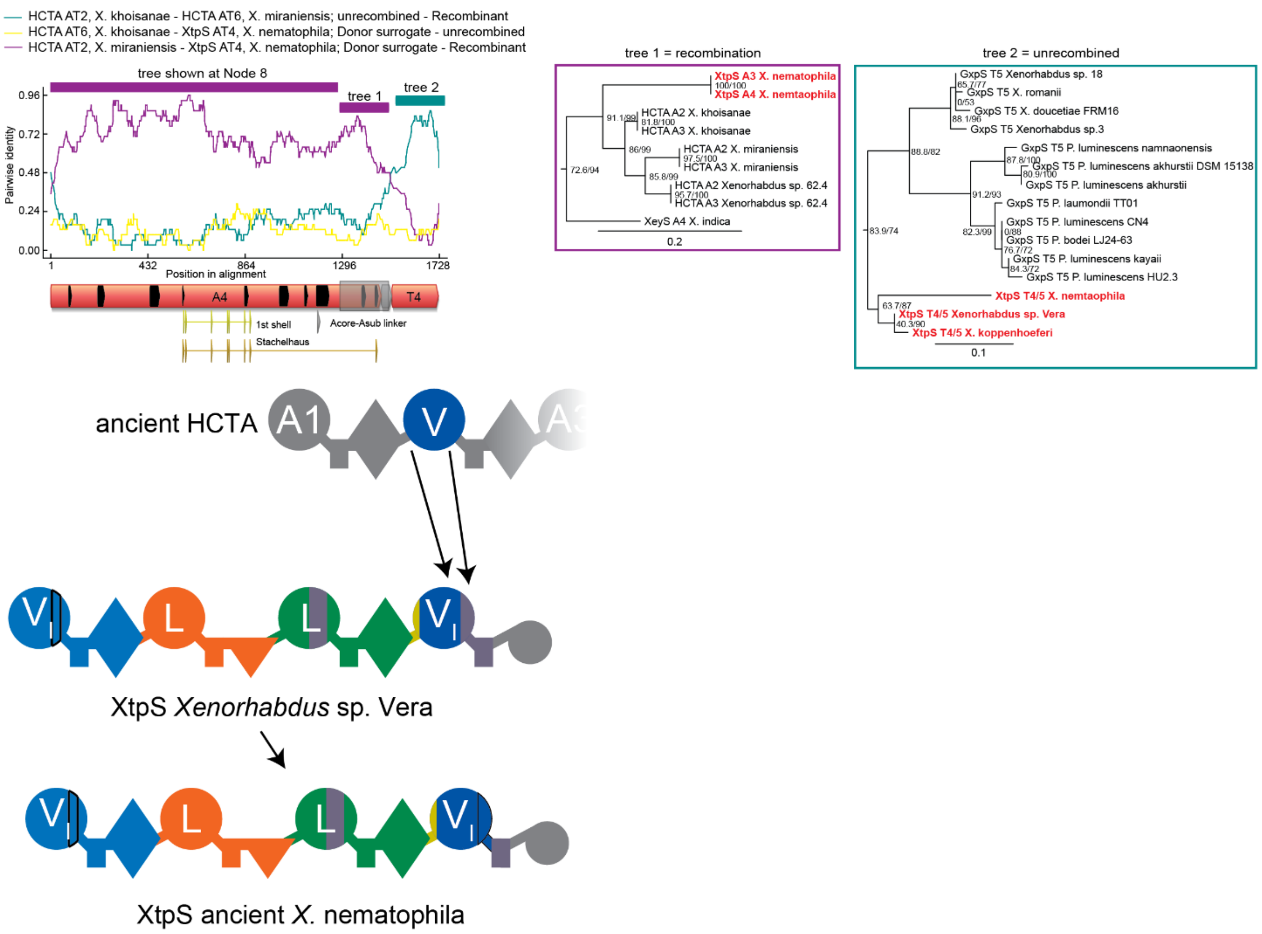


**Figure S70. Recombination analysis of Node 9.** This recombination was discovered through RDP analysis. RDP and tree analysis as above, trees were inferred using nucleotides. This recombination also occurred in what is now the A4 domain of XtpS, specifically *X. nematophila*. This recombination has a border further towards the subsequent T-domain than the one shown in Supplementary Fig. 68. Tree 1 is inferred from only the longer part of this recombination and places that fragment also sister to HCTA [15]. Tree 2 places XtpS T4 inside a clade of T5 domains, because XtpS has lost one module but kept its terminal T5 domain, as described in Supplementary Fig. 68. This implies two possibilities: Either a second synonymous recombination occurred after the first with an HCTA A-domain that overwrote the last part of the A-domain including the two last motifs. Alternatively, GxpS evolved two times independently using the same donor, which we interpret as less likely in the absence of other evidence.

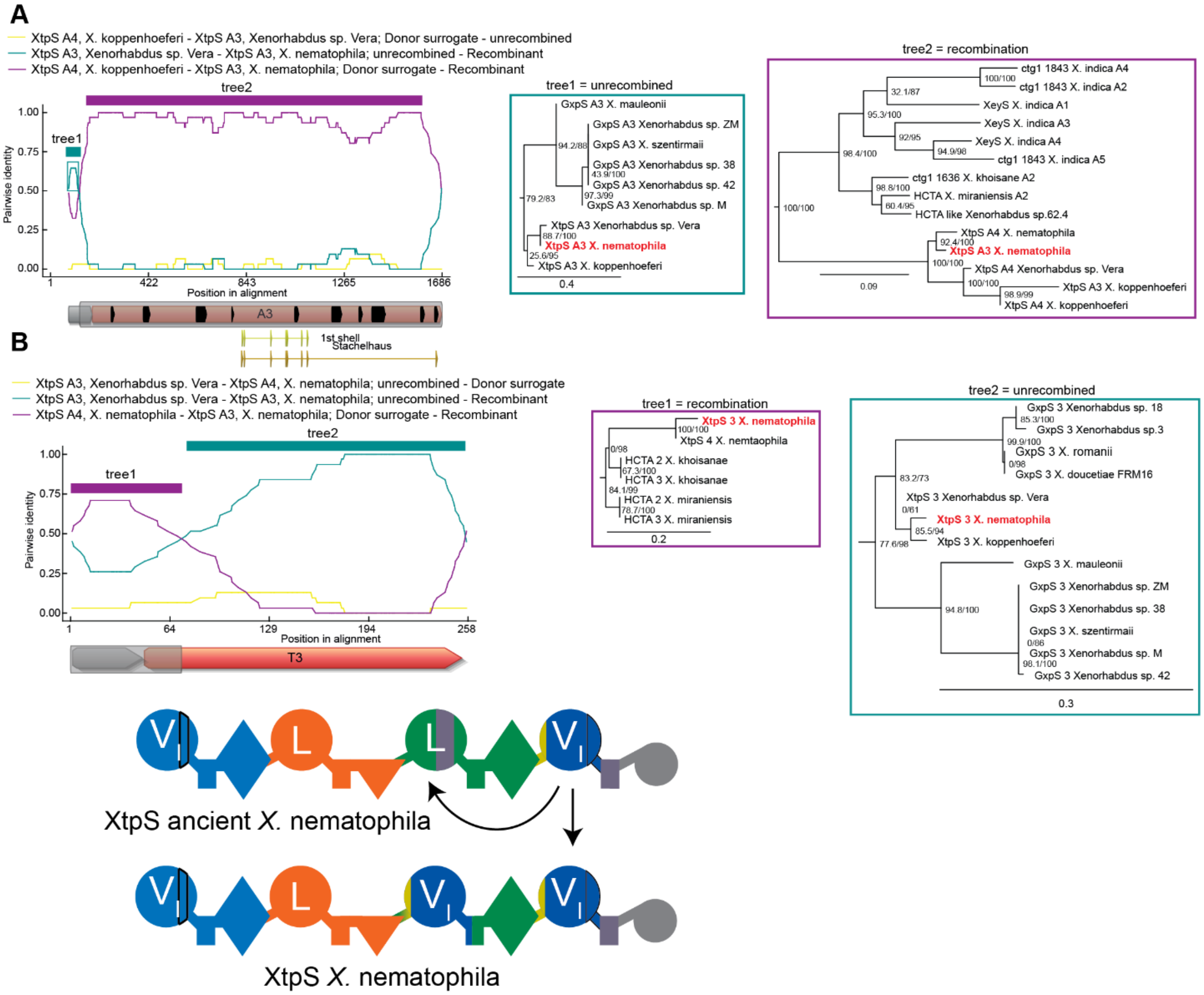


**Figure S71. Recombination analysis of Node 9.** This recombination was identified through the tree in Supplementary Fig. 67. This recombination also affected XtpS of *X. nematophila*. (**A**) Identification of the left breakpoint. RDP and tree analysis as above, trees were inferred with nucleotides. (**B**) Identification of the right breakpoint. RDP and tree analysis as above, trees were inferred using nucleotides. The fourth A-domain of Xtps from *X. nematophila* has non-synonymously recombined into its own A3 domain. It thereby changed its specificity from leucine towards valine.

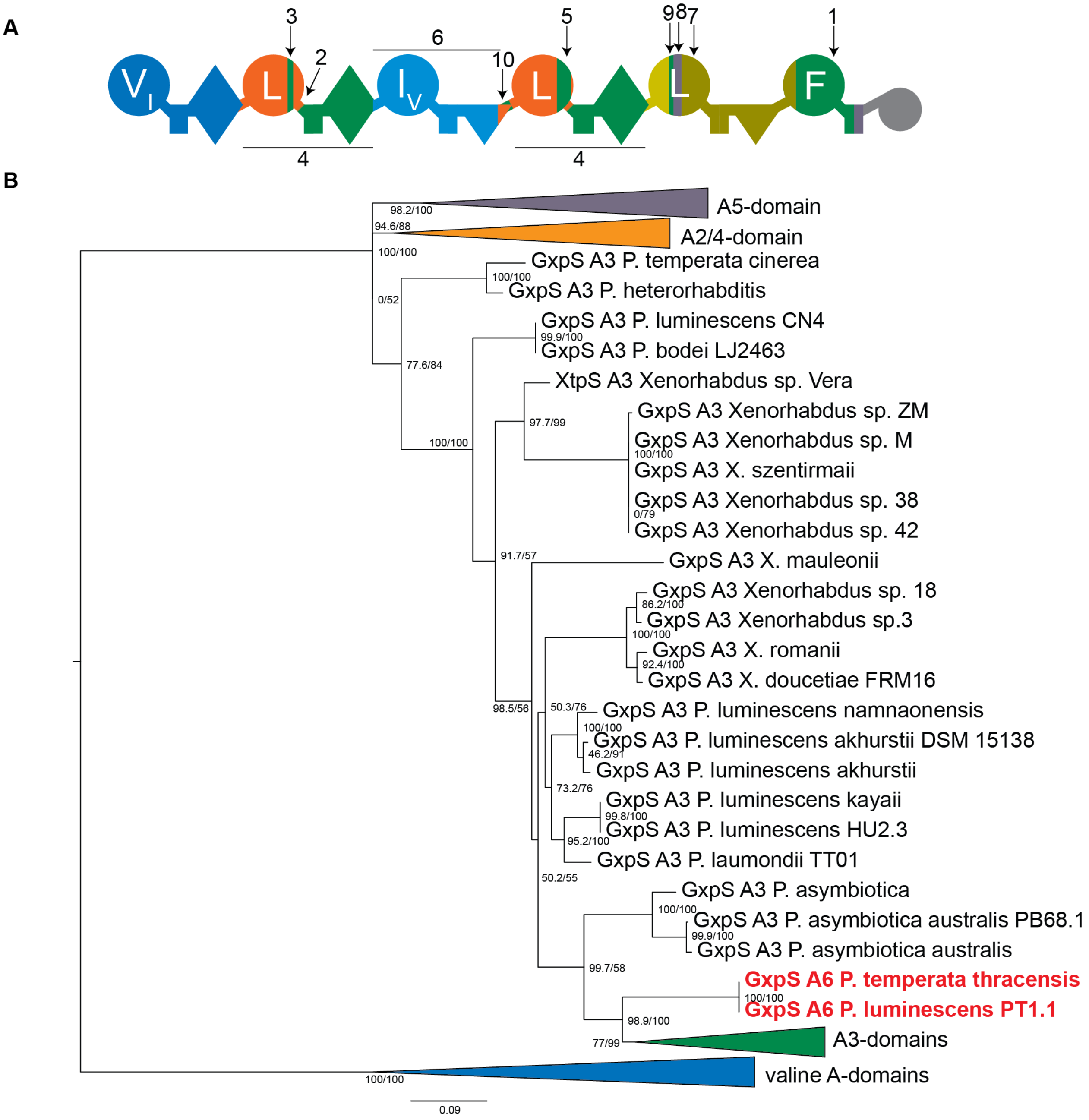


**Figure S72. Evolution of a six-modular GxpS.** (**A**) Domain structure of HexS as found in *P. thracensis* and *P. luminescens* PT1.1. Numbered arrows point to fragments that have recombined in the trajectory leading from a five-modular ancestor to this synthetase. Recombinations 1-10, are shown in Supplementary Fig. 73-78. The numbering is arbitrary and does not refer to the order of these recombinations in time, which we do not know. (**B**) Evidence for the recombination 1 along the trajectory to HexS on a phylogenetic tree of A-domains inferred from amino acids, in which the A6 domain of HexS clades inside the A3 clade of GxpS. This recombination is shown in detail in Supplementary Fig. 73.

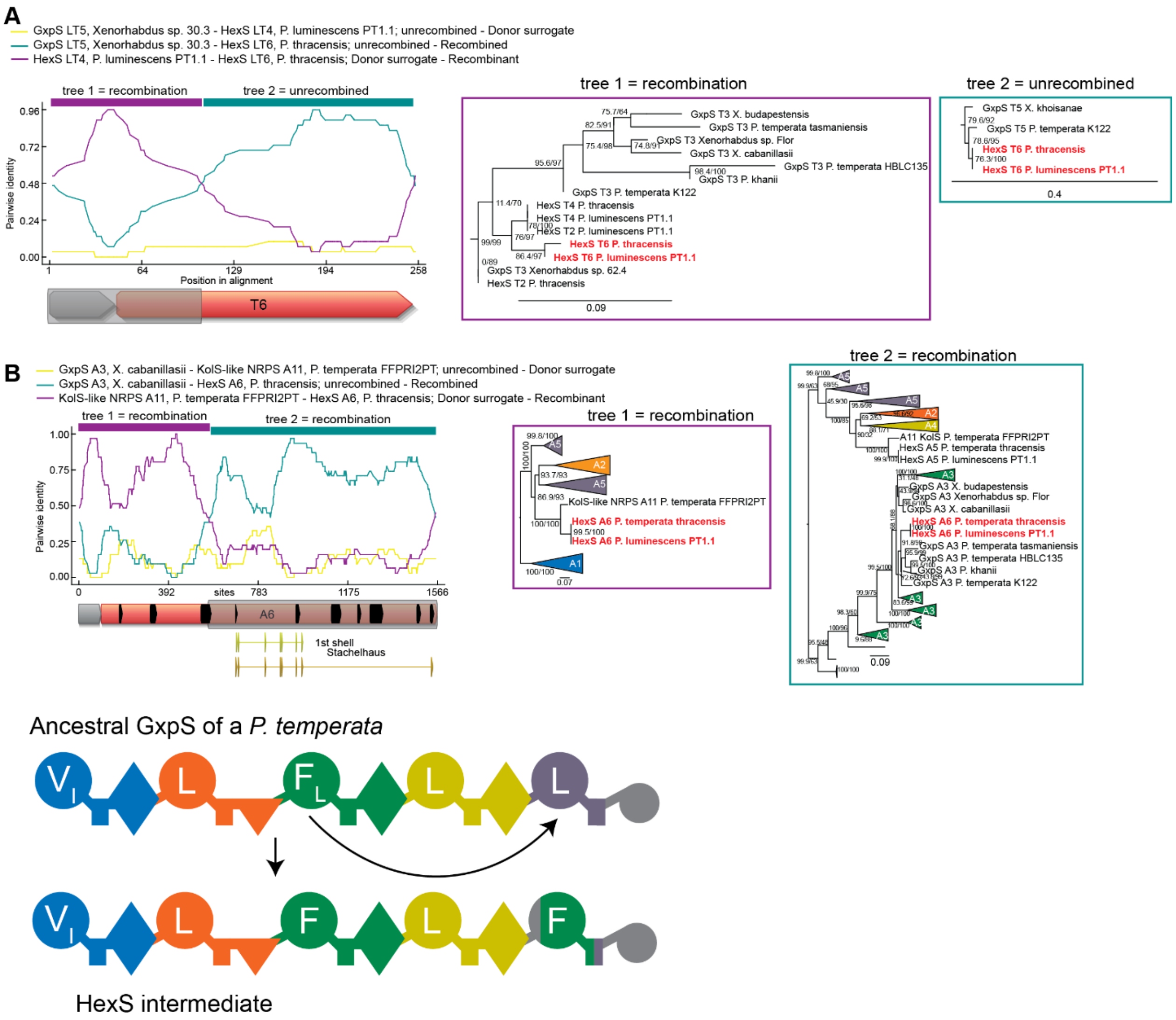


**Figure S73. Recombination 1 at node 10 along the trajectory to HexS.** Recombination analysis of the event described in Supplementary Fig. 72. (**A**) Identification of the right breakpoint using RDP. RDP analysis and tree inference as above, trees were inferred using nucleotides. Tree 2 places the T6 domain of the two HexS synthetases (red) sits in the T5 clade of five modular GxpS, implying that A-domain was inserted and that what was the fifth T-domain GxpS became the sixth in HexS. In tree1 this A-domain sits in the T3 clade, implying that the 3rd T-domain has overwritten what becomes the 6th T-domain. (**B**) Identification of the left breakpoint in the A-domain. RDP analysis and tree inference as above, both trees were inferred using nucleotides. Tree 2 shows the same grouping of HexS A6 domains with A3 GxpS domains as was seen for the subsequent T-domain in (A). This implies that the third A-domain form GxpS overwrote what would become the sixth A-domain in HexS, thereby changing specificity from leucine to phenylalanine. Tree 1 is the result of another recombination event in the same synthetases that is explained in detail in Supplementary Fig. 77. Recombination diagram is shown for five modular synthetases, though this recombination could also have occurred after the module number had been expanded to six.

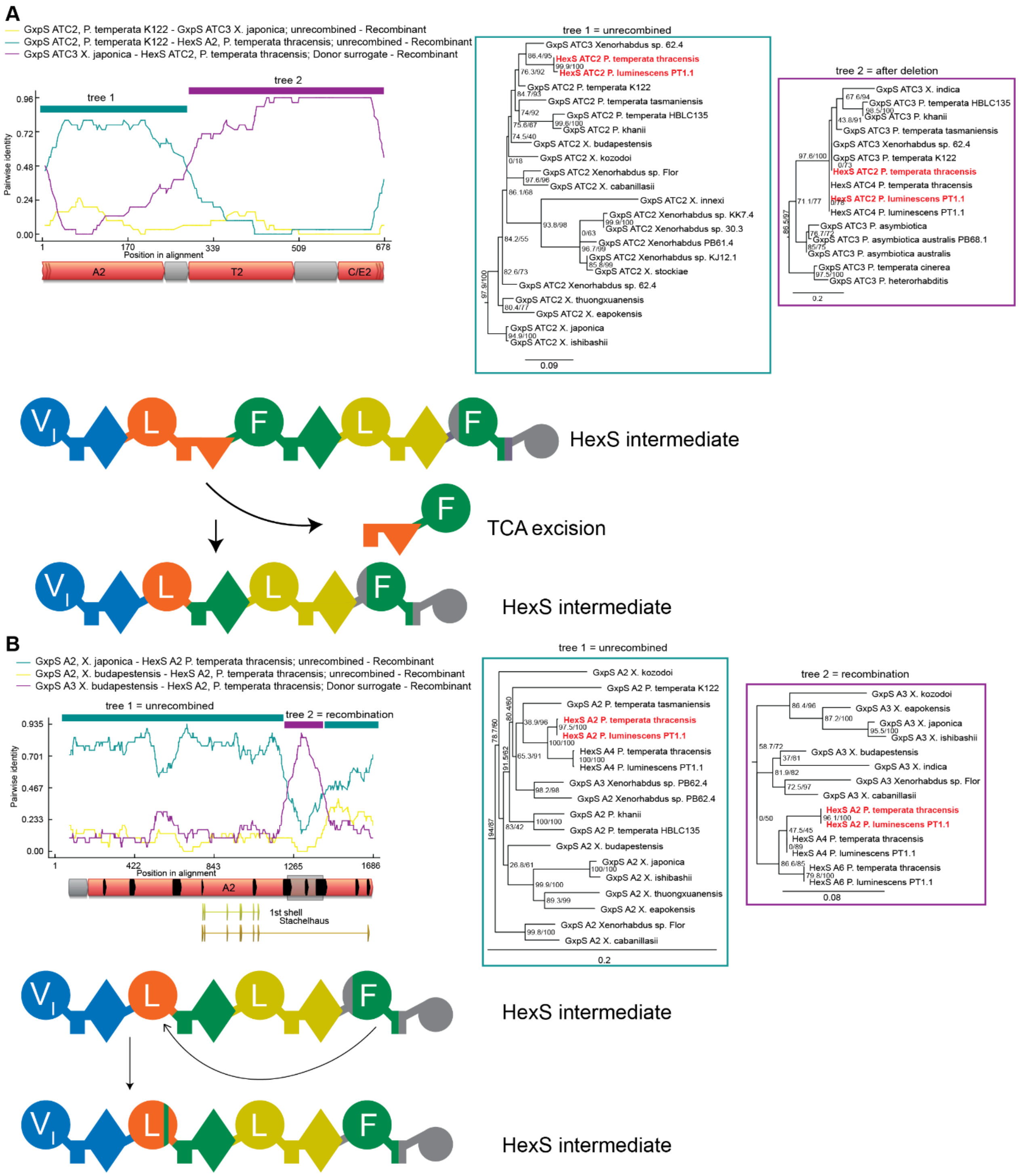

**Figure S74. Recombinations 2 and 3 at node 10 along the trajectory to HexS. (A)** Recombination number two along the trajectory to HexS. RDP analysis and tree inference as above, trees were inferred using nucleotides. Tree 1 shows the A2 domains of HexS inside the A2 GxpS clade. In tree 2, which covers the subsequent T-domain and C-domain the TC2 sequence of HexS clades with the TC3 sequences from GxpS. This pattern indicates that the third A-domain along with the preceding second TC-domain were excised, such that what was the TC3 in GxpS now flanks A2 in HexS. In tree 2, the TC4 domains of HexS are sister to TC2, which is the result of a duplication that we explain in

Supplementary Fig. 75. **(B)** Recombination number two along the trajectory to HexS. RDP analysis and tree inference as above, trees were inferred using nucleotides. In tree 1 the A2 domains form HexS labelled in red clade inside the A2 GxpS clade. In tree 2, they sit sister to HexS A6 clade. The data imply that a small stretch of what becomes the last A-domain of HexS recombined into HexS A2 domain. In both tree 1 and tree 2, the A2 domains in red are sister to the A4 domains from HexS. This is caused by a subsequent ATC-tridomain duplication that is explained in Supplementary Fig. 75.

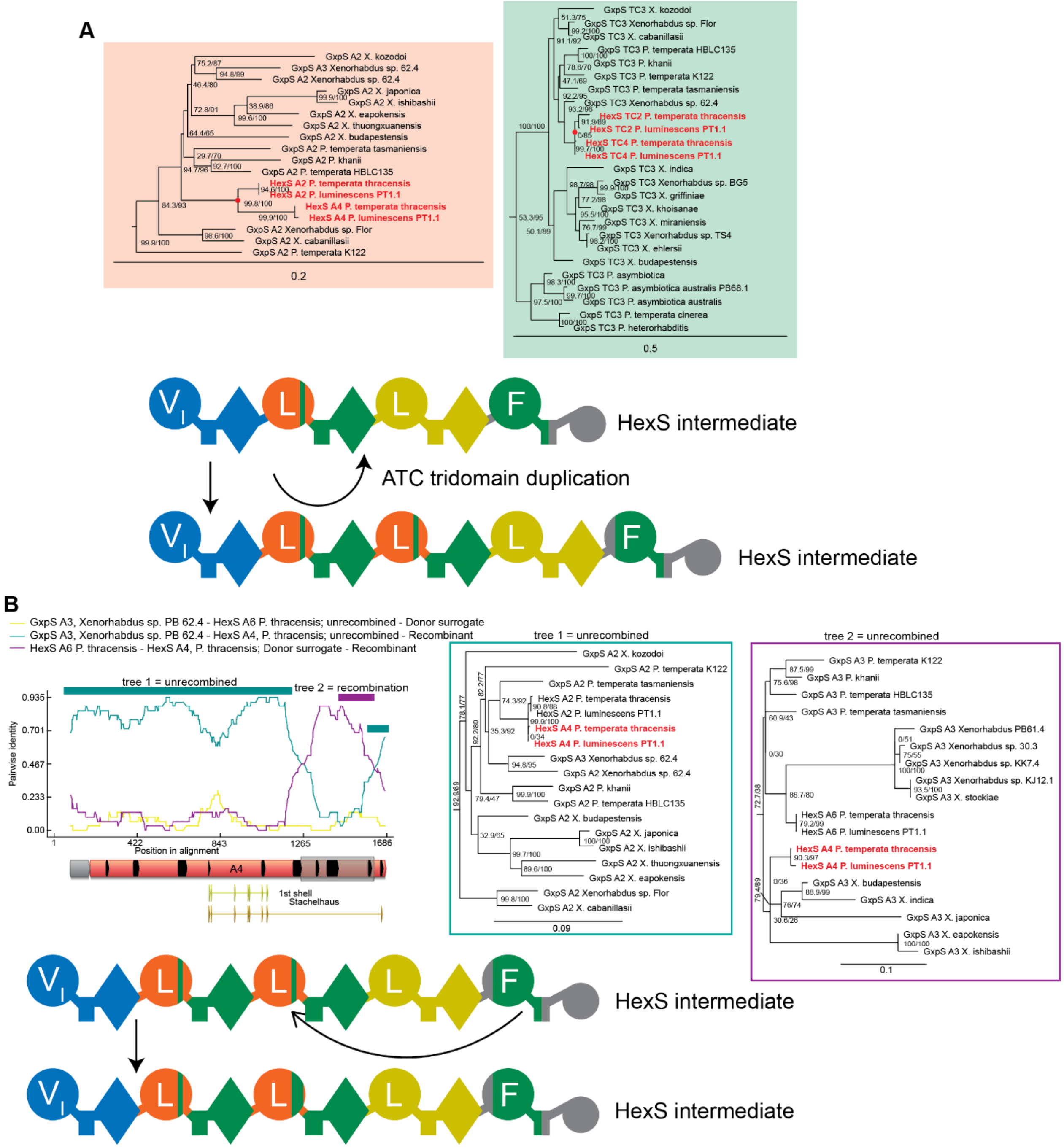

**Figure S75: Recombinations 4 and 5 at node 10 along the trajectory to HexS.** (**A**) Recombination 4 along the trajectory to HexS. A-domain and the TC-domain protein tree and in red the duplication node. The A2 and A4 domain as well as the TC2 and TC4 domains of *P. thracensis* and *P. luminescens* PT1.1, respectively, are sister to each other, indicating that an ancestral tri-domain duplicated. Both the A2 and A4 domains carry the small recombination fragment from the sixth A-domain described in Supplementary Fig. 74, implying that this small recombination happened first, and was thus inherited by the two duplicated domains. Later, an ATC-tridomain was inserted between these two domains, making them A2 and A4, instead of A2 and A3 as shown in the diagram at the bottom (see Supplementary Fig. 76). (**B**) Recombination 5 along the trajectory to HexS. This recombination was discovered using RDP. RDP analysis and tree inference as previously, trees were inferred using nucleotides. Here in additional small fragment originating from what becomes the 6$^{th}$ A-domain of HexS or a phenylalanine specific A3 domain from GxpS recombined into what becomes A4 in HexS. This additional recombination is exclusive to A4: tree 2 for the additional recombined stretch only places HexS A4 domains, but not HexS A2 domains in the phenylalanine clade.

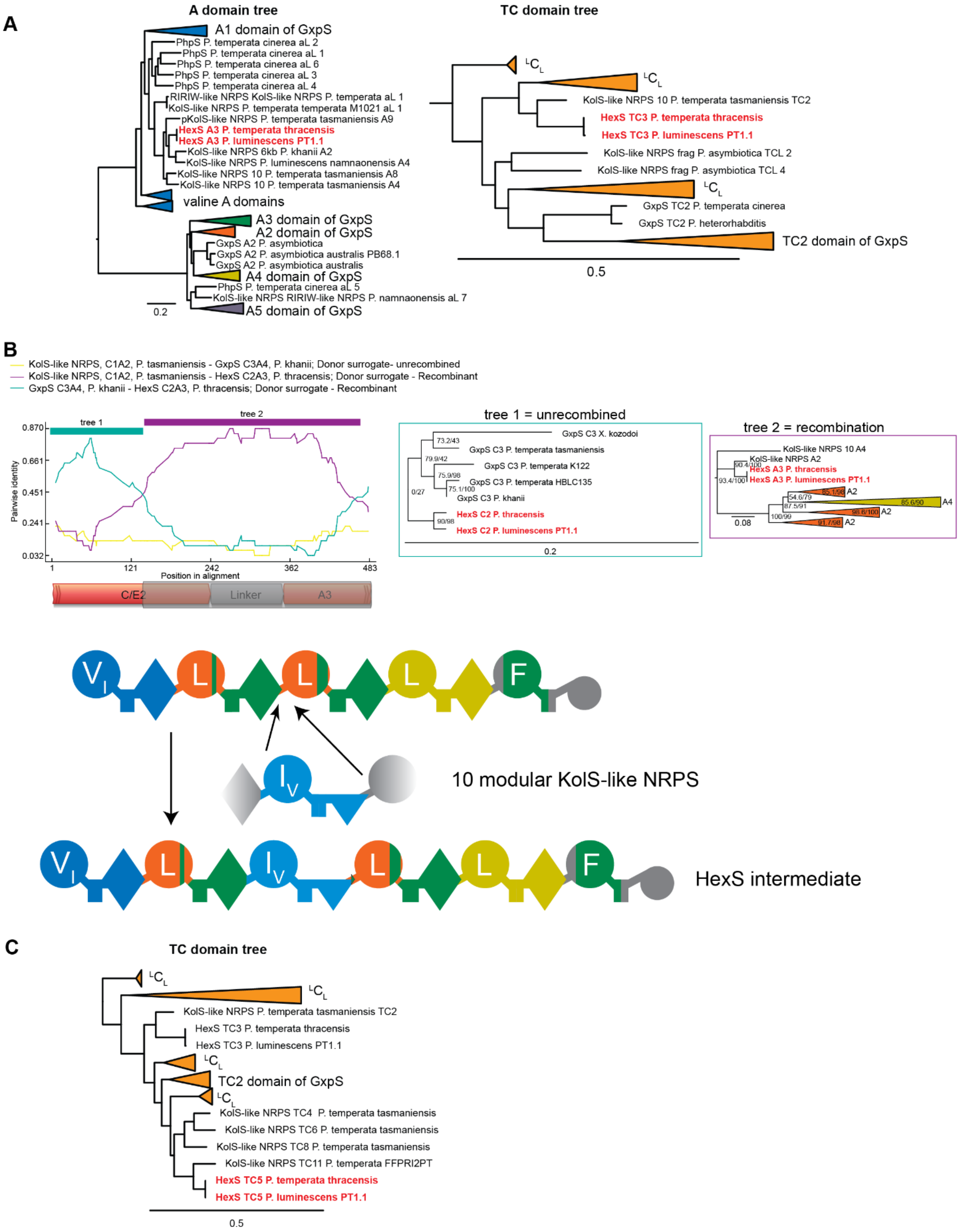


**Figure S76. Recombination 6 at node 10 along the trajectory to HexS. (A)** Discovery of recombination 6 of HexS. Left: The GxpS A-domain tree inferred from amino acids with additional sequences from other NRPS. Shown in red are the third A-domains from HexS, which clade with non-

GxpS A-domains. Right: TC-domain tree inferred from amino acids with additional sequences from other NRPS. The third TC-domains of the HexS (red) also calde with non-GxpS $^{L}C_{L}$s from similar synthetases as on the A-domain tree. The tree topology of both trees indicates that the third ATC-domain of HexS descends from another NRPS. This ATC tri-domain has likely recombined into GxpS to give rise to a six-modular GxpS. (**B**) Analysis of recombination 6 and identification of the instertion site of the event described in A. RDP analysis and tree inference as previous, trees were inferred using nucleotides. Tree 1 shows that HexS C2 sits inside the C3 clade. This is because of the module TCA excision described in Supplementary Fig. 74. Tree 2 places the the subsequent C and A-domain sites in a group of non-GxpS NRPS, indicating the ATC tri-domain inserted into the very end of the C2 domain of HexS. (**C**) TC-domain phylogeny inferred from amino acids that guided us to recombination 7. HexS TC5 groups with TC-domains from another NRPS similar to KolS, not with GxpS TC-domains, indicating recombination. This would have replaced a dual C/E-domain with an $^{L}C_{L}$.

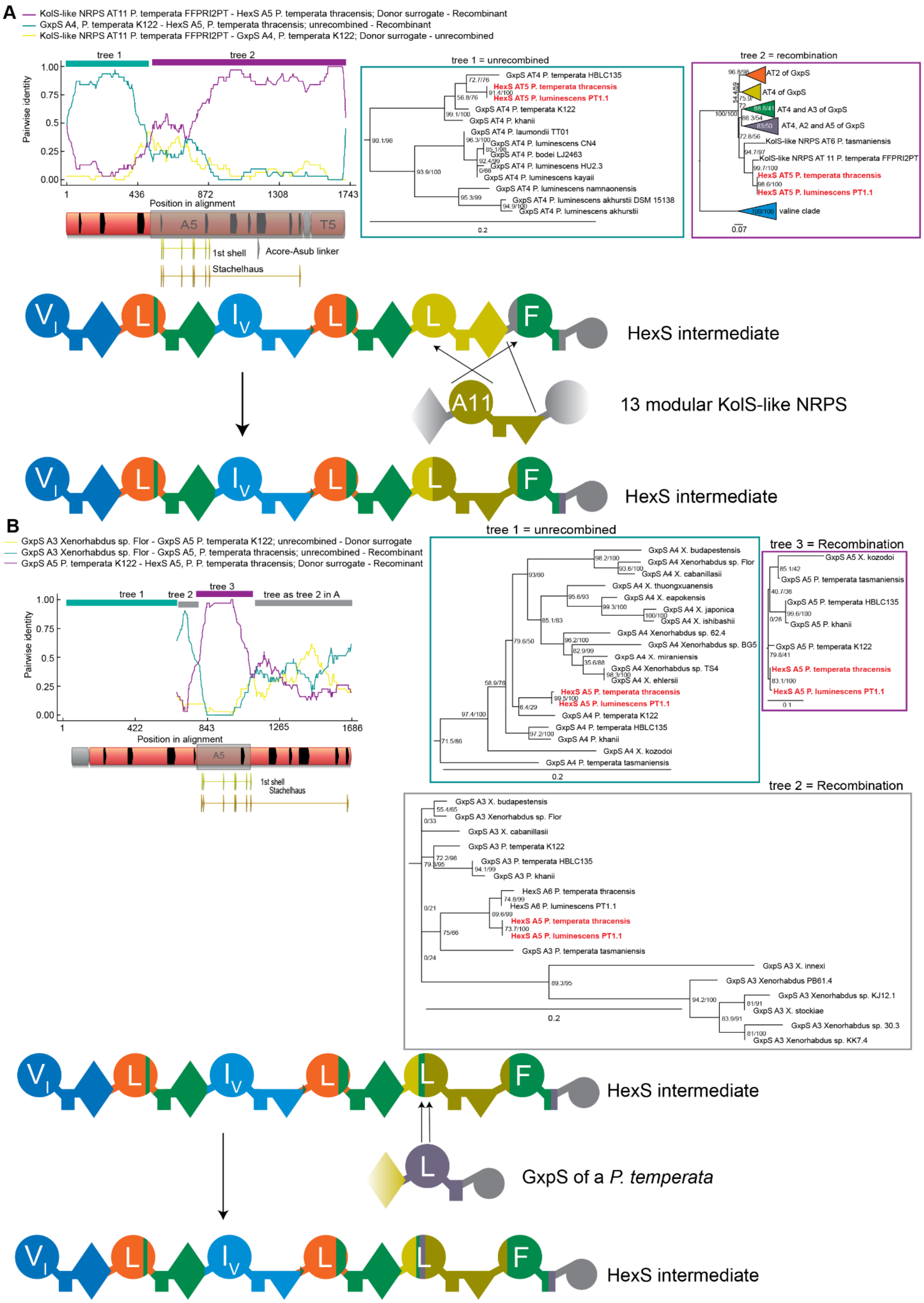
A
KolS-like NRPS AT11 P. temperata FFPRI2PT - HexS A5 P. temperata thracensis; Donor surrogate - Recombinant
GxpS A4, P. temperata K122 - HexS A5, P. temperata thracensis; unrecombined - Recombinant
KolS-like NRPS AT11 P. temperata FFPRI2PT - GxpS A4, P. temperata K122; Donor surrogate - unrecombined
tree 1
tree 2
Pairwise identity
Position in alignment
A5
T5
1st shell
Acore-Asub linker
Stachelhaus
tree 1 = unrecombined
GxpS AT4 P. temperata HBLC135
HexS AT5 P. temperata thracensis
HexS AT5 P. luminescens PT1.1
GxpS AT4 P. temperata K122
GxpS AT4 P. khanii
GxpS AT4 P. laumondii TT01
GxpS AT4 P. luminescens CN4
GxpS AT4 P. bodei LJ2463
GxpS AT4 P. luminescens HU2.3
GxpS AT4 P. luminescens kayaii
GxpS AT4 P. luminescens namnaonensis
GxpS AT4 P. luminescens akhurstii DSM 15138
GxpS AT4 P. luminescens akhurstii
tree 2 = recombination
AT2 of GxpS
AT4 of GxpS
AT4 and A3 of GxpS
AT4, A2 and A5 of GxpS
KolS-like NRPS AT6 P. tasmaniensis
KolS-like NRPS AT 11 P. temperata FFPRI2PT
HexS AT5 P. temperata thracensis
HexS AT5 P. luminescens PT1.1
valine clade
HexS intermediate
A11
13 modular KolS-like NRPS
HexS intermediate
B
GxpS A3 Xenorhabdus sp. Flor - GxpS A5 P. temperata K122; unrecombined - Donor surrogate
GxpS A3 Xenorhabdus sp. Flor - GxpS A5, P. temperata thracensis; unrecombined - Recombinant
GxpS A5 P. temperata K122 - HexS A5, P. P. temperata thracensis; Donor surrogate - Recominant
tree 1
tree 2
tree 3
tree as tree 2 in A
Pairwise identity
Position in alignment
A5
1st shell
Stachelhaus
tree 1 = unrecombined
GxpS A4 X. budapestensis
GxpS A4 Xenorhabdus sp. Flor
GxpS A4 X. cabanillasii
GxpS A4 X. thuongxuanensis
GxpS A4 X. eapokensis
GxpS A4 X. japonica
GxpS A4 X. ishibashii
GxpS A4 Xenorhabdus sp. 62.4
GxpS A4 Xenorhabdus sp. BG5
GxpS A4 X. miranensis
GxpS A4 Xenorhabdus sp. TS4
GxpS A4 X. ehlersii
HexS A5 P. temperata thracensis
HexS A5 P. luminescens PT1.1
GxpS A4 P. temperata K122
GxpS A4 P. temperata HBLC135
GxpS A4 P. khanii
GxpS A4 X. kozodoi
GxpS A4 P. temperata tasmaniensis
tree 3 = Recombination
GxpS A5 X. kozodoi
GxpS A5 P. temperata tasmaniensis
GxpS A5 P. temperata HBLC135
GxpS A5 P. khanii
GxpS A5 P. temperata K122
HexS A5 P. temperata thracensis
HexS A5 P. luminescens PT1.1
tree 2 = Recombination
GxpS A3 X. budapestensis
GxpS A3 Xenorhabdus sp. Flor
GxpS A3 X. cabanillasii
GxpS A3 P. temperata K122
GxpS A3 P. temperata HBLC135
GxpS A3 P. khanii
HexS A6 P. temperata thracensis
HexS A6 P. luminescens PT1.1
HexS A5 P. temperata thracensis
HexS A5 P. luminescens PT1.1
GxpS A3 P. temperata tasmaniensis
GxpS A3 X. innexi
GxpS A3 Xenorhabdus PB61.4
GxpS A3 Xenorhabdus sp. KJ12.1
GxpS A3 X. stockiae
GxpS A3 Xenorhabdus sp. 30.3
GxpS A3 Xenorhabdus sp. KK7.4
HexS intermediate
GxpS of a P. temperata
HexS intermediate

**Figure S77. Recombinations 7, 8 and 9 at node 10 along the trajectory to HexS. (A)** Recombination analysis of the event discovered in Supplementary Fig. 76c. RDP analysis and tree inferred as previous, tree 1 and tree 2 were inferred using amino acids. In tree 1, the A5 domain form HexS groups inside A4 domains of GxpS. This is because of the module insertion described in Supplementary Fig. 76, which shifts the numbering of domains in HexS. In tree 2 the sites spanning parts of the AT5 domains of HexS now group with non-GxpS sequences similar to Kolossin, indicating recombination. The dip in the purple line at the beginning of the sequence that is assigned to tree 2 in the RDP plot is caused by another recombination event, which we explain in (B). The right breakpoint of this recombination event is discovered in Supplementary Fig. 73a, where it is located in the beginning of what is now the A6 domain of HexS. This requires that an ATC tridomain fragment of KolS likely first circularized itself prior to recombination, as shown in our recombination diagram. This is necessary because the 3' end of Kolossin's A11 domain recombined into A6 of HexS, whereas 5' end of Kols A11 recombined into A5 of HexS. Without prior circularization of the recombination donor, we would require multiple independent recombination events with KolS to explain the sequence similarity and phylogenetic patterns. (**B**) Recombination 8 along the trajectory to HexS was discovered using RDP. Trees were inferred using nucleotides. In tree 2, the A5 domain of HexS groups with A5 domains from GxpS. This indicates recombination, because the A5 domain of HexS would normally correspond to the A4 domain of GxpS due to ATC-tridomain insertion in HexS. Tree 1 covers an area that RDP detects as part of the recombination analysis shown in A and therefore ignores. It shows A5 to group with A4, as expected for the A5 domain of HexS. Tree 2 shows that a very small fragment of HexS A5 domain is similar to its own A6 domain, which RDP detects as similarity to A3 domains (from which the phenylalanine-encoding A6 descends). The breakpoints of this recombination are difficult to ascertain, because it is directly buffeted on its right border by recombination 9. Tree 3, which describes this area, places HexS' A5 domain in the GxpS A5 clade. This indicates recombination, because HexS A5 domain normally descends from GxpS' A4 domain. The area to the right of this recombination is affected by the recombination described in (A). In the RDP plot (left), this part of the sequence is therefore similar to neither A3 nor A5 domains from GxpS.

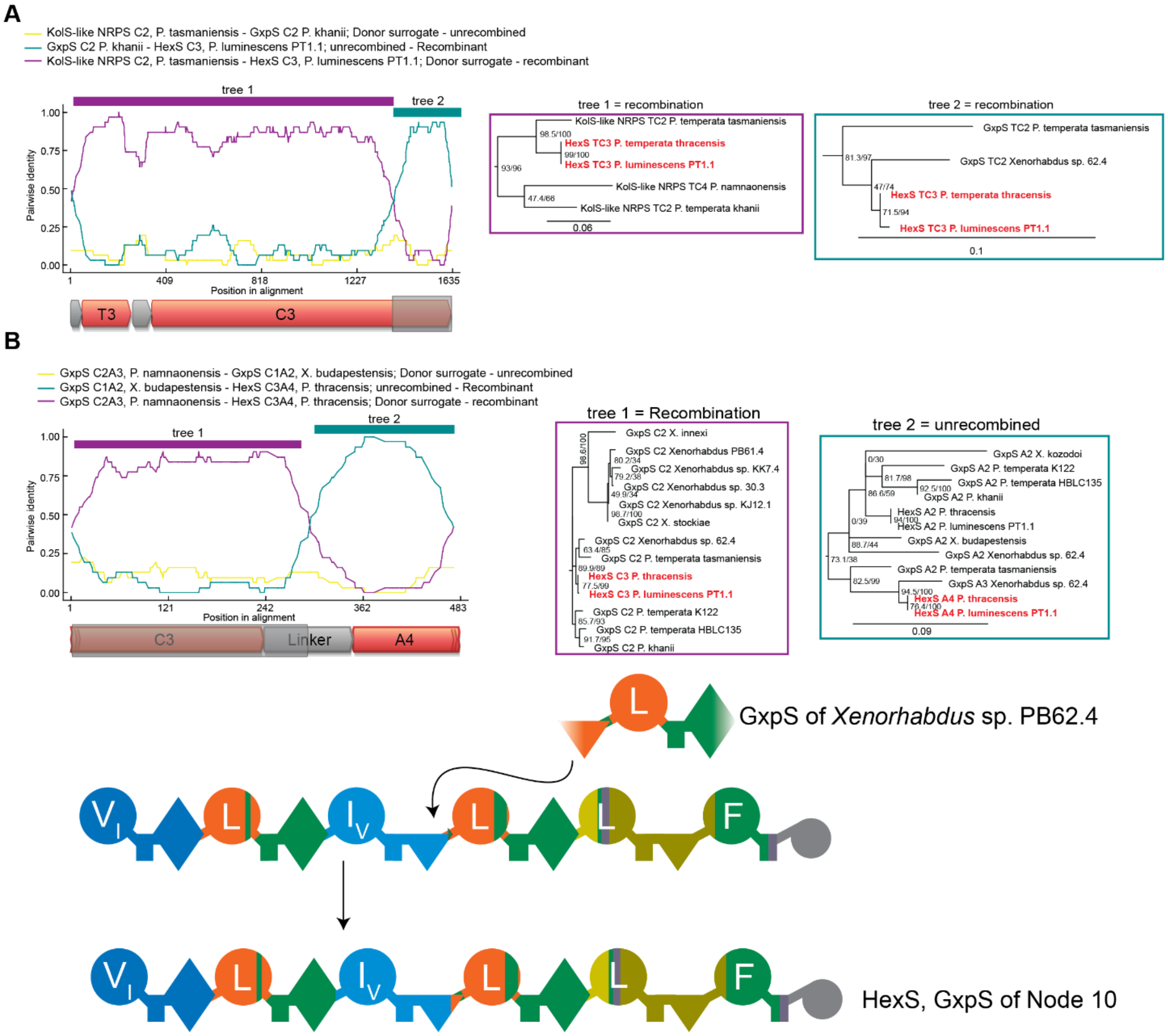


**Figure S78. Recombination 10 at node 10 along the trajectory to HexS.** This recombination was discovered indirectly using RDP. (**A**) Recombination diagram implied by the analysis that follows: A fragment of the C2-A3 sequence of *Xenorhabdus* sp. PB62.4 recombined into the C3-A4 segment of HexS. The *Xenorhabdus sp*. PB62.4 A3 domain was overwritten by its own A2 domain, which we explain in Supplementary Fig. 92. Identification of the left recombination border using RDP. In this case, RDP did not recognize the correct recombination partners and direction, because of the confounding influence of *Xenorhabdus* sp. PB62.4's A3 recombination. Tree 1 (inferred from amino acids) shows that HexS C3 domains (red) sit inside the $^{L}C_{L}$ clade near a sequence from a Kolossin-like NRPS, as described in Supplementary Fig. 76, due to ATC-tridomain insertion. Tree 2 (inferred from nucleotides) shows that the end of HexS' is sister to the C2 domain from *Xenorhabdus sp*. PB62.4's. RDP does not recognize this and instead shows it being similar to a C2 domain from a different GxpS (from *X. khanii*). (**B**) Identification of the right breakpoint. Here RDP again misidentifies the donor and recipient: Towards the end of the C3 domain and through the middle of the linker, HexS is inferred to be similar to a C2 domain and linker from another GxpS. Subsequently, it becomes similar to an A2 domain. The trees from these two areas explain why this is a spurious result: In both trees, the HexS sequences (in red) are close to the sequence from *Xenorhabdus sp*. PB62.4. The data are much better explained if we assume that HexS recombined with the C2-A3 domain from *Xenorhabdus sp*. PB62.4: This makes its sequence similar to C2 domains in the remining part of the HexS' C3 domain. In HexS' sequence it becomes similar to A2 sequences, from which *Xenorhabdus* sp. PB62.4's A3 descends.

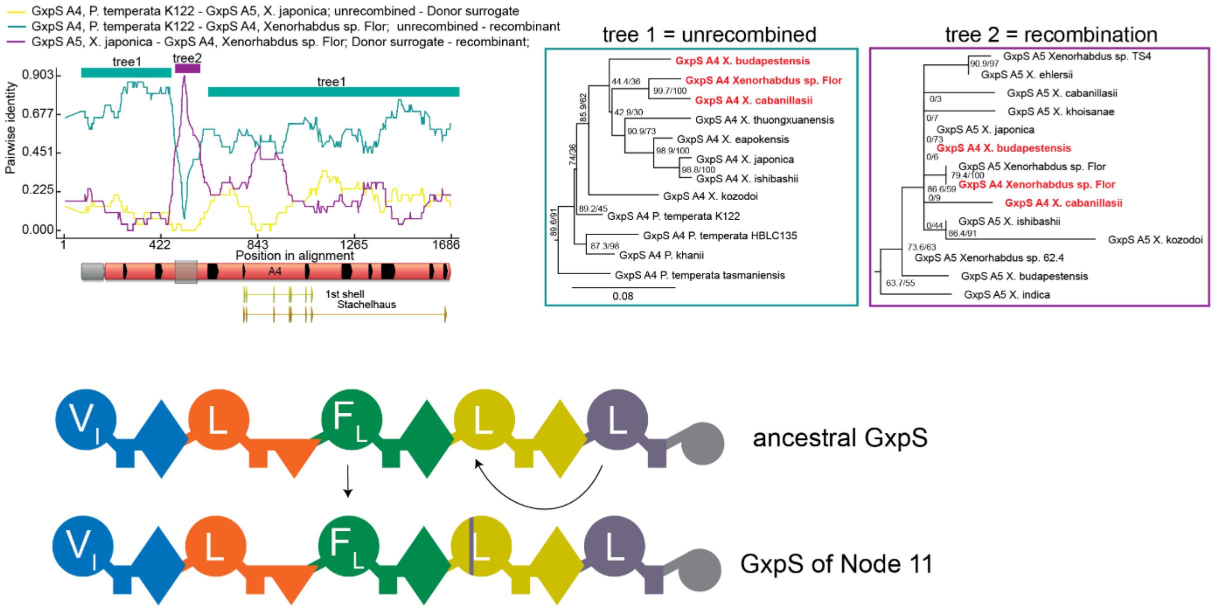


**Figure S79. Recombination analysis of Node 11.** This recombination was discovered using RDP. RDP analysis and tree inference as above, tree 1 was inferred using amino acids, tree 2 using nucleotides. Here a very short stretch of an A5 domain overwrote part of an A4 domain without changing specificity.

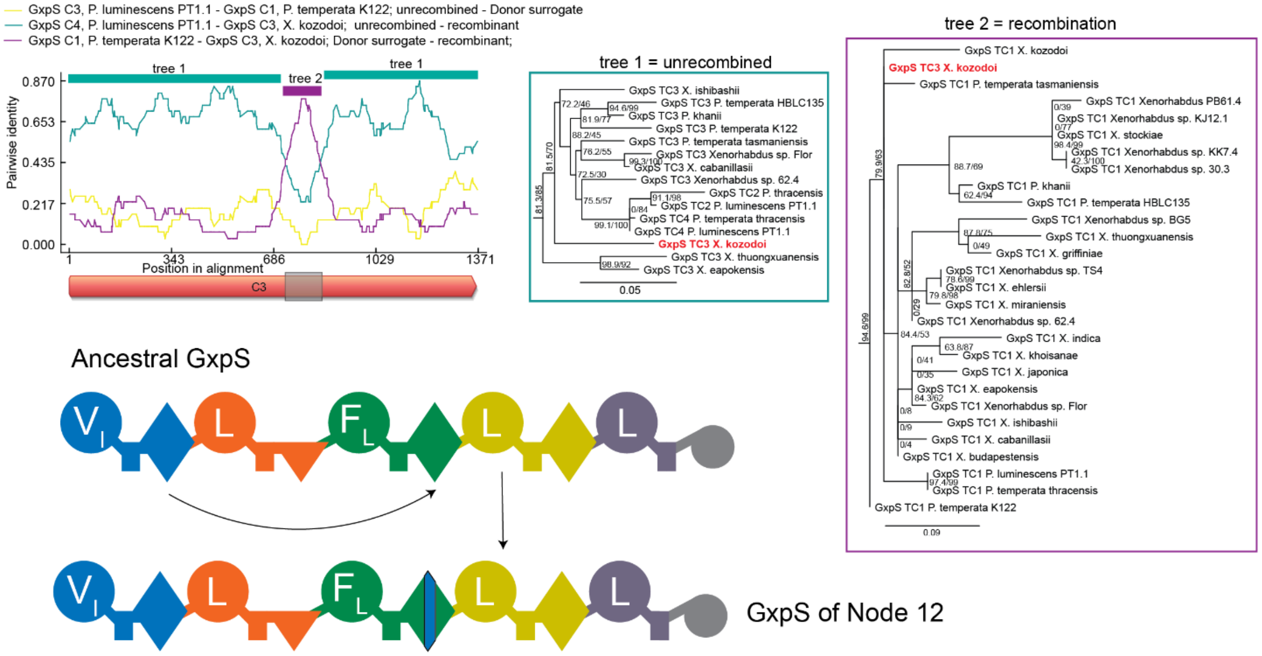


**Figure S80. Recombination analysis of Node 12.** This recombination was discovered using RDP. RDP analysis and tree inference were carried out as above, tree 1 was inferred using amino acids and tree 2 using nucleotides. Here a small part of the first dual C/E-domain overwrote a fragment of the fourth dual C/E-domain. This recombination was synonymous.

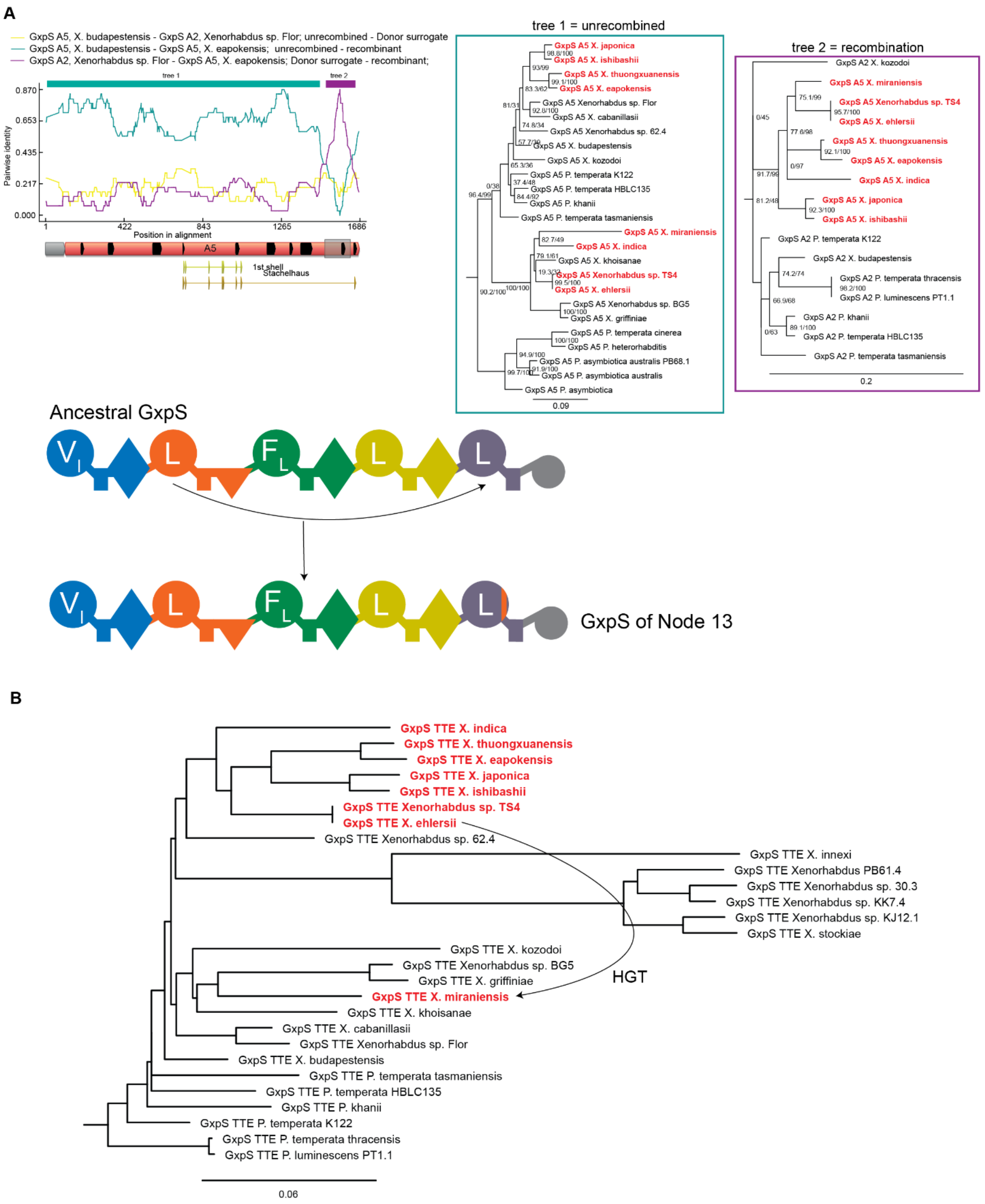


**Figure S81. Recombination analysis of Node 13.** This recombination was discovered through RDP. (**A**) RDP analysis and tree inference as above, tree 1 was inferred using amino acids, tree 2 using nucleotides. In tree 1, the unrecombined sequences are not monophyletic, whereas the recombined sequences are monophyletic in tree 2. This is problematic, because it would imply multiple independent recombinations that all used the same or very similar borders. We suspected this was the case because the A-domain alignment contained other domains that underwent recombinations that distort the topology of tree1. We therefore examined the position of these synthetases on our TTE-domain tree (**B**), where most of the recombinant synthetases are monophyletic, with the exception of the GxpS of *X. miraniensis*. To explain how this synthetase shares the same recombination event with the other ones highlighted in red, we have to propose a subsequent recombination of the A5 domain in *X. miraniensis* with an already recombined A5 domain from GxpS from one of the other species highlighted in red. We could not, however, determine the borders of that second event.

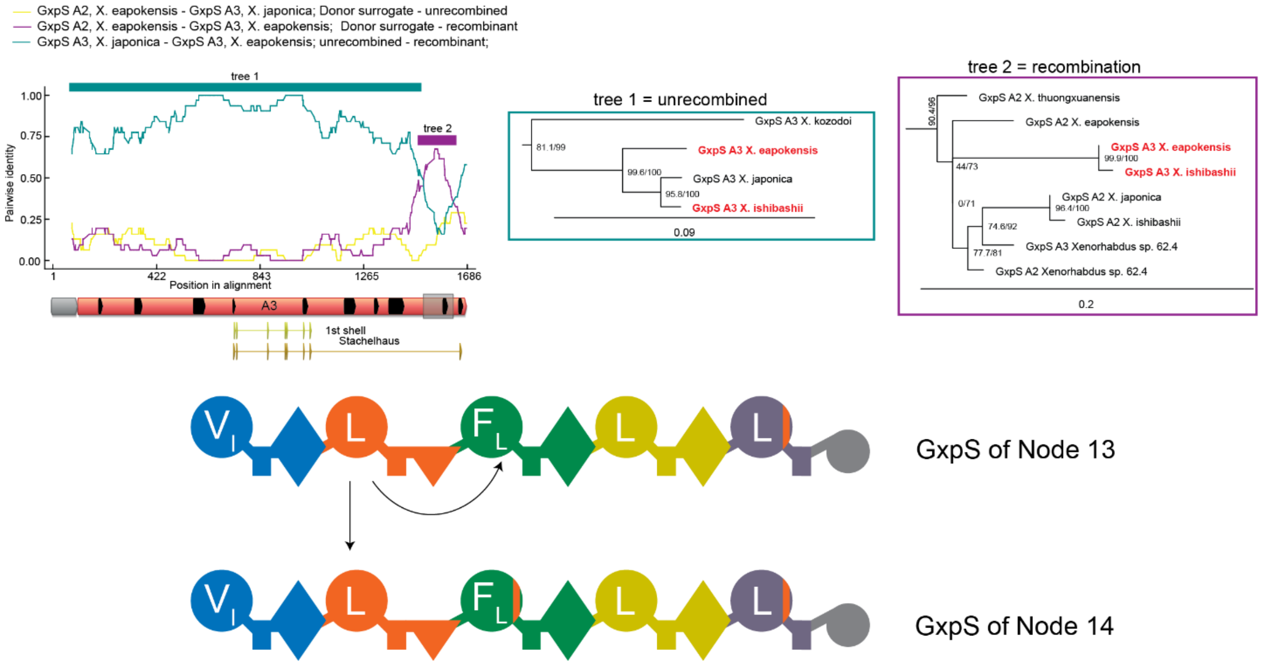


**Figure S82. Recombination analysis of Node 14.** This recombination was discovered using RDP. RDP analysis and tree inference as described above, tree 1 was inferred using amino acids, tree 2 using nucleotides. This is an example of a heterotypic, but synonymous recombination because the recombination occurred outside of the specificity encoding region of the A-domain.

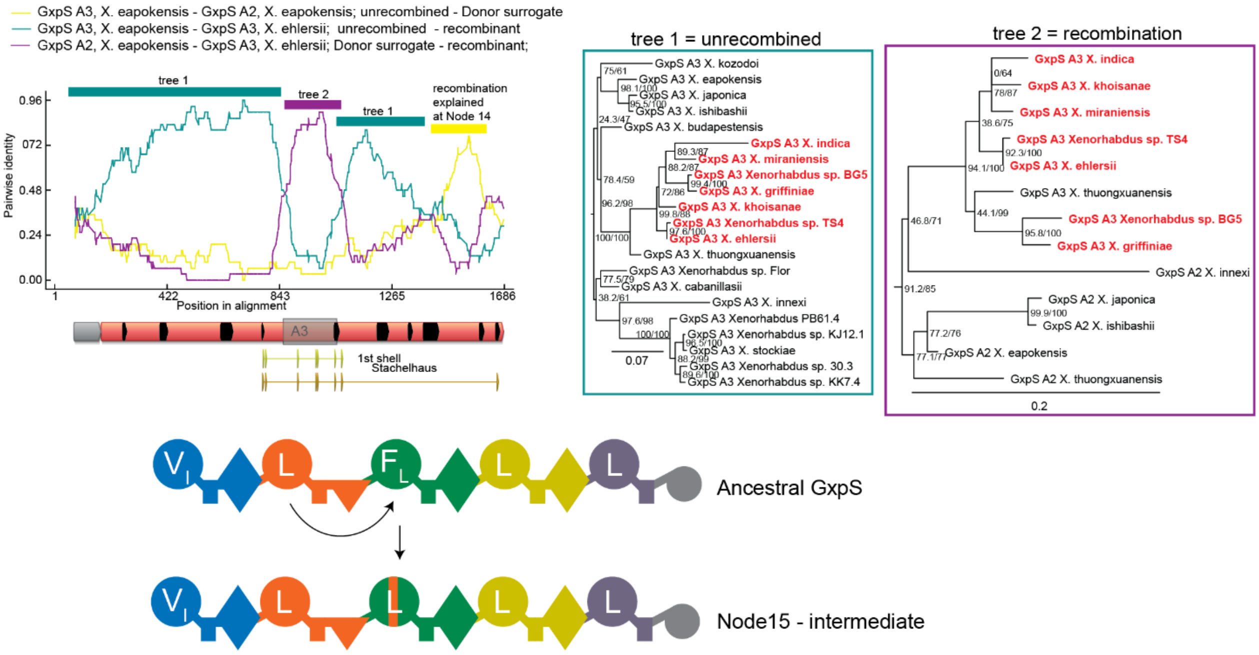


**Figure S83. Evolution of GxpS at Node 15 - part 1.** This recombination was discovered through the incongruent history of A3 domains and the PhyML_Multi analysis in Supplementary Fig. 8. RDP analysis and tree inference as before, tree 1 was inferred using amino acids, tree 2 using nucleotides. In this case a small part of a leucine encoding domain overwrote the third A-domain and changed its specificity from phenylalanine to leucine. The yellow peak in the RDP plot is the result of another recombination that is explained in Supplementary Fig. 82.

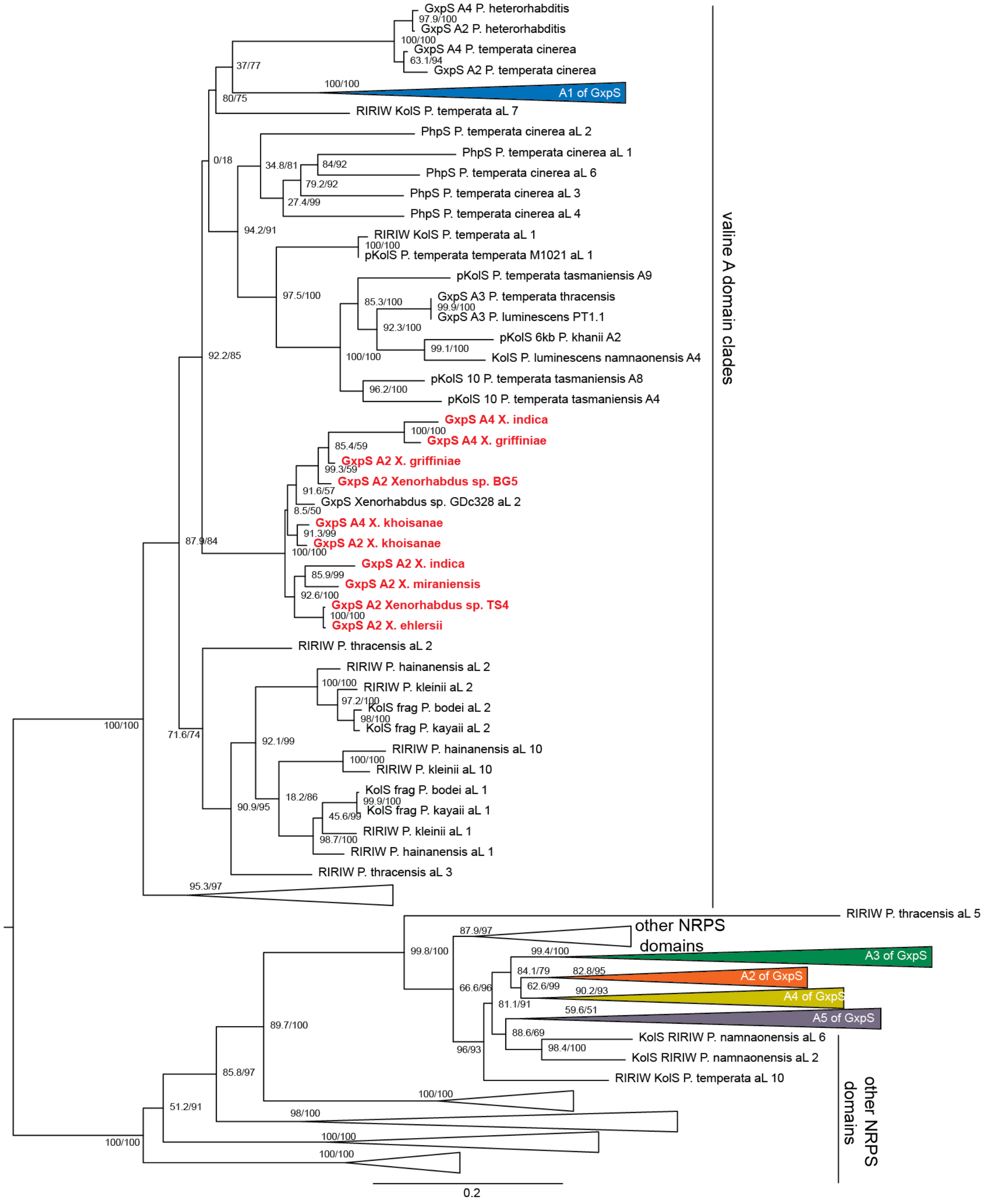


**Figure S84. Evolution of GxpS at Node 15 - part 2.** Discovery A2 and A4 domain recombinations of GxpS. A-domain protein tree of GxpS including A-domains from other synthetases. A2 domains of the GxpS from Node 15 do not sit in the second A-domain clade. Instead, they sit in the valine clade but outside of the GxpS clade, indicating they have recombined with another NRPS that pulled them towards the valine clade.

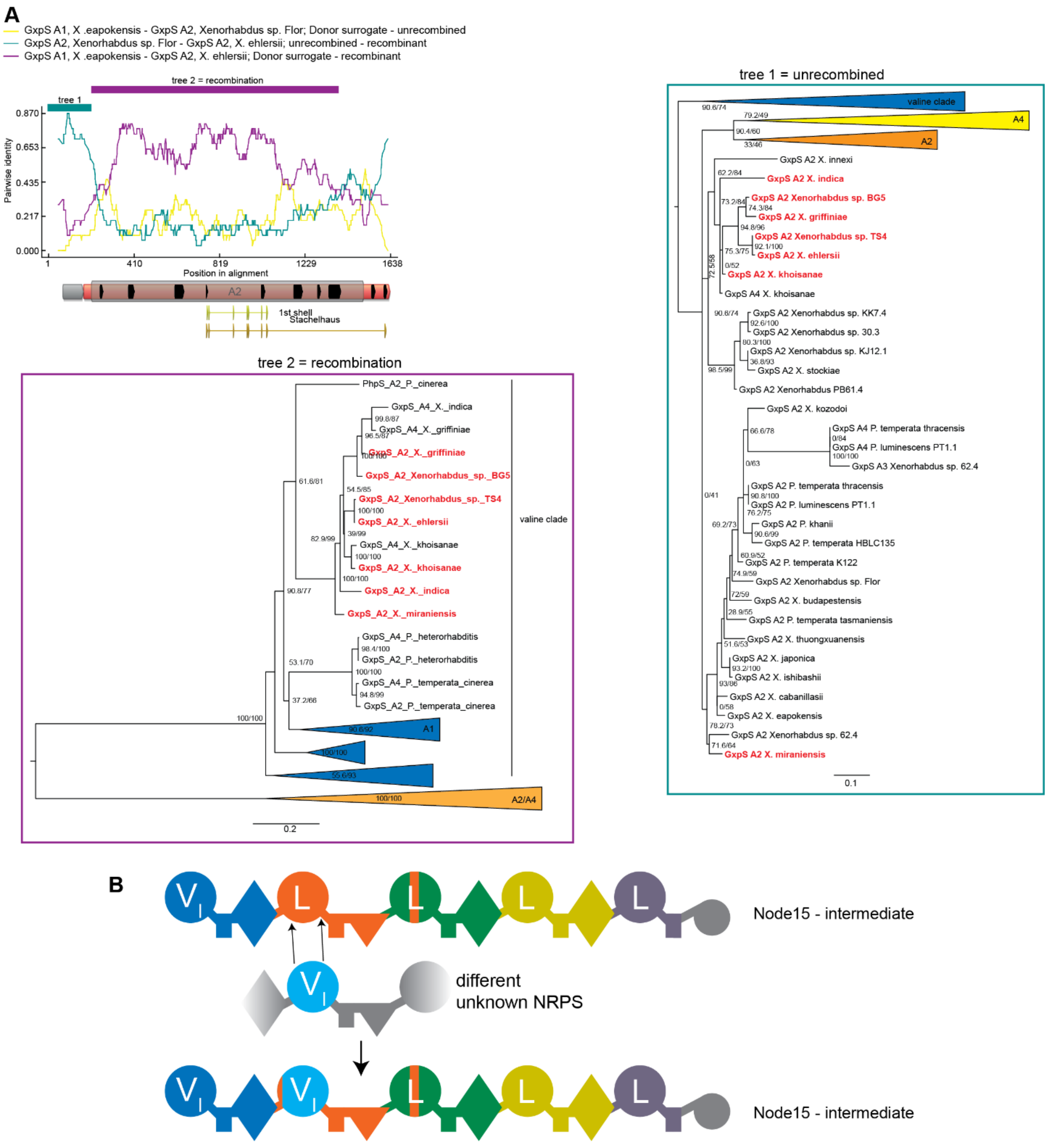


**Figure S85. Evolution of GxpS at Node 15 - part 3.** Analysis of the recombination event discovered in Supplementary Fig. 84. RDP analysis and tree inference as above, both trees were inferred using nucleotides. The recombined tree shows A2 domains clading inside the valine clade but outside the GxpS A-domains, indicating recombination with A-domain from a different NRPS.

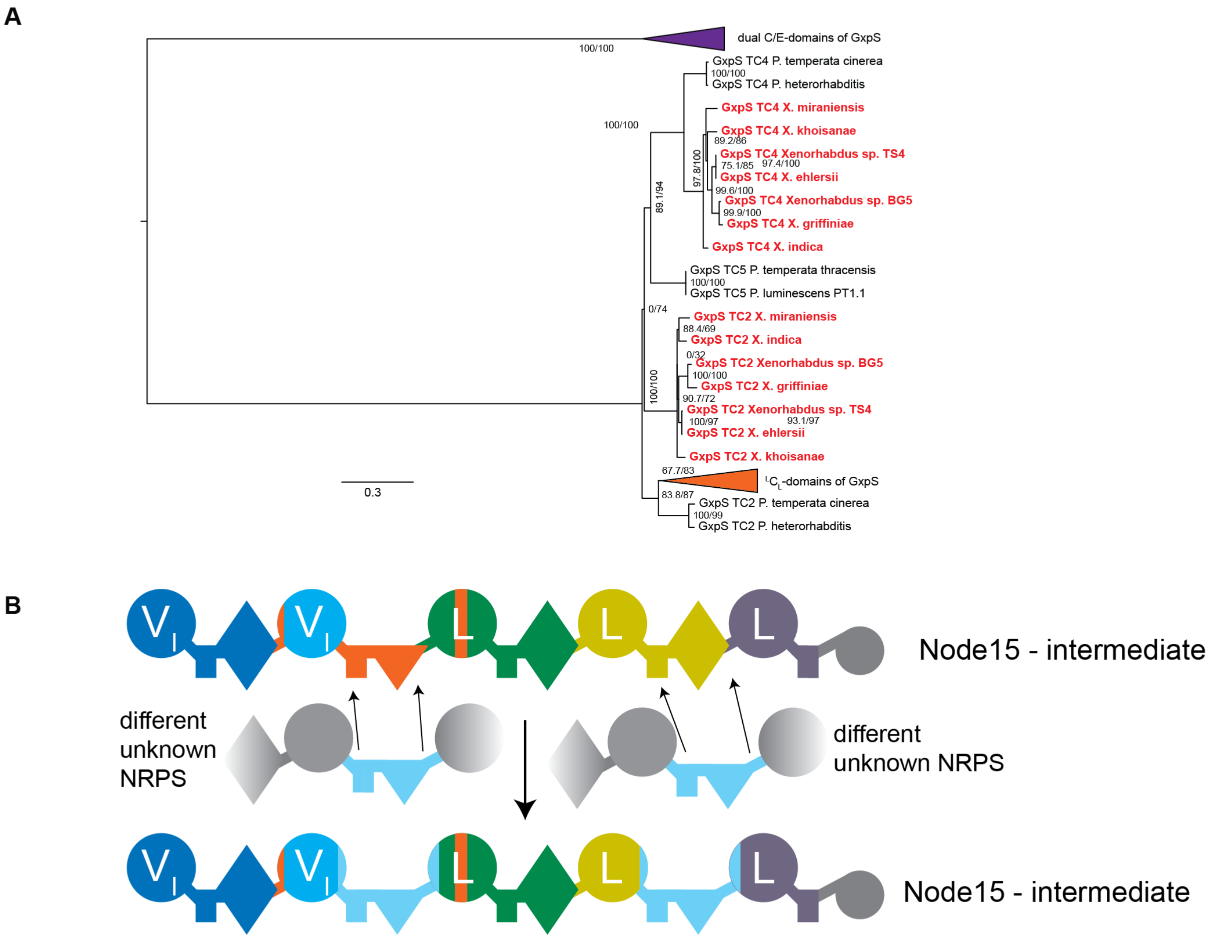


**Figure S86. Evolution of GxpS at Node 15 part 4. (A)** Evidence for a recombination of C-domains on our TC-domain protein tree of GxpS. This tree shows that the TC2- and TC4-domains from several GxpS sit outside of the GxpS clade hinting recombination with another NRPS. TC4 in other GxpS is a dual E/C-domain, so this recombination was non-synonymous. TC2 in other GxpS is an $^{L}C_{L}$, so this recombination was synonymous. We could not identify the breakpoints of these recopmbinations, because they lie outside of the C-domains, possibly in their flanking A-domains. It is possible that these recombinations occurred independently. Alternatively, only one of them recived a fragment from another NRPS. The other C-domain could then have been overwritten within GxpS by the first already-recombined C-domain. The C4 recombination at this node probably allowed early cyclization in the affected GxpS (see also Extended Data Fig. 6). **(B)** Cartoon diagram of these recombinations, where the two C-domain recombinations are arbitrarily shown as independent.

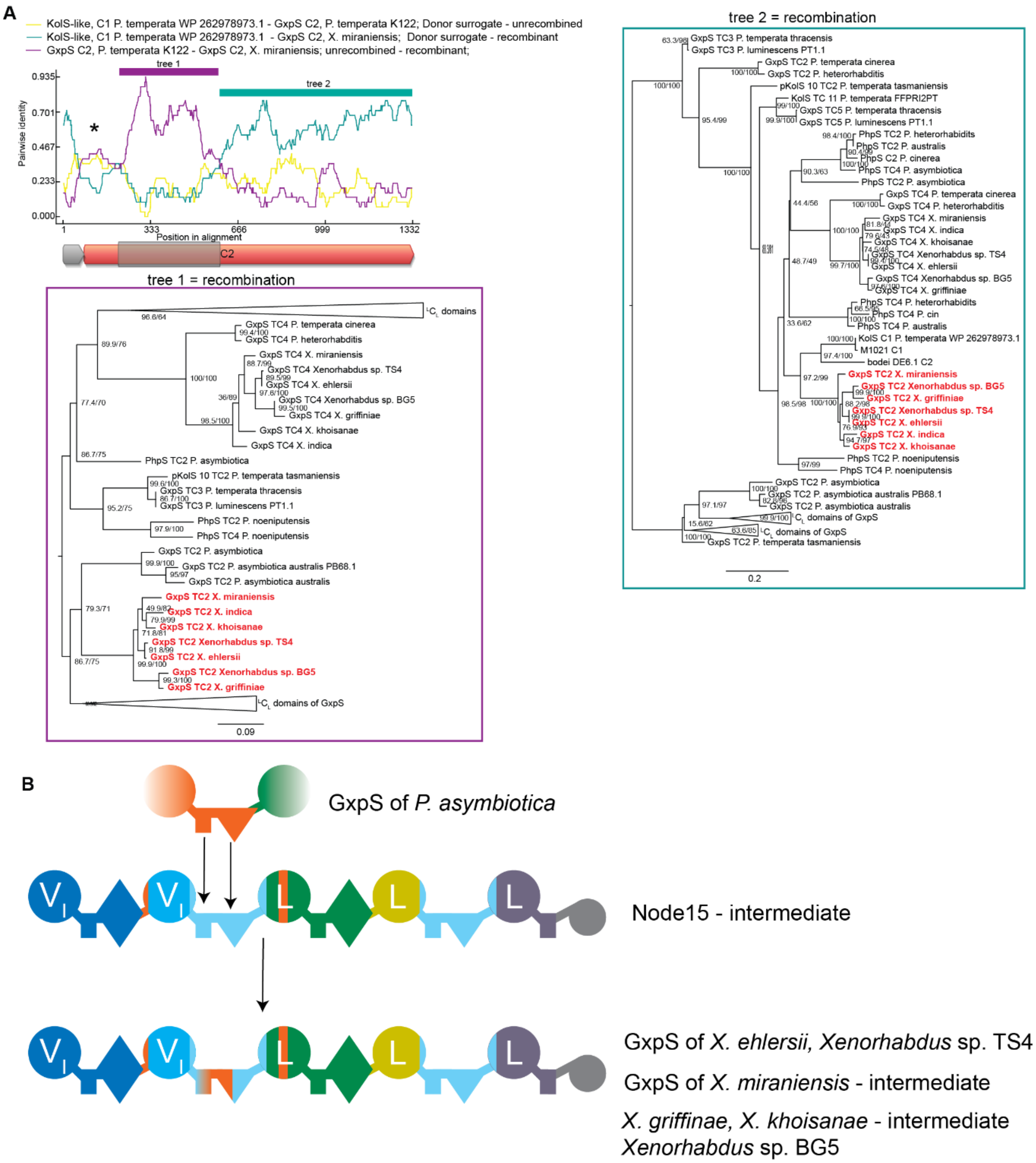


**Figure S87. Evolution of GxpS at Node 15 part 5.** This recombination was discovered using RDP. (**A**) RDP analysis and tree inference as before, trees were inferred using nucleotides. In this case, tree 2 represents the recombined history explained in Supplementary Fig. 86. Tree 1 represents a part of the sequences that underwent an additional recombination into the already recombined 2nd C-domain. The recombined sequences sit very close to TC2s that are far from them on the TTE-domain phylogeny, indicating recombination (**B**) About half of a C-domain recombined homotypically into C2.

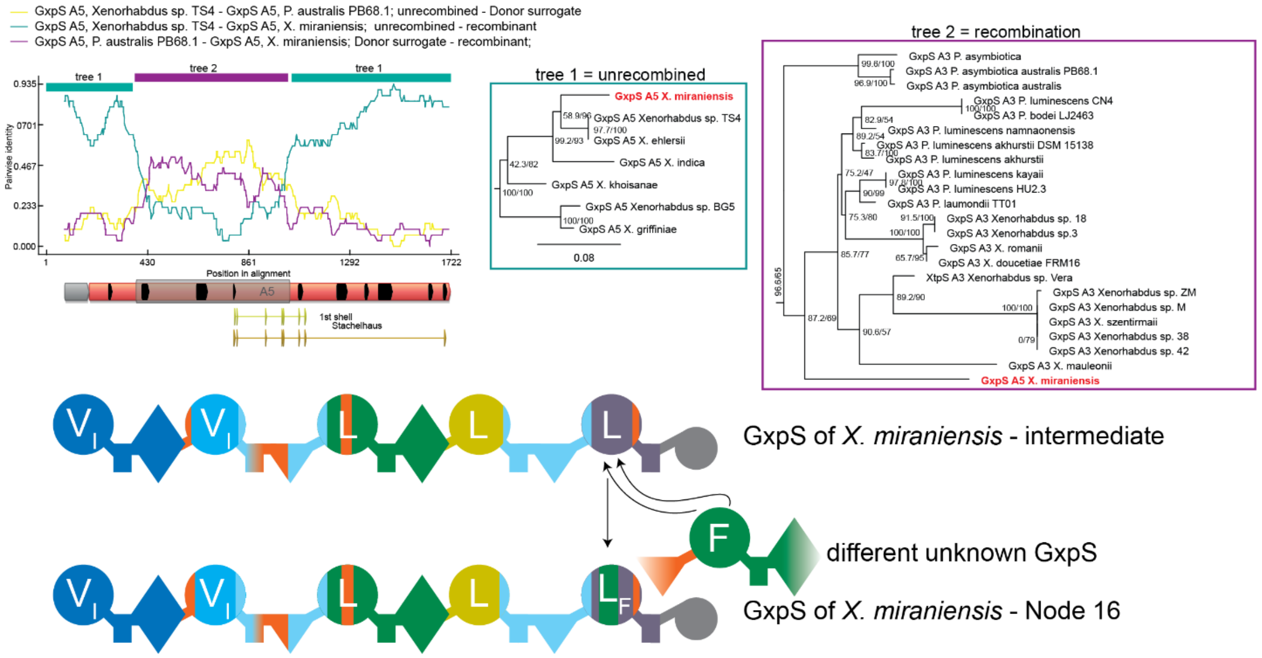


**Figure S88. Evolution of GxpS at Node 16.** This recombination was discovered through RDP. In this case, RDP does recognize a plausible recombination donor: It identifies the donor as another A5 domain. Tree 2, however, shows this sequence to clade inside A3 domains, which implies a non-synonymous change to phenylalanine. We tested this prediction experimentally and find that this synthetase has indeed acquired a side-specificity for phenylalanine in its fifth A-domain. The incorrect assignment of the likely donor by RDP probably arises because of the long branch to the recombined A5 domain (labelled red), which makes it too dissimilar to extant A-domains in our alignment to identify a likely donor.

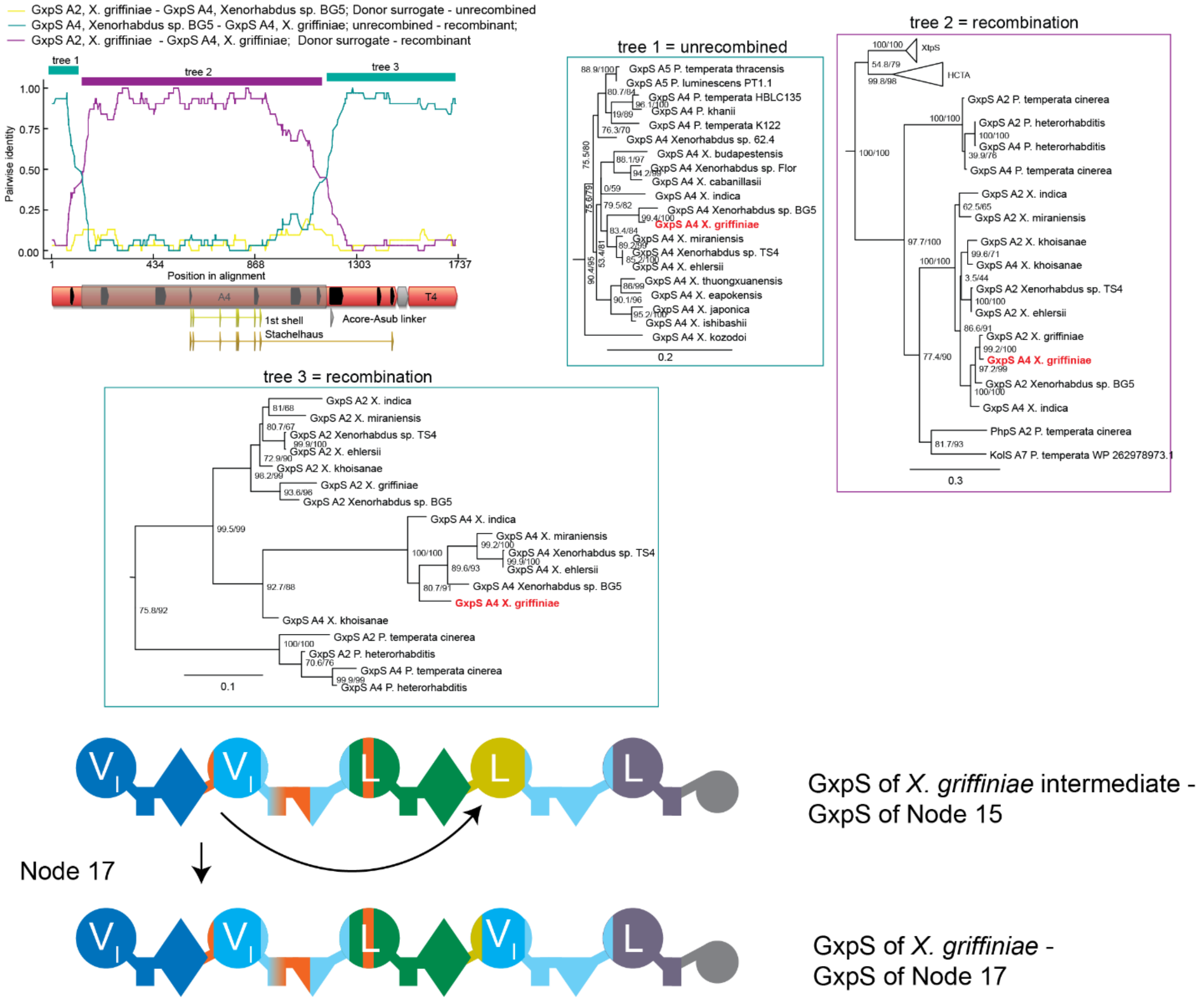


**Figure S89. Evolution of GxpS at Node 17.** This recombination was discovered through the anomalous clading of A4 domains in the valine clade in Supplementary Fig. 84. RDP analysis and tree inference as previous. Here we inferred three trees (all from nucleotides). The first tree is broadly congruent with the TTE-domain phylogeny and therefore this segment has not experienced recombination. The second tree shows that A2 overwrote A4 (highlightd in red). We inferred a third tree for the for the stretch following the recombined segement. This tree is not congruent with the TTE tree, probably because this segment contains sites towards the end of the A-domain (including the T-domain) that experienced the C-domain recombination shown in Supplementary Fig. 86. This results in A2 and A4 domains clading together.

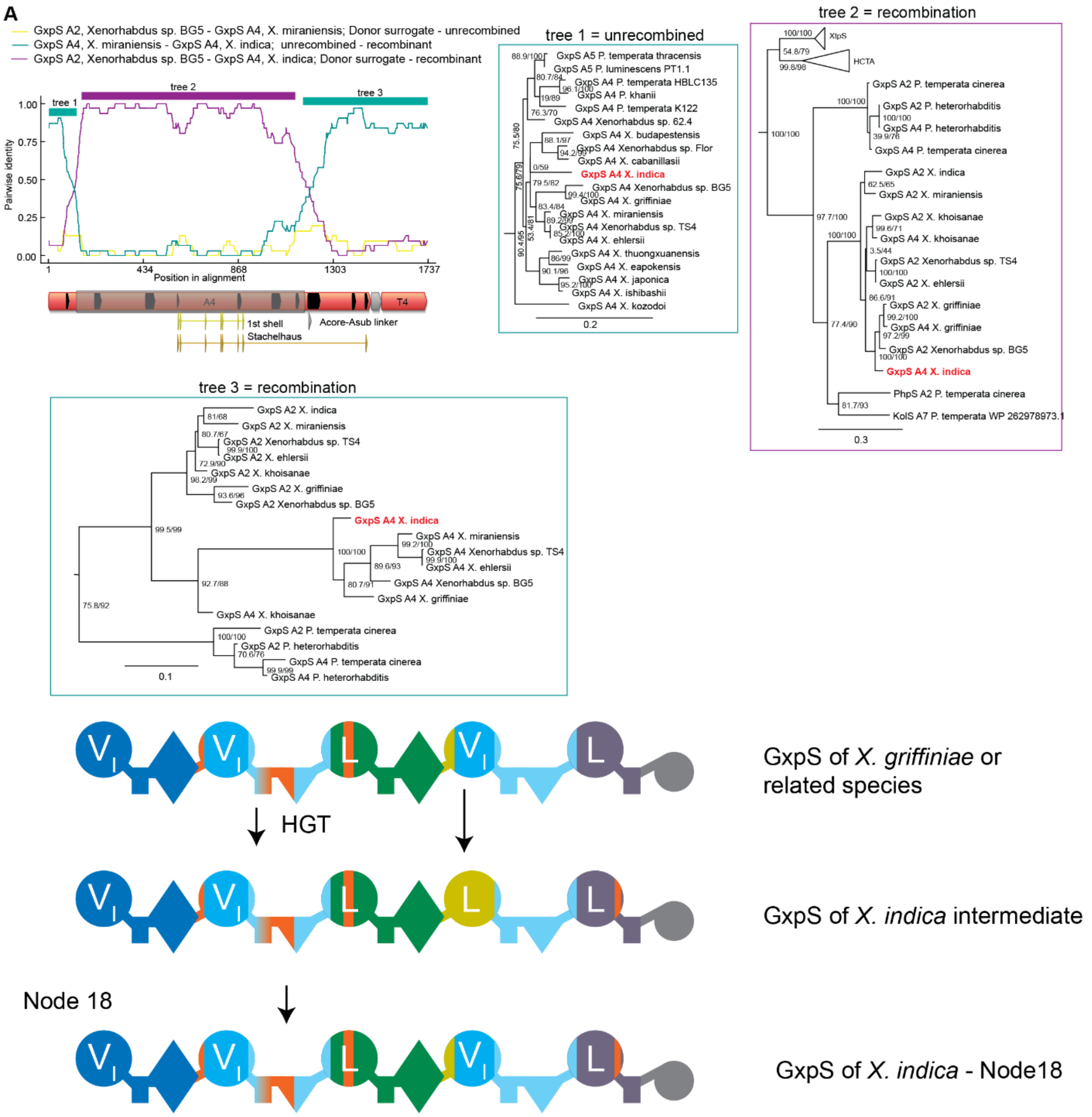


**Figure S90. Evolution of GxpS at Node 18.** This recombination event was discovered through the anomalous position of *X. indica*'s A4 domain in the A-domain phylogeny, as shown in Supplementary Fig. 84. RDP analysis as previous, the trees are identical to the trees in Supplementary Fig. 89. Tree 2 suggests that *X. indica*'s A4 was overwritten by a valine A-domain. The borders seem identical to the recombination in Supplementary Fig. 89. It is unlikely that these patterns were created by a single recombination event that was shared between the GxpS *X. griffinae* and *X. indica*, because the GxpS of these two species are never sister to each other on any of our trees. It seems more likely that *X. indica* received an already recombined valine A-domain (either A2 or A4) from a Xenorhabuds closely related to X *griffinae* or *Xenorhabdus* sp. BG5, probably with borders outside of the original recombination that we failed to detect.

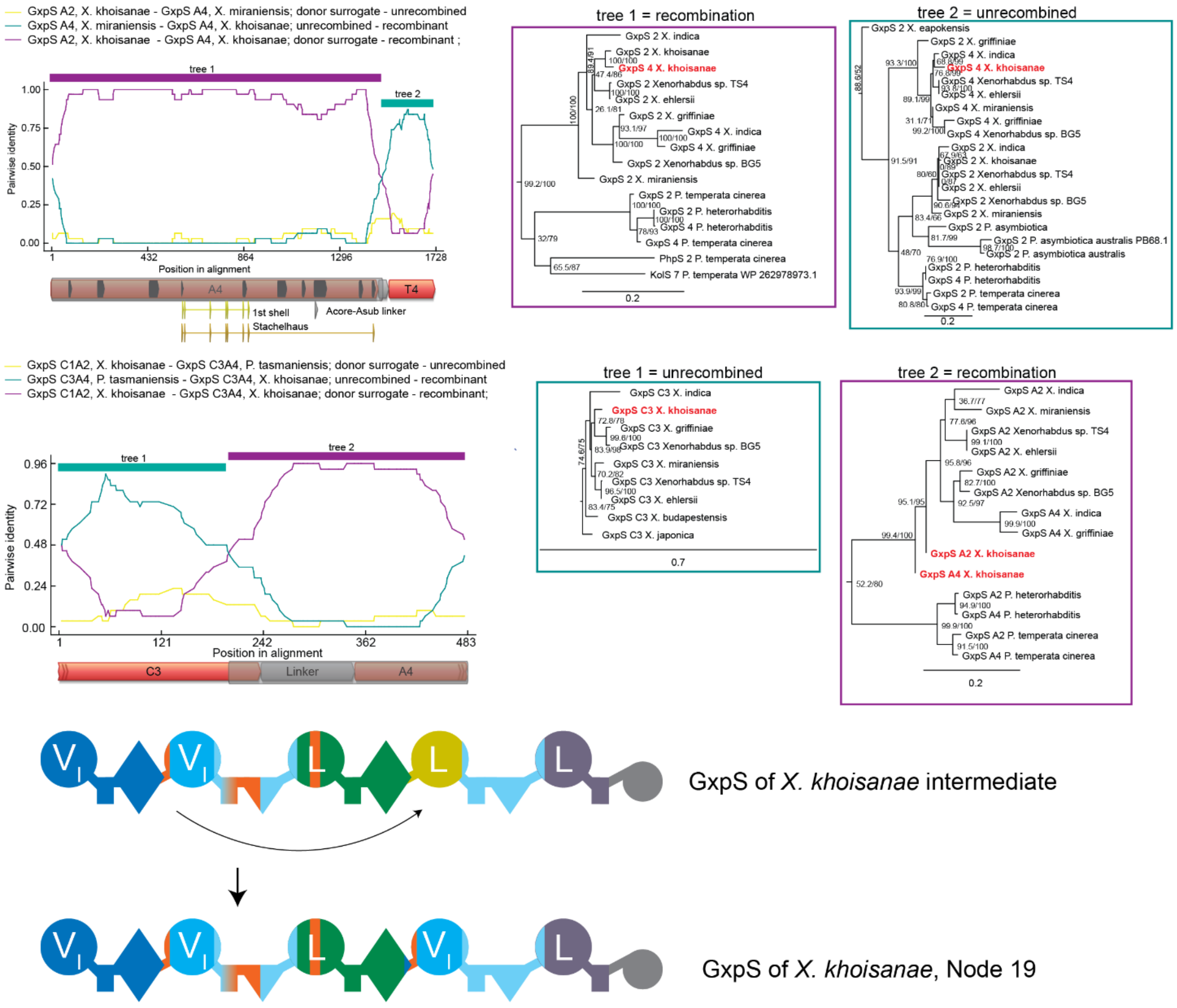


**Figure S91. Evolution of GxpS at Node 19.** This recombination event was discovered through the anomalous position of *X. khoisanae's* A4 domain in the A-domain phylogeny, as shown in Supplementary Fig. 84. (**A**) Identification of the right recombination breakpoint within the A-domain. RDP and tree analysis as previous, trees were inferred using nucleotides. Tree 2 shows that *X. khoisanae's* T4 domain sits with a larger group of A4 domains from related synthetases, which together sit inside the T2 domain clade. This is the result of the C-domain recombination explained in Supplementary Fig. 86. **(B**) Identification of the right recombination breakpoint within the A-domain. RDP and tree analysis as previous, trees were inferred using nucleotides. Overall, this recombination event replaced a leucine A-domain with a valine A-domain independently from two similar events (resulting in the same natural product) shown in Supplementary Fig. 89 and 90. We infer these events to be separate, because they use different recombination borders.

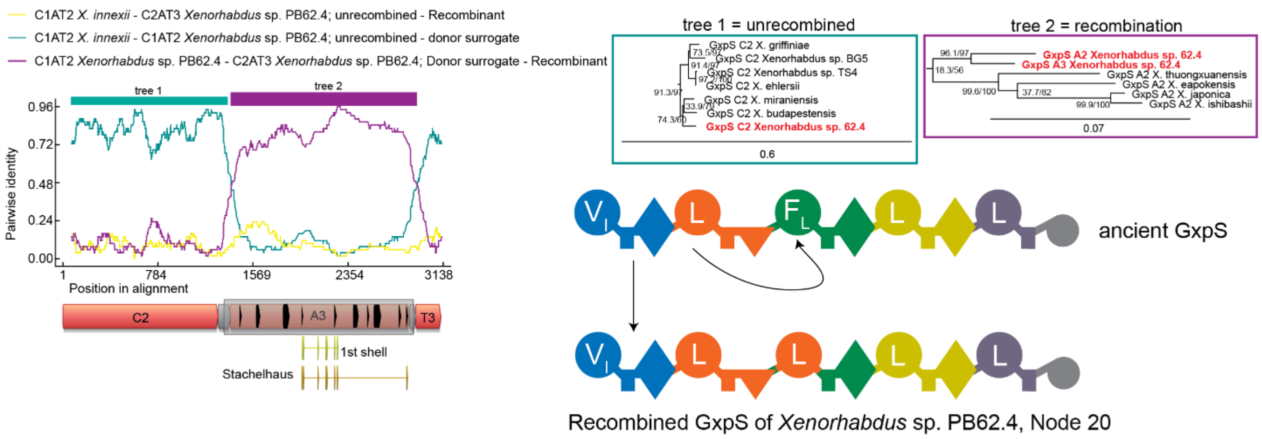


**Figure S92. Evolution of GxpS at Node 20.** This recombination was discovered through the anomalous position of the A3 domain of *Xenorhabdus* sp. PB62.4. RDP and tree analysis as previously, trees were inferred from amino acids. Here the A3 domain was overwritten by a leucine A-domain from the same synthetase.

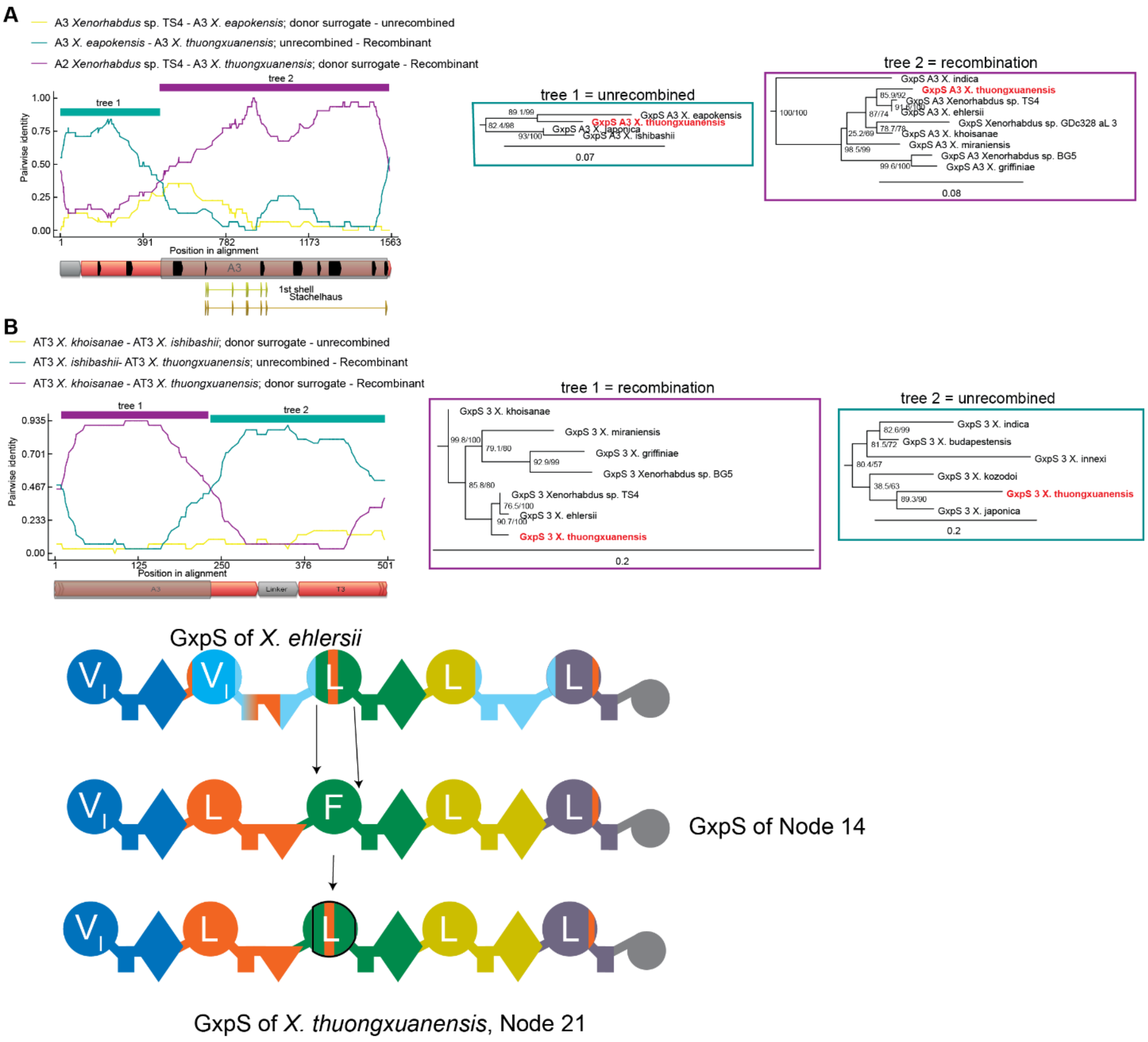


**Figure S93. Evolution of GxpS at Node 21.** This recombination was discovered through the incongruent topology of the A3 domain phylogeny (Supplementary Fig. 8). (**A**) Identification of the left recombination border. RDP analysis and tree inference as previous, trees were inferred from amino acids. (**B**) Identification of the right recombination border using an alignment that uses a smaller part of the A-domain but extends into the T-domain. RDP analysis and tree inference as before, trees were inferred using nucleotide sequences. Here a previously recombined A3 domain that was already specific for leucine overwrote a phenylalanine specific A3 domain.

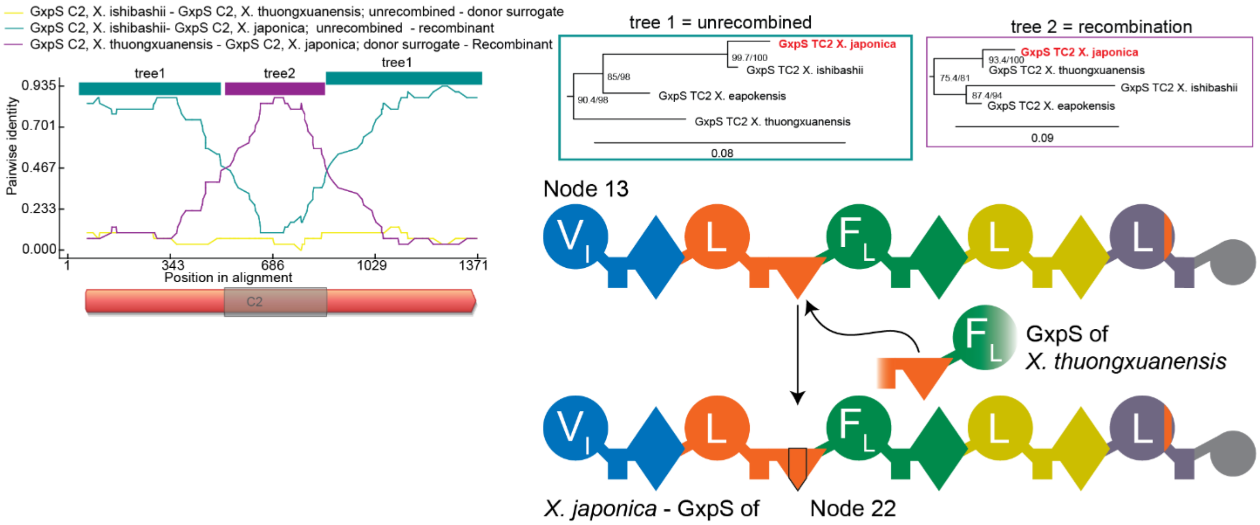


**Figure S94. Evolution of GxpS at Node 22.** This recombination was identified using RDP**.** RDP analysis and tree inference as previous, trees were inferred using protein sequences. Here tree 1 is congruent with the position of *X. japonica* in the TTE phylogeny. Tree 2 places it anomalously as sister to the TC2 domain of a closely related synthetase. We infer a homotypic recombination between the C2 domains of two closely related synthetases.

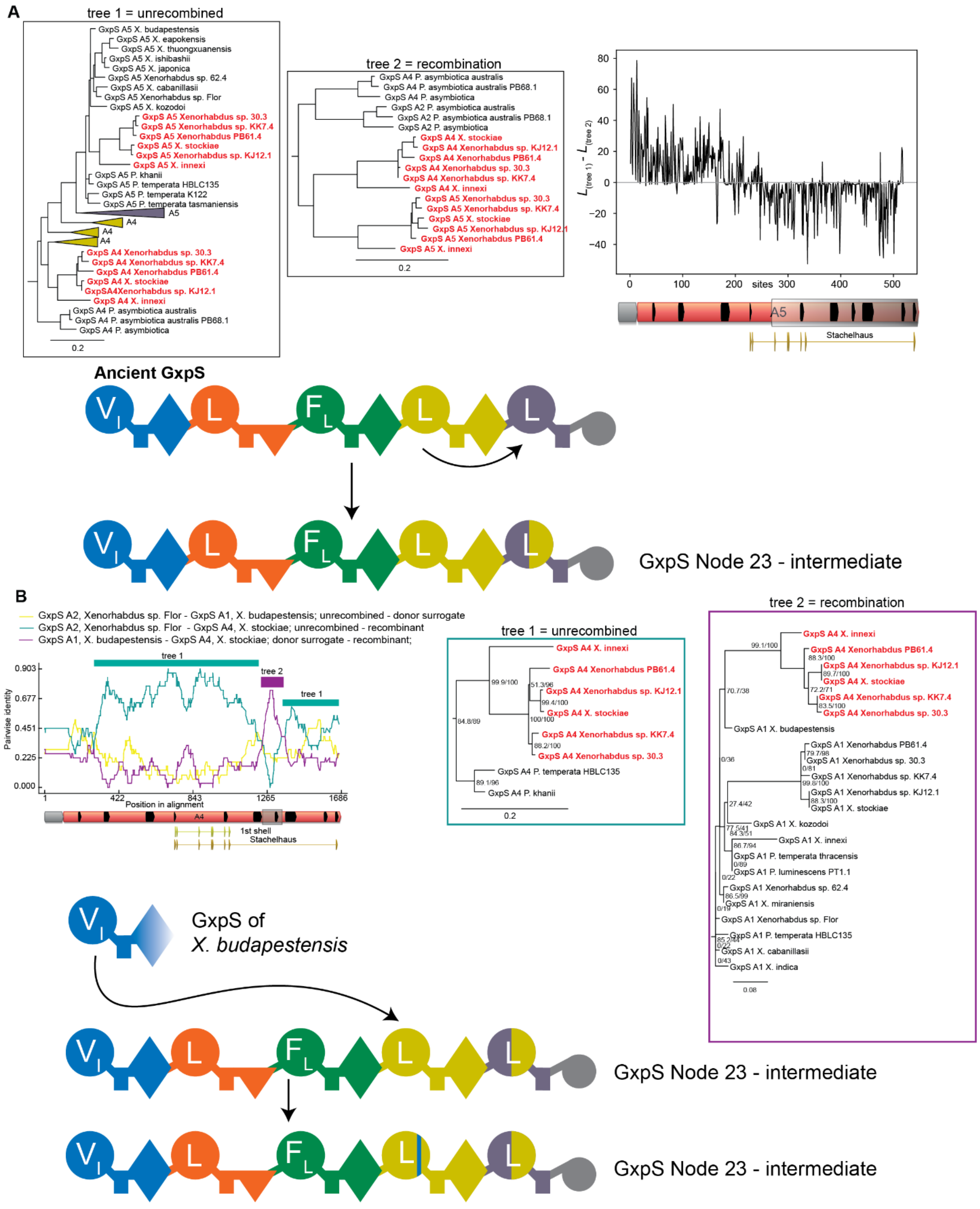


**Figure S95. Evolution of GxpS from Node 23 – part 1.** (**A**) The recombination was identified using a PhyML_Multi analysis of all A-domains from GxpS using amino acids for tree inference. This approach identifies the best two trees which together best describe the alignment. Left: tree 1 as identified by PhyML_Multi. This tree is broadly congruent with the TE-domain phylogeny. Middle: In the other tree inferred by PhyML_Multi, which places A5 domains highlighted in red as sister to the A4 domains of the same synthetases, indicated that A4 overwrote A5. Right: Difference between site-wise likelihoods across sites for the two-tree hypothesis. Sites with positive values favor tree 1, sites with negative values favor tree 2. This analysis was necessary, because RDP did not recognize this relatively deep recombination on the level of nucleotides. The recombination border lies somewhere near the middle of the A-domain but was not clearly identifiable using this approach. (**B**) Another recombination affecting the same set of synthetases as in (A), which we identified using RDP. RDP analysis and tree inference as previous, tree 1 was inferred using amino acids, tree 2 using nucleotides. We infer a small, heterotypic and synonymous recombination between valine A-domain that overwrote a small part of a leucine A-domain. In tree 2 the recombined A4 domains are not sister to the A1 domains from the same synthetases, but the support for this is low. We therefore think it plausible that the recombination occurred between the A1 and A4 domains within one ancestral GxpS (the ancestor of the species highlighted in red).

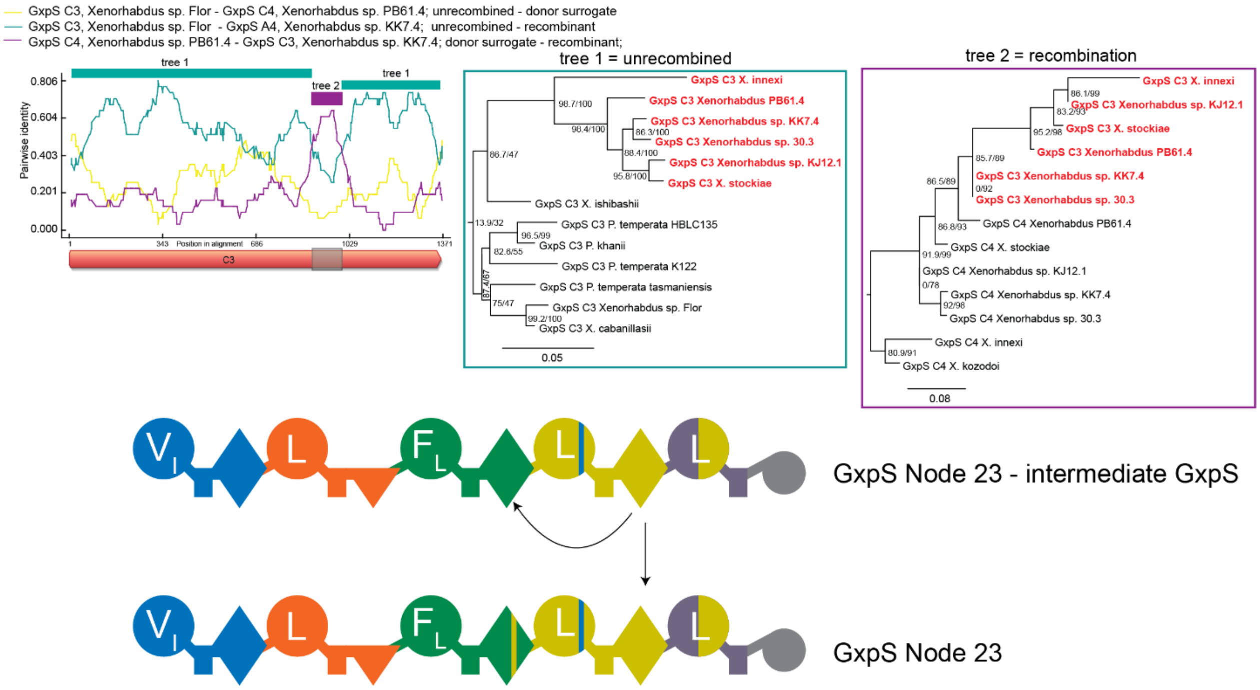


**Figure S96. Evolution of GxpS from Node 23 – part 2.** This recombination was discovered using RDP. RDP analysis and tree inference as previous, tree 1 was inferred using amino acids, tree 2 using nucleotides. Here a very small fragment of the C4 domain overwrote part of the C3 domain in a homotypic recombination.

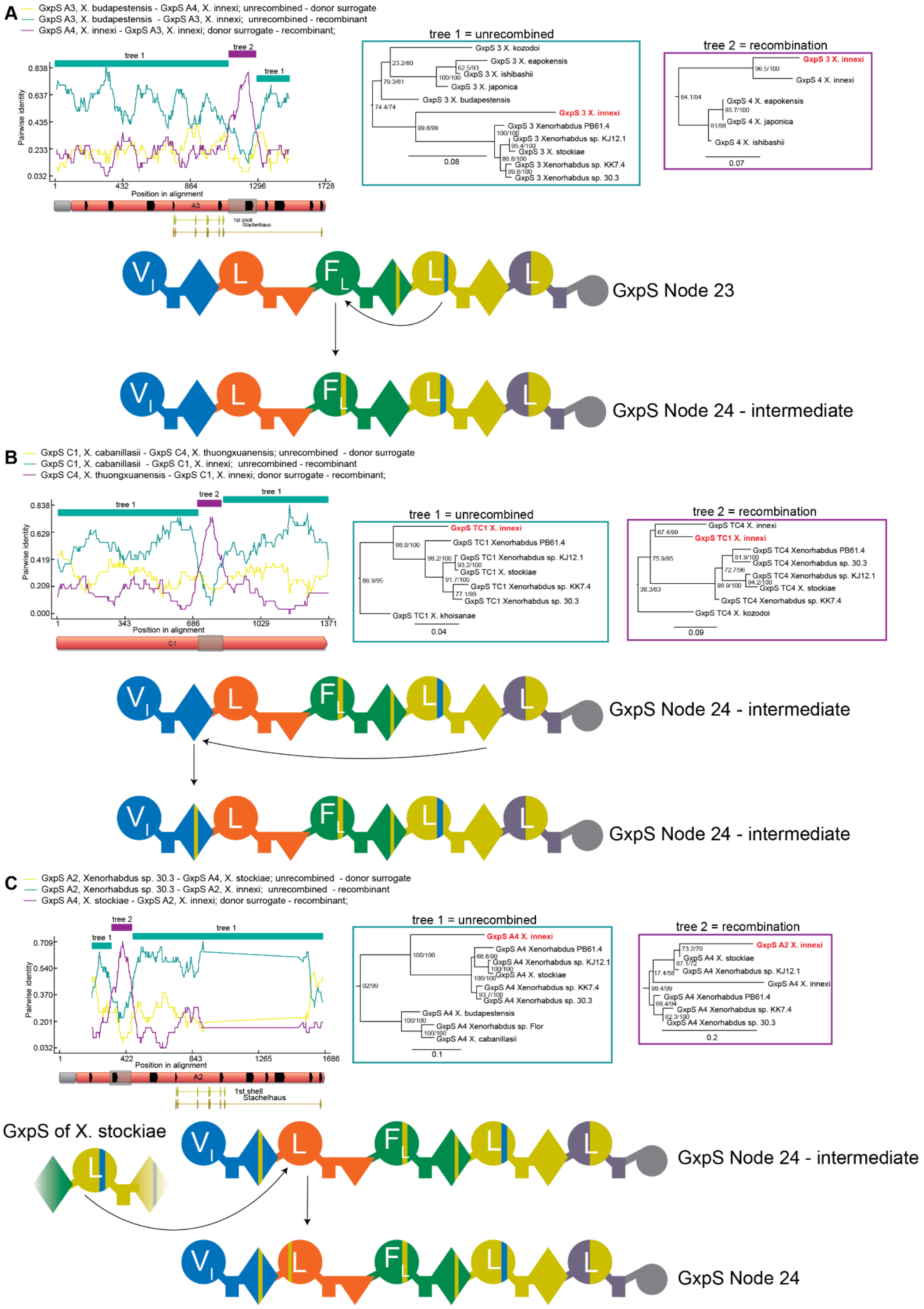

A
GxpS A3, X. budapestensis - GxpS A4, X. innexi; unrecombined - donor surrogate
GxpS A3, X. budapestensis - GxpS A3, X. innexi; unrecombined - recombinant
GxpS A4, X. innexi - GxpS A3, X. innexi; donor surrogate - recombinant;
tree 1 = unrecombined
tree 2 = recombination
Pairwise identity
Position in alignment
GxpS Node 23
GxpS Node 24 - intermediate
B
GxpS C1, X. cabanillasii - GxpS C4, X. thuongxuanensis; unrecombined - donor surrogate
GxpS C1, X. cabanillasii - GxpS C1, X. innexi; unrecombined - recombinant
GxpS C4, X. thuongxuanensis - GxpS C1, X. innexi; donor surrogate - recombinant;
tree 1 = unrecombined
tree 2 = recombination
GxpS Node 24 - intermediate
GxpS Node 24 - intermediate
C
GxpS A2, Xenorhabdus sp. 30.3 - GxpS A4, X. stockiae; unrecombined - donor surrogate
GxpS A2, Xenorhabdus sp. 30.3 - GxpS A2, X. innexi; unrecombined - recombinant
GxpS A4, X. stockiae - GxpS A2, X. innexi; donor surrogate - recombinant;
tree 1 = unrecombined
tree 2 = recombination
GxpS of X. stockiae
GxpS Node 24 - intermediate
GxpS Node 24

**Figure S97. Evolution of GxpS from *X. innexi*.** All recombinations in this figure were identified using RDP (**A**) RDP analysis and tree inference for a heterotypic, synonymous recombination in which the A4 domain overwrote a small fragment of the A3 domain outside of the specificity area [14]. Tree 1 was inferred using amino acids, tree 2 using nucleotides. (**B**) RDP analysis and tree inference for a homotypic recombination in which the C4 domain overwrote a small fragment of the C1 domain. Tree 1 was inferred using amino acids, tree 2 using nucleotides. (**C**) RDP analysis and tree inference for a homotypic recombination in which a A4 domain (possibly from another species) overwrote a small fragment of the A2 domain. Tree 1 was inferred using amino acids, tree 2 using nucleotides.

.

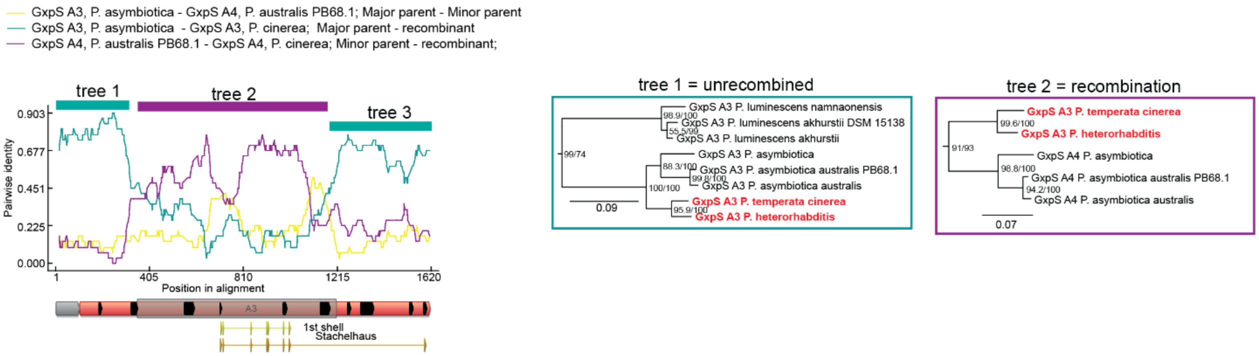


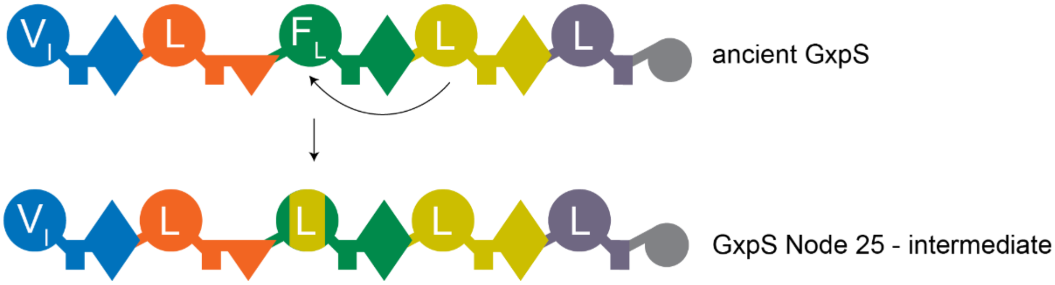


**Figure S98. Evolution of GxpS from Node 25 – part 1.** This recombination was identified through the incongruent topology of the A3 domain phylogeny. RDP analysis and tree inference as previous, trees were inferred using amino acids. Here the A3 domain from the two synthetases highlighted in red were overwritten in a non-synonymous, heterotypic recombination by a leucine-encoding A4 domain.

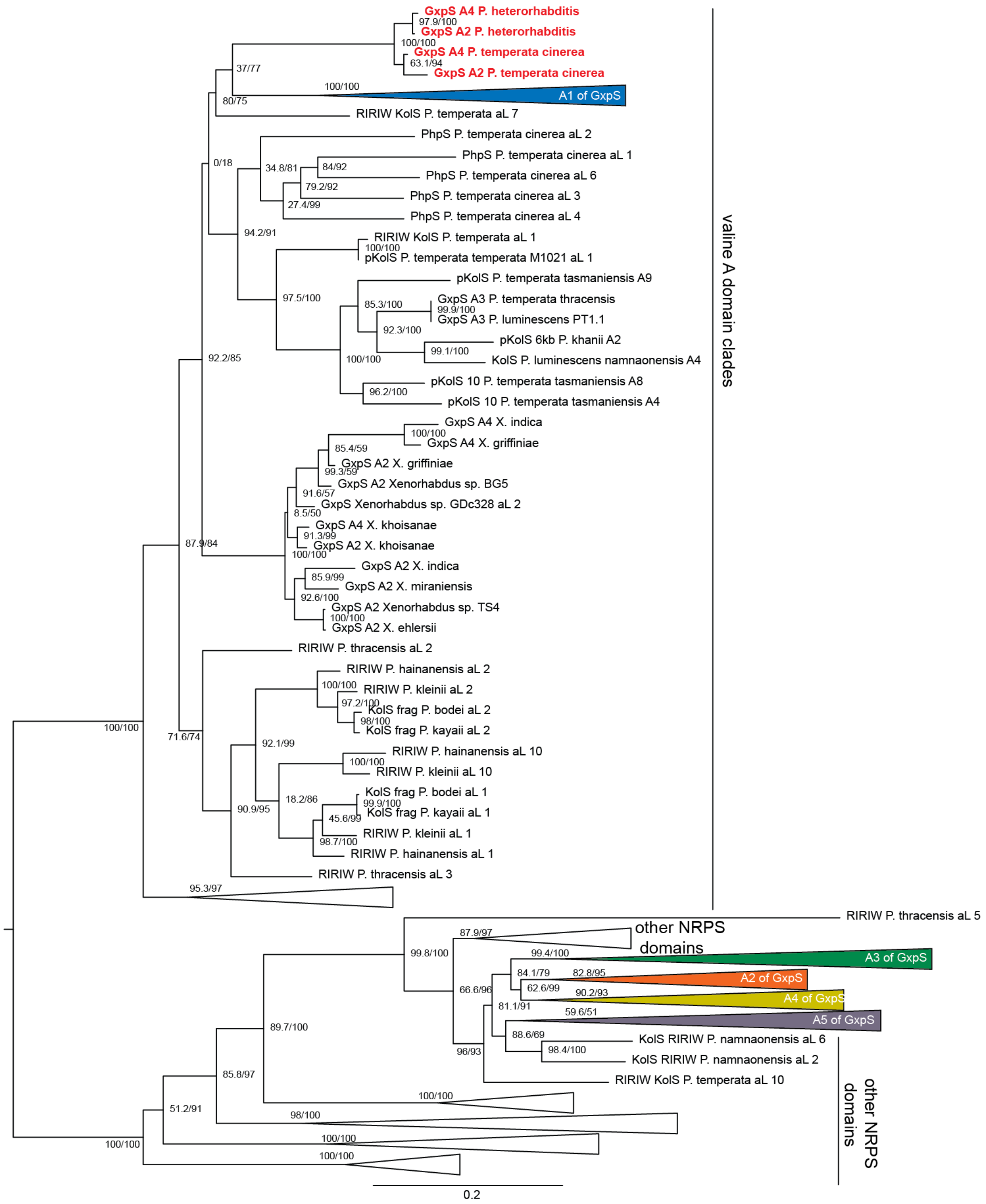


**Figure 99: GxpS-node 25 - part 2.** Identification of an A-domain recombination in the A-domain phylogeny of GxpS. An A-domain tree of GxpS with putative recombination partners was inferred with protein sequences. The second and fourth A-domain of the GxpS from P. *cinerea* and *P. heterorhabditis* do not sit with other A2 of GxpS. Instead, they sit outside of the GxpS clade inside the valine clade. This indicates that another NRPS had recombined with the A2 of the ancestral GxpS that existed at node 25.

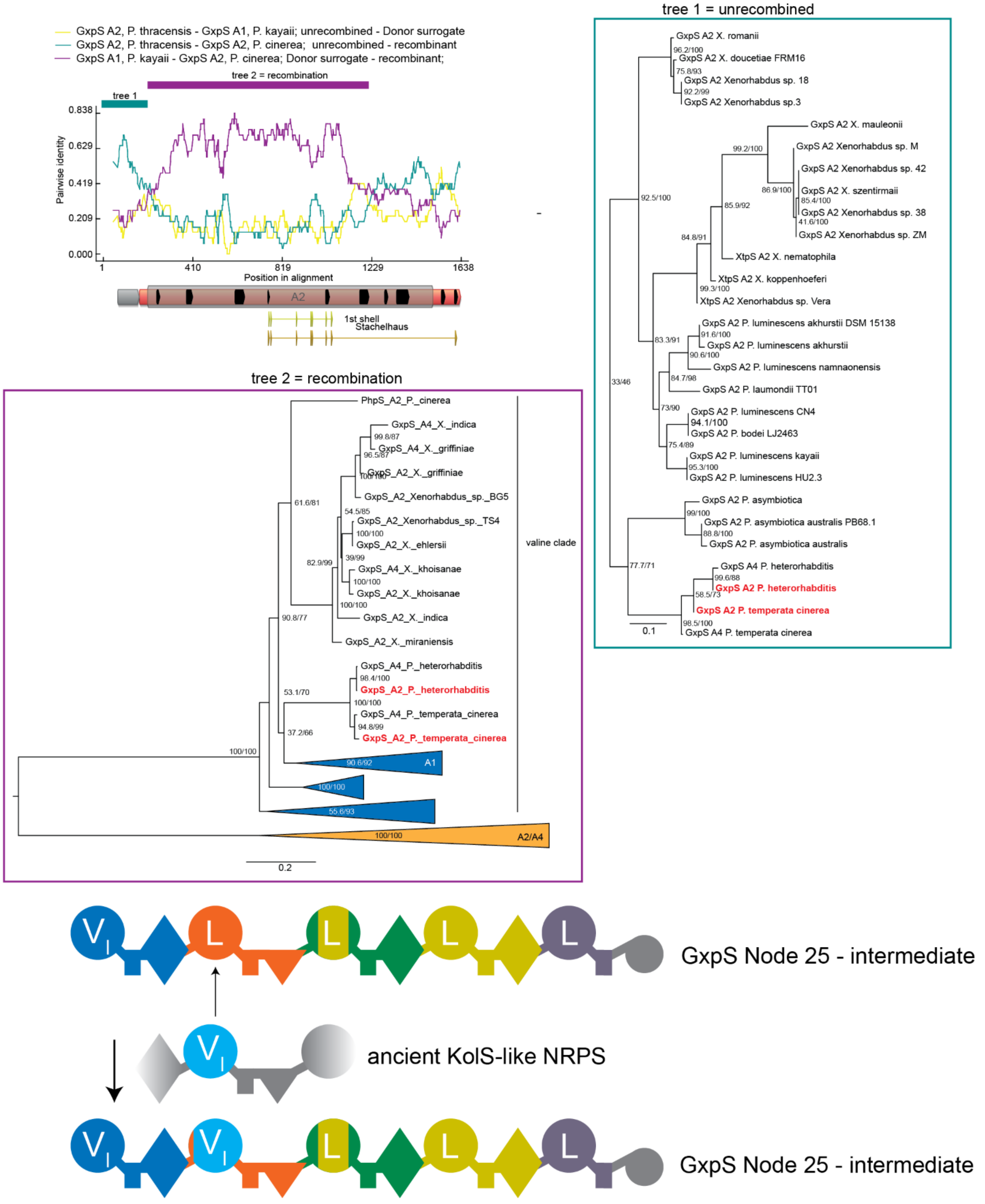


**Figure S100. Evolution of GxpS from Node 25 – part 3.** This recombination was identified through the anomalous clading of A2 domains in the valine clade shown in Supplementary Fig. 99. RDP analysis and tree inference as previous, trees were inferred using nucleotides. In tree 1, A2 domains highlighted in red sit within the A2 clade, consistent with no recombination. Next to them are A4 domains from the same synthetases, which are pulled into the A2 clade through a different recombination event explained in Supplementary Fig. 86. In tree 2, A2 domains highlighted in red sit in the valine clade. Here, a valine A-domain from another NRPS overwrote the A2 domain of the ancestor of *P. heterorhabditis* and *P. cinerea*.

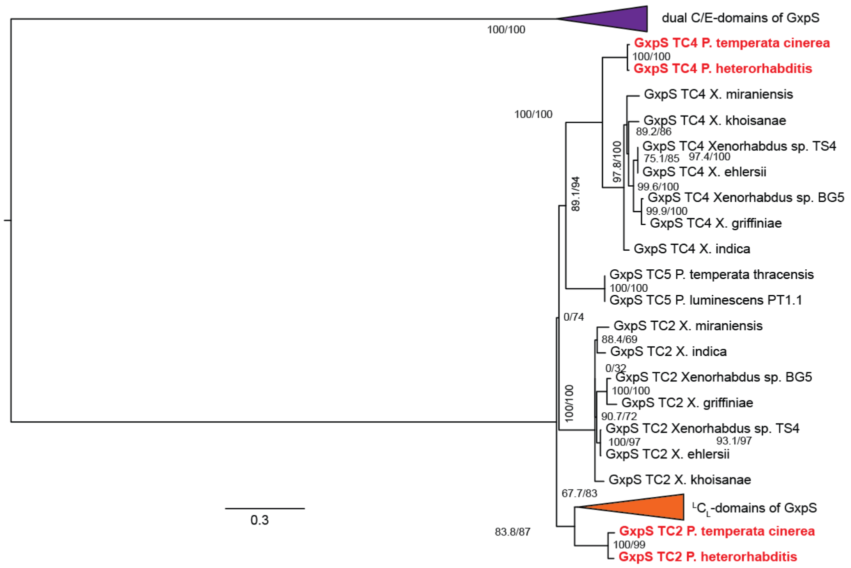


**Figure S101. Evolution of GxpS from Node 25 – part 4.** A TC-domain protein tree from all GxpS. This tree shows obvious C-domain recombinations. The TC2 and TC4 domain of the GxpS from Node 25 sits outside of the GxpS clade which indicates recombination with another NRPS.

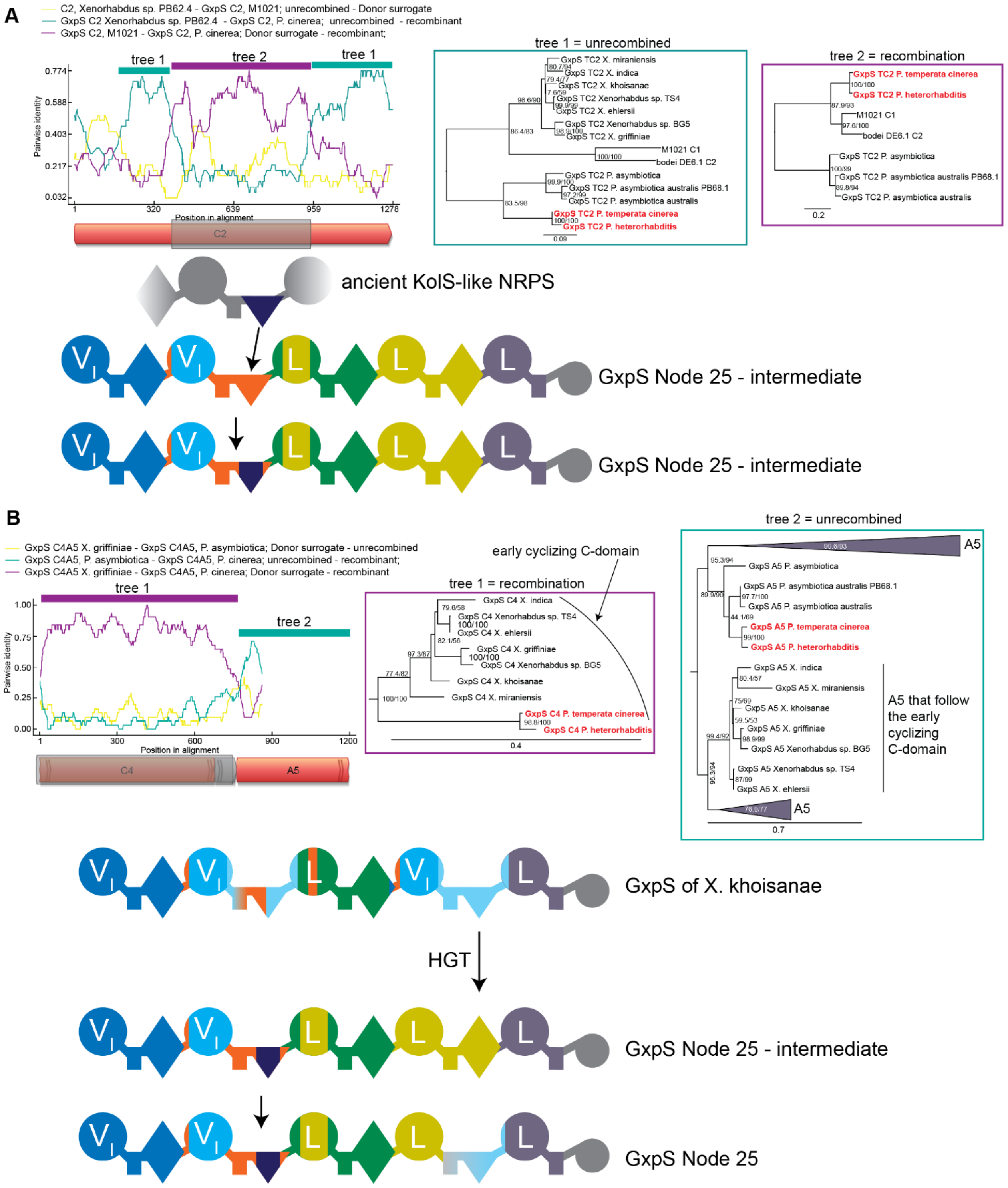


**Figure 102: Evolution of GxpS from Node 25 – part 5. (A)** This recombination was discovered through the anomalous position of the GxpS C2 domains of *P. cinerea* and *P. heterorhabditis* on the C-domain phylogeny in Supplementary Fig. 101. RDP analysis and tree inference as before. Trees were inferred using amino acids. Here the C2 domain was overwritten synonymously by another $^{L}C_{L}$ from an NRPS outside the GxpS clade. This recombination event is superficially similar to the one described in Supplementary Fig. 87 but clearly uses different borders. (**B**) This recombination was

discovered through the anomalous position of the GxpS C4 domains of *P. cinerea* and *P. heterorhabditis* on the C-domain phylogeny in Supplementary Fig. 101. In this case the recombined synthetases (highlighted in red) have had their C4 domain overwritten by an $^{L}C_{L}$ from the group of GxpS capable of early cyclization with their $4^{th}$ C-domain. Consequently, the GxpS from *P. cinerea* and *P. heterorhabditis* should also be capable of early cyclization. We verified this experimentally for the GxpS from *P. cinerea* experimentally. It is extremely implausible that the recombination described in this figure and the one that installed an early cyclizing C4 domain in *Xenorhabdus* GxpS derive from a single event, because these two groups of synthetases are not close to each other on the TTE phylogeny. A single C4 recombination event in their last common ancestor would be very difficult to explain, because most other synthetases that also descend from this ancestor did not experience this exchange of the C4 domain. A much more parsimonious explanation is that the early cyclization C4 domain was first acquired either by the *Xenorhabdus* GxpS and then transferred to the ancestors of *P. cinerea* and *P. heterorhabditis* GxpS. Alternatively, the *P. cinerea* and *P. heterorhabditis* GxpS may have acquired this domain first and transferred it to the last common ancestor of the early cyclizing *Xenorhabdus* GxpS.

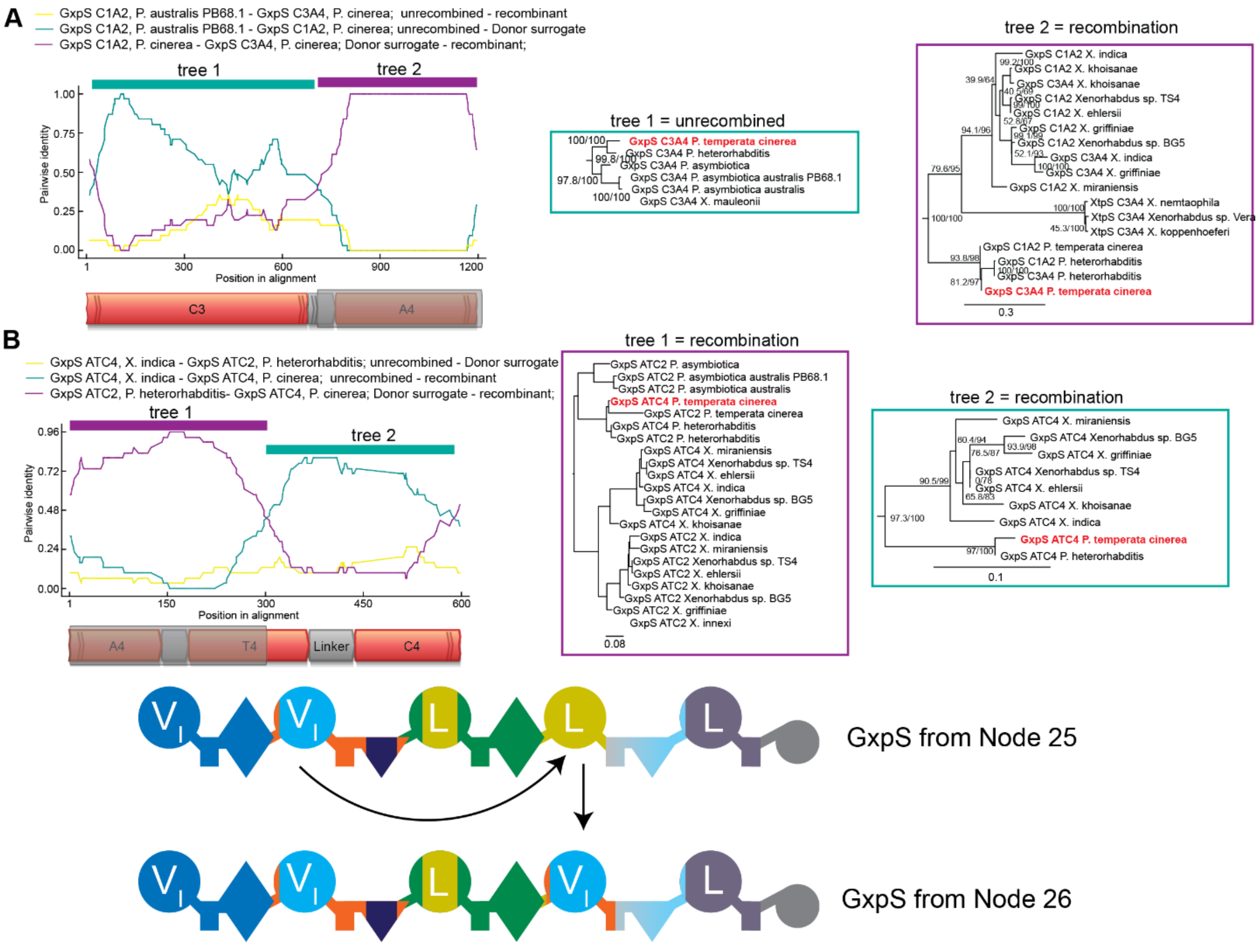


**Figure S103. Evolution of GxpS from Node 26.** This recombination was discovered through the anomalous position of the GxpS A4 domains of *P. cinerea* and *P. heterorhabditis* on the A-domain phylogeny in Supplementary Fig. 99. RDP analysis and tree inference as previous, trees were inferred using nucleotides. Here the A2 domain overwrote the A4 domain and changed the specificity from leucine to valine. (**A**) Identification of the left breakpoint, which appears to be somewhere in the T-domain. On tree 1, the synthetase in red groups with other synthetases according to its linear position in the sequence of GxpS and in agreement with the TTE-domain topology. In tree 2, this sequence is very close to its own C1A2 domain, indicating a very recent recombination. A similar pattern is true for the C3A4 sequence from *P. heterorhabditis*, which appears to have independently undergone the same recombination (sequences highlighted in bold). This is described in Supplementary Fig. 101. (**B**) Identification of the right breakpoint. Here tree 1 shows the same two recent recombinations in *P. cinerea* and *P. heterorhabditis* as tree 2 in (A), with the C3A4 of either species sister to the C1A2 sequence of the GxpS from same species. In tree 2, the position of the ATC4 sequence is affected by the C-domain recombination that *P. cinerea* and *P. heterorhabditis* have undergone, which is described in Supplementary Fig. 101.

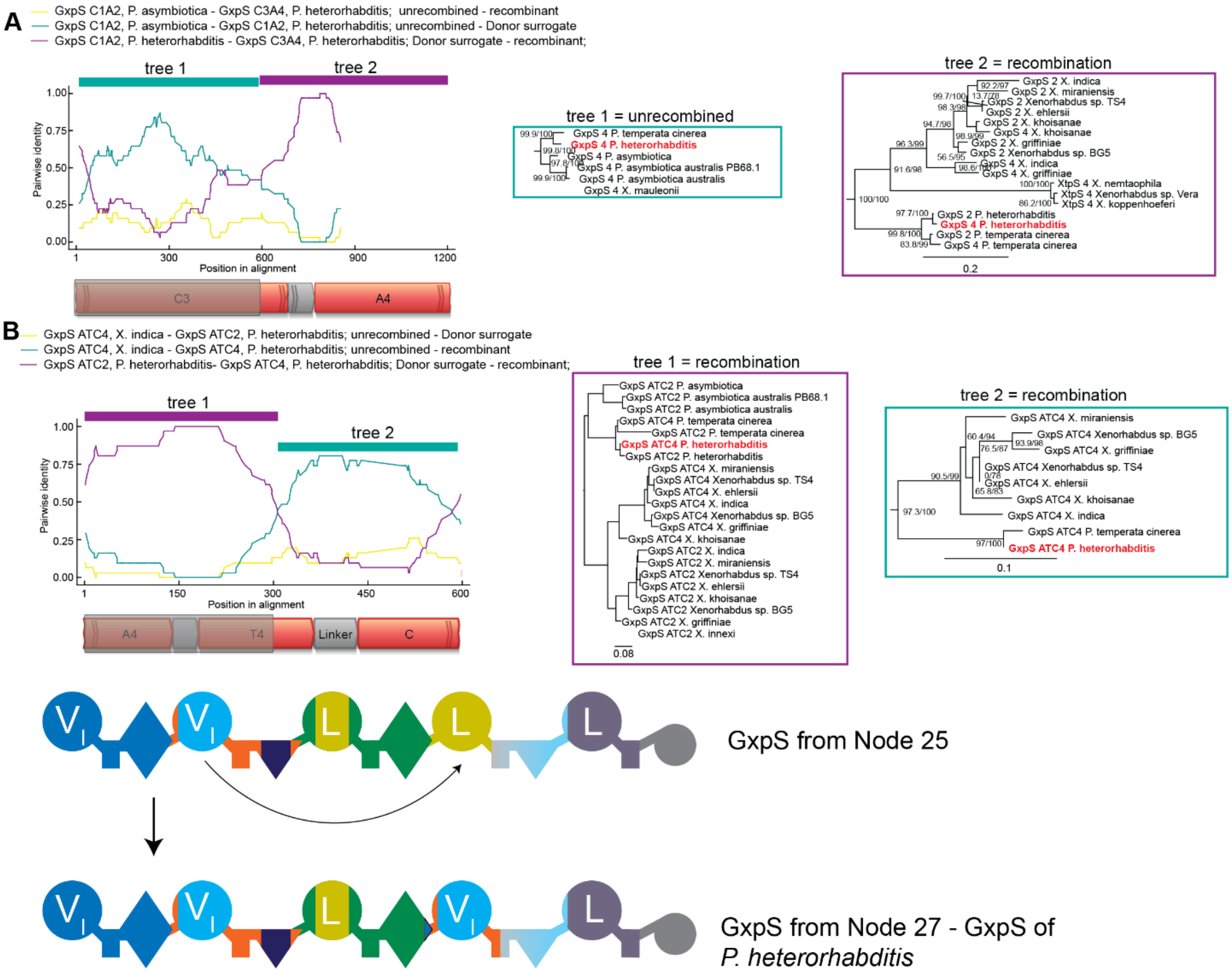


**Figure S104. Evolution of GxpS from Node 27.** This recombination was discovered through the anomalous position of the GxpS A4 domains of *P. cinerea* and *P. heterorhabditis* on the A-domain phylogeny in Supplementary Fig. 99. RDP analysis and tree inference as previous, trees were inferred using nucleotides. This is the same analysis as in Supplementary Fig. 103, but now with *P. heterorhabditis* as the focal synthetase instead of *P. cinerea*. (**A**) Identification of the left breakpoint, which is similar to the one identified in Supplementary Fig. 103. On tree 1, the synthetase in red groups with other synthetases according to its linear position in the sequence of GxpS and in agreement with the TTE-domain topology. In tree 2, this sequence is very close to its own C1A2 domain, indicating a very recent recombination. A similar pattern is true for the C3A4 sequence from *P. cinerea*, which appears to have independently undergone the same recombination (sequences highlighted in bold). This is described in Supplementary Fig. 103. (**B**) Identification of the right breakpoint, which is also similar to the one identified in Supplementary Fig. 103. Tree 1 again shows the same two recent recombinations in *P. cinerea* and *P. heterorhabditis* as tree 2 in (A), with the C3A4 of either species sister to the C1A2 sequence of the GxpS from same species. In tree 2, the position of the ATC4 sequence is affected by the C-domain recombination that *P. cinerea* and *P. heterorhabditis* have undergone, which is described in Supplementary Fig. 101. We interpret these data together with those in Supplementary Fig. 110 as two independent recombination events with very similar borders: in both cases the A2 domain overwrote the A4 domain, thereby introducing valine specificity. This scenario is supported by the fact that the A4 domain of *P. cinerea* is sister to its own A2 domain, and the same is true for the A4 domain from *P. heterorhabditis*. A single recombination event in the last common ancestor of *P. heterorhabditis* and *P. cinerea* should result in the A4 domain from *P. cinerea* as sister to the A4 domain of *P. heterorhabditis*. To explain the phylogenetic pattern, a single recombination event in their ancestor would require that in both species there were subsequent re-recombinations between the two valine A2 and A4 domains that would have evolved in their ancestor. We have no evidence for such a trajectory and therefore interpret our data as two independent recombinations between a valine A2 domain and a leucine A4 domain, as suggested by our trees.

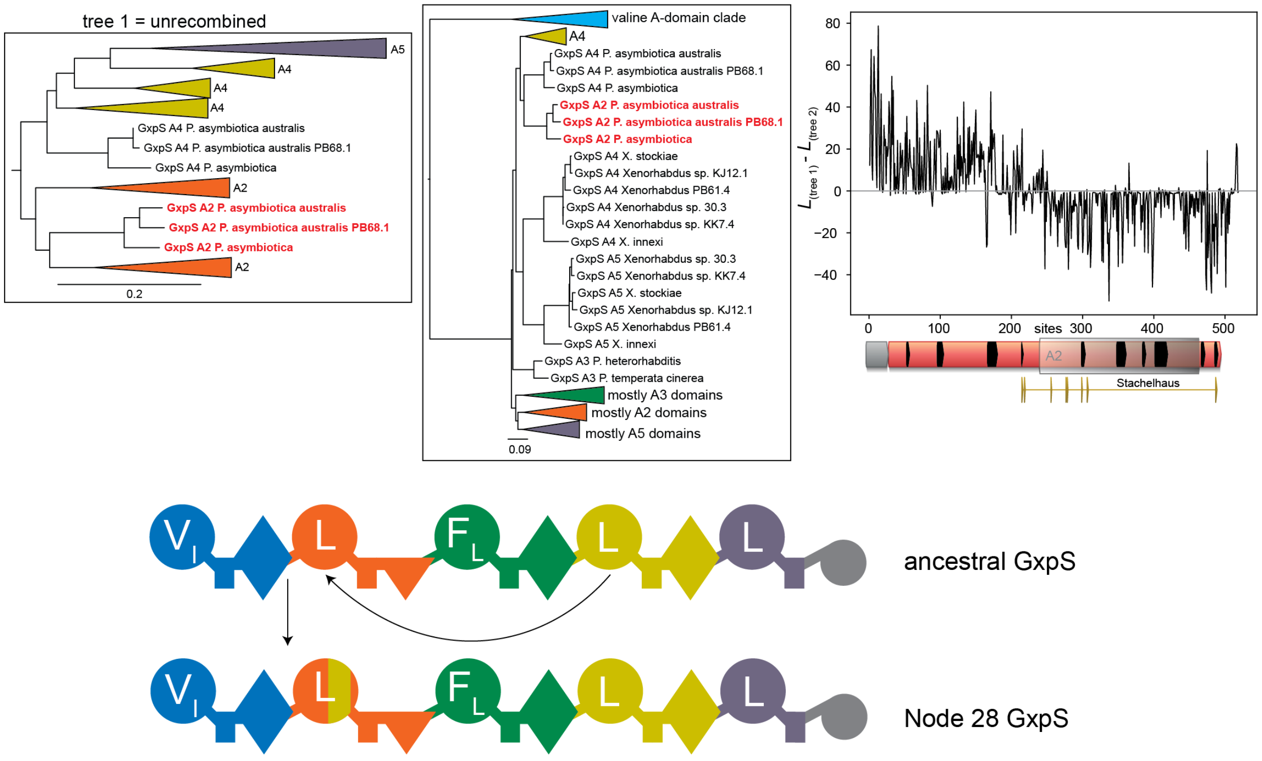


**Figure S105. Evolution of GxpS from Node 28.** This recombination was identified via PhyML_Multi. Left: first tree identified by PhyML_multi of an amino acid alignment of A-domains. A2 and A4 Sequences highlighted in red clade together with other sequences corresponding to their respective linear position in the sequence of GxpS. In this tree, A5 clades inside of A4, likely because our alignment contains A5 recombinations. Middle: second tree identified by PhyML_Multi, in which A2 domains highlighted in red are now sister to the A4 domains of the same synthetases (also in red), indicating that A2 was overwritten by A4 in the last common ancestor of these synthetases. Right: Site-wise log likelihood analysis of the two topologies. Sites with positive values are better described by tree 1, sites with negative values by tree 2. This analysis suggests a breakpoint somewhere in the A-domain, but we could not determine a clear boundary. RDP failed to recognize this relatively deep recombination event, presumably due to too much sequence divergence.

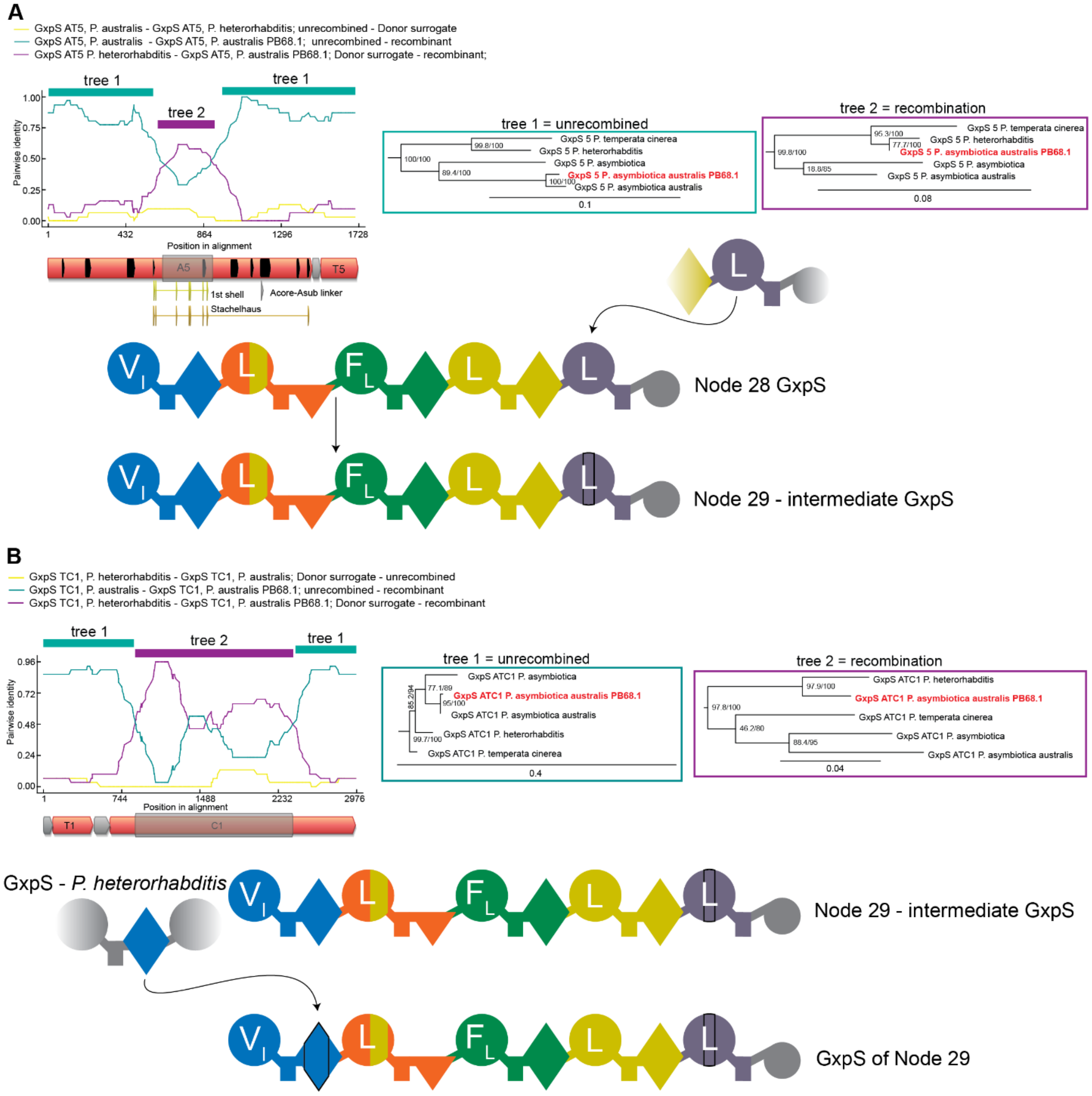


**Figure S106. Evolution of GxpS from Node 29.** These recombinations were identified using RDP. **(A)** RDP analysis and tree inference of a homotypic recombination in the A-domain. Tree 1 was inferred using amino acids, tree 2 using nucleotides. In tree 1 the sequence from *P. asymbiotica australis* PB68.1 (red) is sister to that of *P. australis*, consistent with the TTE-domain tree. In tree 2 it is instead sister to the sequence from *P. heterorhabditis*. These data imply a homotypic recombination between the A5 domains of two GxpS – from an ancestor *P. heterorhabditis* into the ancestor of *P. asymbiotica australis* PB68.1. (**B**) Another recombination between the same two synthetases, this time involving their first C-domains, with the direction of the recombination being the same. These two events may have happened simultaneously.

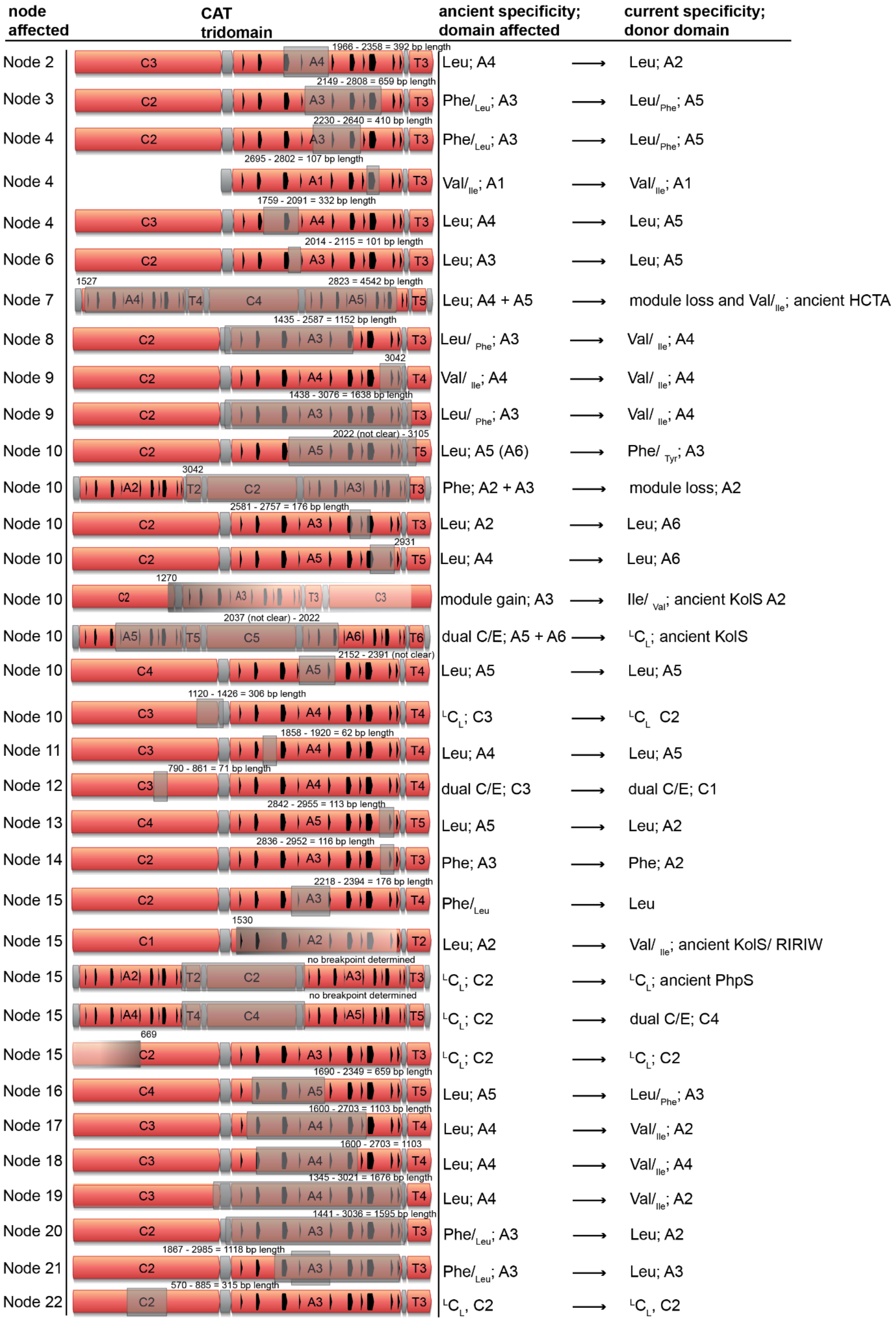
node affected
CAT tridomain
ancient specificity; domain affected
current specificity; donor domain
Node 2
1966 - 2358 = 392 bp length
Leu; A4 → Leu; A2
Node 3
2149 - 2808 = 659 bp length
Phe/Leu; A3 → Leu/Phe; A5
Node 4
2230 - 2640 = 410 bp length
Phe/Leu; A3 → Leu/Phe; A5
Node 4
2695 - 2802 = 107 bp length
Val/Ile; A1 → Val/Ile; A1
Node 4
1759 - 2091 = 332 bp length
Leu; A4 → Leu; A5
Node 6
2014 - 2115 = 101 bp length
Leu; A3 → Leu; A5
Node 7
1527
2823 = 4542 bp length
Leu; A4 + A5 → module loss and Val/Ile; ancient HCTA
Node 8
1435 - 2587 = 1152 bp length
Leu/Phe; A3 → Val/Ile; A4
Node 9
3042
Val/Ile; A4 → Val/Ile; A4
Node 9
1438 - 3076 = 1638 bp length
Leu/Phe; A3 → Val/Ile; A4
Node 10
2022 (not clear) - 3105
Leu; A5 (A6) → Phe/Tyr; A3
Node 10
3042
Phe; A2 + A3 → module loss; A2
Node 10
2581 - 2757 = 176 bp length
Leu; A2 → Leu; A6
Node 10
2931
Leu; A4 → Leu; A6
Node 10
1270
module gain; A3 → Ile/Val; ancient KolS A2
Node 10
2037 (not clear) - 2022
dual C/E; A5 + A6 → LCL; ancient KolS
Node 10
2152 - 2391 (not clear)
Leu; A5 → Leu; A5
Node 10
1120 - 1426 = 306 bp length
LCL; C3 → LCL C2
Node 11
1858 - 1920 = 62 bp length
Leu; A4 → Leu; A5
Node 12
790 - 861 = 71 bp length
dual C/E; C3 → dual C/E; C1
Node 13
2842 - 2955 = 113 bp length
Leu; A5 → Leu; A2
Node 14
2836 - 2952 = 116 bp length
Phe; A3 → Phe; A2
Node 15
2218 - 2394 = 176 bp length
Phe/Leu → Leu
Node 15
1530
Leu; A2 → Val/Ile; ancient KolS/ RIRIW
Node 15
no breakpoint determined
LCL; C2 → LCL; ancient PhpS
Node 15
no breakpoint determined
LCL; C2 → dual C/E; C4
Node 15
669
LCL; C2 → LCL; C2
Node 16
1690 - 2349 = 659 bp length
Leu; A5 → Leu/Phe; A3
Node 17
1600 - 2703 = 1103 bp length
Leu; A4 → Val/Ile; A2
Node 18
1600 - 2703 = 1103
Leu; A4 → Val/Ile; A4
Node 19
1345 - 3021 = 1676 bp length
Leu; A4 → Val/Ile; A2
Node 20
1441 - 3036 = 1595 bp length
Phe/Leu; A3 → Leu; A2
Node 21
1867 - 2985 = 1118 bp length
Phe/Leu; A3 → Leu; A3
Node 22
570 - 885 = 315 bp length
LCL, C2 → LCL, C2

**Figure S107. All recombinations from Node 1 to Node 22 are shown on a CAT-tridomain.** Recombination breakpoints are annotated on every CAT-tridomain and the corresponding length of recombination in bp. Annotated is also every ancient specificity, the domain that was affected by recombination, the donor domain and the current specificity.

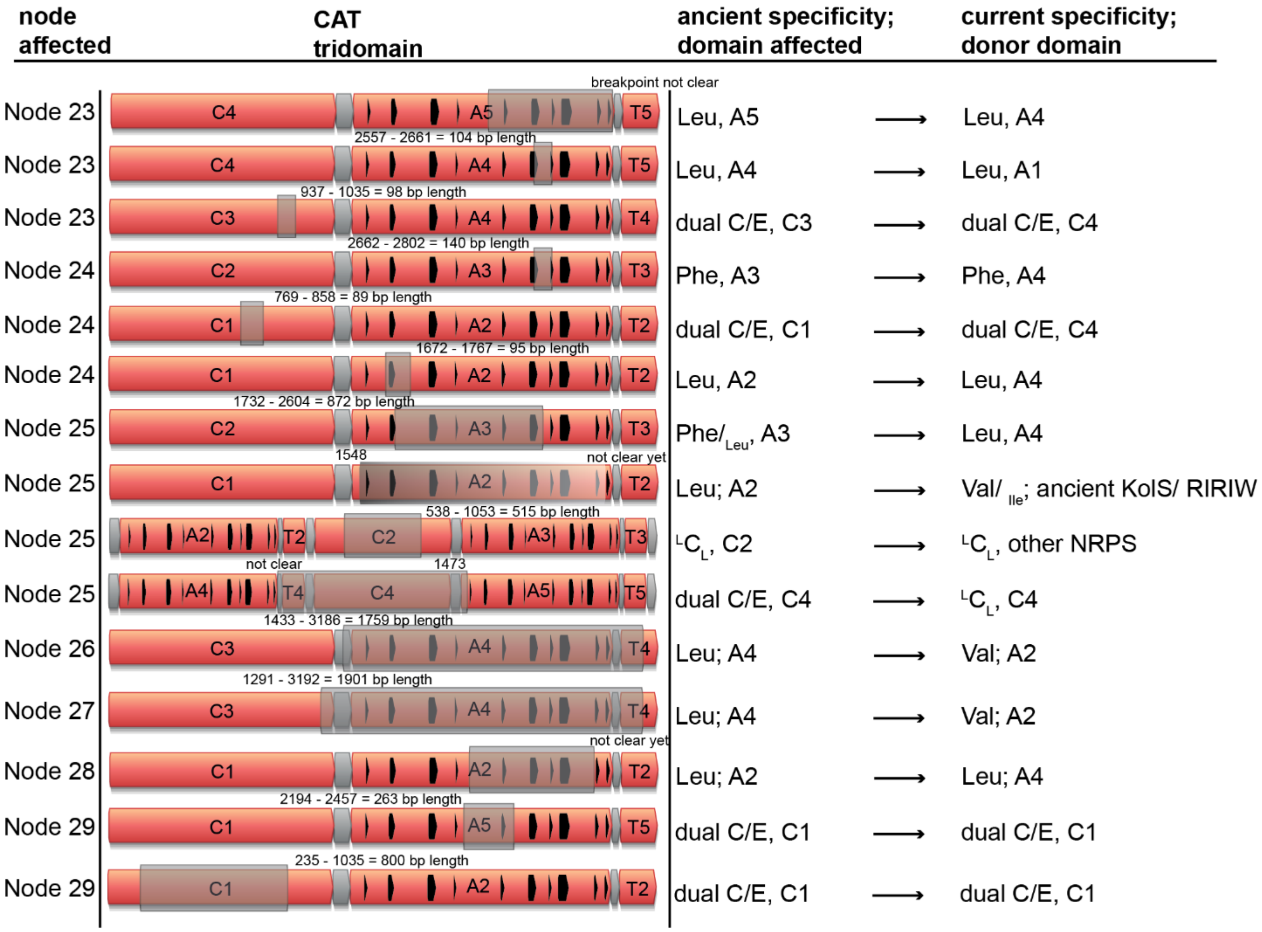


**Figure S108. All recombinations from Node 23 to Node 29 are shown on a CAT-tridomain.** Recombination breakpoints are annotated on every CAT-tridomain and the corresponding length of recombination in bp. Annotated is also every ancient specificity, thE-domain that was affected by recombination, the donor domain and the current specificity.

### References


1 Bode, H. B. *et al.* Determination of the absolute configuration of peptide natural products by using stable isotope labeling and mass spectrometry. *Chemistry* **18**, 2342-2348 (2012). https://doi.org:10.1002/chem.201103479

2 Kegler, C. *et al.* Rapid determination of the amino acid configuration of xenotetrapeptide. *Chembiochem* **15**, 826-828 (2014). https://doi.org:10.1002/cbic.201300602

3 Bian, X., Plaza, A., Yan, F., Zhang, Y. & Muller, R. Rational and efficient site-directed mutagenesis of adenylation domain alters relative yields of luminmide derivatives in vivo. *Biotechnol Bioeng* **112**, 1343-1353 (2015). https://doi.org:10.1002/bit.25560

4 Nollmann, F. I. *et al.* Insect-specific production of new GameXPeptides in photorhabdus luminescens TTO1, widespread natural products in entomopathogenic bacteria. *Chembiochem* **16**, 205-208 (2015). https://doi.org:10.1002/cbic.201402603

5 Schimming, O., Fleischhacker, F., Nollmann, F. I. & Bode, H. B. Yeast homologous recombination cloning leading to the novel peptides ambactin and xenolindicin. *Chembiochem* **15**, 1290-1294 (2014). https://doi.org:10.1002/cbic.201402065

6 Machado, R. A. R. *et al.* Whole-genome-based revisit of Photorhabdus phylogeny: proposal for the elevation of most Photorhabdus subspecies to the species level and description of one novel species Photorhabdus bodei sp. nov., and one novel subspecies Photorhabdus laumondii subsp. clarkei subsp. nov. *Int J Syst Evol Microbiol* **68**, 2664-2681 (2018). https://doi.org:10.1099/ijsem.0.002820

7 Tailliez, P., Pages, S., Ginibre, N. & Boemare, N. New insight into diversity in the genus Xenorhabdus, including the description of ten novel species. *Int J Syst Evol Microbiol* **56**, 2805-2818 (2006). https://doi.org:10.1099/ijs.0.64287-0

8 Goldfarb, T. *et al.* NCBI RefSeq: reference sequence standards through 25 years of curation and annotation. *Nucleic Acids Res* **53**, D243-D257 (2025). https://doi.org:10.1093/nar/gkae1038

9 Palma, L. *et al.* Genome Sequence Analysis of Native Xenorhabdus Strains Isolated from Entomopathogenic Nematodes in Argentina. *Toxins (Basel)* **16** (2024). https://doi.org:10.3390/toxins16020108

10 Lorenzen, W., Ahrendt, T., Bozhuyuk, K. A. & Bode, H. B. A multifunctional enzyme is involved in bacterial ether lipid biosynthesis. *Nat Chem Biol* **10**, 425-427 (2014). https://doi.org:10.1038/nchembio.1526

11 Bozhueyuek, K. A. J., Watzel, J., Abbood, N. & Bode, H. B. Synthetic Zippers as an Enabling Tool for Engineering of Non-Ribosomal Peptide Synthetases*. *Angew Chem Int Ed Engl* **60**, 17531-17538 (2021). https://doi.org:10.1002/anie.202102859

12 Kegler, C. & Bode, H. B. Artificial Splitting of a Non-Ribosomal Peptide Synthetase by Inserting Natural Docking Domains. *Angew Chem Int Ed Engl* **59**, 13463-13467 (2020). https://doi.org:10.1002/anie.201915989

13 Marahiel, M. A., Stachelhaus, T. & Mootz, H. D. Modular Peptide Synthetases Involved in Nonribosomal Peptide Synthesis. *Chem Rev* **97**, 2651-2674 (1997). https://doi.org:10.1021/cr960029e

14 Stachelhaus, T., Mootz, H. D. & Marahiel, M. A. The specificity-conferring code of adenylation domains in nonribosomal peptide synthetases. *Chem Biol* **6**, 493-505 (1999). https://doi.org:10.1016/S1074-5521(99)80082-9

15 Fuchs, S. W. *et al.* Neutral loss fragmentation pattern based screening for arginine-rich natural products in Xenorhabdus and Photorhabdus. *Anal Chem* **84**, 6948-6955 (2012). https://doi.org:10.1021/ac300372p
